# Chemoproteomics-guided development of SLC15A4 inhibitors with anti-inflammatory activity

**DOI:** 10.1101/2022.10.07.511216

**Authors:** Daniel C. Lazar, Wesley W. Wang, Tzu-Yuan Chiu, Weichao Li, Appaso M. Jadhav, Jacob M. Wozniak, Nathalia Gazaniga, Argyrios N. Theofilopoulos, John R. Teijaro, Christopher G. Parker

**Author notes:** Corresponding authors. (J.R.T), (C.G.P.). These authors contributed equally to this work (Daniel C. Lazar, Wesley W. Wang).

## Abstract

SLC15A4 is an endolysosome-resident transporter that is intimately linked with autoinflammation and autoimmunity. Specifically, SLC15A4 is critical for Toll-like receptor (TLR) 7, 8, and 9 as well as the nucleotide-binding oligomerization domain-containing protein (NOD) 2 signaling in several immune cell subsets. Notably, SLC15A4 is essential for the development of systemic lupus erythematosus in murine models and is associated with autoimmune conditions in humans. Despite its therapeutic potential, to our knowledge no pharmacological tools have been developed that target SLC15A4. Here, we use an integrated chemical proteomics approach to develop a suite of chemical tools, including first-in-class functional inhibitors, for SLC15A4. We demonstrate SLC15A4 inhibitors suppress endosomal TLR and NOD functions in a variety of human and mouse immune cells and provide early evidence of their ability to suppress inflammation *in vivo* and in clinical settings. Our findings establish SLC15A4 as a druggable target for the treatment of autoimmune/autoinflammatory conditions.

**One-Sentence Summary:** Discovery and characterization of SLC15A4 inhibitors with anti-inflammatory activity.

## Main Text

The endolysosomal solute carrier gene family 15 member 4 (SLC15A4) is a 12-transmembrane domain protein with high gene expression in antigen presenting cells (APCs), such as plasmacytoid dendritic cells (pDCs) and B cells (*1*). SLC15A4 is a proton-coupled transporter of amino acids, such as histidine, and di- or tri-peptides, including the NOD2 ligand muramyl dipeptide (MDP)(*2–4*). SLC15A4 has also been intimately linked to TLR7, 8 and 9 mediated signaling as well as the production of type-I interferons (IFN-I) and other proinflammatory cytokines (*1, 4–10*). Specifically, immune cells from Slc15a4 loss-of-function mutant (‘*feeble’*) and Slc15a4^-/-^ mice are defective in IFNα as well as TNFα, IL-6 and IL-12 production upon TLR 7-9 stimulation, but otherwise display normal development (*1*) (*4, 10*). Critically, Slc15a4 *feeble* and Slc15a4^-/-^ mice show striking reductions in systemic lupus erythematosus (SLE) manifestations (*9, 10*). Finally, genome-wide association studies (GWAS) have revealed SLC15A4 to be associated with inflammatory diseases such as SLE in human populations (*11–15*). Although these studies establish SLC15A4 as a critical modulator of inflammation and provide a strong therapeutic basis for the development of small molecule inhibitors, to our knowledge, none have yet been reported. Likely contributive to the dearth of chemical tools are general technical challenges associated with the SLC family (*16*), the lack of SLC15A4 structural information, and the absence of clear screenable functional readouts, stemming, in part, from an incomplete molecular understanding of how SLC15A4 mediates TLR-related signaling events.

Considering these challenges, here we opted to take an integrated approach to identify small molecules capable of binding to SLC15A4 directly in immune cells and interfere with SLC15A4-mediated endolysosomal TLR functions (**Fig S1A**). Towards this end, we previously developed a strategy that merges fragment-based ligand discovery (FBLD) with chemical proteomics to broadly map the ligandability of the proteome directly in cells, and demonstrated such ligands can be progressed to selective modulators of protein function (*17, 18*). Using this method, we screened an in-house library of fully-functionalized fragments (FFFs) in human peripheral blood mononuclear cells (PBMCs) to identify SLC15A4 small molecule binders via multiplexed proteomics, as previously described (*18*) (**Fig S1B-C, Table S1**). In parallel, to prioritize SLC15A4 ligands that might perturb SLC15A4-mediated TLR function, we co-screened FFFs for their ability to modulate IFNα production in TLR9-stimulated human pDCs. Among the nine FFFs that were found to substantially bind SLC15A4 from PBMCs, FFF-21 suppressed IFNα levels (**Fig. 1A-B**) to the highest extent, with an IC_50_ of approximately 21 µM while its tag-free analog, 21c, suppressed with a slightly lower IC_50_ of approximately 32 µM (**Fig S2A**). Next, we confirmed FFF-21 and 21c also block TLR 7/8 activation in monocyte-derived THP reporter cells (**Fig. S2B-C**). We further verified that FFF-21 engages SLC15A4 in CAL-1 cells, a pDC-like cell line (*19*), stably expressing lysosomal HA-SLC15A4 (**Fig S2D**) in a dose-dependent fashion which can be effectively blocked when co-treated with excess 21c (**Fig S2E**). Recent studies have shown that SLC15A4 mediates the transport of bacterially derived components, such as the NOD2 ligand MDP, to the cytosol and is required for NOD responses to endosomal trafficked ligands (*2, 3, 20, 21*). To assess whether FFF-21 inhibits NOD-mediated activation, we generated a SLC15A4-dependent NF-κB reporter cell line utilizing the previously reported SLC15A4 (L14A, L15A, L318A, V319A) mutant which localizes to the plasma membrane(*22*). Production of luciferase was dependent upon exposure to MDP and the presence of these mutations (**Fig S3A-B**). Treatment with FFF-21 blocked NF-κB activation in a concentration-dependent fashion, consistent with it blocking SLC15A4-mediated transport of NOD ligands (**Fig S3C**).

**Fig 1.**
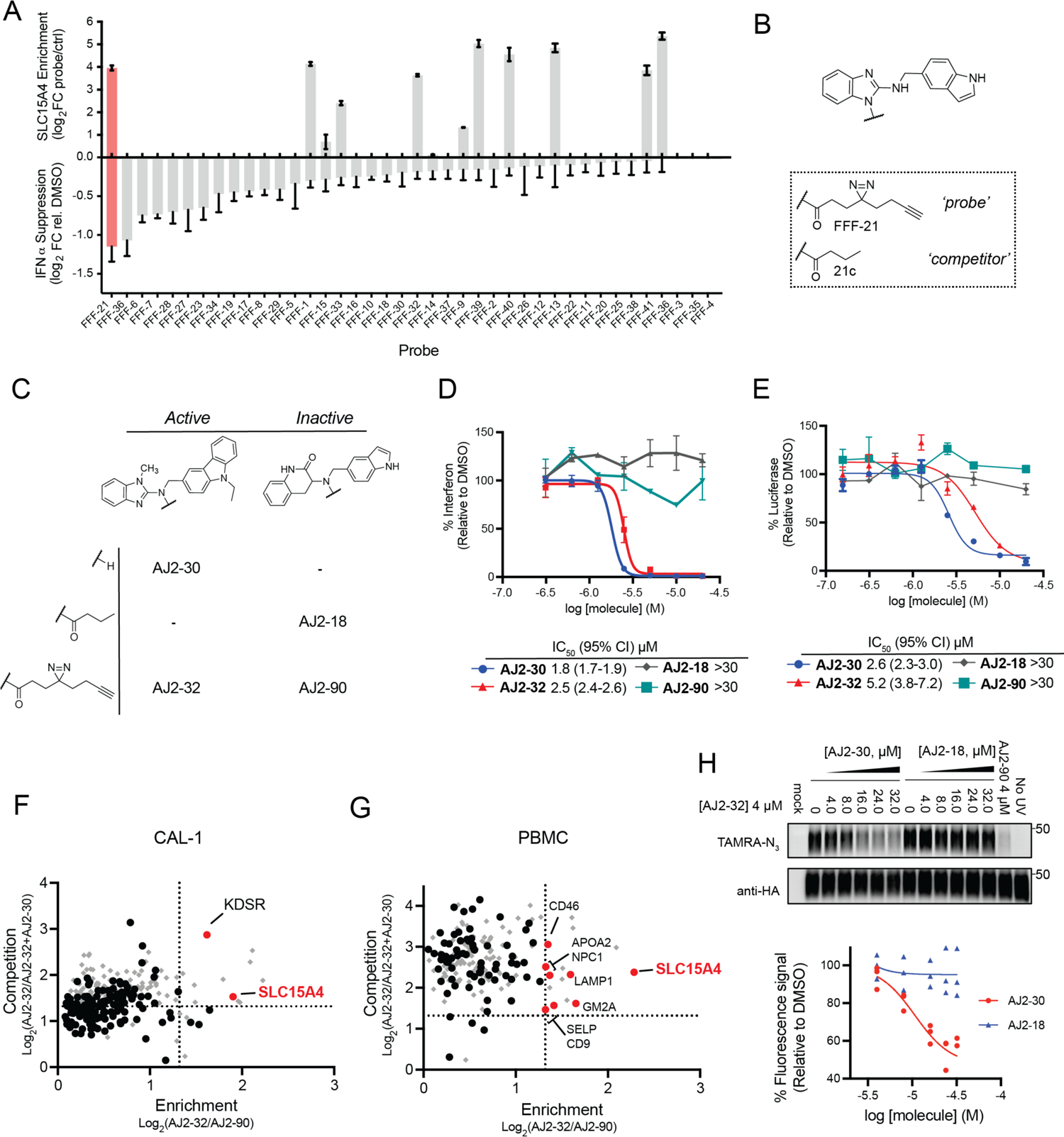
Chemoproteomic development of SLC15A4 inhibitors. (**A**) Integrated screen to identify small molecule SLC15A4 binders that suppress TLR9-mediated IFNα production in primary human pDCs (Fig S1A). Upper graph shows SLC15A4 enrichment by FFF probes (10 μM) over control probe (10 μM) in human PBMCs. Lower graph shows corresponding IFNα suppression in human pDCs after treatment with FFF probes (10 μM) and stimulation with CpG-A (1 μM, 24hrs). (**B**) Structures of hit FFF-21 and tag-free analog 21c (**C**) Structures of optimized analog AJ2-30, inactive analog AJ2-18, and corresponding fully functionalized photoaffinity probes AJ2-32 and AJ2-90, respectively. (**D**) IFNα suppression of CpG-A stimulated human pDCs by of AJ2-30, AJ2-32, AJ2-18 and AJ2-90. (**E**) Inhibition of MDP-mediated NOD2 signaling in A549 NF#x003BA;B reporter cells recombinantly expressing membrane-localized L14A/L15A/L318A/V319A SLC15A4. (**F** and **G**) Chemical proteomic target engagement of AJ2-30 in human CAL-1 cells (**F**) and human PBMCs (**G**). The x-axis shows protein enrichment by active probe AJ2-32 over inactive photoaffinity probe AJ2-90 (4 μM), while the y-axis shows protein competition in cells treated with active photoaffinity probe AJ2-32 (4 μM) and DMSO or the active competitor analog AJ2-30 (24 μM). Dotted lines indicate threshold for proteins to be designated as preferentially enriched by AJ2-32 (vertical) and competed (horizontal) by AJ2-30 (highlighted as red circles). Proteins plotted as gray diamonds are targets that were competed by inactive control competitor compound AJ2-18 (24 μM). Data is presented as the mean of biological replicated experiments (*n* = 2, p < 0.05). (**H**) Confirmation of AJ2-30 engagement of SLC15A4. UV-dependent labeling of HA-tagged SLC15A4 by AJ2-32 in stably expressed in CAL-1 cells is blocked by AJ2-30. Samples treated with PNGase to reduce glycosylation (see Methods). Plot shows quantitation of AJ2-32 probe labeling of SLC15A4 across biological replicates (*n* = 3) via measurement of fluorescence intensity from TAMRA signal.

Given the relatively low potency of FFF-21/21c and the limited structure-activity relationships (SAR) of compounds in our primary screen, we pursued the synthesis and evaluation of additional analogs. We synthesized 24 analogs of 21c, exploring modifications to the benzimidazole, indole and alkyl groups and assessed their relative abilities to suppress IFNα production in TLR9 - stimulated human pDCs and block SLC15A4-mediated MDP NF#x003BA;B activation (**Table S2**). We identified AJ2-30 to be among the most potent in both assays (**Fig 1C**), with an IFNα suppression IC_50_ of 1.8 µM and a MDP transport inhibition IC_50_ of 2.6 µM (**Fig 1D-E**). AJ2-30 displayed similar activity against mouse Slc15a4-mediated NOD activation (**Fig S4A**) and showed no activity against related family member SLC15A3 (**Fig S4B**), which has also been shown to mediate NOD2 ligand transport (*3*). Critically, the AJ2-30 scaffold allowed for derivitization with a photoaffinity tag to enable target engagment studies, with minimal effects on inhibitory potency (AJ2-32, **Fig 1D-E**). We also identified structurally related analogs that displayed no activity and therefore could serve as negative controls (AJ2-18 and corresponding photoaffinity probe AJ2-90; **Fig 1D-E**). We subsequently confirmed that AJ2-30, but not AJ2-18, blocks R848 induced activation of TLR7/8 signalling as well as TriDAP and MDP induced activation of NOD1 and NOD2 signalling, respectively, in pathway-specific reporter THP-1 cells (**Fig S5A-B**). To confirm engagement of SLC15A4 by these new analogs in cells, we profiled the protein targets of AJ2-32 in CAL-1 cells and human PBMCs by quantitative proteomics. From these studies, we found SLC15A4 to be a primary target of AJ2-32 and AJ2-30, as it is substantially enriched over inactive probe AJ2-90 and this binding could be blocked upon co-incubation with AJ2-30 but not inactive AJ2-18 (**Fig 1F-G, Table S3**). We further confirmed AJ2-32 labeling of SLC15A4 can be effectively blocked in a dose-dependent fashion with increasing concentrations of AJ2-30 (**Fig. 1H**), indicative of a stoichometric interaction. Cell free profiling of AJ2-30 at 10 µM across a panel of 468 kinases further supported the selectivity of the compound, with only three kinases (MAPKAPK2, PHKG2, and ERBB3) identified as minimally interacting partners (**Fig S6A**). Finally, these compounds showed no evidence of toxicity (**Fig S6B**) in human PBMCs and AJ2-30 possesssed suitable mouse pharmacokinetic properties for evaluation in short-term *in vivo* models of inflammation (**Fig S6C**).

We next evaluated the ability of AJ2-30 to suppress additional innate signaling pathways. AJ2-30, but not control analog AJ2-18, suppressed IFNα production following stimulation with Class A (CpG2216) and Class B (CpG2006) TLR9 agonists, the TLR7/8 stimulus R848, the TLR7-specific agonist R837 and TLR9 agonistic LL37:DNA complexes (*23*) in human pDCs (**Fig. 2A**). Additionally, when human pDCs were challenged with influenza virus, IFNα production was significantly inhibited by AJ2-30 but not AJ2-18 (**Fig. 2B**). Interestingly, additional pDC derived cytokines and chemokines, including IL6, TNF-α, G-CSF, CCL3, and CCL4, were also significantly reduced upon treatment of AJ2-30 after TLR9 stimulation, but not after TLR7/8 stimulation (**Fig. S7A** and **Fig. S7B**). Significant expression of SLC15A4 is also observed in monocytes, macrophages, and B cells, and genetic disruption of *Slc15a4* in these cell types interferes with their activation following endosomal TLR and NOD signaling (*2, 3, 8, 9*).

**Fig 2.**
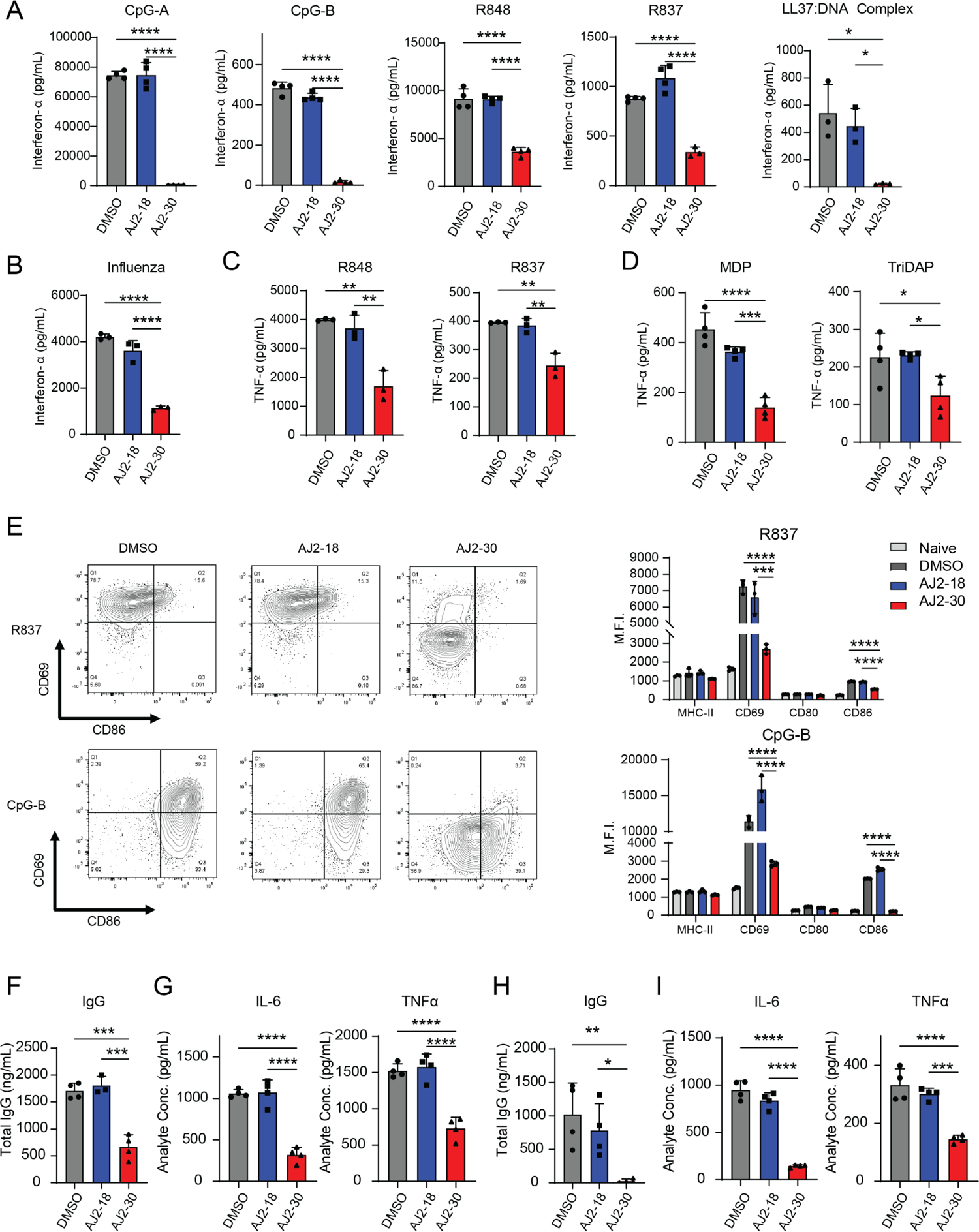
Pharmacological inhibition of SLC15A4 suppresses multiple innate signaling pathways. (**A**) AJ2-30 inhibits TLR7, TLR7/8, and TLR9-induced pDC IFN-α production. pDCs were isolated from human PBMCs and treated with AJ2-18 or AJ2-30 (5 μM, 24 hrs) along with CpG-A (1 μM), CpG-B (1 μM), R848 (5 μg/mL), R837 (5 μg/mL), or LL37:DNA complex (10 μg/mL). (**B**) AJ2-30 inhibits influenza mediated IFN-α production in human pDCs. pDCs were isolated from human PBMCs and AJ2-18 or AJ2-30 (5 μM, 24hr) along with influenza virus (TLR7 agonist; MOI = 1). (**C**) AJ2-30 inhibits TLR7 and TLR7/8 mediated production of TNF-α in primary human monocytes. (**D**) AJ2-30 inhibits NOD1 and NOD2 signaling mediated by the bacterial dipeptides MDP and TriDAP. Primary human derived macrophages were treated with AJ2-18 or AJ2-30 (5 μM), polarized with IFNγ, and then challenged with the bacterial dipeptides MDP (150 ng/ml) or TriDAP (2000 ng/mL). (**E**) Primary human B cells isolated from PBMCs were treated with AJ2-18 or AJ2-30 (5 μM, 24hr) while stimulated with either R837 (μg/mL) or CpG-B (1 μM). B cell activation was assessed by measuring the surface expression of CD69, CD80, CD86, and MHC-II on live cells. (**F** and **H**) *In vitro* IgG secretion from primary human B cells stimulated with CpG-B (**F**) or R837 (**H**) for 6 days in the presence of either AJ2-18 and AJ2-30 (5 μM). (**G** and **I**) *In vitro* secretion of IL-6 and TNF-α from isolated primary human B cells in the presence of either AJ2-18 or AJ2-30 (5 μM) when stimulated by either (**G**) CpG-B or (**H**) R837 after 24 hrs. Results are representative of at least 3 independent experiments and values indicate mean ± SD. Statistical analysis was performed using ANOVA analysis followed by multiple comparisons test * p ≤ 0.05; ** p ≤ 0.01; *** p ≤ 0.001; **** p ≤ 0.0001.

We found AJ2-30, but not AJ2-18 inhibited TLR7/8-induced production of TNFα in human monocytes (**Fig. 2C**). However, AJ2-30 does not suppress production of TNFα with other TLR stimulation, including TLR2 agonist Pam3CSK4 and TLR4 agonist LPS (**Fig. S7C)**, or IFN-β production upon STING stimulation (**Fig S7D**), together suggesting AJ2-30 is not broadly immunosuppressive. Consistent with our observations in NOD reporter assays described above, treatment with AJ2-30 also suppressed MDP and TriDAP-mediated TNFα production in human macrophages (**Fig. 2D**). However, as was observed in *Slc15a4^-/-^* derived macrophages (*9*), we observe no suppressive effects of AJ2-30 in human macrophages upon TLR7/8 stimulation (**Fig S7E**). SLC15A4 inhibition also suppresses B cell function. Flow cytometry analysis of cell surface activation markers showed that AJ2-30 also suppresses B cell activation following stimulation with either CpG or R837 (**Fig. 2E**). In addition, AJ2-30 but not AJ2-18 reduced total IgG (**Fig 2F-H**), IL-6, TNFα and IL-10 production (**Fig 2G-I and Fig S7F-G**) upon TLR9 (**Fig 2F-G**) or TLR7/8 (**Fig 2 H-I**) stimulation. Finally, AJ2-30 also displays cross-species reactivity, suppressing IFNα production in mouse pDCs (**Fig. S8A**) as well as mouse B cell IL-6 production (**Fig S8B**) with TLR7-9 stimulation but not with the TLR4 agonist LPS. Consistent with B cells derived from *Slc15a4*^-/-^ mice (*8*), we also observe AJ2-30-mediated suppression of IgG, but not IgM production, of TLR7-9 stimulated WT B cells (**Fig S8C-D**).

Previously it has been shown that B cells from *Slc15a4*^-/-^ mice display reduced CD86 upregulation following R837 stimulation compared to wild-type B cells (*8*). Likewise, we observed reduced levels of CD86 induction in *feeble* B cells compared to WT controls, however AJ2-30 treatment did not further suppress activation following TLR7/8 or 9 stimulation, nor did it further affect IL-6 production in these cells (**Fig S8E-F**). Similarly, bone marrow derived macrophages (BMDMs) derived from *feeble* mice exhibited reduced IL-6 production following MDP stimulation and AJ2-30 did not have additional suppressive effects (**Fig S8G-H**). Further treatment of GM-CSF derived cDCs, which in contrast to pDCs, have significantly lower expression of Slc15a4 (*1*), with AJ2-30 produce equivalent amounts of TNFα in response to various TLR agonists (**Fig S8I**). Taken together, these data demonstrate that AJ2-30 inhibits SLC15A4-dependent TLR 7-9 as well as NOD functions in both human and mouse derived pDCs, B cells, and macrophages.

We next sought to characterize how SLC15A4 inhibition with AJ2-30 disrupts endolysosomal TLR signaling. SLC15A4 deletion was previously shown to increase lysosomal pH in B cells, potentially through the disruption v-ATPase activity (*8*). Interestingly, we observe a small increase in lysosomal acidity when B cells were treated with AJ2-30 (**Fig. S9**), likely the result of inhibition of SLC15A4-mediated proton export from the lysosome. Despite this pH change, AJ2-30 did not disrupt normal IRAK1, IKKα/β, and I#x003BA;Bα activation upon stimulation with R848, suggesting that SLC15A4 inhibition does not disrupt proximal TLR7/8 signaling (**Fig S10A**), consistent with observations in Slc15a4^-/-^ B cells (*9*). In contrast, we observed that AJ2-30 disrupted proximal TLR9 signaling upon stimulation with CpG (**Fig S10B**). Amplification of IFNα is primarily driven by IRF7, while IRF5 generally induces expression of other proinflammatory cytokines such as TNFα, IL-6 and IL-12 (*24–26*). SLC15A4 loss has been shown to disrupt the mTOR signaling pathway (*7, 8, 27*), a critical component of the IRF7-IFNα regulatory circuit (*28*). To further characterize AJ2-30 mediated SLC15A4 inhibition, we next examined whether it interferes with mTOR signaling in immune cells. Indeed, we observe a strong, dose-dependent impairment of mTOR pathway activation in human B cells stimulated with either TLR7/8 or TLR9 agonists by AJ2-30, but not with control AJ2-18 (**Fig 3A**). Similar inhibitory effects were also observed in human pDCs (**Fig S11A-B**). However, AJ2-30 does not block mTOR pathway activation upon stimulation with the TLR2 agonist Pam3CSK4, suggesting it does not directly target mTOR itself and inhibition is restricted to endolysosomal TLR7-9-mediated mTOR activation (**Fig 3A**). mTOR is critical for TLR-mediated expression of type I IFNs in various immune cells, and disruption of mTOR signaling impairs IRF5 activation and the phosphorylation, translation, and nuclear translocation of IRF7 (*28, 29*). When pDCs are stimulated with either CpG or R848, we observe a strong blockade of IRF7 phosphorylation (**Fig S11C-D**) as well as IRF5 and IRF7 nuclear translocation with AJ2-30, but not with inactive control AJ2-18 (**Fig 3B-C**). Furthermore, we observe that AJ2-30, but not AJ2-18, reduced both TLR7/8 and TLR9-induced IRF5 and IRF7 protein expression in human B cells (**Fig 3D**).

**Fig 3.**
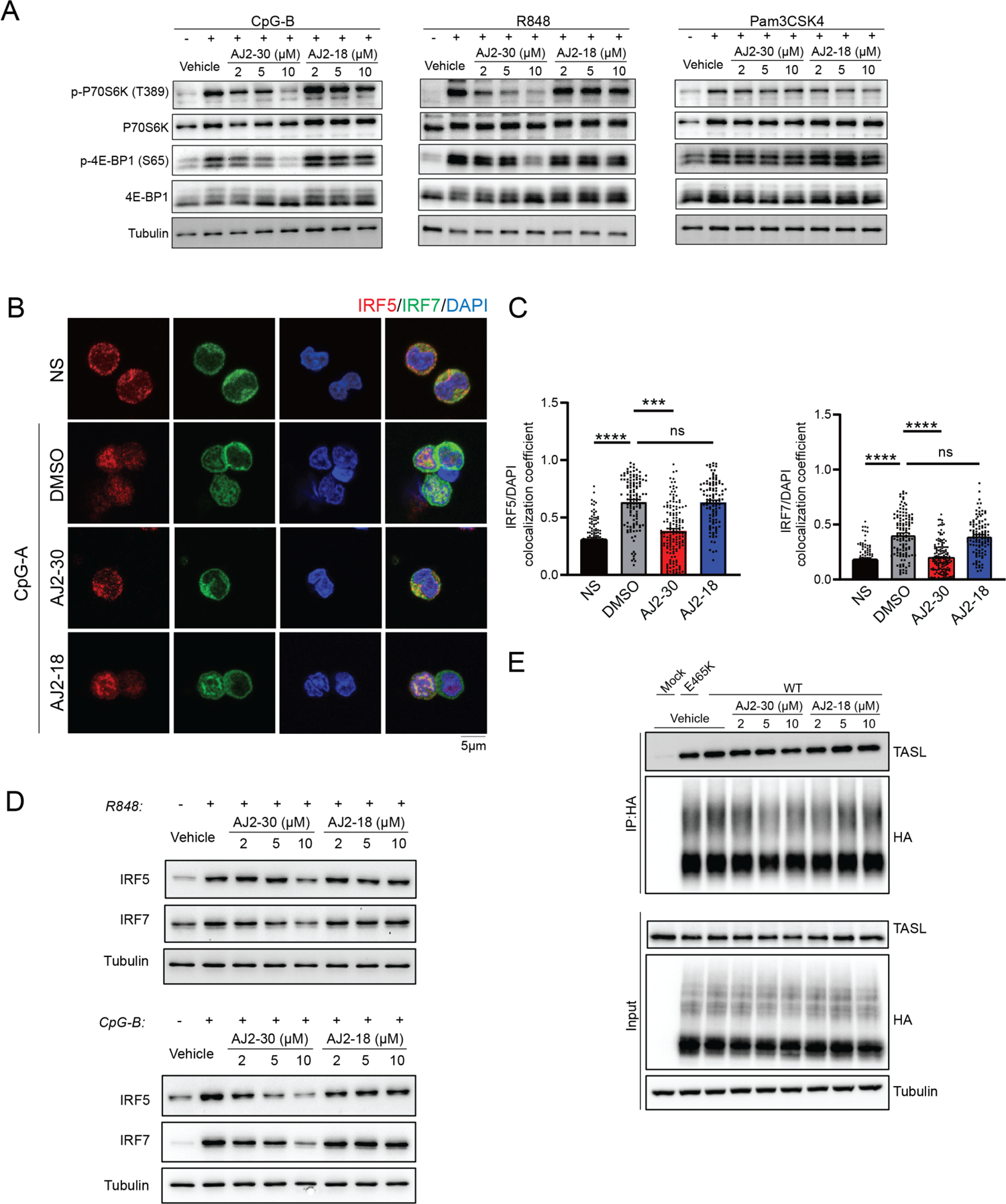
TLR7-9 mediated signaling changes impaired by SLC15A4 inhibition. (**A**) Immunoblot analysis of mTOR substrates in B cells isolated from human PBMCs. Cells were co-treated with compounds/DMSO and stimuli (1 μM CpG-B, 5 μg/ml R848, 1 μg/ml Pam3CSK4). Data are representative of two independent experiments. (**B**) SLC15A4 inhibition blocks IRF5 and IRF7 nuclear translocation in human pDCs as measured by immunofluorescence microscopy. Human pDCs were co-treated with compounds (5 μM) or DMSO and stimuli for 4 hrs before fixation. (**C**) Quantitation of the colocalization coefficient of IRF5 or IRF7 with DAPI (*n* = 30-50 cells; mean ± SD). Images are representative of three independent experiments. Statistical analysis was performed using ANOVA analysis followed by multiple comparisons test. ***p <0.001, **** p ≤ 0.0001 (**D**) AJ2-30 suppresses TLR7/8 or TLR9-induced IRF5 and IRF7 translation in human B cells. Cells were co-treated with compounds (5 μM) or DMSO and stimuli (1 μM CpG-B or 5 μg/ml R848) for 8 hrs. Data are representative of two independent experiments. (**E**) SLC15A4/TASL interaction analysis by HA-immunoprecipitation in CAL-1 cells stably overexpressing EGFP, HA-SLC15A4, and transport-deficient mutant HA-SLC15A4 (E465K) (*22, 33*). Cells were treated with escalating doses of compounds or DMSO for 1 hour before lysis and immunoprecipitation using anti-HA magnetic beads. After washing, proteins were eluted and analyzed by SDS-PAGE-western blot. Data are representative of three independent experiments.

Together, these data suggest that pharmacological inhibition of SLC15A4 with AJ2-30 impairs endolysosomal TLR7-9-mediated mTOR activation, IRF5 and 7 activation, and subsequent cytokine production. Recently, it has also been shown that SLC15A4 interacts with the adaptor protein TASL which serves as a scaffold for TLR7-9 mediated IRF5 activation (*6*). Considering this, we also examined whether AJ2-30 perturbs this interaction. Using CAL-1 cells expressing HA-tagged SLC15A4, we confirmed its interaction with endogenous TASL, but we detected no disruption by AJ2-30 (**Fig 3E**). Similarly, using tagged recombinant TASL as bait, again no disruption with SLC15A4 was observed (**Fig S12**).

Finally, we sought to assess the therapeutic potential of small molecule SLC15A4 inhibition. Towards this end, we first investigated immunosuppressive effects of AJ2-30 in PBMCs isolated from a cohort of lupus patients both with and without a therapeutic regime for treatment of their disease (**Table S4**). As expected, stimulation of PBMCs from these patients with TLR7/8 or TLR9 agonists (**Fig 4A-B**) produced significant levels of IFN-α and other inflammatory cytokines, which were substantially blunted by AJ2-30 treatment (**Fig 4A-B**).

**Fig 4.**
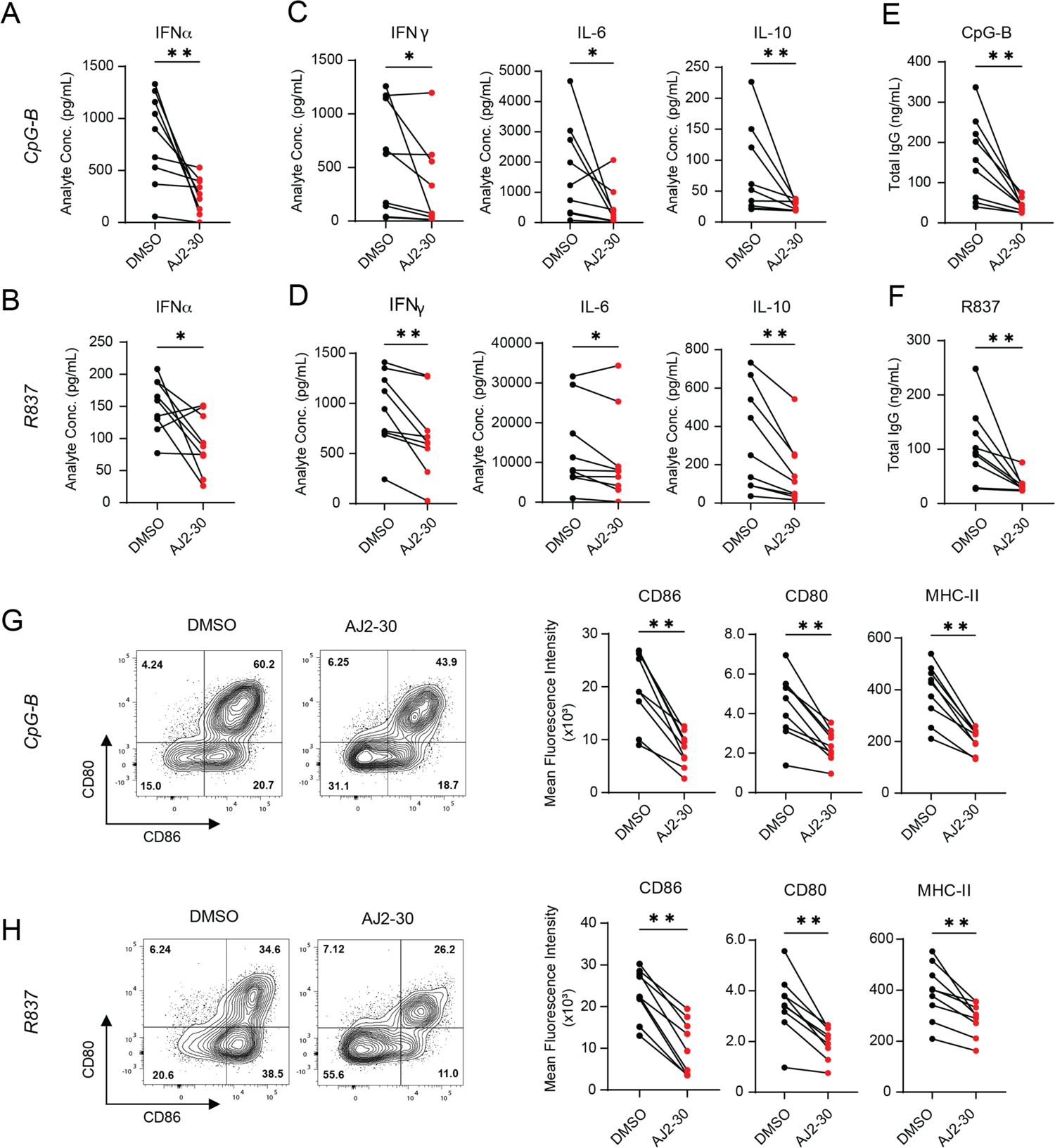
Pharmacological inhibition of SLC15A4 suppresses inflammatory cytokine production in lupus patient derived PBMCs. (**A** and **B**) Secretion of IFN-α and other inflammatory cytokines from PBMCs of lupus patients in the presence of either DMSO or AJ2-30 (5 μM) when stimulated by either CpG-A (**A**, 1 μM) or R837 (**B**, 10 μg/mL) after 24 hrs. (**C** and **D**) Total IgG from PBMCs stimulated with R837 or CpG-B in the presence of AJ2-30 (5 μM) or DMSO was measured by ELISA after 6 days of stimulation. (**E** and **F**) Expression of the costimulatory molecules CD80, CD86, and MHC-II on B cells following R837 or CpG-B stimulation when treated with either DMSO or AJ2-30 (5 μM) for 24hrs. Significance was determined using a Wilcoxon matched-pairs signed rank test. * p ≤ 0.05; ** p ≤ 0.01.

Additionally, AJ2-30 significantly reduced IgG antibody levels (**Fig 4C-D**) as well as the expression of the costimulatory molecules CD80, CD86, and MHC-II on B cells following TLR7/8 or TLR9 stimulation (**Fig 4E-F**). Elevated IFN-I signatures, cytokine production and cellular activation is observed in lupus patients (*30–32*). Treatment of unstimulated lupus patient-derived PBMCs with AJ2-30 significantly suppressed production of IFNγ, IL6, and IL10 (**Fig S13A**). Moreover, the elevated levels of B cell co-stimulatory molecules and IgG were reduced following treatment with AJ2-30 (**Fig S13B-D**), demonstrating that the heightened activation state of lupus PBMCs can be suppressed by AJ2-30. Finally, we sought to obtain preliminary *in vivo* evidence that SLC15A4 inhibition results in disruption of type 1 and 2 IFN production.

Towards this end, IFN production was stimulated with DOTAP-complexed CpG-A, as previously described (*9*), and serum cytokine levels were measured following treatment with AJ2-30. Consistent with findings in Slc15a4-deficient mice (*1, 9, 10*) and our cellular studies, AJ2-30 significantly reduced IFNα, IFNγ, IL-6 and IL-10 levels (**Fig S14**), supporting that AJ2-30 can mimic SLC15A4 functional loss *in vivo*.

Here, we report the development of, to our knowledge, the first SLC15A4 inhibitors. We demonstrate that our lead compound, AJ2-30, engages SLC15A4 directly in primary immune cells, inhibits TLR7-9 signaling pathways and cytokine production as well as SLC15A4-mediated NOD 1/2 activation in primary human and mouse immune cells, key functions regulated by SLC15A4 (*1, 4, 5, 8-10*). Our findings suggest that pharmacological inhibition of SLC15A4 results in disruption of an mTOR-IRF circuit essential for IFN-I and other inflammatory cytokine production, however the molecular details of how SLC15A4 modulates this pathway are not yet elucidated. AJ2-30 also substantially blunts inflammatory cytokine production in lupus patient derived immune cells and *in vivo*, underscoring the therapeutic potential of SLC15A4 inhibitors. More broadly, our study highlights the general utility of the integrated chemical proteomic strategy described herein to identify useful chemical probes for therapeutically compelling but technically challenging targets.

## Acknowledgments

We thank Belharra Therapeutics for their kind assistance with *in vivo* and kinome profiling studies. We also thank B.F. Cravatt for his invaluable discussions and insights.

## Author contributions

Conceptualization: CGP, JRT, DCL, WWW

Methodology: DCL, WWW, WL, TC

Investigation: DCL, WWW, WL, TC, AMJ, JMW, NG

Funding acquisition: CGP, JRT

Project administration: CGP, JRT

Supervision: CGP, JRT, ANT

Writing – original draft: CGP, JRT

Writing – review & editing: CGP, JRT, DCL, WWW, WL, TC, AMJ, JMW, ANT

## Data and materials availability

All data are available in the manuscript or in the supplementary materials. Noncommercial reagents described in this manuscript are available from CGP or JRT under a material transfer agreement with The Scripps Research Institute. All proteomics data will be deposited to the ProteomeXchange Consortium server upon acceptance of publication

## Supplementary Materials

### MATERIALS AND METHODS

#### Purchased reagents

| REAGENT or RESOURCE | SOURCE | IDENTIFIER |
| --- | --- | --- |
| **Antibodies** |  |  |
| Anti-CXorf21(TASL) antibody (1:1000) | Millipore-Sigma | #HPA001185 |
| HA-Tag mAb (1:2000) | Cell Signaling Technology | #3724 |
| p70 S6 Kinase mAb (1:1000) | Cell Signaling Technology | #34475 |
| P-p70 S6 Kinase (T389) mAb (1:500) | Cell Signaling Technology | #9234 |
| IRF-5 mAb (IB 1:500, IF 1:200) | Cell Signaling Technology | #96527 |
| IRF-7 mAb (IF 1:200) | Santa Cruz Biotechnology | sc-74472 |
| IRF-7 mAb (IB:1:1000) | Abcam | Ab238137 |
| 4E-BP1 mAb (1:1000) | Cell Signaling Technology | #9644 |
| P-4E-BP1 (S65) mAb (1:1000) | Cell Signaling Technology | #13443 |
| IκBα (1:500) | Cell Signaling Technology | #4814 |
| p-IκBα (S32) mAb (1:500) | Cell Signaling Technology | #2859 |
| p- IKKα/$\beta$ (Ser176/180) (1:500) | Cell Signaling Technology | #2697 |
| IKKβ mAb (1:1000) | Cell Signaling Technology | #8943 |
| IRAK1 (1:300) | Cell Signaling Technology | #4359 |
| Anti-Flag mAb (1:1000) | Cell Signaling Technology | #14793 |
| Anti-Tubulin hFAB(1:3000) | Bio-Rad | #12004165 |
| Donkey anti-Mouse IgG, Alexa Fluor Plus 488 | Thermo Fisher Scientific | A32766TR |
| Donkey anti-Rabbit IgG, Alexa Fluor Plus 647 | Thermo Fisher Scientific | A32795TR |
| Rabbit anti-Mouse IgG, HRP | Thermo Fisher Scientific | PA1-28568 |
| Goat anti-Rabbit IgG, HRP | Thermo Fisher Scientific | 31460 |
| NucBlue™ Fixed Cell ReadyProbes™ Reagent (DAPI) | Thermo Fisher Scientific | R37606 |
| LysoSensor Green DND-189 | Thermo | L7535 |
| eBioscience Fixable Viability Dye eFluor™ 780 | Thermo | 65-0865-18 |
| BV421 anti-human CD80 (Clone 2D10) | Biolegend | 305222 |
| AlexaFluor488 anti-human HLA-DR (Clone LN3) | Biolegend | 327010 |
| PE anti-human CD69 (Clone FN50) | Biolegend | 310906 |
| APC anti-human CD86 (Clone IT2.2) | Biolegend | 305412 |
| PE/Cy7 anti-human CD19 (Clone HIB19) | Biolegend | 302216 |
| BV421™ anti-mouse/human CD45R/B220 Antibody (Clone RA3-6B2) | Biolegend | 103240 |
| PE anti-mouse CD69 (Clone H1.2F3) | Biolegend | 104508 |
| APC anti-mouse CD86 (Clone GL-1) | Biolegend | 105012 |
| PerCP/Cy5.5 anti-mouse CD80 (Clone 16-10A1) | Biolegend | 104722 |
| PerCP/Cy5.5 anti-human CD123 (Clone 6H6) | Biolegend | 306016 |
| PE anti-human CD303 (BDCA-2) (Clone 201A) | Biolegend | 354204 |
| IRF-7 pS477/pS479 Antibody, anti-human, REAfinity™ | Miltenyi Biotec | 130-104-702 |
| Anti-hu/mo phospho-mTOR (Ser2448)-PE | Thermo | 12-9718-42 |
| phospho-S6ribosomal Protein (S235/236) XP-PeCy7 | Cell Signaling Technology | 34411S |
| phospho-4E-BP1 (T37/46) AlexaFluor647 | Cell Signaling Technology | 5123S |
| **Cell Lines** |  |  |
| 293T | ATCC | CRL-3216 |
| A549 | ATCC | CCL-185 |
| CAL-1 | S. Kamihira, (Nagasaki University ) | PMID: 15765784 |
| CAL-1::EGFP | In this study | CAL-1 stably overexpressing EGFP |
| CAL-1::HA-SLC15A4 | In this study | CAL-1 stably overexpressing HA tagged SLC15A4 |
| CAL-1::HA-SLC15A4<sup>E465K</sup> | In this study | CAL-1 stably overexpressing HA tagged SLC15A4<sup>E465K</sup> |
| THP-1 Dual | InvivoGen | thpd-nfis |
| **Critical commercial assays** |  |  |
| Quanti-Luc™-Gold | InvivoGen | rep-qlc1 |
| Quanti-Blue™ | InvivoGen | rep-qbs |
| Trans-Blot Turbo RTA transfer Kit, LF, PVDF | BioRad | #1704275 |
| CellTiter-Glo Luminescent Cell Viability Assay | Promega | G7570 |
| CytoTox 96® Non-Radioactive Cytotoxicity Assay | Promega | G1780 |
| Human IFN-Beta ELISA Kit | pbl Assay Science | 41410-2 |
| Human IFN-Alpha ELISA Kit | pbl Assay Science | 41100-2 |
| Mouse IFN-Alpha ELISA Kit | pbl Assay Science | 42120-2 |
| ELISA MAX™ Deluxe Set Human TNF-α | Biolegend | 430204 |
| ELISA MAX™ Deluxe Set Mouse IL-6 | Biolegend | 431304 |
| ELISA MAX<sup>TM</sup> Deluxe Set<br>Mouse TNF-α | Biolegend | 430904 |
| IgG (Total) Human Uncoated<br>ELISA Kit with Plates | Thermo | 88-50550-22 |
| IgG2c Mouse Uncoated ELISA<br>Kit with Plates | Thermo | 88-50670-22 |
| IgM Mouse Uncoated ELISA<br>Kit with Plates | Thermo | 88-50470-22 |
| Bio-Plex Pro Mouse Cytokine<br>23-plex assay | Bio-Rad | M60009RDPD |
| Bio-Plex Pro Human Cytokine<br>27-plex Assay | Bio-Rad | M500KCAF0Y |
| **Reagents, Peptides and Recombinant proteins** |  |  |
| HA peptide | GenScript | RP11735 |
| Rhodamine-azide | Synthesized in lab | - |
| Triptolide | AdipoGen Life Sciences | 501146334 |
| GSK538, RIPK2i | MedChem | 1346547-00-9 |
| NOD2 Signaling Inhibitor II,<br>GSK717 | 5.33718.0001 | EMD Millipore |
| R848 (Resiquimod) | InvivoGen | tlrl-848 |
| Imiquimod (R837) | InvivoGen | tlrl-imq |
| VACV-70/ LyoVec<sup>TM</sup> | InvivoGen | tlrl-vav70c |
| Pam3CSK4 | InvivoGen | tlrl-pms |
| Standard LPS, E. coli K12 | InvivoGen | tlrl-eklps |
| ODN 2216 | InvivoGen | tlrl-2216-5 |
| ODN 2006 (ODN 7909) | InvivoGen | tlrl-2006-5 |
| LL-37 (Antimicrobial peptide) | InvivoGen | tlrl-l37 |
| Muramyldipeptide (L-D<br>isoform, active) | InvivoGen | tlrl-mdp |
| Tri-DAP | InvivoGen | tlrl-tdap |
| DNeasy Blood & Tissue Kit | Qiagen | 69504 |
| QIAprep Spin Miniprep Kit<br>(250) | Qiagen | 27106 |
| EndoFree Plasmid Maxi Kit | Qiagen | 12362 |
| Pierce Anti-HA Magnetic Beads | Thermo Scientific | 88837 |
| Sequencing Grade Modified<br>Trypsin | Promega | V5111 |
| DOTAP Liposomal Transfection<br>Reagent | Roche | 11202375001 |
| CD304 (BDCA-4/Neuropilin-1)<br>MicroBead Kit, human | Miltenyi Biotec | 130-090-532 |
| EasySep<sup>TM</sup> Human B Cell<br>Isolation Kit | StemCell Technologies | 17954 |
| EasySep<sup>TM</sup> Mouse B Cell<br>Isolation Kit | StemCell Technologies | 19854 |
| EasySep<sup>TM</sup> Human Monocyte<br>Isolation Kit | StemCell Technologies | 19359 |
| ACK lysing buffer (RBC Lysis) | Thermo | 118-156-101 |
| Recombinant Human IFN-gamma | R&D Systems | Catalog 285-IF |
| Human Recombinant M-CSF | StemCell Technologies | 78057.1 |
| FLt3L-Ig BioXcell | BioXcell | BE0098 |
| Mouse M-CSF | Fisher | 416ML010 |
| GM-CSF | Fisher | 41-5ML0-05CF |
| **Recombinant DNA** |  |  |
| pDONR221 SLC15A4 | Addgene | #131910 |
| pLJM1-EGFP | Addgene | #19319 |
| Cxorf21 (TASL) | GPS core UF Scripps | 476835 |
| Human SLC15A4 (Codon Optimized) | IDT |  |
| Human Slc15A3 (Codon Optimized) | IDT |  |
| Mouse SLC15A4 (Codon Optimized) | IDT |  |
| Human NOD2 (Codon Optomized) | IDT |  |
| **Software and Algorithms** |  |  |
| Proteome Discoverer | Thermo Scientific | Opton |
| GraphPad | Dotmatics | - |
| FlowJo™ v10 | FlowJo | - |

#### Cell line culture

HEK293T, and A549 cells were purchased from ATCC. THP1-Dual^TM^ Cells were purchased from InvivoGen. CAL-1 cells were obtained from S. Kamihira, Nagasaki University (PMID: 15765784).(*1*) HEK293T cells were maintained in DMEM media supplemented with 10% (v/v) fetal bovine serum (FBS), penicillin (100 U/mL), streptomycin (100 μg/mL) and L-glutamine (2 mg/mL). A549 cells were maintained in DMEM supplemented with 2 mM L-glutamine, 25 mM HEPES, 10% (v/v) heat-inactivated FBS, 1X non-essential amino acids, and 55 μM β-Mercaptoethanol. CAL-1 cells were maintained in RPMI1640 media supplemented with 10% (v/v) FBS, penicillin (100 U/mL), streptomycin (100 μg/mL) and L-glutamine (2 mg/mL). THP1-Dual^TM^ Cells were cultured according to manufacturer’s instructions. All cell lines were grown at 37℃ in a humidified 5% CO2 atmosphere.

#### Plasmids and cDNA fragments

A template for cloning human SLC15A4 was obtained from Addgene (pDONR221_SLC15A4, #131910). cDNA for TASL (Cxorf21:476835) was obtained from Genetic Perturbation Screening core in UF Scripps. For transient transfection, SLC15A4 or TASL cDNA was cloned to pRK5 backbone in frame with either an N-terminal HA tag or a C-terminal FLAG tag. For lentiviral transduction, SLC15A4 cDNA was cloned into pLEX305 backbone in frame with a N-terminal HA tag. To construct transport assay cell line, gBlocks containing the codon-optimized human SLC15A3, human SLC15A4, mouse SLC15A4, and human NOD2 sequences were ordered from Integrated DNA Technologies. Human SLC15A3, human SLC15A4, mouse SLC15A4 and human NOD2 sequences were cloned into the BamH1 site of pLPC retrovirus backbone containing an mCherry protein in frame with a Glycine-Serine bridge using the NEBuilder HiFi Assembly kit (New England Biolabs).

Mutational disruption of the two dileucine motifs to induce plasma membrane localization was performed in both the mouse and human SLC15A4 constructs using using Q5® Site-Directed Mutagenesis Kit (New England Biolabs), which lead to the generation of the following mutants: human SLC15A4 (L14A, L15A, L318A, V319A) and mouse SLC15A4 (L8A, L9A, L319A, V320A). Primers: SLC15A4 Mut1 F (AAGGGCACCTgcggccGGCGCAAGACGGGCAGCCGC); SLC15A4 Mut1 R (TCCCCAGCCCCACCGCCG) (Undercase represents site of mutation); SLC15A4 Mut2 F (TGTGAAAGCTgcggccAAGATTGTCC); SLC15A4 Mut2 R (TCTTCCACTTTCTCTTCTG); mouse SLC15A4 Mut F1 (GAGAGCTCCTgcggccGGTAGTCGCCGACCC); mouse SLC15A4 Mut R1 (TCTCCCTCCATGGTTCTG); mouse SLC15A4 Mut F1 (TGTAAAGGCTgcggccAAAATCGTAC); mouse SLC15A4 Mut R1 (TCCTCTACTTTGTCTTCTG).

#### Statistical analysis

All fluorescent gel scanning and western blots were performed with a minimum of two biological replicates. Cytokine production, THP1-Dual^TM^ reporter signals, B cell activation, and transport assays were performed with minimum of three biological replicates. Statistical analysis for comparison of AJ2-30 treated samples to DMSO and AJ2-18 treated samples was done using an ANOVA followed by multiple comparisons with significance reported for P values of <0.05. Statistical significance for lupus patient samples was assessed using Wilcoxon matched-pairs signed-rank test with significance reported for P values of <0.05. Compound effects at different doses were normalized to vehicle treated cells and fitted with Sigmoidal 4PL curve. All statistical analysis was performed using Graphpad Prism 9. Proteomic analysis was performed with the processing software Proteome Discoverer 2.4 (Thermo Fisher Scientific). Peptide sequences were identified by matching theoretical spectra derived from proteome databases with experimental fragmentation patterns via the SEQUEST HT algorithm(*2*). Fragment tolerances were set to 0.6 Da, and precursor mass tolerances set to 10 ppm with one missed cleavage site allowed. Carbamidomethyl (C, +57.02146) and TMT-tag (K and N-terminal, +229.1629 for 10plex, +304.2071 for 16plex) were specified as static modifications while oxidation (M, +15.994915) was specified as variable. Spectra were searched against the *Homo Sapiens* proteome database (Uniprot, 2018, 42,358 sequences) using a false discovery rate of 1% (Percolator)(*3, 4*). MS^3^ peptide quantitation was performed with a mass tolerance of 20 ppm. Abundances in each channel were normalized to the endogenously biotinylated protein PCCA. The final list of reported proteins was required to have at least two unique peptides. TMT ratios obtained by Proteome Discoverer were transformed with log_2_(x), and p-values were calculated via Student’s two-tailed t-tests with two biological replicates (significant if p <0.05). Quantitative data are listed in the supplementary proteomics data spreadsheet. The mass spectrometric datasets will be deposited after acceptance for publication.

### Isolation of peripheral blood mononuclear cells (PBMCs) and primary cells

#### Healthy donors

All studies with non-diseased primary human cells were performed on samples obtained from Scripps Research’s Normal Blood Donor Service following protocols approved by The Scripps Research Institute Institutional Review Board (IRB). Peripheral blood mononuclear cells (PBMCs) were isolated from heparinized whole blood using a Lymphoprep (STEMCELL Technologies) gradient and washed twice with fluorescence-activated cell sorting (FACS) buffer (Phosphate-buffered saline (PBS), 1X without calcium and magnesium; 2% FBS, and 1 mM EDTA).

Plasmacytoid dendritic cells (pDCs) were isolated via positive selection using the CD304 (BDCA-4/Neuropilin-1) MicroBead Kit (Miltenyi Biotec) following the manufacturers’ instructions. From the flow-through of the pDC isolation, primary human B cells or monocytes were isolated using EasySep™ Human B Cell Isolation Kit (StemCell Technologies, negative selection), or EasySep™ Human Monocyte Isolation Kit (StemCell Technologies, negative selection), respectively, according to manufacturer’s instructions.

#### Patient Samples

Lupus patient samples were obtained through StemCell Technologies Diseased Human Peripheral Blood Mononuclear Cells Catalog. Donor cells were collected following IRB-approved consent forms and protocols. Donors are screened for HIV-1, HIV-2, hepatitis B, and hepatitis C, and medical information of donors is available (**Table S4**). PBMCs were stored in liquid nitrogen upon arrival and thawed with RPMI 1640, 2 mM L-glutamine, 25 mM HEPES, 10% heat-inactivated FBS, 1X non-essential amino acids, and 55 μM β-Mercaptoethanol. Once thawed, cells are cultured at a concentration of 1×10^6^ PBMCs/mL. To assess IFN-α production, cells were stimulated with 1 μM CpG-A, 1 μM CpG-B, or 10 μg/mL R837. 24 hours (hrs) post stimulation, supernatant was collected and stored at −80°C until cytokine quantitation was performed. To assess the effect of compounds on the B cell activation of the lupus patients, cells were stimulated with 10 μg/mL R837 or 1 μM CpG-B. 24 hrs post stimulation, activation markers were assessed by flow cytometry and supernatant was collected and stored at −80°C until cytokine quantitation was performed. To assess antibody production, PBMCs were cultured for 6 days, at which point supernatant was collected.

#### Differentiation and treatment of primary macrophages

Human monocytes were isolated using EasySep™ Human Monocyte Isolation Kit, 50,000 cells are plated in 200 μL differentiation culture media (RPMI with L-Glutamine, 10% Heat Inactivated FBS, penicillin (100 U/mL), streptomycin (100 μg/mL), 10 ng/mL recombinant Human M-CSF Protein (StemCell Technologies)) in a 96 well, flat-bottom plate. Cells were allowed to differentiate for 5 days. On day 5, differentiation media was replaced, and cells were polarized using IFN-γ (100 ng/mL) for 1 hr, Polarized macrophages were then treated with 150 ng/mL MDP, 2000 ng/mL TriDAP, or 5 μg/mL R848. Macrophages were treated for 24 hrs at 37°C and 5% CO_2_, after which supernatant was collected and stored at −80°C until cytokine quantitation.

#### *In vitro* mouse studies

Mouse cells were isolated from a single cell suspension of mouse splenocytes, coming from either Wild-type C57BL/6J or *feeble* mice (PMID: 21045126)(*5*) which had been back-crossed onto the C57BL/6J line. Mouse immune cells were isolated using corresponding isolation kits from StemCell Technologies. Isolated B cells were cultured in complete medium consisting of RPMI 1640, 2 mM L-glutamine, 25 mM HEPES, 10% heat-inactivated FBS, 1X non-essential amino acids, and 55 μM β-Mercaptoethanol.

Bone marrow was harvested from murine femurs and tibias from both Wild-type C57BL/6J or *feeble* mice. Briefly, bones were flushed with RPMI-1640, erythrocytes were lysed with ACK lysis buffer (Fisher), and subsets were obtained through various differentiation protocols. Dendritic cells were generated by culturing bone marrow cells in DMEM supplemented with 2 mM L-glutamine, 25 mM HEPES, 10% heat-inactivated FBS, 1x non-essential amino acids, and 55 μM β-Mercaptoethanol. 10 ng/mL granulocytes macrophage colony-stimulating factor (GM-CSF) (Thermo Fisher Scientific) was added to the culture and cells were allowed to differentiate for 7 days, with GM-CSF containing medium being replenished on Day 3 and Day 6 to derive mouse dendritic cells. Bone marrow-derived macrophages (BMDMs) were generated from bone marrow cells derived in the manner described above in growth medium consisting of RPMI 1640, 2 mM L-glutamine, 25 mM HEPES, 10% heat-inactivated FBS, 1x non-essential amino acids, and 55 μM β-Mercaptoethanol, supplemented with M-CSF (50 ng/mL) (Fisher) for 6 days, with the M-CSF containing medium being replaced on Day 3. Flt3-generated pDCs were differentiated from bone marrow cells as follows. Bone marrow cells were cultured at a concentration of 1.5 x 10^6^ cells/mL in growth medium consisting of DMEM, 2 mM L-glutamine, 25 mM HEPES, 10% heat-inactivated FBS, 1X non-essential amino acids, and 55 μM β-Mercaptoethanol, supplemented with Flt3 (200 ng/mL) (BioXcell) for 8 days respectively.

#### *In vivo* challenge of mice with CpG-A

Prior to challenge, C57BL/6J mice were treated by intraperitoneal injection with AJ2-30 (50 mg/kg; 10% DMSO, 10% Tween-80, 80% 1x PBS by volume) or vehicle (10% DMSO, 10% Tween-80, 80% 1x PBS by volume). Mice were subsequently challenged with 2 μg of CpG-A complexed with 12 μL DOTAP Liposomal transfection reagent (Roche), administered by intravenous tail vein injection as described previously.(*5*) 6 hours following CpG-A/DOTAP administration, blood was drawn by a retinal orbital bleed, and serum was isolated using centrifugation. Serum cytokine levels were quantitated by either Mouse IFN-Alpha ELISA Kit (pbl Assay Science) or the Bio-Plex Pro Mouse Cytokine 23-plex assay (Bio-Rad).

#### *In vivo* pharmacokinetic analysis of AJ2-30

3.96 mg of AJ2-30 was dissolved by sequential vortexing and sonication in 0.132 mL of DMA (N, N-dimethylacetamide), 0.132 mL of Solutol, and 1.056 mL of water to reach concentration of 3 mg/mL. 30 mg/kg of AJ2-30 was dosed in male Balb-c mice through Intraperitoneal (IP) administration. Blood was drawn at different time points (0.25 hrs, 0.5 hrs, 1 hrs, 2 hrs, 4 hrs, 8 hrs, 24 hrs). The desired serial concentrations of working solutions were achieved by diluting stock solution of analyte with 50% acetonitrile in water solution. 10 µL of working solutions (0.5, 1, 2, 5, 10, 50, 100, 500, 1000 ng/mL) were added to 10 µL of the blank mouse plasma to achieve calibration standards of 0.5∼1000 ng/mL (0.5, 1, 2, 5, 10, 50, 100, 500, 1000 ng/mL) in a total volume of 20 µL. Four quality control samples at 1 ng/mL, 2 ng/mL, 50 ng/mL, and 800 ng/mL for plasma were prepared independently of those used for the calibration curves. These QC samples were prepared on the day of analysis in the same way as calibration standards. 20 µL of standards, 20 µL of QC samples and 20 µL of unknown samples (10 µL of mouse plasma with 10 µL of blank solution) were added to 200 µL acetonitrile and proteins were precipitated respectively. Then the samples were vortexed for 30 seconds. After centrifugation at 4 °C, 4000 rotations per minutes (rpm) for 15 minutes (min), the supernatant was diluted 5 times with water, 3 µL of diluted supernatant was injected into the LC/MS/MS system for quantitative analysis. PK parameters were estimated by non-compartmental model using WinNonlin 8.3.

#### Cytotoxicity evaluation

To assess the cytotoxicity of the compounds, multiple concentrations of AJ2-18, AJ2-30, AJ2-32, and AJ2-90 were added to unstimulated PBMCs. PBMCs were plated at a concentration of 1 x 10^6^ cells/mL in medium consisting of RPMI-1640, 2 mM L-glutamine, 25 mM HEPES, 10% heat-inactivated FBS, 1x non-essential amino acids, and 55 μM β-Mercaptoethanol. PBMCs cultured with the aforementioned compounds, necessary control compounds or DMSO were prescribed by the CytoTox 96® Non-Radioactive Cytotoxicity Assay kit, were incubated for 24 hrs at 37°C and 5% CO_2_. Samples were then centrifuged (500 x g, 5 min) and supernatant was transferred to a new plate. CytoTox 96® Non-Radioactive Cytotoxicity Assay was used, following manufacturer’s instructions, to assess cytotoxicity of the samples. Cytotoxicity levels for AJ2-18, AJ2-30, AJ2-32, and AJ2-90 were compared to that of PMBCs treated with DMSO.

#### Culturing and stimulation of immune cells

Synthesized oligodeoxynucleotides (ODNs) and toll-like receptor (TLR) agonists for stimulation of immune cells were purchased from InvivoGen (See Reagents Table). Isolated cells were maintained in growth medium consisting of RPMI 1640, 2 mM L-glutamine, 25 mM HEPES, 10% heat-inactivated (FBS), 1x non-essential amino acids, and 55 μM β-Mercaptoethanol. Human primary pDCs and Flt3 derived mouse pDCs were stimulated with 1 μM CpG-A, 1 μM CpG-B, 5 μg/mL R837, or 5 μg/mL R848 for 24 hrs. Human and mouse pDCs were stimulated with one multiplicity of infection of influenza for 24 hrs. For LL37:DNA complex formation, whole cell genomic DNA was isolated from CAL-1 cells using Qiagen Blood and Tissue Kit. 150 µg of genomic DNA was incubated with 100 µg of LL37 protein and allowed to complex for 1 hour in 250 µL complete RPMI growth medium (see below). Complexed DNA was added to primary pDCs to obtain a final concentration of 10 µg/mL of LL37. Human primary monocytes were stimulated with the following concentrations of reagents: 5 μg/mL R848 or R837, 100 ng/mL LPS, 100 ng/ml Pam3CSK4, or 5 μg/mL VACV-70/ LyoVec™ for 24 hrs. Both mouse and human B-cells were stimulated at the following concentrations unless otherwise noted: R837, 10 μg/mL; LPS, 10 μg/mL; CpG-B 1 μM; and Pam3CSK4, 1 μg/mL. For quantitation of cytokines and analysis of B cell activation, cells were stimulated for 24 hrs. For quantitation of *in vitro* immunoglobin production, cells were stimulated for 6 days. Following treatment, cells were centrifuged (500 x g for 6 min) and the supernatant was transferred to new plate. Supernatants were stored at −80°C until cytokine quantitation was performed.

#### Quantitation of cytokines and immunoglobulin levels

Quantitation of interferon levels were determined by ELISAs using the Human and Mouse IFN-α and IFN-β kits according to manufacturer’s instructions. Broad cytokine analysis of cellular supernatants was performed using the Bio-Plex Pro Human Cytokine 27-plex Assay and the Bio-Plex Pro Mouse Cytokine 23-plex Assay. Analysis of individual analytes and immunoglobulin levels was performed using individual ELISA kits following manufacturer’s instructions. ELISAs were read using the ClarioStar plate reader (BMG Labtech). Bio-plex cytokine assays were run on Bio-plex 200 System (Bio-Rad).

#### *In situ* labeling of living cells by FFF probes

CAL-1 cells or PBMCs were incubated with serum-free media containing indicated probes, and, if applicable, competitors or DMSO vehicle for 30 mins at 37 °C under a 5% CO_2_ atmosphere. The cells were then irradiated with UV light (365 nm, Stratagene, UV Stratalinker 1800 with 5 × Hitachi F8T5 UV bulbs) for 20 min, scraped, washed twice with ice-cold DPBS (1,400 x g, 4 °C, 3 min), and collected into 15 mL Falcon tubes. Cell pellets were either directly processed or stored at −80 °C until the next stage of processing.

#### Preparation of samples for chemoproteomic analysis

Cell pellets were resuspended in 500 µL ice-cold DPBS and lysed by sonicating three times using a Branson Sonifier probe (15 milliseconds (msec) on, 40 msec off, 15% amplitude, 1 second (sec) total on). Protein concentrations were normalized (2 mg/mL in 500 µL with ice-cold DPBS) using a DC Protein Assay (Bio-Rad) with absorbance measured using a Tecan, Infinite F500 plate reader following manufacturer’s instructions. For “click-chemistry”, a mix of the following solutions was added to each 500 μL sample, tris((1-benzyl-4-triazolyl)methyl)amine (TBTA, 30 μL, 1.7 mM in DMSO/*t*-BuOH 1:4 v/v), tris(2-carboxyethyl)phosphine (TCEP, 10 µL, 50 mM), biotin-PEG3-azide (5 µL, 100 µM), and CuSO_4_ (10 µL, 50 mM) and samples were shaken at room temperature (RT) for 1 hr. After the “click-chemistry” reaction, 2.5 mL of ice-cold 4:1 MeOH/CHCl_3_ solution and 1 mL ice-cold DPBS was added to each sample to precipitate proteins. The samples were then vortexed and centrifuged (3,200 x *g,* 4 °C, 10 min), forming a protein disc. The supernatant was removed, and the samples were washed with 4:1 MeOH/CHCl_3_, resuspended by sonication and centrifuged as above. Pellets were resuspended in freshly prepared urea solution (500 µL DPBS, 6 M) with SDS (10 µL of 10% *w/v*), Protein reduction was carried out by the addition of a freshly prepared 1:1 solution (50 µL total, 25 µL each solution) of TCEP (200 mM in DPBS) and K_2_CO_3_ (600 mM in DPBS) and incubating the samples for 30 min at 37 °C. Followed reduction, proteins were alkylated by the addition of freshly prepared iodoacetamide (70 µL, 400 mM in DPBS) and incubated for 30 min at RT in the dark. A solution of SDS (130 µL, 10% in DPBS *w/v*) was added, and samples were diluted with DPBS (5.5 mL) for streptavidin enrichment. A streptavidin-agarose slurry (100 µL, 50%, Pierce) was added to each tube and rotated for 1.5 hr at RT. The beads were pelleted by centrifugation (750 x *g*, 2 min, 4 °C) and washed four times (1 x 5 mL 0.2% *w/v* SDS in DPBS, 1 x 5 mL DPBS, 1 x 5 mL of H_2_O, 1 x 5 mL 100 mM TEAB, pH 8.5) before being transferred to 1.5 mL LoBind microcentrifuge tubes with TEAB (1 mL, 100 mM). Beads were resuspended in sequencing-grade modified trypsin (100 µL, 100 mM TEAB pH 8.5, 100 µM CaCl_2_) and incubated at 37 °C overnight with shaking for on-bead digestion. The digest was separated by centrifugation (300 x *g*, 5 min, 4 °C) and transferred to new LoBind centrifuge tubes. The beads were washed with TEAB (50 µL, 100 mM, pH 8.5), and the washing was combined with the previous supernatant. MS-grade acetonitrile (60 µL) was added to each sample before labeling with respective tandem-mass-tags (TMT – Thermo Fisher Scientific) 10plex or 16plex reagents (8 µL, 20 µg/µL) for 1 hr with occasional vortexing at RT. To quench the labeling reaction, hydroxylamine (6 µL, 5% *v/v*) was added to each sample and incubated for 15 min at RT. Formic acid (6 µL) was added to each tube to acidify the samples before drying under vacuum centrifugation. The samples were combined by redissolving the contents of one tube in a solution of trifluoroacetic acid (TFA, 200 µL, 0.1% in water) and transferred into each sample of a respective multiplexed experiment until all samples were redissolved. This stepwise process was repeated with a second volume of TFA solution (100 µL in water) for a final volume of 300 µL. The samples were fractionated using the Pierce high pH Reversed-Phase Fractionation Kit (Thermo Fisher Scientific 84868) according to manufacturer’s instructions. The peptide fractions were eluted from the spin column with solutions of 0.1% triethylamine containing an increasing concentration of MeCN (5 - 95% MeCN; 12 fractions). The fractions were combined into 6 final fractions in a pairwise fashion (fraction 1 and fraction 7, fraction 2 and fraction 8, etc.), dried via vacuum centrifugation, and stored at −80 °C until ready for mass spectrometry analysis.

#### LC-MS analysis of TMT samples

TMT labeled peptides were resuspended in MS buffer A (20 µL, 0.1% formic acid in water) prior to LC-MS analysis. 3 µL of each sample was loaded onto an Acclaim PepMap 100 precolumn (75 µm x 2 mm) and eluted on an Acclaim PepMap RSLC analytical column (75 µm x 15 cm) using an UltiMate 3000 RSLCnano system (Thermo Fisher Scientific). Buffer A was prepared as described above and buffer B (0.1% formic acid in MeCN) were used in a 220 min gradient (flow rate 0.3 mL min, 35 °C) of 2% buffer B for 10 min, followed by an incremental increase to 30% buffer B over 192 min, 60% buffer B for 5 min, 60-95% buffer B for 1 min, hold at 95% buffer B for 5 min, followed by descent to 2% buffer B for 1 min followed by re-equilibration at 2% buffer B for 6 min. The eluents were analyzed with a Thermo Fisher Scientific Orbitrap Fusion Lumos mass spectrometer with a cycle time of 3 sec and nano-LC electrospray ionization source applied voltage of 2.0 kV. MS^1^ spectra were recorded at a resolution of 120,000 with an automatic gain control (AGC) value of 1×10^6^ ions, maximum injection time of 50 msec (dynamic exclusion enabled, repeat count 1, duration 20 sec). The scan range was specified from 375 to 1,500 m/z. Peptides selected for MS^2^ analysis were isolated with the quadrupole (isolation window 1.6 m/z) and fragmented using collision-induced dissociation (CID 30%) with resultant fragments recorded in the ion trap (AGC 1.8×10^4^, maximum inject time 120 msec). A list of TMT 10plex-labeled SLC15A4 peptide molecular weights (573.6666, 872.9958, 463.8029, 688,3862, 1239.1694, 1012.5444, 705.4477, 459.2593, 582.3318, 859.9966, 517.7844, 721.8893, 470.6345, 675.3653) were employed for parallel reaction monitoring (PRM). MS^3^ spectra were generated by high-energy collision-induced dissociation (HCD) with collision energy of 65%. Synchronous precursor selection(*6*) (SPS) was employed to isolate up to 10 MS^2^ ions for MS^3^ analysis to improve sensitivity of MS^3^-based quantitation.

#### Generation of CAL-1 cells stably overexpressing EGFP or HA-SLC15A4

cDNA ORF for EGFP (#19319) and SLC15A4 (#131910) were obtained from Addgene and cloned into pLEX_305 vector (#41390). Lentiviral particles were produced by co-transfecting HEK293T cells with pMD2.G (Addgene, #12259), psPAX2 (Addgene, #12260), and pLEX305 vectors carrying genes of interest using PEI (Polysciences Inc). Media containing viral particles were collected and filtered through a 0.45 μm membrane 72 hrs after transfection. Lentivirus was added to CAL-1 cell culture in a range of dilutions (1:5, 1:10, 1:20, 1:100, 1:500) with 10 µg/mL polybrene and centrifuged in 15 mL falcon tubes at 800 g for 30 min at 32℃. Transduced cells were selected with 2 µg/mL puromycin and cultured in RPMI1640 complete media containing 1 µg/mL puromycin.

#### Gel-based analysis of crosslinked HA tagged proteins in cells

Probe labelled cell pellets were lysed in Dulbecco’s PBS (DPBS) buffer supplemented with 1% NP-40 (v/v) and 1X halt^TM^ protease inhibitors cocktail (Thermo Fisher Scientific). Protein concentration was normalized to 1.5-2.0 mg/mL in 1 mL volume with lysis buffer. Anti-HA magnetic beads (Thermo Fisher Scientific) was equilibrated with lysis buffer and add to normalized lysate at dose of 20 μl original slurry/1 mg lysate. HA-tagged proteins were enriched by rotating overnight at 4℃. Magnetic beads were washed with 500 μl lysis buffer for 6 times. HA-tagged proteins were eluted by adding 50 μL DPBS buffer supplemented with 1% NP-40 (v/v) and 2 mg/mL HA peptide and shaking in incubators at 37℃ for 30 min. Deglycosylation of eluted protein was performed by incubating with 1 μg of recombinant PNGase F at 37℃ (shown in the figure below). 3 μl of freshly prepared “click” reagent mixture containing 0.1 mM tris(benzyltriazolylmethyl)amine (TBTA) (1.5 μL/sample, 1.7 mM in 1:4 DMSO:*t-*ButOH), 1 mM CuSO_4_ (0.5 μL/sample, 50 mM in H_2_O), 25 μM Rhodamine-azide (0.5 μL/sample, 1.25 mM in DMSO), and 1 mM tris(2-carboxyethyl)phosphine HCl (TCEP) (0.5 μL/sample, 50 mM in H_2_O) was added to 25 μl deglycosylated sample and incubated for 1 hr at room temperature in the dark. 10 μl SDS sample buffer (4X stock) was added to quench the reaction. 10-20 μl sample was resolved by 10% SDS-PAGE and visualized for in-gel fluorescence on Bio-Rad ChemiDoc Imaging System. Proteins were transferred to a polyvinylidene fluoride (PVDF) membrane using Trans-Blot Turbo RTA Mini 0.2 μm PVDF transfer kit and blotted with anti-HA antibody in 5% (w/v) non-fat milk in TBST (50 mM Tris, 150 mM NaCl, 0.2% (v/v) Tween-20, pH 7.5) buffer.

#### SLC15A4-mediated NOD2 activation (transport) reporter assay

To generate the SLC15A4 transport assay, the plasmid pGL4.32[luc2P NF-kB-RE Hygro] (Promega) was stably expressed in A549 cells. The various codon optimized mutants of SLC15A4 were cloned in frame with the mCherry protein, bridged by glycine-serine bridge into the pLPC retrovirus backbone using IDT gBlock fragments to enable expression within the reporter containing A549 cells. NOD2 was overexpressed in these cells using the pHAGE lentiviral backbone. Reporter cells were cultured in DMEM supplemented with 2 mM L-glutamine, 25 mM HEPES, 10% heat-inactivated FBS, 1X non-essential amino acids, and 55 μM β-Mercaptoethanol. Prior to use, 2.5 x 10^4^ cells/mL cells were seeded in a 96 well flat-bottom dish and then treated with either compound or vehicle (DMSO). Following treatment, cells were stimulated with MDP (500 ng/mL, unless otherwise indicated) and luminescence is read using the ClarioStar plate reader (BMG Labtech) after 24 hrs.

#### Flow Cytometry

Flow cytometry was performed in the Scripps Research Flow Core using either the NovoCyte Analyzers (ACEA Biosciences, Inc) or the LSRII Analyzers (BD Biosciences). Cells were resuspended in 1X PBS and stained using eBioscience Fixable Viability Dye eFluor™ 780 (Invitrogen) according to manufacturer’s instructions for 20 minutes. The Live/Dead cell stain was quenched, and cells were washed once using 1X FACS buffer. Surface marker staining was then performed in FACS buffer for 30 minutes. Following the staining for surface markers, intracellular staining to assess phosphorylation status of signaling proteins was performed using BD Cytofix/Cytoperm™ Fixation/Permeabilization Solution Kit (BD Biosciences) following manufacturer’s instructions. All FACS data was analyzed using FlowJo version v10.

#### THP1-Dual reporter assay

1 × 10^5^ THP1-Dual cells (InvivoGen, #thpd-nfis) were plated per 96 well, incubated with varying concentration of compounds, and stimulated with 5 μg/mL R848 for 18 hrs. Plates were then centrifuged at 300 x g for 5 mins and supernatants were collected for NF-κB and IRF reporter analysis. 20 μL of supernatants were first added to each well of a fresh white bottom 96 well plate, followed by 50 μL of Quanti-Luc gold (InvivoGen) reagent. Plates were gently tapped for around 20 times to ensure mixing, then luminescence was measured with ClarioStar plate reader (BMG Labtech). For NF-κB reporter analysis, 20 μL of supernatants were first added to each well of another clear bottom 96 well plate, followed by 180 μL of Quanti-Blue (InvivoGen) reagent. Plate was incubated at 37℃ for 5 to 10 mins, optical density was measured at 620-655 nM using ClarioStar plate reader (BMG Labtech).

#### Immunofluorescent imaging (CAL-1 stable cell lines, IRF translocation analysis)

Primary pDC cells were seeded on 0.02% poly-L-lysine (Sigma, P1274) coated coverslips at RT for 15 min. Cells were then fixed in 2% PFA (Electron Microscopy Sciences) for 10 min, and permeabilized with ice-cold methanol at −20°C for 10 min. After fixation/permeabilization, cells were washed for 5 min with PBS, and then blocked with 3% bovine serum albumin (BSA)/PBST (PBS with 0.05% TX-100) for 10 min. Primary antibodies at indicated dilution were prepared with blocking buffer and incubated with cells for 1 hrs. After washing three times in PBST, cells were incubated with Alexa Fluor-labeled secondary antibodies (1:500) for additional 1 hrs. Cells were then washed three times with PBST. Finally, cells were stained with NucBlue Probe (Thermo Fisher Scientific, R37606) for 5 min to visualize nucleus, and then mounted onto slides with ProLong Diamond Antifade Mountant (Thermo Fisher Scientific, P36965). Confocal images were carried out on a Zeiss LSM 780 using a 100× objective (NA = 1.40). 1 Airy Unit was set as a pinhole for each channel. ImageJ and ZEN (Zeiss) software were utilized for image processing and analysis. For IRFs nuclear translocation, around 40-50 cells were chosen randomly from each condition, and thresholds for each channel were set to analyze the colocalization coefficient between IRFs and DAPI. ZEN software was used to quantify the coefficient.

#### Western blot and co-immunoprecipitation

After treatment with compounds and immune agonists for designated time, cells were lysed in RIPA buffer (Thermo Fisher Scientific, 89900) supplemented with Halt protease and phosphatase inhibitor (Thermo Fisher Scientific, 78841) for 20 mins on ice. After protein normalization with BioRad DC protein assay (#5000112), lysates were mixed with 4X SDS sample buffer and resolved by regular SDS-PAGE or SuperSep^TM^ Phos-tag^TM^ gel (WAKO chemicals, #192-18001). Proteins were transferred to PVDF membrane, blocked with 3% BSA or SuperBlock™ T20 (TBS) Blocking Buffer (Thermo Fisher Scientific, 37536) in TBST buffer, sequentially incubated with primary and secondary antibody, and visualized with a Bio-Rad ChemiDoc Imaging System.

For co-immunoprecipitation of overexpressed protein, cell pellets were resuspended in buffer (50 mM HEPES, 250 mM NaCl, 5 mM EDTA, 0.3% (w/v) CHAPS, 1X Halt protease inhibitor (Thermo Fisher Scientific #78438) supplemented with vehicle (DMSO) or the required dose of compound, incubated on ice for 20 minutes, vortexed every 5 minutes, and sonicated (15 milliseconds (msec) on, 40 msec off, 15% amplitude, 1 second (sec) total on). Lysates were cleared by centrifuge (14,000 x g, 10 min, 4℃) and normalized to 1.5-2.0 mg/mL using lysis buffer. HA magnetic beads (20 μL slurry per 1 mg lysates) and corresponding concentrations of compounds were added to lysates, HA tagged proteins (and their interacting partners) and interactome were enriched by incubate overnight at 4℃ and washed three times by 500 μL lysis buffer. For western blot analysis, proteins were eluted by adding 50 μL 1X SDS sample buffer for every 20 μL of HA magnetic beads slurry and shaking at 37℃ for 30 mins. Eluted proteins were analyzed by SDS-PAGE and western blot.

## SUPPLEMENTAL FIGURES

**Fig.S1.**
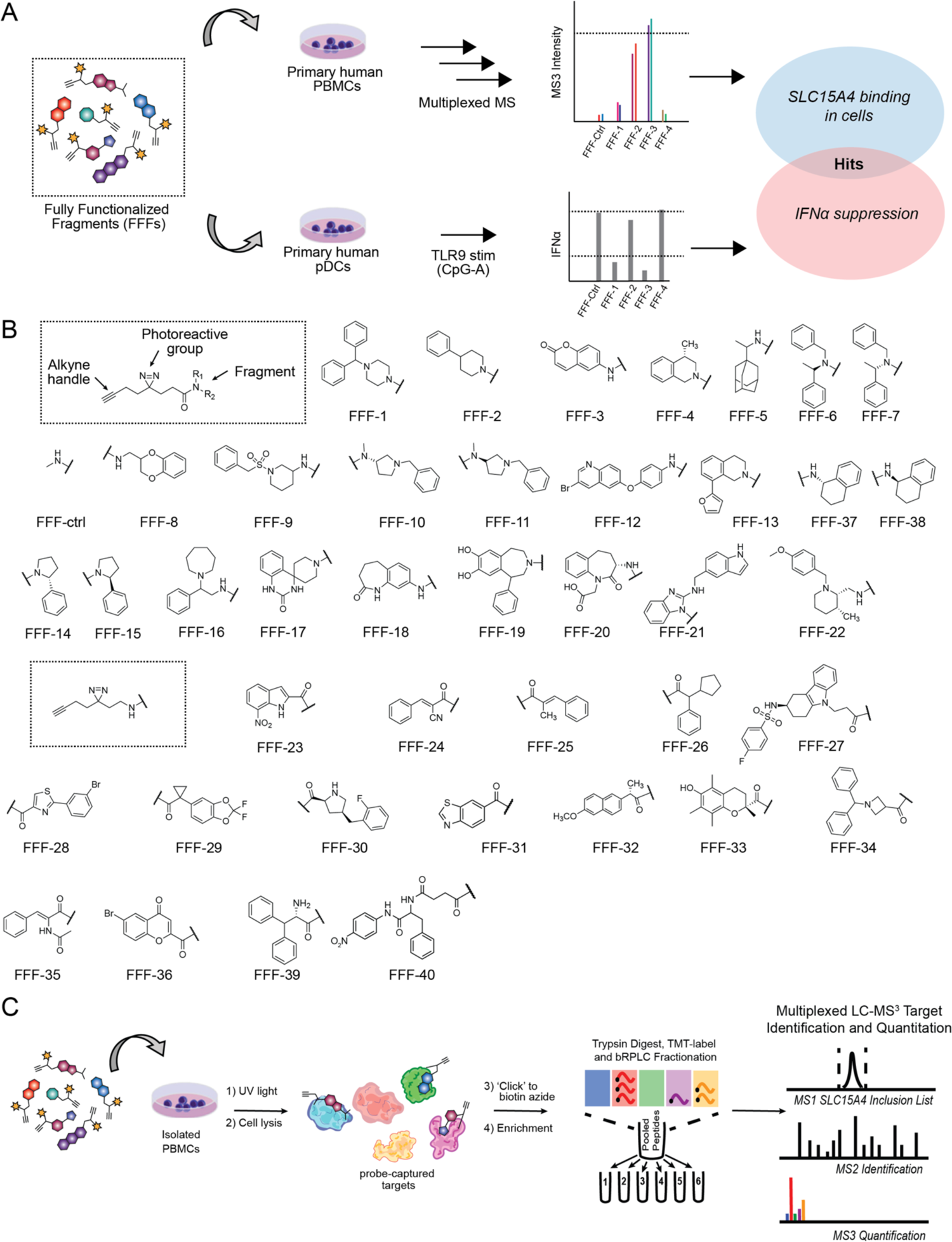
Schematic description of SLC15A4 screening strategy and fully-functionalized fragment (FFF) library. (**A**) Integrated screening approach to identify SLC15A4 inhibitors from a FFF probe library. *In situ* SLC15A4 binding in human PBMCs was analyzed by chemoproteomics. IFNα suppression was screened in human pDCs stimulated with CpG-A. Fragments were considered hits if they enrich SLC15A4 (>5 fold) over a control probe (FFF-ctrl) and suppress IFNα production by more than 50%. (**B**) Chemical structures of FFF library. Each member of the FFF probe library possesses: 1) a ‘variable’ recognition element consisting of structurally diverse small-molecule fragments intended to promote interactions with distinct proteins in human cells; and (2) a structurally minimized ‘constant’ region bearing a photoactivatable diazirine group and alkyne handle, which together enabled UV-light-induced covalent modification and detection, enrichment, and identification of fragment-interacting protein targets. (**C**) Chemoproteomic profiling of SLC15A4 binding by FFF probes. PBMCs isolated from human blood were incubated with 10 μM probe, then underwent UV radiation to induce crosslinking between probe and its bound protein. After a copper-catalyzed azide-alkyne cycloaddition (CuAAC) or click reaction with biotin-azide, proteins bound by probe were enriched by streptavidin beads for trypsinization. Resulting peptides were labelled with TMT10-plex reagents and subjected to multiplexed LC-MS^3^ target identification and quantitation.

**Fig S2.**
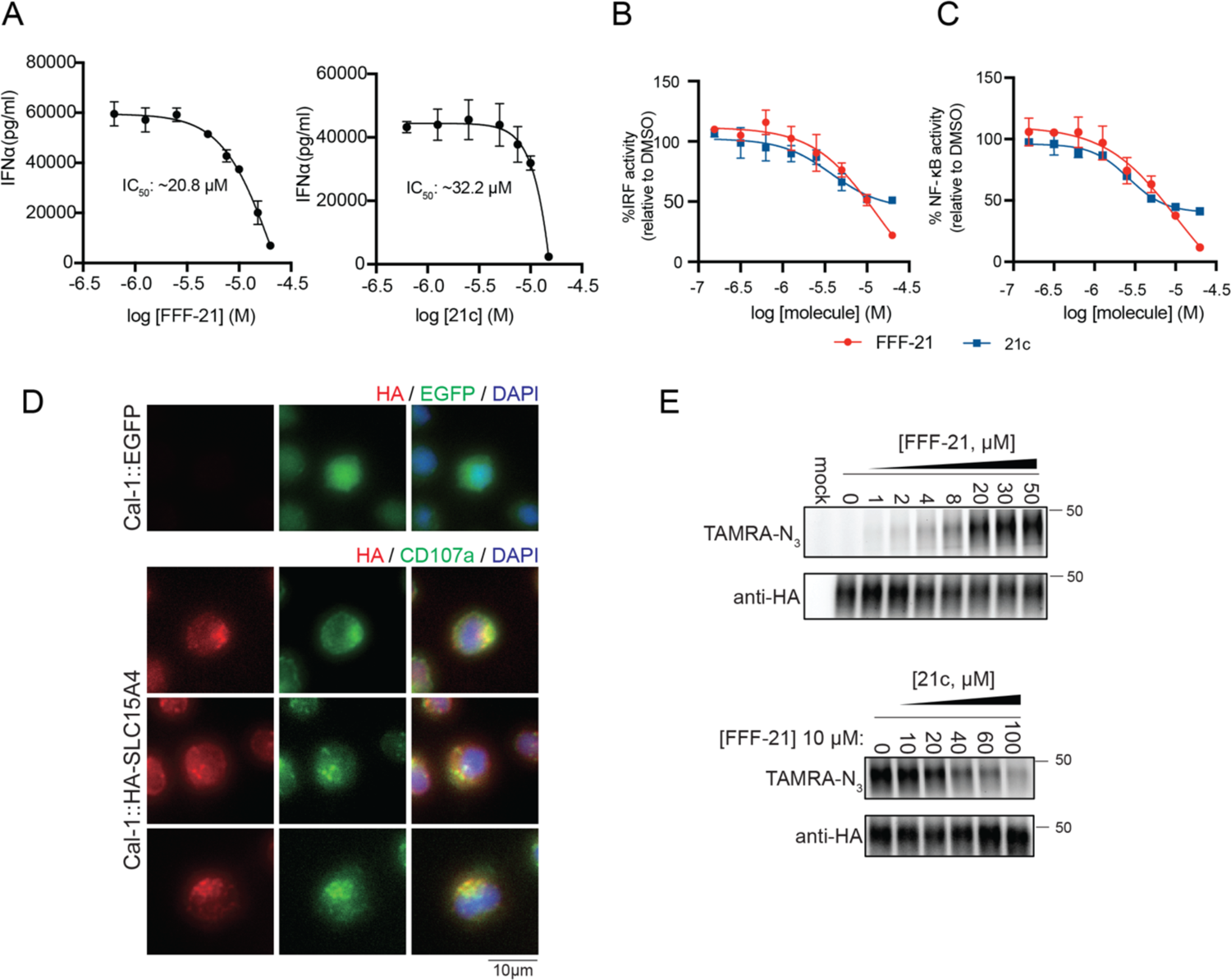
Functional characterization of initial hit FFF and validation of SLC15A4 engagement in cells. (**A**) FFF-21 and 21c inhibit TLR9 mediated IFN production in primary pDCs. Primary human pDCs stimulated with 1μM CpG-A while being treated with escalating doses of compounds for 20 hrs. (**B** and **C**) FFF-21 and 21c inhibit TLR7/8 mediated signaling. THP1 Dual reporter cells were co-incubated with R848 (5 μg/mL) and escalating doses of FFF-21 or 21c. IRF inhibition was monitored by measuring activity of secreted luciferase. NF-kB inhibition was monitored by measuring activity of secreted alkaline phosphatase. (**D**) Immunostaining of SLC15A4 in CAL-1 cells stably overexpressing EGFP or HA-tagged SLC15A4. Cells were co-stained with CD107a as a lysosome marker. Scale bar: 10 μm. (**E**) Gel profiling of HA-SLC15A4 engagement by FFF-21 and 21c in CAL-1 cells stably overexpressing HA-tagged SLC15A4. Cells were treated with DMSO or FFF-21 and binding was visualized by in-gel fluorescence scanning after PNGase treatment (see Methods), click reaction with TAMRA-azide. Anti-HA immunoblot was used as loading control. For competition experiments (bottom), CAL-1 cells were treated with FFF-21 (10 μM) and escalating doses of 21c. Data are representative of two independent replicates.

**Fig S3.**
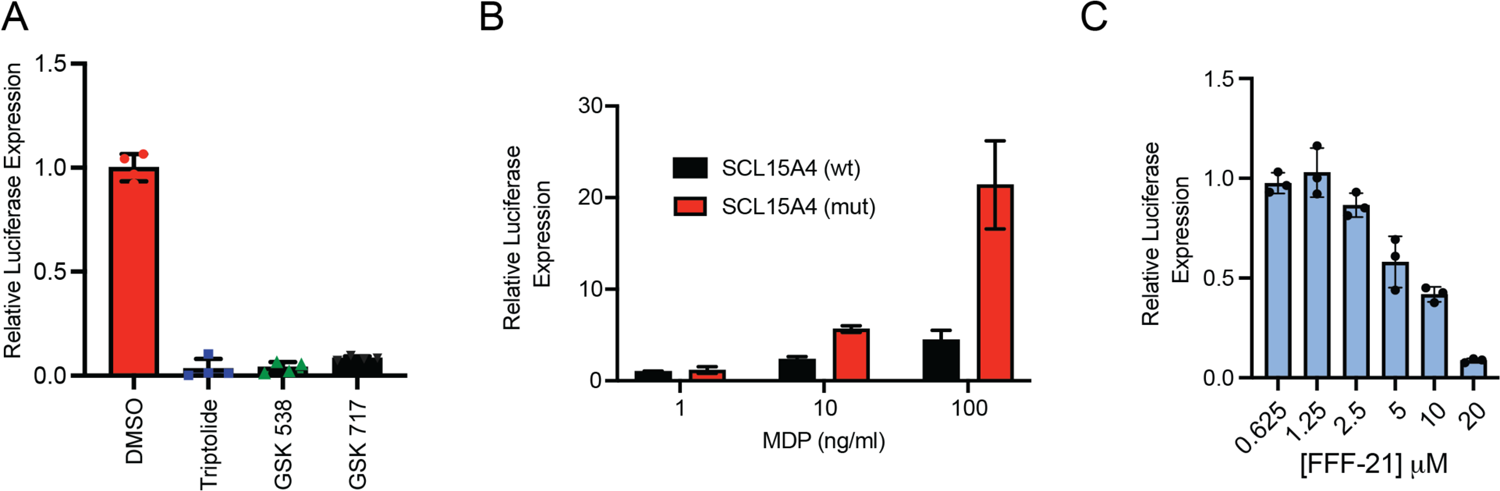
FFF hit suppresses NOD-mediated NF#x003BA;B activation on exposure to NOD ligands in SLC15A4 L14A/L15A A549 cells. A549 cells engineered to express an NF-#x003BA;B-luciferase reporter, NOD2, and either membrane localizing L14A/L15A, L318A, and V319A mutations (*21*) of SLC15A4 or wildtype SLC15A4.(**A**) Chemical intervention against NOD2 signaling pathways by the controls triptolide (an NF#x003BA;B inhibitor), GSK538 (RIPK2 inhibitor), or GSK717 (NOD2 inhibitor). (**B**) Mutant SLC15A4 results in amplified MDP induced NOD activation relative to WT (24 hrs). (**C**) Concentration dependent inhibition of SLC15A4-dependent MDP transport by FFF-21. SLC15A4 transport cells were treated with escalating doses of FFF-21 and MDP for 24 hrs. Results are representative of at least 3 independent experiments and values indicate mean ± SD. Statistical analysis was performed using ANOVA analysis followed by multiple comparisons test, ****p <0.0001.

**Fig S4.**
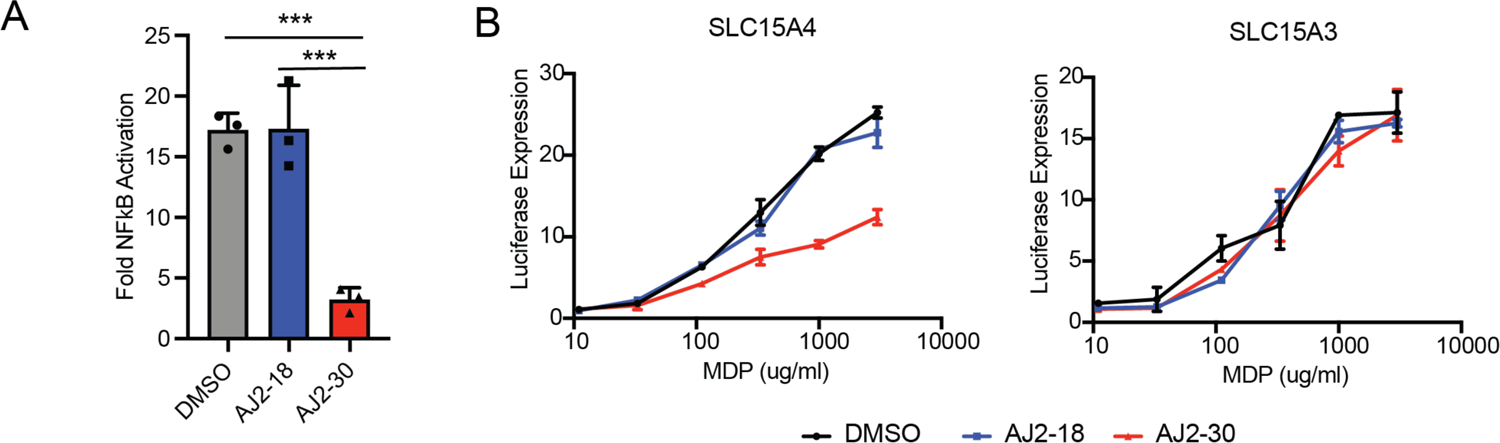
AJ2-30 blocks murine SLC15A4 NOD2 activation and does not affect human SLC15A3 NOD2 activation. (**A**) A549 cells transfected with membrane trafficked murine Slc15a4 as well as NOD2, and the NF-κB-luciferase reporter were stimulated with MDP and treated with AJ2-18 or AJ2-30 (5 μM) for 24 hrs. Luciferase activity was measured in the cells at 24 hrs post-stimulation. (B) A549 cells transfected with an NF-kB-luciferase and NOD2 with either human L14A/L15A SLC15A3 or SLC15A4. Cells were treated with AJ2-18 or AJ2-30 (5 μM) for 24 hr along with escalating quantities of MDP. Results are representative of at least 3 independent experiments and values indicate mean ± SD. Statistical analysis was performed using ANOVA analysis followed by multiple comparisons test. ***p <0.001

**Fig S5.**
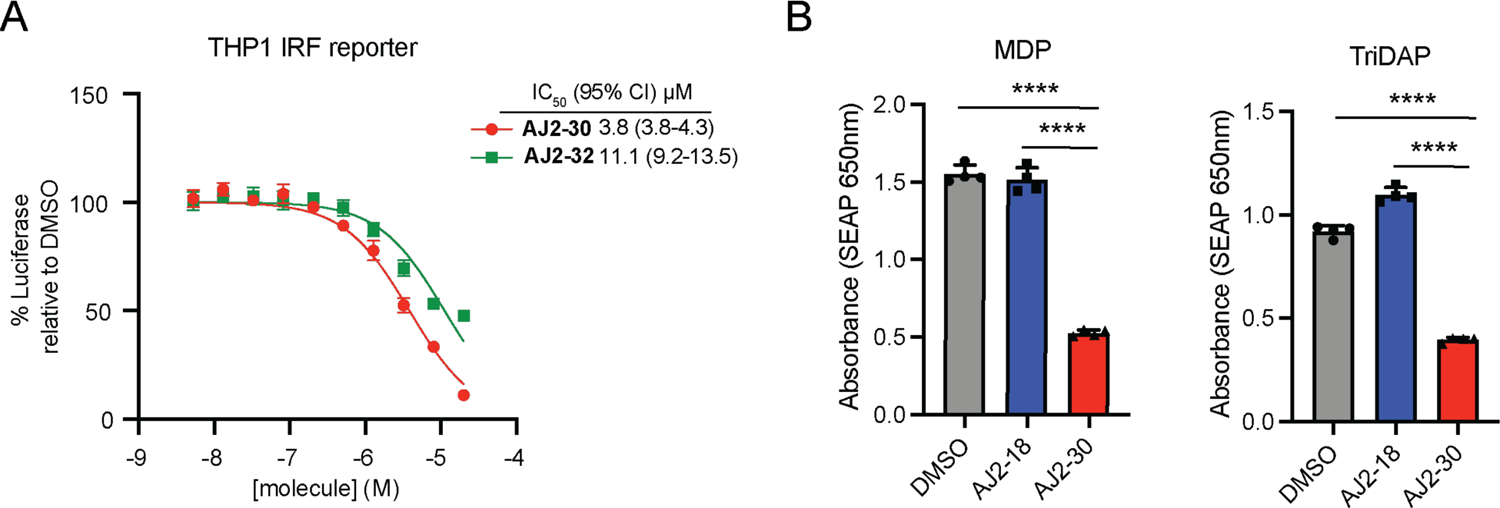
AJ2-30 and AJ2-32 inhibit TLR7/8 mediated signaling. (A). THP1 Dual reporter cells were co-incubated with 5 μg/ml R848 and escalating doses of AJ2-30 or AJ2-32. IRF inhibition was monitored by measuring activity of secreted luciferase. (B). THP1 NF#x003BA;B reporter cells were stimulated with MDP and TriDAP in the presence of AJ2-30 or AJ2-18 and SEAP levels were assessed at 24 hrs in the supernatant. Results are representative of at least 3 independent experiments and values indicate mean ± SD. Statistical analysis was performed using ANOVA analysis followed by multiple comparisons test. **** p ≤ 0.0001

**Fig S6.**
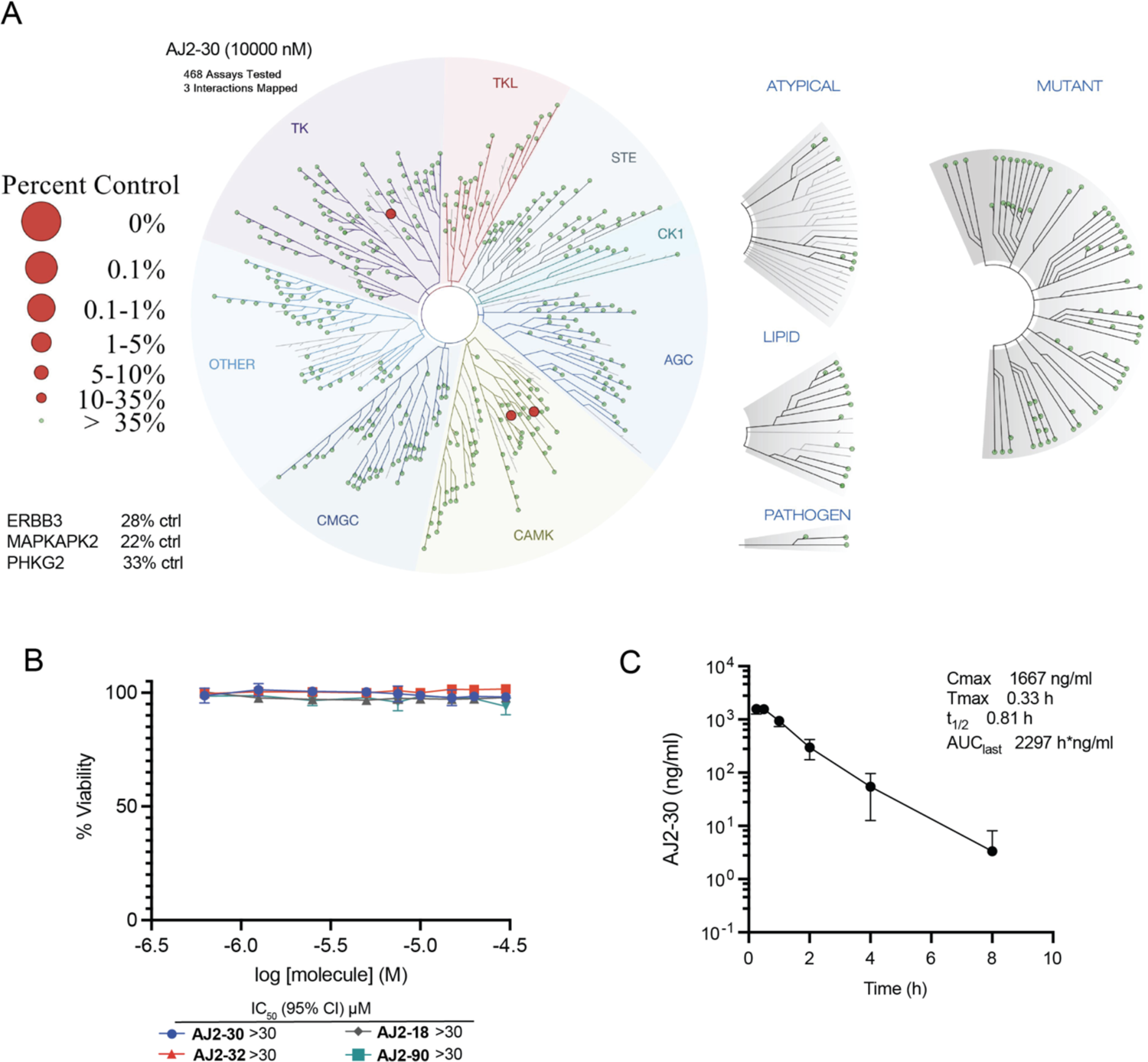
Toxicity and pharmacokinetic characterization of SLC15A4 chemical probes. (**A**) AJ2-30 (10 μM) was submitted for a KinomeScan (DiscoveryX) profiling to quantify interactions with 468 human kinases. Results are displayed as a TREESPOT interaction map. (**B**) Cytotoxicity profiling of compounds AJ2-30, AJ2-32, AJ2-18, and AJ2-90 in human PBMCs. (**C**) Pharmacokinetics of AJ2-30 following intraperitoneal administration (30 mg/kg) to male BALB/c mice. Plasma concentration of AJ2-30 was measured at different time points by LC-MS.

**Fig S7.**
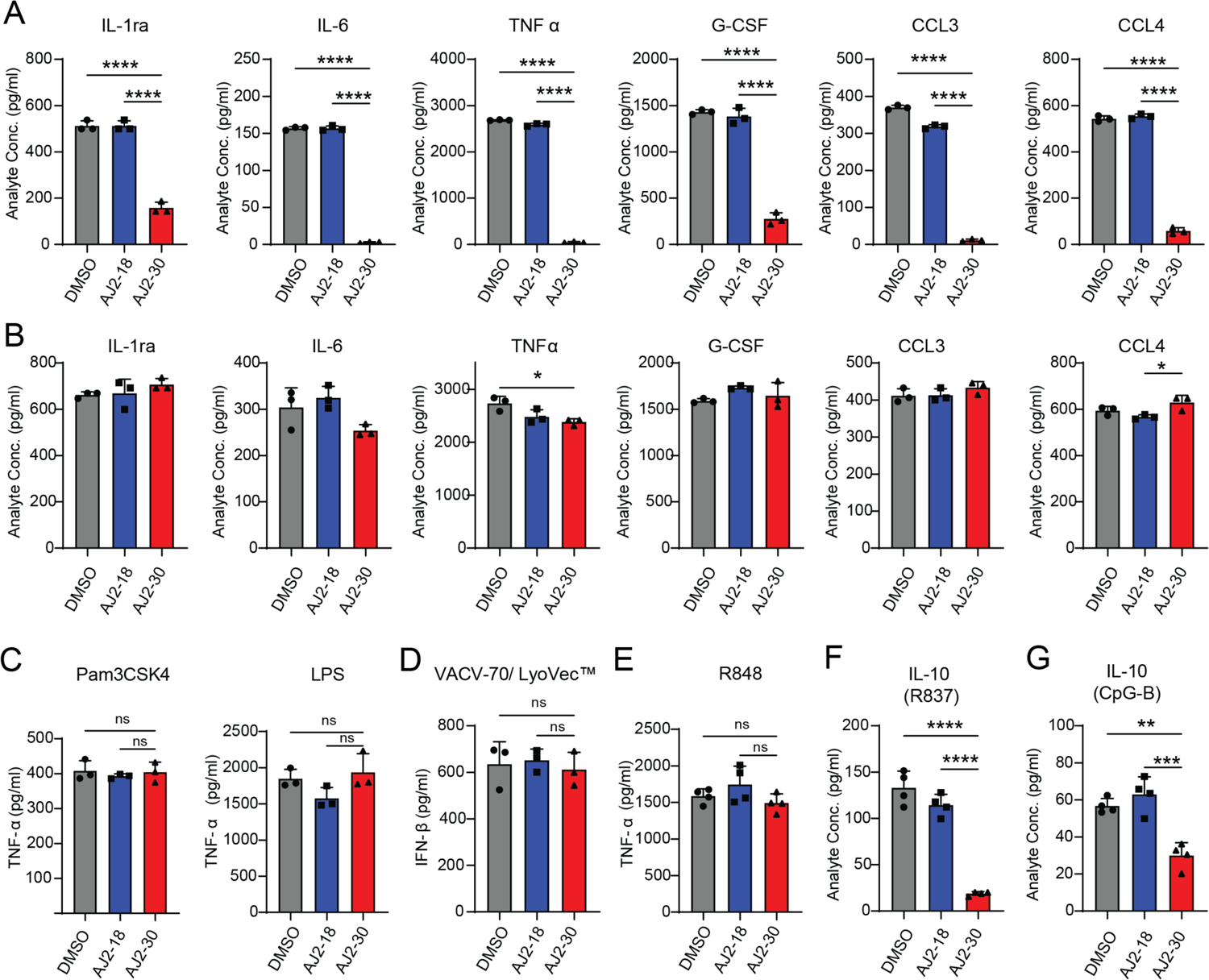
Pharmacological inhibition of SLC15A4 suppresses multiple innate signaling pathways. (**A** and **B**) Secretion of cytokines and chemokines by primary human pDCs in the presence of either AJ2-18 or AJ2-30 in response to stimulation with CpG-A (1 μM) (**A**) or R848 (5 μg/mL) (**B**). (**C** and **D**) Secretion of cytokines from primary monocytes is not inhibited by treatment of AJ2-30 when stimulated by non-endosomal TLR (Pam3CSK4 or LPS) and STING (VACV-70/LyoVec) agonists. (**E**) AJ2-30 does not inhibit TLR7/8 signaling in primary macrophages. Primary human derived macrophages were treated with AJ2-18 or AJ2-30 (5 μM), polarized with IFNγ, and then stimulated with R848 (5 μg/mL) for 22 hrs. (**F** and **G**) *in vitro* secretion of IL-10 from isolated primary human B cells in the presence of either AJ2-18 (5 μM) or AJ2-30 (5 μM) when stimulated by either R837 (10 μg/mL) or CpG-B (1 μM). Cytokine and chemokine levels in supernatants were assessed by Luminex after 24 hrs of stimulation. Statistical analysis was performed using ANOVA analysis followed by multiple comparisons test. * p ≤ 0.05; ** p ≤ 0.01; *** p ≤ 0.001; **** p ≤ 0.0001

**Fig S8.**
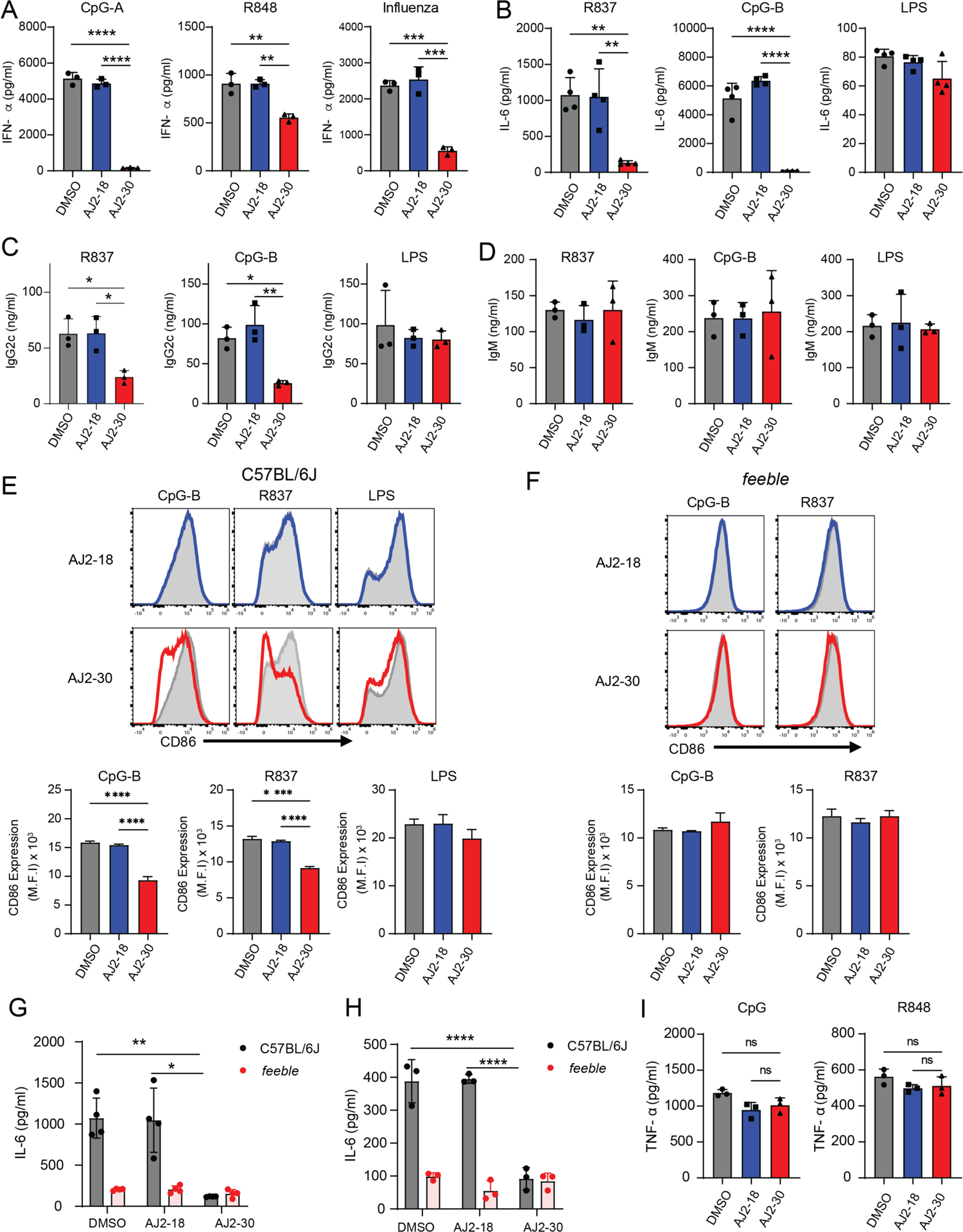
AJ2-30 suppresses TLR7, 8 and 9 activation of mouse immune cells in a *Slc15a4*-dependent manner. (**A**) AJ2-30 inhibits TLR7/8, TLR9, and influenza-induced IFNα production in Flt3L derived pDCs. pDCs were derived from bone marrow using Flt3 ligand and treated with 5 μM AJ2-18 or AJ2-30 for 24 h along with 1 μM CpG-A, 5 μg/ml R-848, or challenged with influenza (MOI=1). The levels of IFNα were quantified by ELISA. (**B**). AJ2-30 inhibits production of IL-6 induced by WT mouse B cells 24hrs after TLR7 and TLR9 stimulation but not TLR4 stimulation. (**C-D**) AJ2-30 inhibits production of IgG2c but not IgM from mouse B cells. *In vitro* IgG2c and IgM secretion from mouse B cells stimulated with TLR ligands for 6 days and quantified by ELISA. (**E** and **F**) AJ2-30 suppresses WT mouse B cell activation in WT but not *feeble* B cells. CD86 expression was detected by flow cytometry on WT (**E**) or *feeble* (**F**) B cells treated with either AJ2-18 (5 μM) or AJ2-30 (5 μM) 24 hrs after TLR4, TLR7, and TLR9 stimulation. Results are representative of two independent experiments. (**G** and **H**) AJ2-30 inhibits production of IL-6 induced by R837 in mouse WT but not *feeble* B cells (**G**) and following stimulation with the bacterial dipeptide MDP in BMDMs (**H**) 24hrs after stimulation. (**I**) AJ2-30 does not inhibit TNFα production from GM-CSF-differentiated dendritic cells. Bone marrow cells were harvested from wild-type C57BL/6J mice and differentiated into conventional dendritic cells with GM-CSF for 10 days. Dendritic cells were then stimulated with CpG-A or R848 for 24 hrs in the presence of DMSO, AJ2-18 or AJ2-30. Statistical analysis was performed using ANOVA analysis followed by multiple comparisons test. * p ≤ 0.05; ** p ≤ 0.01; *** p ≤ 0.001; **** p ≤ 0.0001.

**Fig S9.**
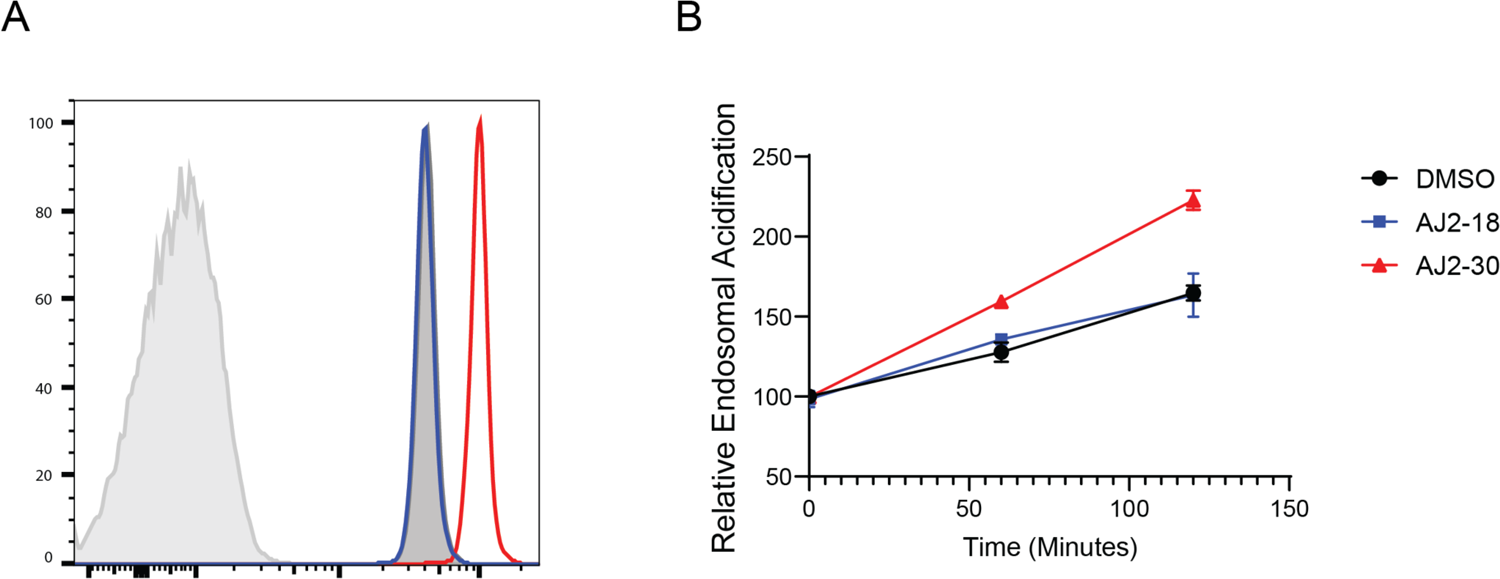
AJ2-30 affects pH value in the endolysosome. (**A**) Endolysosome pH was determined using Lysosensor DND-189. Histogram overlay of mouse B cells in in presence of DMSO (dark gray overlay), AJ2-18 (blue overlay), AJ2-30 (red overlay), 2 hrs post CpG-B stimulation. Unstimulated cells shown in light gray trace. (**B**) Acidification of lysosomes upon CpG-B stimulation in B cells. B cells isolated from wild-type C57BL/6J were treated with either DMSO, AJ2-18 (5 μM), or AJ2-30 (5 μM), stimulated with CpG-B (1 μM) for the indicated periods and labeled with LysoSensor DND-189. The fluorescence was analyzed by flow cytometry.

**Fig S10.**
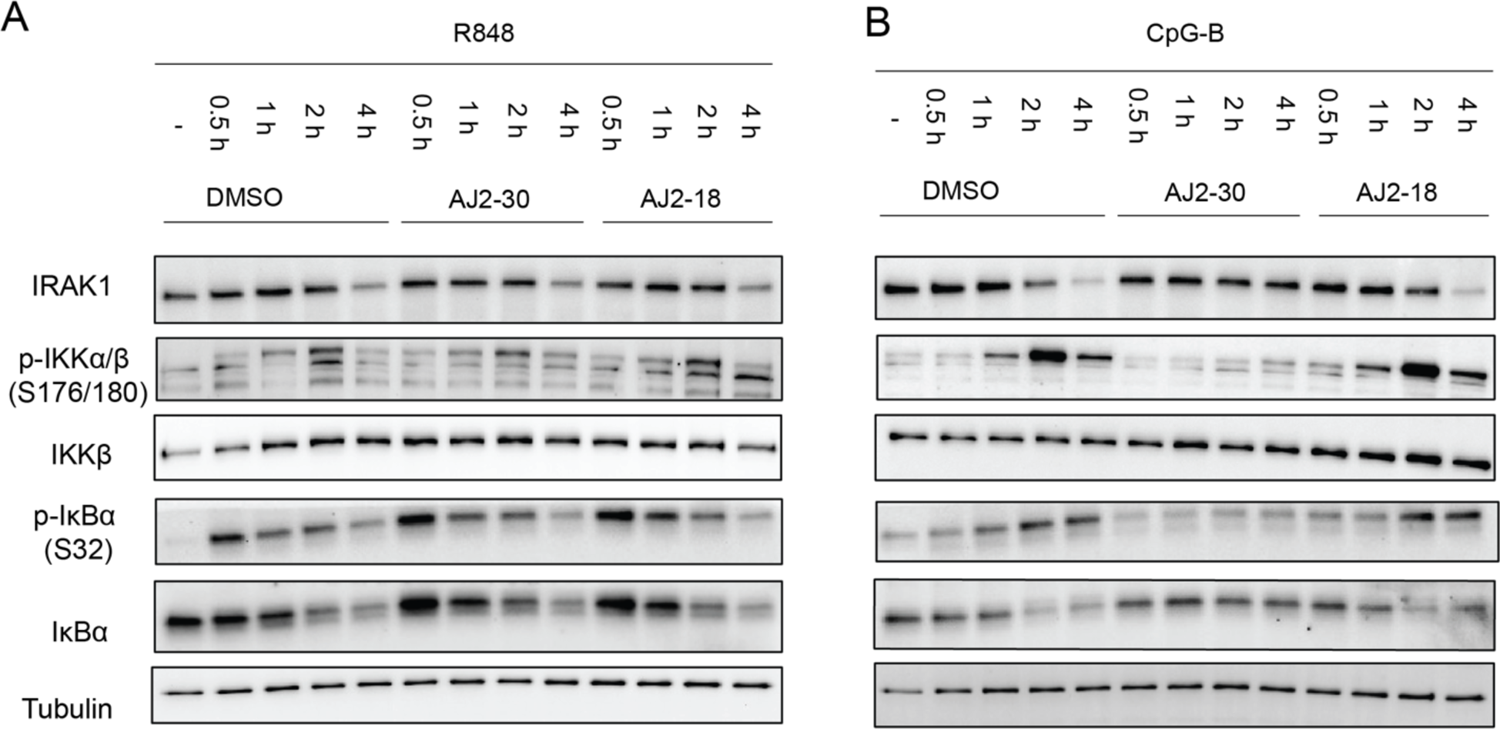
AJ2-30 affects TLR9 proximal signaling in human B cells stimulated but not TLR7/8. (**A, B**) Immunoblot analysis of TLR proximal signaling in B cells isolated from human blood. Cells were co-treated with compounds/DMSO and stimuli (**A**) 5 μg/ml R848 (**B**) 1 μM CpG-B for indicated time points before lysis. Data are representative of two independent experiments.

**Fig S11.**
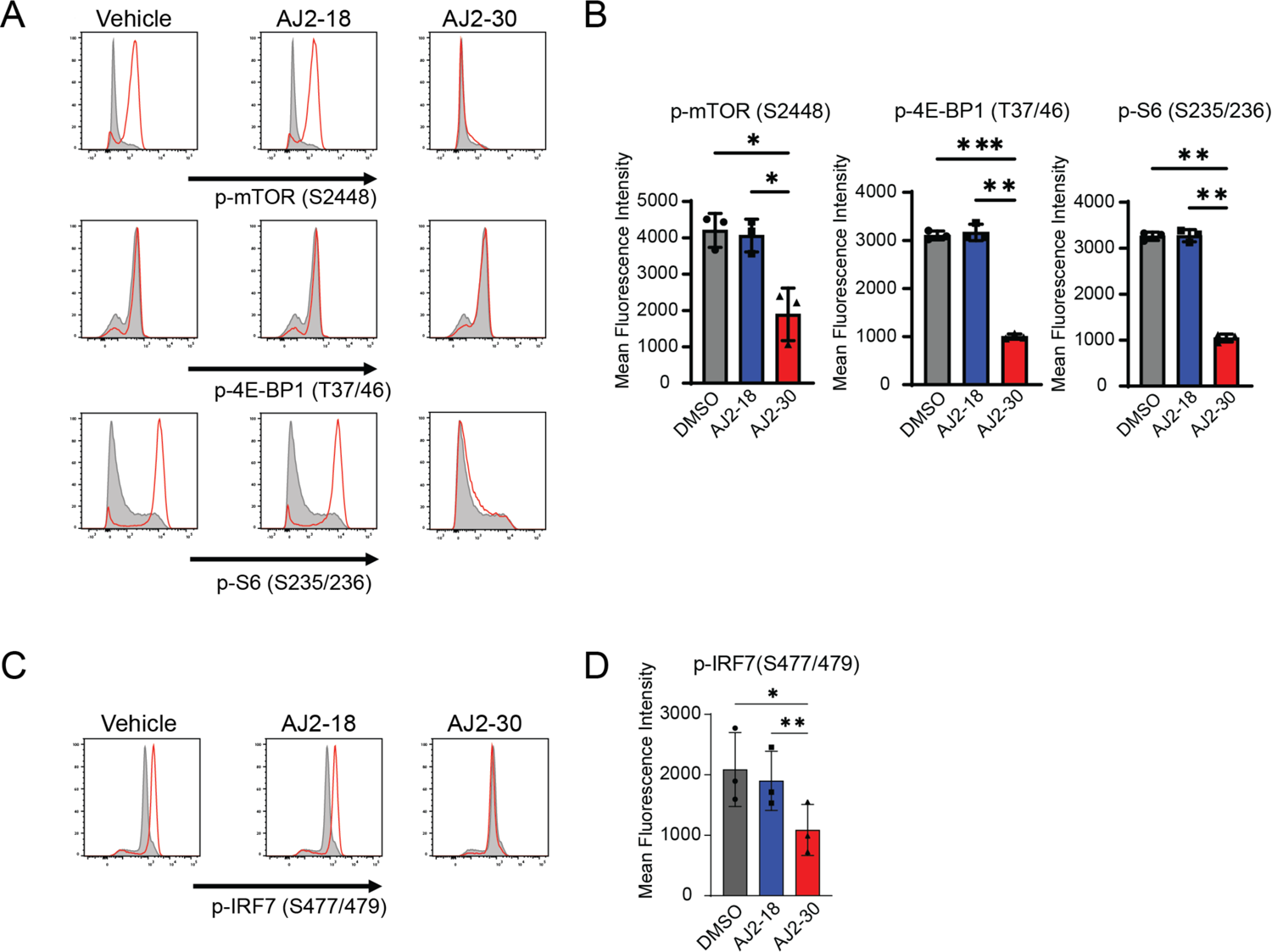
AJ2-30 attenuates mTORC1-IRF7 pathway in human pDCs. Primary human pDCs were treated with AJ2-18 or AJ2-30 while stimulated with CpG-A (1 μM). (**A-B**) mTORC1 activation was assessed by measuring the phosphorylation of mTOR (S2448), 4E-BP1 (T37/46), and ribosomal protein S6 (S235/236) via flow cytometry. (**C-D**) IRF7 activation was assessed by measuring the phosphorylation of IRF7 (S477/479) by flow cytometry. Error bars indicate mean ± SD (*n* = 3). Statistical analysis was performed using ANOVA analysis followed by multiple comparisons test. *p ≤ 0.05, **p ≤ 0.005, ***p < 0.001.

**Fig S12.**
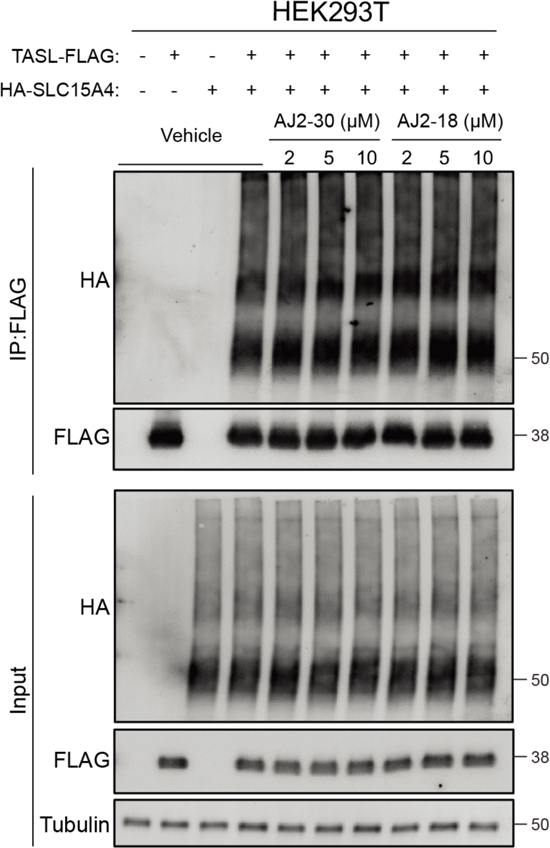
AJ2-30 does not disrupt the interaction between SLC15A4 and TASL. SLC15A4/TASL interaction analysis by immunoprecipitation in HEK293T cells transiently overexpressing HA-SLC15A4 and FLAG-TASL. Cells were treated with escalating doses of compounds or DMSO for 1 hr before lysis and immunoprecipitation by anti-FLAG antibody. After washing, proteins were eluted and analyzed by SDS-PAGE-western blot. Data are presentative of two independent experiments.

**Fig S13.**
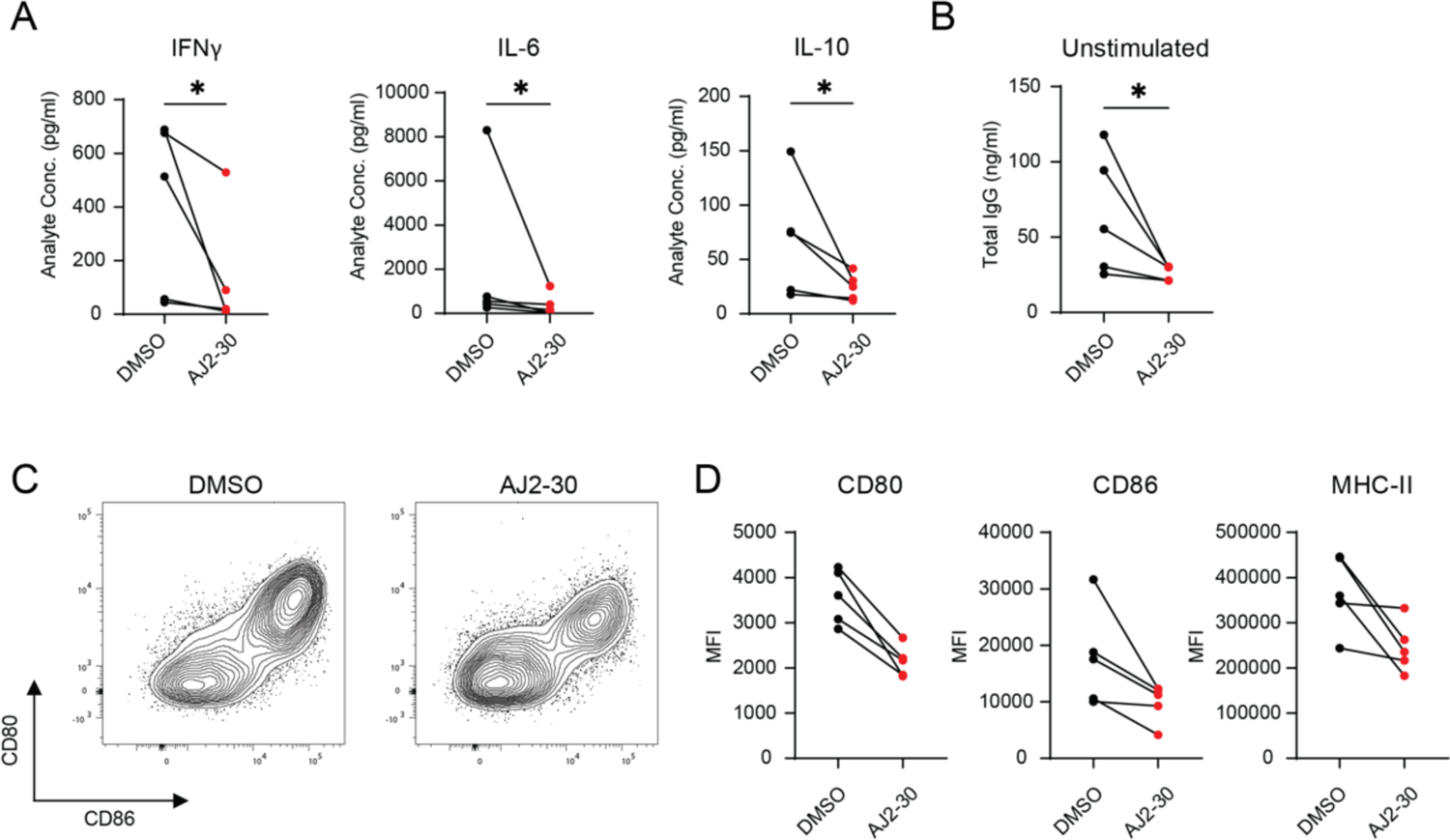
AJ2-30 suppresses steady state cytokine levels and B cell activation markers in lupus patients. (**A**) The levels of secreted cytokines were measured in the supernatants of unstimulated PBMCs in the presence of either AJ2-30 or DMSO. Cytokine and chemokine levels in supernatants were assessed by Luminex multiplex cytokine bead assay after 24 hrs of stimulation. (**B**) Total IgG levels were measured by ELISA to assess the effect of AJ2-30 on suppression of antibody production from unstimulated PBMCs after 6 days of stimulation. (**C-D**) Expression of the costimulatory molecules CD80, CD86, and MHC-II on unstimulated B cells when treated with either DMSO or AJ2-30. Significance was determined using a Wilcoxon matched-pairs signed rank test. *p ≤ 0.05.

**Fig S14.**
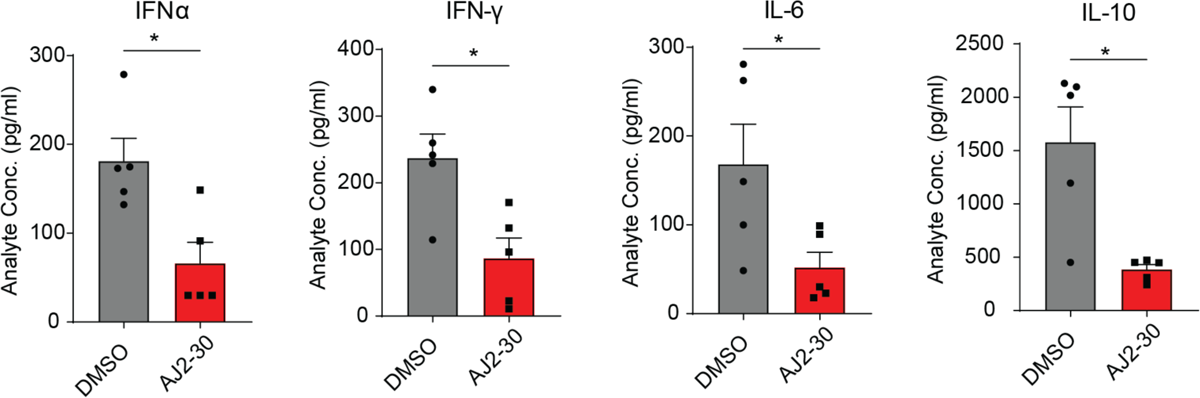
AJ2-30 reduced proinflammatory cytokine levels in mice. Mice were treated with Compound AJ2-30 (50 mpk) or vehicle by intraperitoneal injection and then mice were challenged with the TLR9 ligand CpG-A complexed with DOTAP, administered via tail vein injection and 6 hrs post injection, blood was harvested and cytokine levels were measured. Statistical analysis was performed comparing vehicle and AJ2-30 using an unpaired t-test. *p ≤ 0.05.

**Fig S15.**
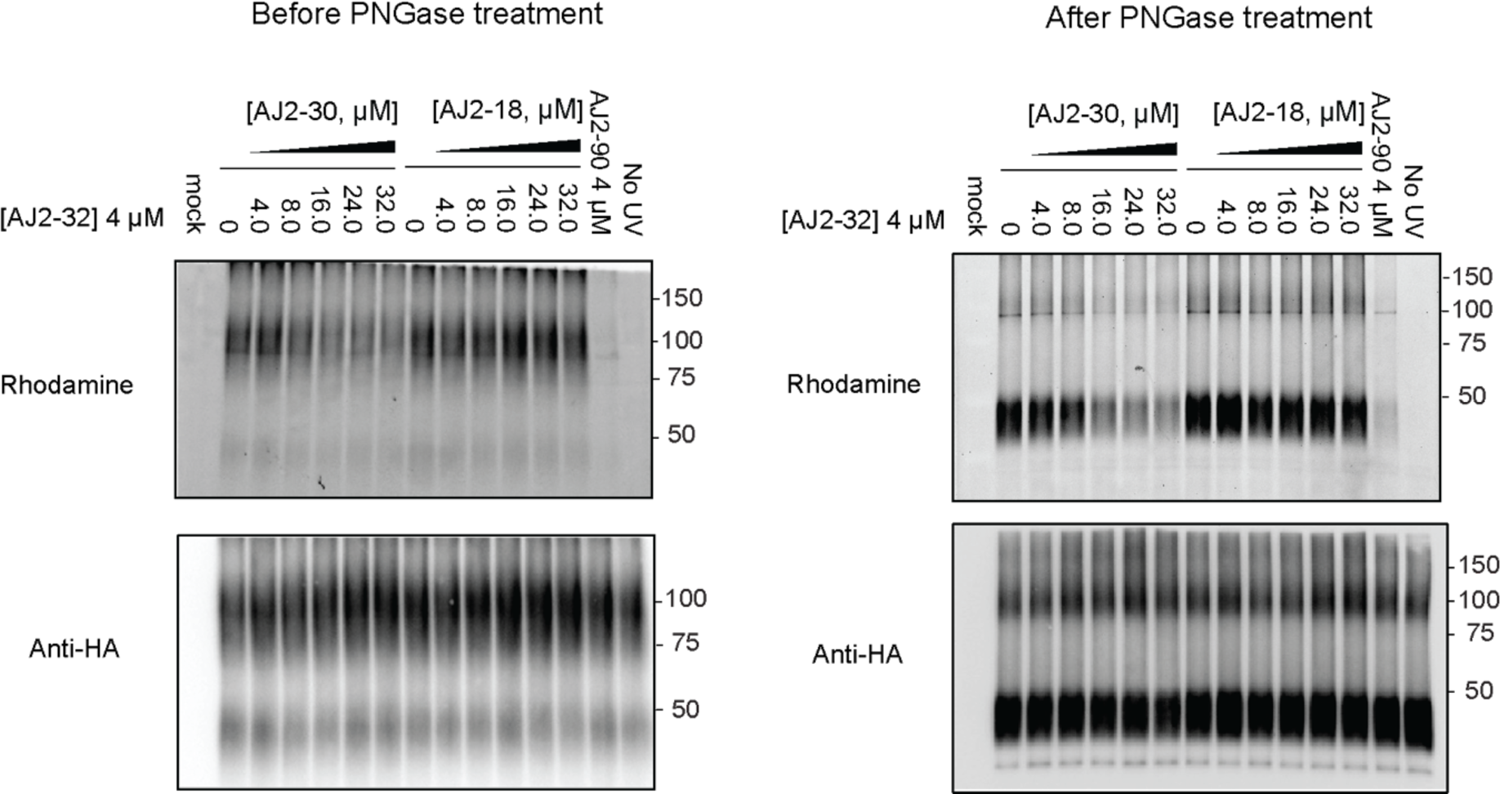
Effect of PNGase treatment on immunoblot profiles of CAL-1 cells stably expressing HA-SLC15A4. SLC15A4 glycosylation reduced when lysates are treated with PNGase as described.

## SUPPLEMENTAL TABLES

**Table S1.** Chemoproteomic profiling of FFF probes in PBMCs. See attached excel table.

**Table S2.**
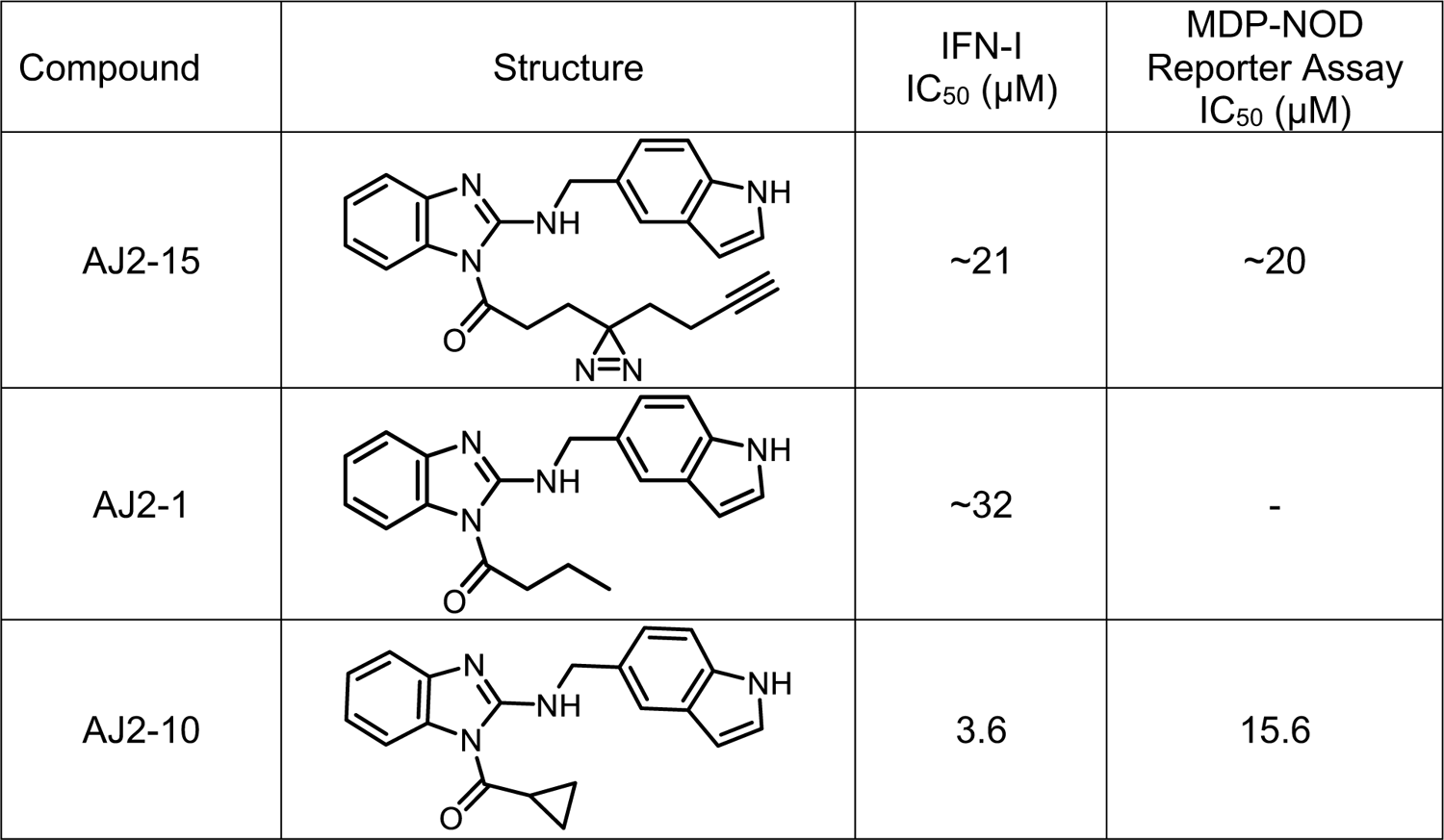
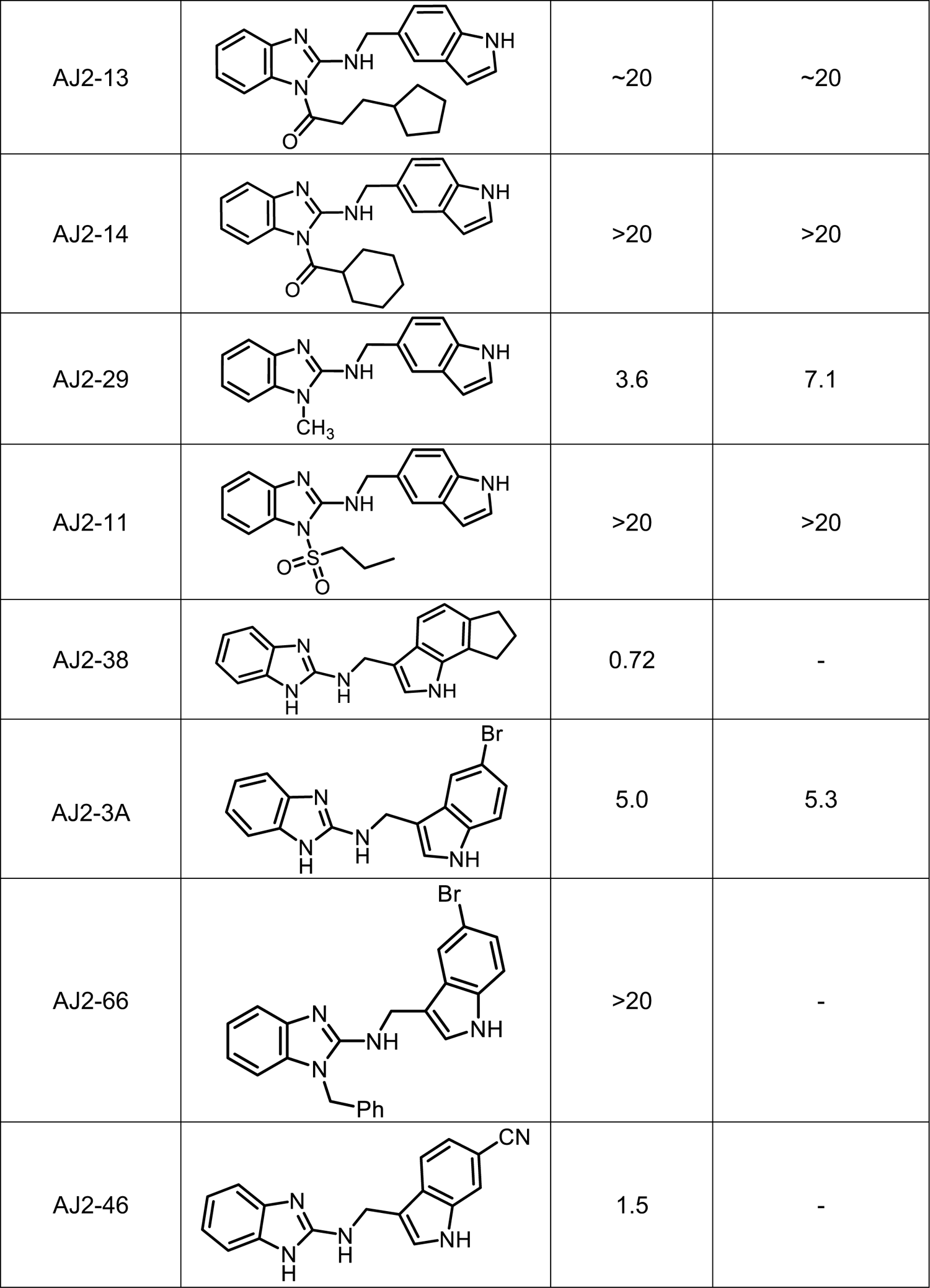
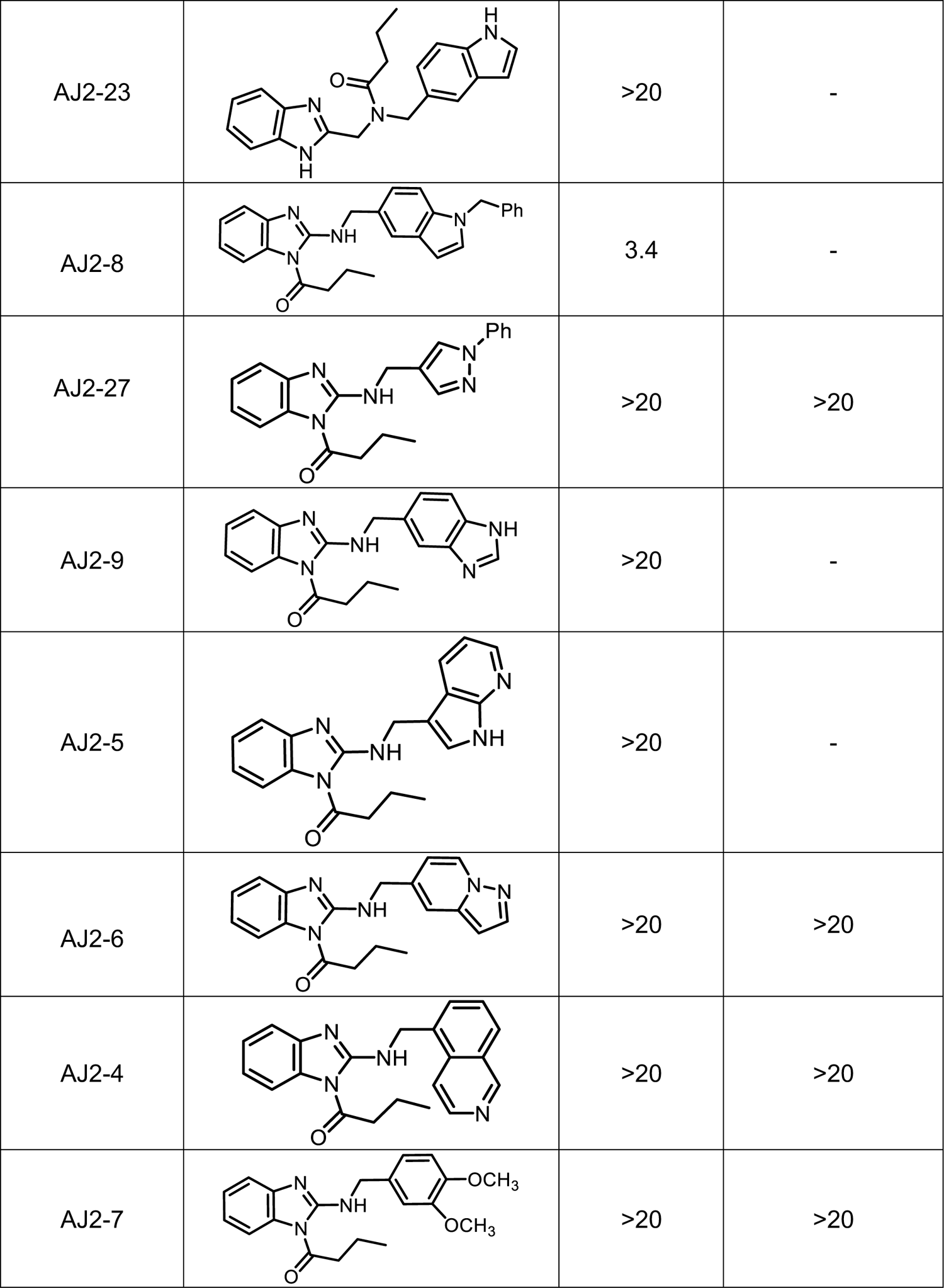
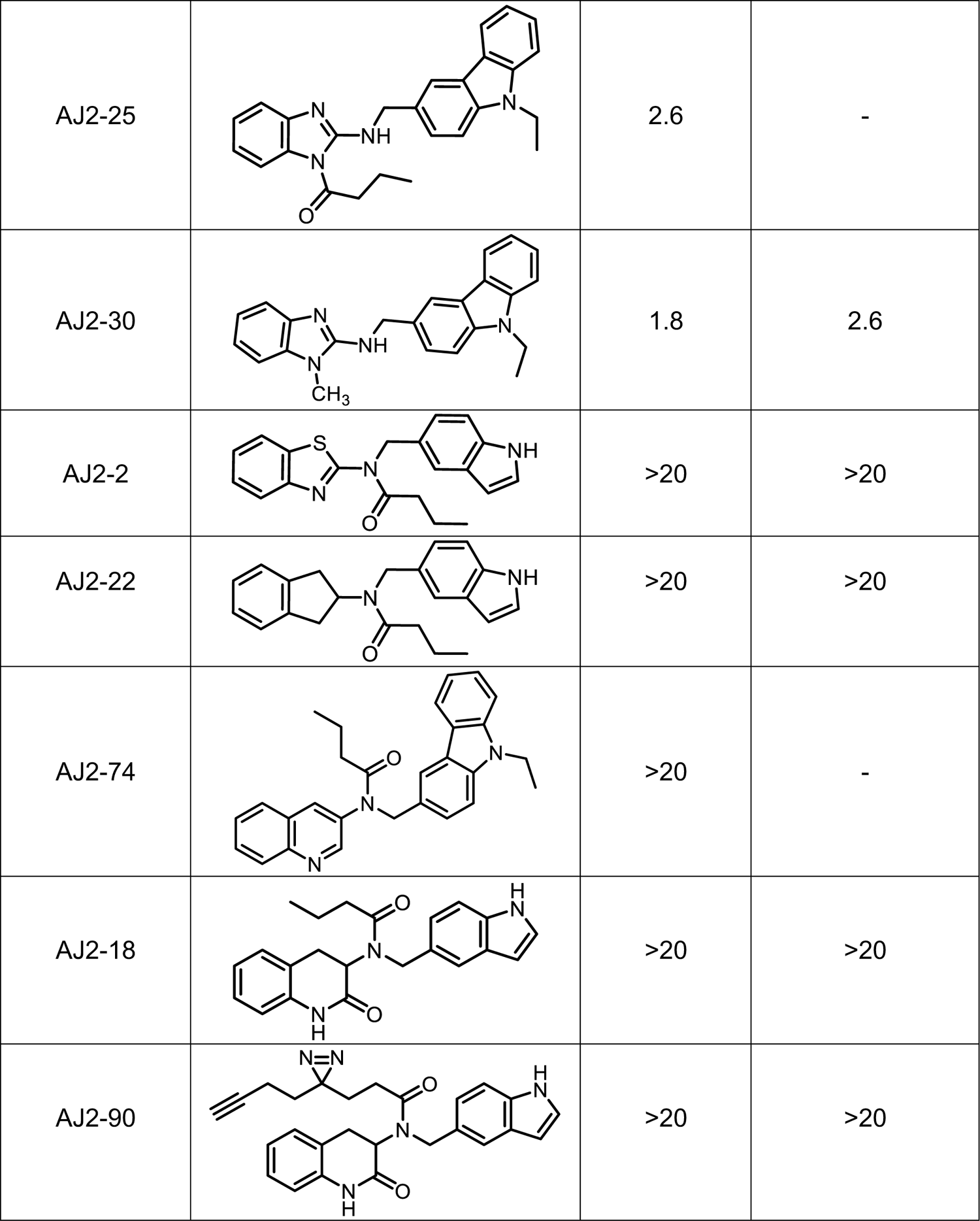
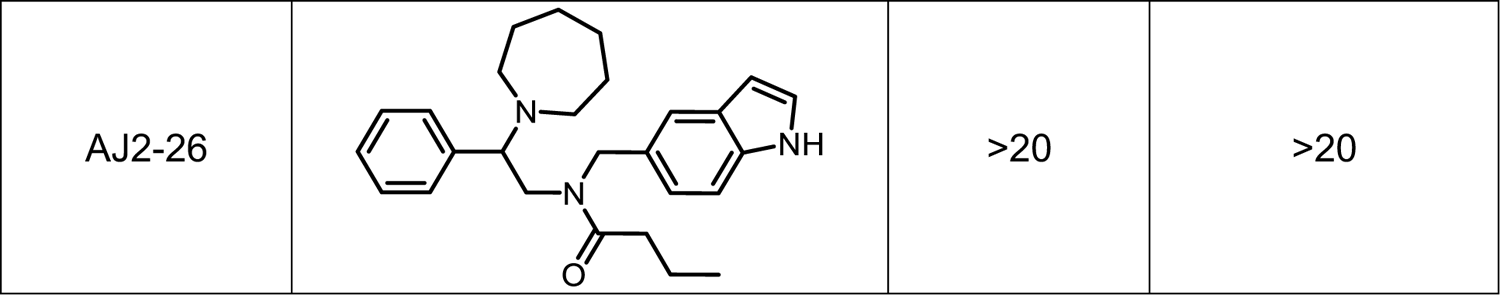
List of Analogs’ inhibition data of IFN-α production in human pDCs and SLC15A4 transport activity.

| Compound | Structure | IFN-I<br>IC<sub>50</sub> (μM) | MDP-NOD<br>Reporter Assay<br>IC<sub>50</sub> (μM) |
| --- | --- | --- | --- |
| AJ2-15 |  | ~21 | ~20 |
| AJ2-1 |  | ~32 | - |
| AJ2-10 |  | 3.6 | 15.6 |
| AJ2-13 |  | ~20 | ~20 |
| AJ2-14 |  | >20 | >20 |
| AJ2-29 |  | 3.6 | 7.1 |
| AJ2-11 |  | >20 | >20 |
| AJ2-38 |  | 0.72 | - |
| AJ2-3A |  | 5.0 | 5.3 |
| AJ2-66 |  | >20 | - |
| AJ2-46 |  | 1.5 | - |
| AJ2-23 |  | >20 | - |
| AJ2-8 |  | 3.4 | - |
| AJ2-27 |  | >20 | >20 |
| AJ2-9 |  | >20 | - |
| AJ2-5 |  | >20 | - |
| AJ2-6 |  | >20 | >20 |
| AJ2-4 |  | >20 | >20 |
| AJ2-7 |  | >20 | >20 |
| AJ2-25 |  | 2.6 | - |
| AJ2-30 |  | 1.8 | 2.6 |
| AJ2-2 |  | >20 | >20 |
| AJ2-22 |  | >20 | >20 |
| AJ2-74 |  | >20 | - |
| AJ2-18 |  | >20 | >20 |
| AJ2-90 |  | >20 | >20 |
| AJ2-26 |  | >20 | >20 |

**Table S3.** Chemoproteomic profiling of AJ2-30, AJ2-32, AJ2-18, and AJ2-90 in both PBMCs and CAL-1 cell line. See attached excel table.

**Table S4.** SLE Patients information

| Donor ID | Date of Draw | Age | Sex | Ethnicity | Weight (kg) | Height (cm) | Smoking Status | Blood Type | Year of Diagnosis | Current Medication |
| --- | --- | --- | --- | --- | --- | --- | --- | --- | --- | --- |
| **RG1817** | 9/17/20 | 47 | F | Caucasian | 83 | 157 | Smoker | A- | 2016 | Levothyroxine |
| **RG1814** | 10/29/20 | 55 | M | Caucasian | 95 | 178 | Non-smoker | A+ | 2013 | Topro-XL |
| **RG2551** | 10/30/20 | 67 | F | Caucasian | 96 | 168 | Non-smoker | A+ | 2010 | Cymbalta,<br>Hydroxychloroquine,<br>Levothyroxine,<br>Lisinopril, Mobic,<br>Prilosec |
| **RG1342** | 11/9/20 | 54 | F | Caucasian | 57 | 165 | Non-smoker | O+ | 2013 | Insulin, Loestrin |
| **RG3041** | 2/5/21 | 48 | F | Caucasian | 83 | 168 | Non-smoker | N/A | 2019 | Gabapentin, Lexapro,<br>Methotrexate |
| **RG3195** | 5/19/21 | 42 | F | Caucasian | 128 | 155 | Smoker | N/A | 2020 | Allergy Shot, Folic acid, Hydrocodone, Levothyroxine, Linzess, Methotrexate, Tizanidine |
| **RG1166** | 2/4/22 | 49 | F | Caucasian | 85 | 157 | Non-smoker | A+ | 2015 | Crestor, Fenofibrate, Lyrica, Metformin, Vascepa, Zyrtec, Estradile |
| **RG3502** | 2/8/22 | 30 | F | Caucasian | 131 | 165 | Non-smoker | A+ | 2006 | Albuterol inhaler, Iron Supplement, Prenatal Vitamins, Zinc, Vitamin A, C, and D |
| **RG2261** | 5/3/22 | 59 | F | Caucasian | 77 | 157 | Non-smoker | O+ | 2015 | None |

## Supplementary Information (Compound synthesis and characterization data)

### I. Synthetic Methods

#### (A) Chemistry material

Chemicals and reagents were purchased from commercial vendors, including Sigma-Aldrich, Fisher Scientific, Combi-Blocks, MedChemExpress, Alfa Aesar and AstaTech, and were used as received without further purification, unless otherwise noted. Anhydrous solvents were purchased from Sigma-Aldrich in Sure/Seal™ formulations. All reactions were monitored by thin-layer chromatography (TLC, Merck silica gel 60 F-254 plates). The plates were stained either with *p*-anisaldehyde (2.5% *p*-anisaldehyde, 1% AcOH, 3.5% H_2_SO_4_ (conc.) in 95% EtOH), ninhydrin (0.3% ninhydrin (w/v), 97:3 EtOH-AcOH), KMnO_4_ (1.5g of KMnO_4_, 10g K_2_CO_3_, and 1.25mL 10% NaOH in 200mL water), iodine or directly visualized with UV light. Reaction purification was carried out using Flash chromatography (230 – 400 mesh silica gel), and Biotage® or preparative thin layer chromatography (pTLC, Analtech, 500-2000 μm thickness). NMR spectra were recorded on Bruker DPX-400 MHz, Bruker AV-500 MHz, Bruker AV-600 MHz spectrometers in the indicated solvent. Multiplicities are reported with the following abbreviations: s singlet; d doublet; t triplet; q quartet; p pentet; m multiplet; br broad; dd doublet of doublets; dt doublet of triplets; td triplet of doublets. Chemical shifts are reported in ppm relative to the residual solvent peak and J values are reported in Hz. Mass spectrometry data were collected on an Agilent 6120 single-quadrupole LC/MS instrument (ESI, low resolution) or an Agilent ESI-TOF instrument (ESI-TOF, HRMS).

#### (B) Synthetic Procedures

##### 1) General synthetic procedure of diazirine containing FFF (Fully Functionalized Fragments) Scheme 1

###### General Procedure a

To a solution of corresponding commercially available carboxylic acid (0.113 mmol) in 3 ml DCM, the corresponding diazirine amine (0.118 mmol), DIPEA (0.354 mmol), EDC-HCl (0.177 mmol), and HOBt (0.177 mmol) were added. The reaction mixtures were stirred at room temperature for 14 to 16 hr. After completion (monitored by TLC) the crude reaction mixture was diluted with DCM (20 mL) and washed first with saturated aqueous NH_4_Cl (10 mL) and saturated aqueous NaHCO_3_ (10 mL) solution, dried over anhydrous Na_2_SO_4_ and volatiles removed by rotary evaporation. Crude products were purified by PTLC or Biotage® SNAP Cartridge KP-Sil Snap 10g with linear gradient of ethyl acetate and hexane over 20 column volumes (CV).

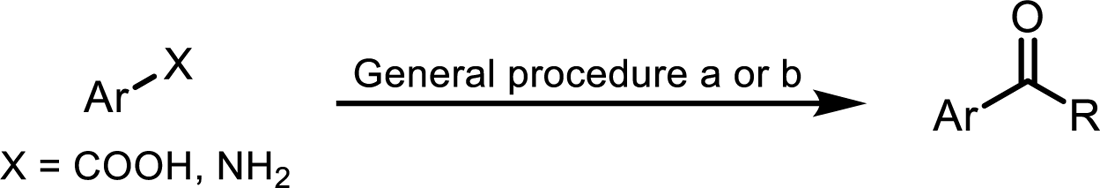

###### General Procedure b

To a solution of corresponding commercially available amine (0.113 mmol) in 3 ml DCM, the corresponding diazirine acid (0.118 mmol), DIPEA (0.354 mmol), EDC-HCl (0.177 mmol), and HOBt (0.177 mmol) were added. The reaction mixtures were stirred at room temperature for 14 to 16 hr. After completion (monitored by TLC) the crude reaction mixture was diluted with DCM (20 mL) and washed first with saturated aqueous NH_4_Cl (10 mL) and saturated aqueous NaHCO_3_ (10 mL) solution, dried over anhydrous Na_2_SO_4_ and volatiles removed by rotary evaporation. Crude products were purified by PTLC or Biotage® SNAP Cartridge, KP-Sil, 10g with linear gradient of ethyl acetate and hexane over 20 column volumes (CV).

##### 2) General Synthetic Scheme 2

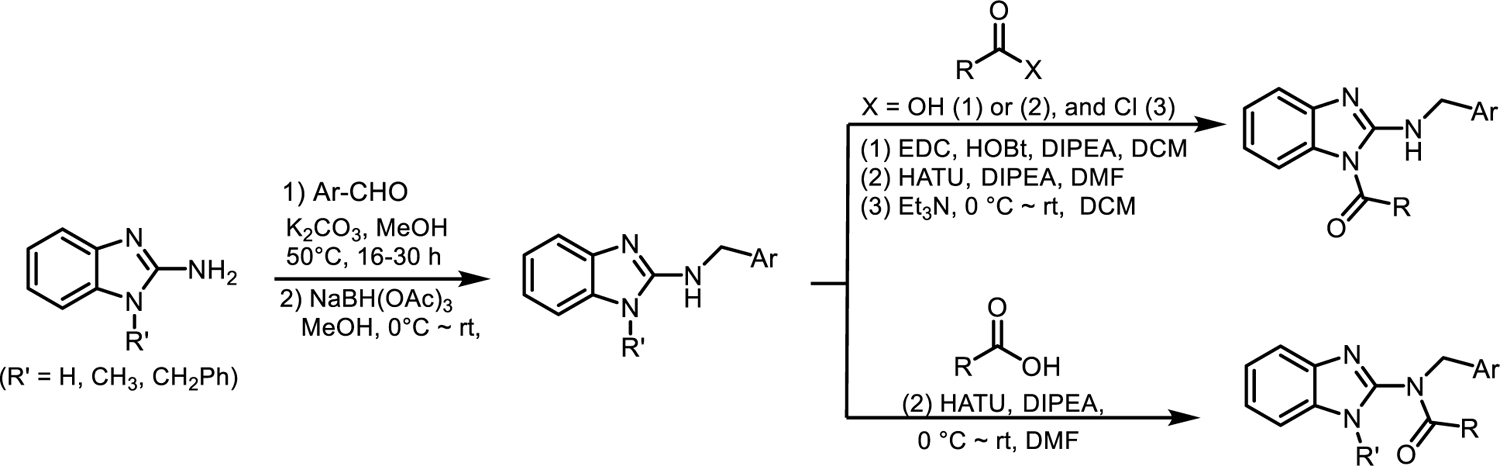

###### General Procedure 1: coupling procedure for the synthesis of benzo[*d*]imidazole amine intermediate (S1)

Synthesized according to reported procedure^3^, to a dried round bottom flask containing solution of commercially available 2-aminobenzimidazole, or 2-amino-1-methylbenzimidazole, or 2-aminoindane derivatives (1.0 eq.) and corresponding aldehyde (1.0 eq.) in dry methanol, potassium carbonate (3.0 eq.) was added, and the reaction mixture was heated at 50 °C for 16 to 30 h. The solvent was filtered to remove the excess potassium carbonate and sodium triacetoxyborohydride (2.0 eq.) was added at 0°C to the solution and resulting mixture was stirred for 8 h at room temperature. After completion (monitored by TLC) the solvent was removed by rotary evaporation, crude mixture was diluted with water and washed with saturated aqueous NaHCO_3_ solution extracted in ethyl acetate, the combined extract was dried over Na_2_SO_4,_ filtered and concentrated in vacuum, purified by flash column chromatography on Biotage^®^ isolera one instrument to give corresponding amines intermediate (**S1**).

###### General Procedure 2: Coupling of amines intermediate with acid

To a vial containing corresponding amine intermediate (1 eq.) in dichloromethane (60 mM relative to S1), commercially available butyric acid or 3-(3-(but-3-yn-1-yl)-3H-diazirin-3-yl)propanoic acid (1.1 eq.), N, N-diisopropylethylamine (3.0 eq.), EDC-HCl (1.5 eq.) and HOBt (1.5 eq.) were added and stirred at room temperature for 4 h to overnight. After completion (monitored by TLC) the crude mixture was diluted with dichloromethane, saturated aqueous NH_4_Cl solution was added and extracted in dichloromethane. The combined extract was washed with saturated aqueous NaHCO_3_ solution, extracted in dichloromethane dried over anhydrous Na_2_SO_4,_ and volatiles removed by rotary evaporation. Crude products were purified by PTLC or flash column chromatography on Biotage^®^ isolera one instrument to give the corresponding products.

###### General Procedure 3: Coupling of amines intermediate with acid

To a solution of corresponding butyric acid or 3-(3-(but-3-yn-1-yl)-3H-diazirin-3-yl)propanoic acid (1.1 eq.) and amine intermediate (1.0 eq.) in dimethylformamide (60 mM relative to S1), the N, N-diisopropylethylamine (3.0 eq.) and HATU (1.1 eq.) were added at 0°C and resulting mixture was stirred at room temperature until amines were fully consumed, as indicated by TLC. The crude mixture was diluted with cold water and extracted in ethyl acetate then combined extract were dried over anhydrous Na_2_SO_4_ and volatiles removed by rotary evaporation. Crude materials were purified by PTLC or flash column chromatography on Biotage^®^ isolera one instrument to give the corresponding products.

###### General Procedure 4: Coupling procedure for synthesis of amide with acid chloride

To a solution of corresponding amine (1.0 eq.) and triethylamine (1.1 eq.) in dichloromethane (0.1 M), and corresponding acid chloride (1.0 eq.) solution in dichloromethane was added over 10 minutes at 0°C, and resulting mixture was allowed to stir at room temperature until starting amines was fully consumed, as indicated by TLC. The crude reaction mixture was diluted with dichloromethane, washed with saturated aqueous NH_4_Cl solution followed by NaHCO_3_ solution, the combined dichloromethane solution dried over anhydrous Na_2_SO_4,_ and volatiles removed by rotary evaporation. The crude materials were purified by PTLC or flash column chromatography on Biotage^®^ isolera one instrument to give the corresponding products.

###### General synthetic Scheme 3

To a solution of 3-amino-3,4-dihydroquinolin-2(1H)-one (1.0 eq.) and corresponding 5-Formylindole (1.0 eq.) in dry methanol, potassium carbonate (3.0 eq.) was added, and the reaction mixture was stirred at room temperature for 18 h. The reaction mixture was filtered on Whatman® filter paper to remove the excess potassium carbonate washed with methanol and sodium triacetoxyborohydride (2.0 eq.) was added at 0°C to the filtered solution and resulting mixture was stirred for 6 h at room temperature. After completion (monitored by TLC) the solvent was removed by rotary evaporation, crude mixture was diluted with water and washed with saturated aqueous NaHCO_3_ solution extracted in ethyl acetate, the combined extract was dried over Na_2_SO_4,_ filtered and concentrated in vacuum, purified by flash column chromatography on Biotage^®^ isolera one with 5-80 % ethyl acetate gradient in hexane to obtain corresponding 3-(((1H-indol-5-yl)methyl)amino)-3,4-dihydroquinolin-2(1H)-one (**S7**) as a white solid.

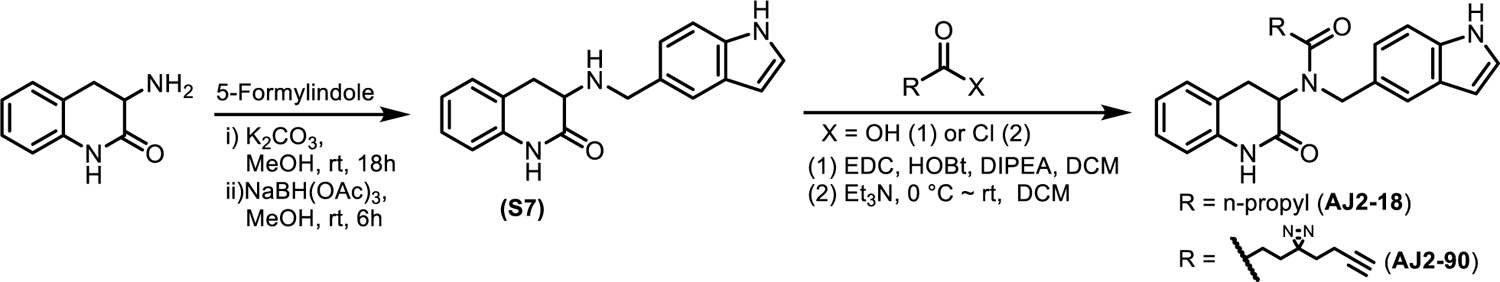

### ii) Characterization data of FFF (Fully Functionalized Fragments) probes

**FFF-1, 2, 3, 6, 7, 10, 11, 14, 15, 37**, and **38** are synthesized according to previously reported procedure.^1, 2^

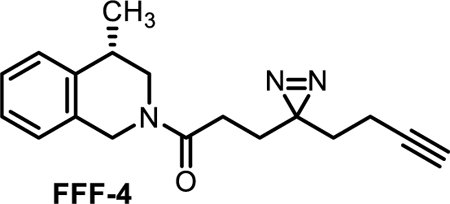

#### (S)-3-(3-(but-3-yn-1-yl)-3H-diazirin-3-yl)-1-(4-methyl-3,4-dihydroisoquinolin-2(1H)-yl)propan-1-one (FFF-4)

Synthesized according to general procedure 2, the crude residue was purified by biotage (Hexanes/EtOAc, 6:4) to afford FFF-4 as a colorless liquid (12 mg, 74%). ^1^H NMR (400 MHz, CDCl_3_) δ 7.25 – 7.06 (m, 7H), 4.95 (d, *J* = 17.2 Hz, 1H), 4.53 (dd, *J* = 34.2, 18.0 Hz, 2H), 3.70 (t, *J* = 5.1 Hz, 1H), 3.57 (d, *J* = 4.2 Hz, 1H), 3.44 (d, *J* = 5.9 Hz, 1H), 3.10 – 2.93 (m, 2H), 2.21 – 2.11 (m, 3H) 2.02 – 2.07 (m, 3H) 1.91 (dd, *J* = 7.1, 1.8 Hz, 3H), 1.69 (t, *J* = 7.5 Hz, 3H), 1.30 (d, *J* = 7.0 Hz, 3H), 1.26 (d, *J* = 7.0 Hz, 2.55H). *Note: rotameric mixtures observed*, LCMS *calcd for* C_18_H_22_N_3_O, 296.18 (M+H^+^), *found*: 296.2.

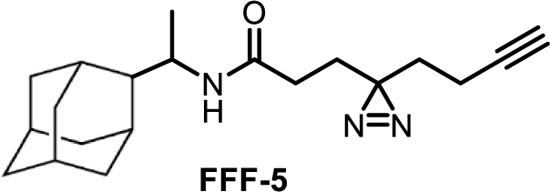

#### N-(1-((3r,5r,7r)-adamantan-1-yl)ethyl)-3-(3-(but-3-yn-1-yl)-3H-diazirin-3-yl)propanamide (FFF-5)

Synthesized according to general procedure 1, purified by biotage (Hexanes/EtOAc, 8:2) to afford FFF-5 as a white solid (11 mg, 67%). ^1^H NMR (400 MHz, CDCl_3_) δ 5.26 (d, J = 9.8 Hz, 1H), 3.71 (dq, J = 9.5, 6.8 Hz, 1H), 2.06 – 1.96 (m, 6H), 1.97 – 1.81 (m, 4H), 1.76 – 1.58 (m, 9H), 1.58 – 1.43 (m, 7H), 1.06 – 0.98 (m, 4H). ^13^C NMR (151 MHz, CDCl_3_) δ 170.76, 83.08, 69.52, 53.41, 38.73, 37.37, 36.07, 32.82, 31.03, 28.82, 28.63, 28.27, 14.89, 13.67.

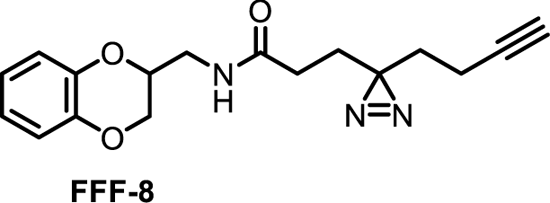

#### 3-(3-(but-3-yn-1-yl)-3H-diazirin-3-yl)-N-((2,3-dihydrobenzo[b][1,4]dioxin-2-yl)methyl)propenamide (FFF-8)

Synthesized according to general procedure 2, purified by biotage (Hexanes/EtOAc, 6:4) to afford FFF-8 as a white solid (16 mg, 74%). ^1^H NMR (400 MHz, CDCl_3_) δ 6.88 – 6.82 (m, 4H), 5.91 (s, 1H), 4.33 – 4.17 (m, 2H), 3.94 (dd, *J* = 11.9, 7.7 Hz, 1H), 3.67 (ddd, *J* = 14.4, 6.5, 3.9 Hz, 1H), 3.47 (dt, *J* = 14.3, 6.1 Hz, 1H), 2.03 – 1.94 (m, 5H), 1.85 (t, *J* = 7.4 Hz, 2H), 1.63 (t, *J* = 7.4 Hz, 2H). ^13^C NMR (151 MHz, CDCl_3_) δ 171.71, 143.06, 142.73, 121.68, 117.28, 117.21, 82.73, 71.91, 69.39, 65.68, 39.81, 32.33, 30.18, 28.25, 27.86, 13.26. LCMS *calcd for* C_17_H_20_N_3_O_3_, 314.1 (M+H^+^), *found*: 314.1.

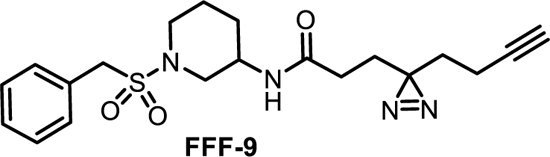

#### N-(1-(benzylsulfonyl)piperidin-3-yl)-3-(3-(but-3-yn-1-yl)-3H-diazirin-3-yl)propanamide (FFF-9)

Synthesized according to general procedure 2, the crude residue was purified by biotage (Hexanes/EtOAc, 8:2) to afford FFF-9 as a colorless liquid (14 mg, 76%). ^1^H NMR (400 MHz, CDCl_3_) δ 7.39 (q, *J* = 2.6 Hz, 5H), 5.81 (d, *J* = 7.7 Hz, 1H), 4.24 (s, 2H), 4.02 – 3.89 (m, 1H), 3.46 – 3.42 (m, 1H), 3.28 – 3.20 (m, 1H), 2.94 (dd, *J* = 12.7, 2.8 Hz, 1H), 2.75 – 1.98 (m, 1H), 2.07 – 1.98 (m, 3H), 1.98 – 1.88 (m, 2H), 1.86 – 1.77 (m, 3H), 1.68 – 1.60 (m, 3H). LCMS *calcd for* C_20_H_27_N_4_O_3_S, 403.18 (M+H^+^), *found*: 403.2.

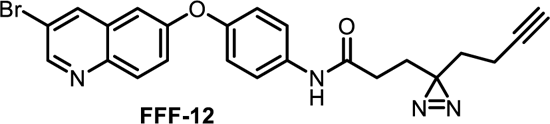

#### N-(4-((3-bromoquinolin-6-yl)oxy)phenyl)-3-(3-(but-3-yn-1-yl)-3H-diazirin-3-yl)propanamide (FFF-12)

Synthesized according to general procedure 2, purified by biotage (Hexanes/EtOAc, 6:4) to afford FFF-12 as a off white solid (15 mg, 74%). ^1^H NMR (600 MHz, CDCl_3_) δ 8.79 (d, *J* = 2.3 Hz, 1H), 8.14 (d, *J* = 2.3 Hz, 1H), 8.03 (d, *J* = 9.2 Hz, 1H), 7.58 – 7.52 (m, 2H), 7.47 (dd, *J* = 9.2, 2.7 Hz, 1H), 7.43 (s, 1H), 7.10 – 7.01 (m, 3H), 2.14 (dd, *J* = 8.2, 6.8 Hz, 2H), 2.05 (td, *J* = 7.4, 2.7 Hz, 2H), 2.00 (t, *J* = 2.6 Hz, 1H), 1.97 (dd, *J* = 8.2, 6.8 Hz, 2H), 1.70 (t, *J* = 7.4 Hz, 2H). ^13^C NMR (151 MHz, CDCl_3_) δ 169.46, 157.00, 152.28, 149.83, 143.05, 136.31, 134.22, 131.41, 130.11, 123.10, 121.93, 120.67, 117.97, 110.94, 82.72, 69.35, 32.48, 31.21, 28.20, 27.86, 13.33. LCMS *calcd for* C_23_H_20_BrN_4_O_2_, 463.1 (M+H^+^), *found*: 463.1.

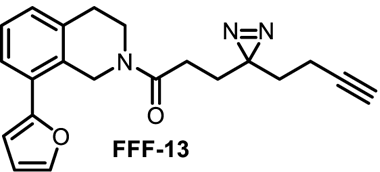

#### 3-(3-(but-3-yn-1-yl)-3H-diazirin-3-yl)-1-(8-(furan-2-yl)-3,4-dihydroisoquinolin-2(1H)-yl)propan-1-one (FFF-13)

Synthesized according to general procedure 1, purified by biotage (Hexanes/EtOAc, 5:5) to afford FFF-13 as a colorless liquid (15 mg, 65%). ^1^H NMR (400 MHz, CDCl_3_) δ 7.37 (d, J = 1.9 Hz, 1H), 7.31 (s, 1.25 H), 7.27 – 7.16 (m, 7.68 H), 6.82 – 6.80 (m, 1.32 H), 6.27 – 6.24 (m, 2.27 H), 6.08 – 5.99 (m, 2.29 H), 5.98 (s, 1H), 4.60 – 4.55 (m, 1H), 3.81 – 3.76 (m 1.33 H), 3.60 – 3.52 (m, 1.35 H)), 3.06 – 2.85 (m, 5H), 2.80 – 2.75 (m, 1.12 H), 2.65 – 2.49 (m, 2.81 H), 2.36 – 2.28 (m, 1.06), 2.23 – 2.07 (m, 2.43 H), 2.09 – 2.00 (m, 3.47 H), 1.98 – 1.95 (m, 2.53 H), 1.94 – 1.86 (m, 4.31H), 1.70 – 1.65 (s, 3.74H). Note: rotameric mixtures observed, LCMS *calcd for* C_21_H_22_N_3_O_2_, 348.2 (M+H^+^), *found*: 348.2.

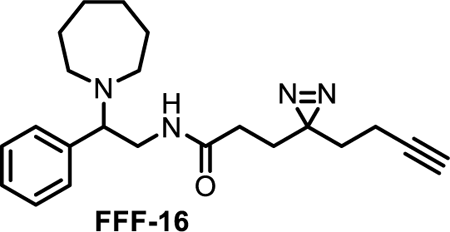

#### N-(2-(azepan-1-yl)-2-phenylethyl)-3-(3-(but-3-yn-1-yl)-3H-diazirin-3-yl)propenamide (FFF-16)

Synthesized according to general procedure 1, purified by biotage (Hexanes/EtOAc, 4:6) to afford FFF-16 as a colorless liquid (9 mg, 74%). ^1^H NMR (400 MHz, CDCl_3_) δ 7.38 – 7.28 (m, 3H), 7.23 (dd, *J* = 7.9, 1.7 Hz, 2H), 6.21 (s, 1H), 3.78 (t, *J* = 7.7 Hz, 1H), 3.55 – 3.53 (m, 2H), 2.80 – 2.66 (m, 2H), 2.59 – 2.53 (m, 2H), 2.07 – 1.91 (m, 5H), 1.90 – 1.78 (m, 2H), 1.69 – 1.53 (m, 10H). LCMS *calcd for* C_22_H_31_FN_4_O, 367.2 (M+H^+^), *found*: 367.2.

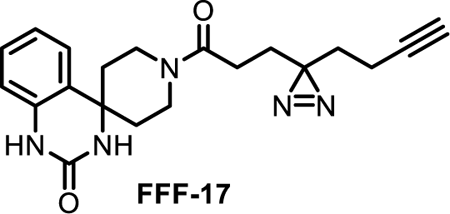

#### N-(2-(3-(but-3-yn-1-yl)-3H-diazirin-3-yl)ethyl)-2’-oxo-2’,3’-dihydro-1’H-spiro[piperidine-4,4’-quinazoline]-1-carboxamide (FFF-17)

Synthesized according to general procedure 2, purified by biotage (Hexanes/EtOAc, 8:2) to afford FFF-17 as an off white solid (12 mg, 67 %). ^1^H NMR (400 MHz, CDCl_3_) δ 8.91 (d, *J* = 2.0 Hz, 1H), 7.18 – 7.09 (m, 2H), 7.05 (d, *J* = 2.1 Hz, 1H), 6.98 (dd, *J* = 7.6, 1.2 Hz, 1H), 6.78 (dd, *J* = 7.9, 1.2 Hz, 1H), 4.68 – 4.54 (m, 1H), 3.75 – 3.61 (m, 1H), 3.53 – 3.46 (m, 1H), 3.02 – 2.98 (m, 1H), 2.12 – 1.93 (m, 8H), 1.93 – 1.79 (m, 4H), 1.72 – 1.62 (m, 2H). ^13^C NMR (101 MHz, CDCl_3_) δ 169.53, 154.96, 135.65, 128.61, 125.12, 123.84, 122.87, 114.80, 82.82, 69.27, 55.00, 40.52, 38.13, 36.78, 32.55, 28.03, 27.96, 26.86, 13.34. LCMS *calcd for* C_20_H_25_N_6_O_2_, 381.2 (M+H^+^), *found*: 381.2.

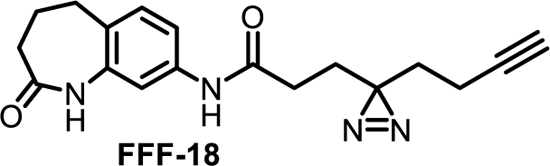

#### 3-(3-(but-3-yn-1-yl)-3H-diazirin-3-yl)-N-(2-oxo-2,3,4,5-tetrahydro-1H-benzo[b]azepin-8-yl)propenamide (FFF-18)

Synthesized according to general procedure 1, purified by biotage (Hexanes/EtOAc, 6:4) to afford FFF-18 as a white solid (9 mg, 64%).^1^H NMR (400 MHz, CDCl_3_) δ 8.27 (s, 1H), 8.00 (s, 1H), 7.33 – 7.23 (m, 2H), 7.13 (d, *J* = 8.1 Hz, 1H), 2.73 (t, *J* = 7.2 Hz, 2H), 2.33 (t, *J* = 7.3 Hz, 2H), 2.26 – 2.09 (m, 4H), 2.06 – 1.98 (m, 3H), 1.92 (dd, *J* = 8.4, 6.7 Hz, 2H), 1.67 (t, *J* = 7.3 Hz, 2H). ^13^C NMR (101 MHz, CDCl_3_) δ 175.59, 169.87, 138.14, 137.26, 130.21, 130.10, 117.02, 113.51, 82.74, 69.36, 32.89, 32.40, 31.20, 29.77, 28.50, 28.17, 27.92, 13.32. LCMS *calcd for* C_18_H_21_N_4_O_2_, 325.2 (M+H^+^), *found*: 325.2.

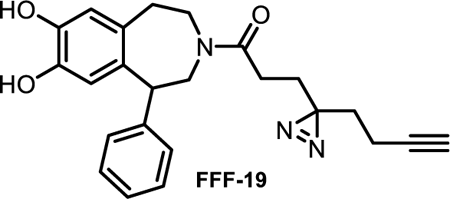

#### 3-(3-(but-3-yn-1-yl)-3H-diazirin-3-yl)-1-(7,8-dihydroxy-1-phenyl-1,2,4,5-tetrahydro-3H-benzo[d]azepin-3-yl)propan-1-one (FFF-19)

Synthesized according to general procedure 1, purified by biotage (Hexanes/EtOAc, 6:4) to afford FFF-19 as a white solid (8 mg, 65%).^1^H NMR (400 MHz, CDCl_3_) δ 7.39 – 7.16 (m, 5H), 7.13 – 7.03 (m, 1H), 6.99 – 6.85 (m, 2H), 6.72 (s, 1H), 6.39 (s, 1H), 4.48 – 4.43 (m, 1H), 4.37 – 4.32 (s, 1H), 4.03 – 3.88 (m, 1H), 3.70 – 3.61 (m, 1H), 3.59 – 3.49 (m, 2H), 3.38 – 3.19 (m, 2H), 2.84 – 2.67 (m, 1H), 1.96 – 1.81 (m, 7H), 1.62 – 1.52 (m, 2H), 1.52 – 1.36 (m, 4H). Note: rotameric mixtures observed, LCMS *calcd for* C_24_H_26_N_3_O_3_, 404.2 (M+H^+^), *found*: 404.2.

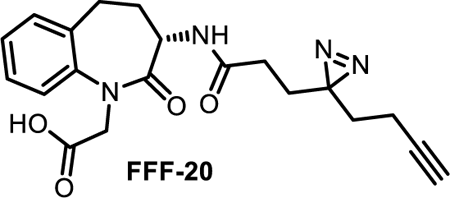

#### (S)-2-(3-(3-(3-(but-3-yn-1-yl)-3H-diazirin-3-yl)propanamido)-2-oxo-2,3,4,5-tetrahydro-1H-benzo[b]azepin-1-yl)acetic acid (FFF-20)

Synthesized according to general procedure 1, purified by biotage (Hexanes/EtOAc, 8:2) to afford FFF-20 as a brown solid (11 mg, 64 %). ^1^H NMR (400 MHz, CDCl_3_) δ 7.31 (ddd, *J* = 7.8, 6.6, 2.4 Hz, 1H), 7.26 – 7.19 (m, 2H), 7.14 (dd, *J* = 7.8, 1.3 Hz, 1H), 6.64 (d, *J* = 7.3 Hz, 1H), 4.65 (d, *J* = 17.4 Hz, 1H), 4.58 – 4.51 (m, 1H), 4.47 (d, *J* = 17.5 Hz, 1H), 3.31 – 3.19 (m, 1H), 2.73 – 2.56 (m, 2H), 2.10 (s, 1H), 2.03 – 1.89 (m, 5H), 1.82 – 1.68 (m, 2H), 1.66 – 1.53 (m, 2H). LCMS *calcd for* C_20_H_23_N_4_O_4_, 383.17 (M+H^+^), *found*: 383.2.

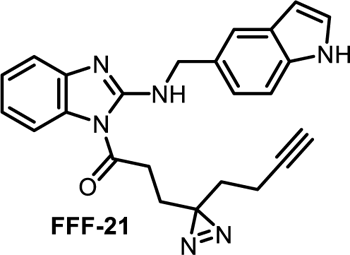

#### 1-(2-(((1H-indol-5-yl)methyl)amino)-1H-benzo[d]imidazol-1-yl)-3-(3-(but-3-yn-1-yl)-3H-diazirin-3-yl)propan-1-one (FFF-21)

Synthesized according to scheme 1, general procedure 1 and following general procedure 2, purified on Biotage^®^ with hexane: ethyl acetate (6:4) to afford AJ2-15 as colorless liquid (12 mg, 42 %) ^1^H NMR (400 MHz, CDCl_3_) δ 8.36 (s, 1H), 8.06 (t, *J* = 5.2 Hz, 1H), 7.67 (s, 1H), 7.45 (d, *J* = 7.8 Hz, 1H), 7.35 (d, *J* = 8.4 Hz, 1H), 7.30 – 7.20 (m, 4H), 7.06 (t, *J* = 7.8 Hz, 1H), 6.52 (s, 1H), 4.84 (d, *J* = 5.1 Hz, 2H), 2.76 (t, *J* = 7.4 Hz, 2H), 2.10 – 1.98 (m, 5H), 1.73 (d, *J* = 7.3 Hz, 2H). ^13^C NMR (151 MHz, CDCl_3_) δ 174.13, 149.86, 140.04, 135.33, 132.21, 128.20, 127.64, 124.96, 122.19, 122.11, 120.66, 118.41, 111.64, 110.60, 102.69, 82.67, 69.19, 50.27, 32.34, 29.72, 29.48, 27.57, 13.22. HRMS (ESI-TOF) calcd for C_24_H_23_N_6_O, 411.1928 (M+H^+^), found 411.1938.

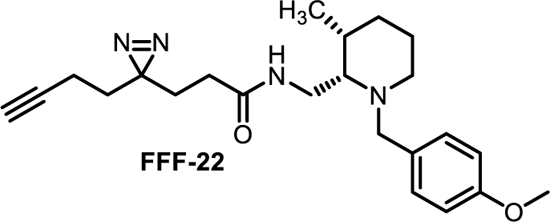

#### 3-(3-(but-3-yn-1-yl)-3H-diazirin-3-yl)-N-(((2S,3R)-1-(4-methoxybenzyl)-3-methylpiperidin-2-yl)methyl)propanamide (FFF-22)

Synthesized according to general procedure 2. The residue obtained was purified by biotage (Hexanes/EtOAc, 8:2) to afford FFF-22 as a colorless liquid (13 mg, 62%). ^1^H NMR (400 MHz, CDCl_3_) δ 7.20 (d, J = 8.2 Hz, 2H), 6.92 – 6.86 (m, 2H), 6.24 – 6.11 (m, 1H), 3.82 (d, J = 1.4 Hz, 3H), 3.76 (s, 2H), 3.43 – 3.31 (m, 1H), 3.10 (ddd, J = 13.2, 11.0, 1.8 Hz, 1H), 2.79 – 2.69 (m, 1H), 2.67 – 2.50 (m, 2H), 2.09 (tt, J = 7.8, 4.2 Hz, 1H), 2.06 – 1.97 (m, 3H), 1.92 – 1.72 (m, 5H), 1.68 – 1.56 (m, 3H), 1.37 – 1.27 (m, 2H), 0.84 (dd, J = 7.0, 1.1 Hz, 3H). ^13^C NMR (126 MHz, CDCl_3_) δ 170.96, 129.86, 113.91, 82.72, 69.19, 56.71, 55.30, 55.29, 45.11, 34.31, 32.36, 30.53, 29.70, 28.57, 27.90, 27.01, 20.49, 18.17, 13.31. HRMS (ESI-TOF) *calcd for* C_23_H_33_N_4_O_2_, 397.5425 (M+H^+^), *found* 397.2549.

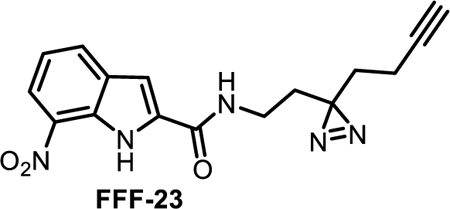

#### N-(2-(3-(but-3-yn-1-yl)-3H-diazirin-3-yl)ethyl)-7-nitro-1H-indole-2-carboxamide (FFF-23)

Synthesized according to general procedure 2, purified by biotage (Hexanes/EtOAc, 6:4) to afford FFF-23 as a white solid (10 mg, 68%). ^1^H NMR (400 MHz, CDCl_3_) δ 10.51 (s, 1H), 8.24 (dd, *J* = 8.1, 1.0 Hz, 1H), 8.01 (d, *J* = 7.8 Hz, 1H), 7.32 – 7.19 (m, 1H), 6.58 (t, *J* = 6.0 Hz, 1H), 3.39 (q, *J* = 6.4 Hz, 2H), 2.13 – 1.99 (m, 3H), 1.88 (t, *J* = 6.6 Hz, 3H), 1.71 (t, *J* = 7.0 Hz, 2H). ^13^C NMR (101 MHz, CDCl_3_) δ 160.41, 133.44, 133.14, 131.24, 130.22, 129.29, 121.60, 120.11, 103.31, 82.80, 69.60, 34.76, 32.55, 32.09, 26.90, 13.20. LCMS *calcd for* C_16_H_15_N_5_O_3_, 326.1 (M+H^+^), *found*: 326.1.

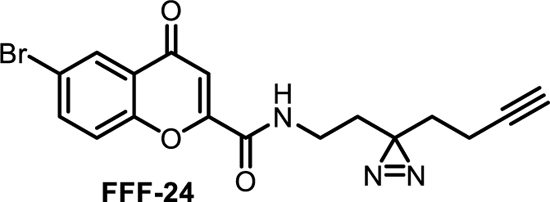

#### 6-bromo-N-(2-(3-(but-3-yn-1-yl)-3H-diazirin-3-yl)ethyl)-4-oxo-4H-chromene-2-carboxamide (FFF-24)

Synthesized according to general procedure 1. The residue obtained was purified by biotage (Hexanes/EtOAc, 7:3) to afford FFF-36 as a white solid (12 mg, 71%). ^1^H NMR (400 MHz, CDCl_3_) δ 8.36 (d, *J* = 2.4 Hz, 1H), 7.84 (dd, *J* = 8.9, 2.5 Hz, 1H), 7.47 (d, *J* = 8.9 Hz, 1H), 7.17 (s, 1H), 6.99 (s, 1H), 3.37 (q, *J* = 6.5 Hz, 2H), 2.12 – 1.99 (m, 3H), 1.88 (t, *J* = 6.7 Hz, 2H), 1.71 (t, *J* = 7.1 Hz, 2H). LCMS *calcd for* C_17_H_14_BrN_3_O_3_, 388.02 (M+H^+^), *found*: 388.02.

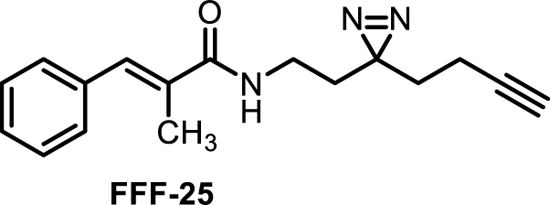

#### (E)-N-(2-(3-(but-3-yn-1-yl)-3H-diazirin-3-yl)ethyl)-2-methyl-3-phenylacrylamide (FFF-25)

Synthesized according to general procedure 2, the crude residue was purified by biotage (Hexanes/EtOAc, 7:3) to afford FFF-25 as a colorless liquid (15 mg, 71%). ^1^H NMR (400 MHz, CDCl_3_) δ 7.43 – 7.28 (m, 6H), 6.01 (s, 1H), 3.25 (td, *J* = 6.6, 5.9 Hz, 2H), 2.12 (d, *J* = 1.5 Hz, 3H), 2.10 – 1.98 (m, 3H), 1.80 (t, *J* = 6.6 Hz, 2H), 1.69 (t, *J* = 7.2 Hz, 2H). LCMS *calcd for* C_17_H_19_N_3_O, 281.16 (M+H^+^), *found*: 281.2.

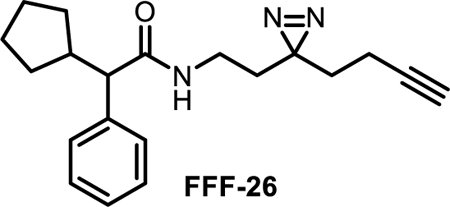

#### N-(2-(3-(but-3-yn-1-yl)-3H-diazirin-3-yl)ethyl)-2-cyclopentyl-2-phenylacetamide (FFF-26)

Synthesized according to general procedure 2, the crude residue was purified by biotage (Hexanes/EtOAc, 9:1) to afford FFF-26 as a colorless liquid (13 mg, 68%). ^1^H NMR (600 MHz, CDCl_3_) δ 7.39 – 7.34 (m, 2H), 7.34 – 7.30 (m, 2H), 7.28 – 7.23 (m, 1H), 5.65 (s, 1H), 3.13 – 3.03 (m, 2H), 3.02 – 3.00 (m, 1H), 2.63 – 2.59 (m, 1H), 2.03 – 1.95 (m, 2H), 1.93 – 1.89 (m, 2H), 1.70 – 1.58 (m, 5H), 1.56 – 1.51 (m, 2H), 1.49 – 1.42 (m, 1H), 1.28 (d, *J* = 8.5 Hz, 2H), 1.07 – 0.95 (m, 1H). ^13^C NMR (151 MHz, CDCl_3_) δ 173.49, 139.98, 128.59, 128.00, 127.12, 82.73, 69.35, 59.91, 43.17, 34.26, 32.43, 32.07, 31.74, 30.95, 26.77, 25.19, 24.84, 13.16. LCMS *calcd for* C_20_H_26_N_3_O, 324.20 (M+H^+^), *found*: 324.2.

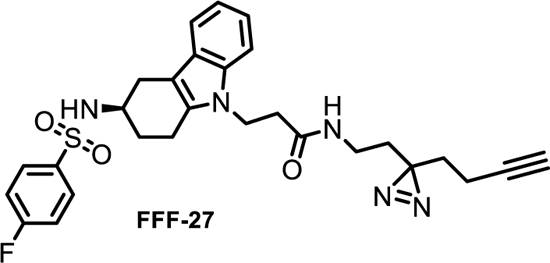

#### (R)-N-((3-(but-3-yn-1-yl)-3H-diazirin-3-yl)methyl)-3-(3-((4-fluorophenyl)sulfonamido)-1,2,3,4-tetrahydro-9H-carbazol-9-yl)propanamide (FFF-27)

Synthesized according to general procedure 2, purified by biotage (Hexanes/EtOAc, 6:4) to afford FFF-27 as a colorless liquid (15 mg, 78%). ^1^H NMR (400 MHz, CDCl_3_) δ 7.86 – 7.79 (m, 2H), 7.30 – 7.22 (m, 2H), 7.16 – 7.08 (m, 3H), 7.06 – 7.00 (m, 1H), 5.60 (t, *J* = 5.8 Hz, 1H), 5.37 (d, *J* = 8.4 Hz, 1H), 4.38 (dt, *J* = 14.8, 6.9 Hz, 1H), 4.25 (dt, *J* = 14.8, 6.6 Hz, 1H), 3.80 (d, *J* = 8.3, Hz, 1H), 3.04 – 2.87 (m, 2H), 2.90 – 2.78 (m, 2H), 2.78 – 2.67 (m, 1H), 2.57 – 2.43 (m, 3H), 2.00 – 1.93 (m, 3H), 1.87 (td, *J* = 7.1, 2.5 Hz, 2H), 1.51 – 1.35 (m, 4H), LCMS *calcd for* C_27_H_29_FN_5_O_3_S, 521.2 (M+H^+^), *found*: 521.2.

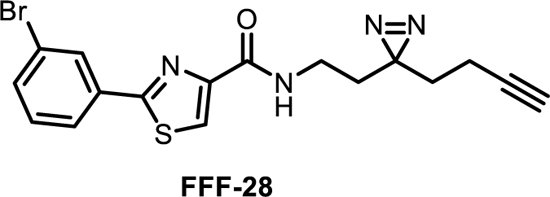

#### 2-(3-bromophenyl)-N-(2-(3-(but-3-yn-1-yl)-3H-diazirin-3-yl)ethyl)thiazole-4-carboxamide (FFF-28)

Synthesized according to general procedure 2, purified by biotage (Hexanes/EtOAc, 7:3) to afford FFF-28 as an off white solid (12 mg, 71%). ^1^H NMR (600 MHz, CDCl_3_) δ 8.13 (t, *J* = 1.8 Hz, 1H), 8.10 (s, 1H), 7.84 (dd, *J* = 7.8, 1.7, Hz, 1H), 7.57 (dd, *J* = 8.0, 2.0, Hz, 1H), 7.52 (d, *J* = 6.1 Hz, 1H), 7.32 (t, *J* = 7.9 Hz, 1H), 3.35 (td, *J* = 7.0, 6.1 Hz, 2H), 2.04 (td, *J* = 7.4, 2.6 Hz, 2H), 2.00 (t, *J* = 2.6 Hz, 1H), 1.82 (t, *J* = 7.0 Hz, 2H), 1.70 (t, *J* = 7.3 Hz, 2H). ^13^C NMR (151 MHz, CDCl_3_) δ 166.34, 160.96, 150.75, 134.59, 133.50, 130.61, 129.41, 125.27, 123.55, 123.23, 82.63, 69.45, 34.32, 32.85, 26.86, 13.29. LCMS *calcd for* C_17_H_16_BrN_4_OS, 403.01 (M+H^+^), *found*: 403.1.

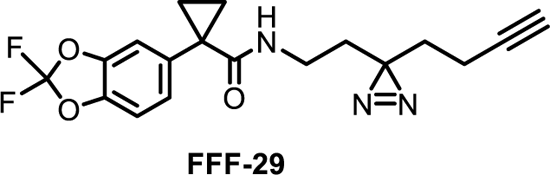

#### N-(2-(3-(but-3-yn-1-yl)-3H-diazirin-3-yl)ethyl)-1-(2,2-difluorobenzo[d][1,3]dioxol-5-yl)cyclopropane-1-carboxamide (FFF-29)

Synthesized according to general procedure 2, purified by biotage (Hexanes/EtOAc, 6:4) to afford FFF-29 as a white solid (15 mg, 78%).^1^H NMR (500 MHz, CDCl_3_) δ 7.23 – 7.16 (m, 2H), 7.09 (d, J = 8.1 Hz, 1H), 5.39 (d, J = 6.2 Hz, 1H), 3.08 (q, J = 6.4 Hz, 2H), 2.00 – 1.93 (m, 3H), 1.60 (m, 6H), 1.05 (q, J = 3.8 Hz, 2H). ^13^C NMR (126 MHz, CDCl_3_) δ 173.24, 143.12, 143.51, 135.90, 133.80 (t, *J* = 256.0 Hz), 126.63, 112.50, 109.95, 82.65, 69.34, 35.26, 32.63, 32.13, 30.46, 26.84, 16.14, 16.12, 13.17. HRMS (ESI-TOF) *calcd for* C_18_H_18_F_3_N_3_O_3_, 362.1311 (M+H^+^), *found* 362.1312.

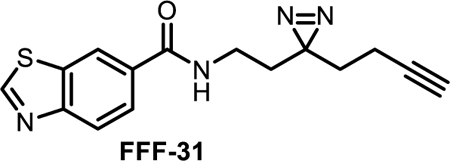

#### N-(2-(3-(but-3-yn-1-yl)-3H-diazirin-3-yl)ethyl)benzo[d]thiazole-6-carboxamide (FFF-31)

Synthesized according to general procedure 2, purified by biotage (Hexanes/EtOAc, 6:4) to afford FFF-31 as a white solid (16 mg, 76%). ^1^H NMR (400 MHz, CDCl_3_) δ 9.07 (s, 1H), 8.42 (dd, *J* = 1.8, 0.6 Hz, 1H), 8.09 (dd, *J* = 8.5, 0.6 Hz, 1H), 7.86 (dd, *J* = 8.5, 1.8 Hz, 1H), 6.75 (t, *J* = 5.9 Hz, 1H), 3.31 (td, *J* = 6.7, 5.8 Hz, 2H), 2.06 – 1.94 (m, 3H), 1.81 (t, *J* = 6.7 Hz, 2H), 1.65 (t, *J* = 7.1 Hz, 2H). ^13^C NMR (101 MHz, CDCl_3_) δ 167.01, 156.67, 155.04, 134.12, 131.80, 124.69, 123.52, 121.69, 82.73, 69.54, 35.18, 32.42, 32.06, 26.98, 13.22. LCMS *calcd for* C_15_H_15_N_4_OS, 298.09 (M+H^+^), *found*: 298.1.

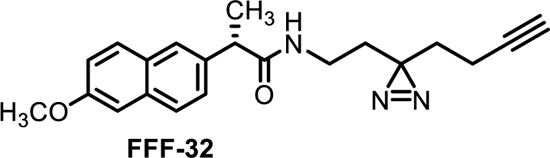

#### (S)-N-(2-(3-(but-3-yn-1-yl)-3H-diazirin-3-yl)ethyl)-2-(6-methoxynaphthalen-2-yl)propanamide (FFF-32)

Synthesized according to general procedure 1. The residue obtained was purified by biotage (Hexanes/EtOAc, 7:3) to afford FFF-32 as a white solid (14 mg, 76%). ^1^H NMR (400 MHz, CDCl_3_) δ 7.78 – 7.65 (m, 3H), 7.39 (dd, *J* = 8.5, 1.9 Hz, 1H), 7.21 – 7.09 (m, 2H), 5.46 (s, 1H), 3.92 (s, 3H), 3.70 (q, *J* = 7.2 Hz, 1H), 3.05 (t, *J* = 6.7 Hz, 2H), 1.93 – 1.80 (m, 3H), 1.61 (d, *J* = 7.2 Hz, 3H), 1.59 – 1.55 (m, 2H), 1.54 – 1.46 (m, 2H). LCMS *calcd for* C_21_H_22_N_3_O_2_, 350.2 (M+H^+^), *found*: 350.2.

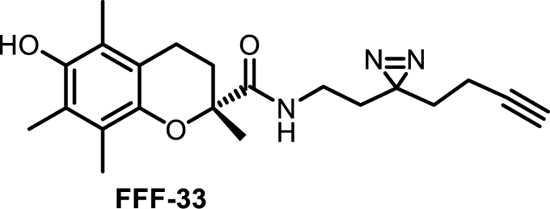

#### (R)-N-(2-(3-(but-3-yn-1-yl)-3H-diazirin-3-yl)ethyl)-6-hydroxy-2,5,7,8-tetramethylchromane-2-carboxamide (FFF-33)

Synthesized according to general procedure 1. The residue obtained was purified by biotage (Hexanes/EtOAc, 7:3) to afford FFF-33 as a white solid (10 mg, 68%). ^1^H NMR (400 MHz, CDCl_3_) δ 6.67 – 6.52 (m, 1H), 4.49 (d, *J* = 7.3 Hz, 1H), 3.07 (qdt, *J* = 14.0, 7.0, 6.1 Hz, 2H), 2.71 – 2.49 (m, 2H), 2.37 (dt, *J* = 13.5, 5.9 Hz, 1H), 2.24 (s, 3H), 2.18 (s, 3H), 2.10 (s, 3H), 1.95 (t, *J* = 2.6 Hz, 1H), 1.92 – 1.83 (m, 3H), 1.63 – 1.83 (m, 2H), 1.52 (s, 3H), 1.50 – 1.46 (m, 1H). LCMS *calcd for* C_21_H_27_N_3_O_3_, 370.2 (M+H^+^), *found*: 370.2.

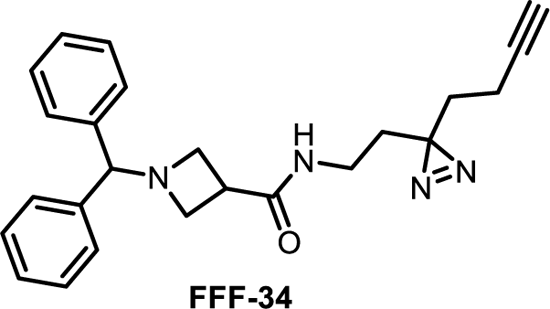

#### 1-benzhydryl-N-(2-(3-(but-3-yn-1-yl)-3H-diazirin-3-yl)ethyl)azetidine-3-carboxamide (FFF-34)

Synthesized according to general procedure 2, purified by biotage (Hexanes/EtOAc, 6:4) to afford FFF-27 as a colorless liquid (14 mg, 78%). ^1^H NMR (400 MHz, CDCl_3_) δ 7.48 – 7.38 (m, 4H), 7.32 – 7.26 (m, 4H), 7.24 – 7.16 (m, 2H), 6.48 (s, 1H), 4.45 (s, 1H), 3.46 – 3.29 (m, 4H), 3.15 (q, *J* = 6.3 Hz, 2H), 3.07 (q, *J* = 6.5 Hz, 1H), 2.03 (td, *J* = 7.3, 2.8 Hz, 2H), 1.98 (t, *J* = 2.6 Hz, 1H), 1.75 (t, *J* = 6.6 Hz, 2H), 1.66 (t, *J* = 7.3 Hz, 2H). ^13^C NMR (101 MHz, CDCl_3_) δ 173.40, 141.51, 128.55, 127.47, 127.29, 82.65, 77.63, 77.28, 69.49, 56.41, 35.83, 34.18, 32.42, 32.27, 26.91, 13.25. LCMS *calcd for* C_24_H_27_N_4_O, 387.2 (M+H^+^), *found*: 387.2.

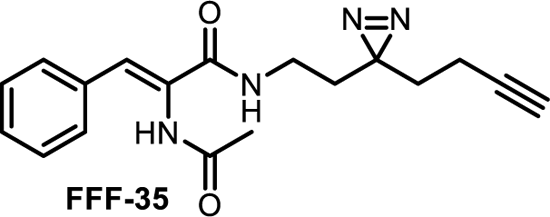

#### (Z)-2-acetamido-N-(2-(3-(but-3-yn-1-yl)-3H-diazirin-3-yl)ethyl)-3-phenylacrylamide (FFF-35)

Synthesized according to general procedure 2, purified by biotage (Hexanes/EtOAc, 8:2) to afford FFF-35 as an off white solid (12 mg, 71 %). ^1^H NMR (600 MHz, CD_3_OD_SPE) δ 7.52 (dd, *J* = 7.7, 1.7 Hz, 2H), 7.45 – 7.38 (m, 2H), 7.38 – 7.32 (m, 1H), 7.14 (s, 1H), 3.22 (t, *J* = 7.3 Hz, 2H), 2.31 (d, *J* = 2.7, 0.9 Hz, 1H), 2.12 (d, *J* = 0.9 Hz, 3H), 2.07 (tdd, *J* = 7.4, 2.7, 0.9 Hz, 2H), 1.74 – 1.65 (m, 4H). ^13^C NMR (151 MHz, CD_3_OD_SPE) δ 171.94, 166.70, 133.86, 129.17, 129.14, 129.06, 128.70, 128.37, 82.32, 69.06, 48.09, 47.94, 47.80, 47.66, 47.52, 47.38, 47.24, 34.58, 32.07, 31.84, 26.56, 21.36. LCMS *calcd for* C_18_H_21_N_4_O_2_, 325.2 (M+H^+^), *found*: 325.2.

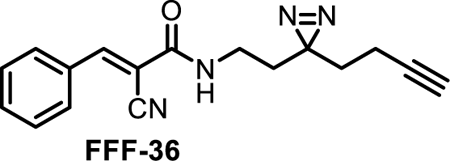

#### (E)-N-(2-(3-(but-3-yn-1-yl)-3H-diazirin-3-yl)ethyl)-2-cyano-3-phenylacrylamide (FFF-36)

Synthesized according to general procedure 2, purified by biotage (Hexanes/EtOAc, 6:4) to afford FFF-36 as an off white solid (14 mg, 74 %). ^1^H NMR (400 MHz, CDCl_3_) δ 8.33 (s, 1H), 7.99 – 7.86 (m, 2H), 7.59 – 7.44 (m, 3H), 6.51 (s, 1H), 3.31 (td, *J* = 6.9, 5.9 Hz, 2H), 2.10 – 1.99 (m, 3H), 1.81 (t, *J* = 6.9 Hz, 2H), 1.74 – 1.64 (m, 2H). ^13^C NMR (101 MHz, CDCl_3_) δ 160.34, 153.19, 132.90, 131.75, 130.70, 129.28, 116.91, 103.76, 82.55, 69.61, 35.50, 32.48, 32.14, 26.66, 13.25. LCMS *calcd for* C_17_H_17_N_4_O, 293.2 (M+H^+^), *found*: 293.2.

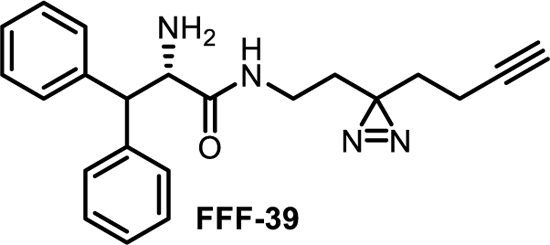

#### (S)-2-amino-N-(2-(3-(but-3-yn-1-yl)-3H-diazirin-3-yl)ethyl)-3,3-diphenylpropanamide (FFF-39)

Synthesized according to general procedure 2, purified by biotage (Hexanes/EtOAc, 8:2) to afford FFF-39 as a white solid (8 mg, 58 %). ^1^H NMR (400 MHz, CDCl_3_) δ 7.35 – 7.26 (m, 8H), 7.26 – 7.19 (m, 2H), 6.70 (s, 1H), 4.59 (d, *J* = 6.5 Hz, 1H), 4.08 (d, *J* = 6.5 Hz, 1H), 3.05 – 2.88 (m, 2H), 2.01 – 1.90 (m, 3H), 1.53 (d, *J* = 7.4 Hz, 2H), 1.42 (q, *J* = 6.9 Hz, 2H). LCMS *calcd for* C_22_H_25_N_4_O, 361.20 (M+H^+^), *found*: 361.2.

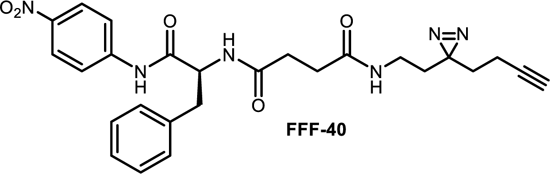

#### (S)-N1-(2-(3-(but-3-yn-1-yl)-3H-diazirin-3-yl)ethyl)-N4-(1-((4-nitrophenyl)amino)-1-oxo-3-phenylpropan-2-yl)succinamide (FFF-40)

Synthesized according to general procedure 2, purified by biotage (Hexanes/EtOAc, 4:6) to afford FFF-40 as a white solid (8 mg, 65%). ^1^H NMR (400 MHz, CDCl_3_) δ 9.22 (s, 1H), 8.24 – 8.14 (m, 2H), 7.97 – 7.88 (m, 2H), 7.37 – 7.28 (m, 3H), 7.23 (dd, *J* = 8.0, 1.6 Hz, 3H), 6.17 (d, *J* = 8.2 Hz, 1H), 5.72 (t, *J* = 5.9 Hz, 1H), 4.92 (d, *J* = 8.3, Hz, 1H), 3.38 – 3.18 (m, 2H), 3.08 – 2.96 (m, 2H), 2.91 – 2.76 (m, 1H), 2.42 – 2.37 (m, 2H), 2.05 – 1.94 (m, 3H), 1.71 – 1.67 (m, 2H), 1.63 – 1.55 (m, 3H). ^13^C NMR (101 MHz, CDCl_3_) δ 172.89, 172.39, 170.31, 144.10, 143.46, 136.27, 129.19, 128.90, 127.27, 124.75, 119.77, 82.66, 69.65, 54.86, 36.96, 34.60, 32.27, 32.05, 31.49, 31.11, 26.72, 13.17. LCMS *calcd for* C_26_H_29_N_6_O_5_, 505.2 (M+H^+^), *found*: 505.2.

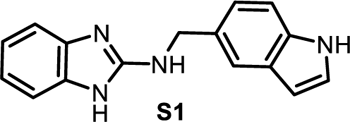

#### N-((1H-indol-5-yl)methyl)-1H-benzo[d]imidazol-2-amine

Synthesized according to scheme 1 purified on Biotage^®^ with 0 to 7 % methanol gradient in dichloromethane to afford S1 as an light brown solid (450 mg, 62 %). ^1^H NMR (400 MHz, DMSO) δ 11.02 (s, 1H), 7.53 (d, J = 1.6 Hz, 1H), 7.34 (d, J = 8.3 Hz, 1H), 7.30 (t, J = 2.7 Hz, 1H), 7.15 – 7.09 (m, 3H), 7.05 (t, J = 6.1 Hz, 1H), 6.85 (dd, J = 5.8, 3.2 Hz, 2H), 6.37 (t, J = 2.6 Hz, 1H), 4.56 (d, J = 5.9 Hz, 2H). HRMS (ESI-TOF) calcd for C_16_H_15_N_4_, 263.1291 (M+H^+^), found 263.1287.

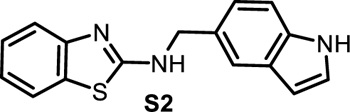

#### N-((1H-indol-5-yl)methyl)benzo[d]thiazol-2-amine (S2)

To a solution of 2-Aminobenzothiazole (1.0 eq.) and 5-Formylindole (1.0 eq.) in dry ethanol, sodium triacetoxyborohydride (2.0 eq.) and acetic acid (2.0 eq.) was added at 0°C to the solution and resulting mixture was stirred for 14 h at room temperature. After completion (monitored by TLC) the solvent was removed by rotary evaporation, crude mixture was diluted with water and washed with saturated aqueous NaHCO_3_ solution extracted in ethyl acetate, the combined extracts was dried over Na_2_SO_4,_ filtered and concentrated in vacuum, purified by flash column chromatography on Biotage^®^ isolera one with 5-70 % ethyl acetate gradient in hexane to obtain corresponding N-((1H-indol-5-yl)methyl)benzo[d]thiazol-2-amine (**S2**) as a light brown solid (12 mg, 22 %). ^1^H NMR (400 MHz, CD_3_OD_SPE) δ 7.58 – 7.53 (m, 2H), 7.43 (dd, J = 8.1, 1.2, Hz, 1H), 7.36 (dd, J = 8.3, 0.9 Hz, 1H), 7.27 – 7.19 (m, 2H), 7.14 (dd, J = 8.4, 1.7 Hz, 1H), 7.03 (td, J = 7.6, 1.2 Hz, 1H), 6.41 (d, J = 3.1, Hz, 1H), 4.66 (s, 2H). HRMS (ESI-TOF) calcd for C_16_H_14_N_3_S, 280.0903 (M+H^+^), found 280.0901.

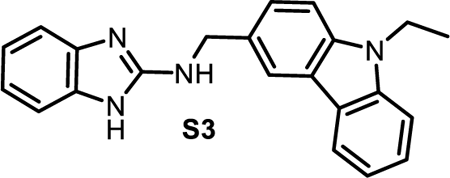

#### N-((9-ethyl-9H-carbazol-3-yl)methyl)-1H-benzo[d]imidazol-2-amine (S3)

Synthesized according to scheme 1 purified on Biotage^®^ with 0 to 6 % methanol gradient in dichloromethane to afford S3 as an light yellow solid (210 mg, 72 %). ^1^H NMR (400 MHz, DMSO) δ 11.00 (s, 1H), 8.15 (dd, J = 1.7, 0.7 Hz, 1H), 8.10 (dd, J = 7.8, 1.0 Hz, 1H), 7.62 – 7.55 (m, 2H), 7.51 (dd, J = 8.4, 1.7 Hz, 1H), 7.43 (dd, J = 8.3, 1.2 Hz, 1H), 7.27 (s, 1H), 7.21 – 7.11 (m, 3H), 6.88 (dd, J = 5.7, 3.2 Hz, 2H), 4.67 (d, J = 5.4 Hz, 2H), 4.42 (q, J = 7.1 Hz, 2H), 1.28 (t, J = 7.1 Hz, 3H). HRMS (ESI-TOF) calcd for C_22_H_20_N_4_, 341.1761 (M+H^+^), found 341.1762.

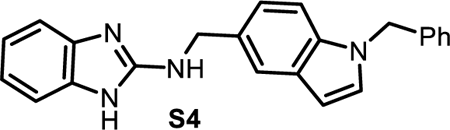

#### N-((1-benzyl-1H-indol-5-yl)methyl)-1H-benzo[d]imidazol-2-amine (S4)

Synthesized according to scheme 1 purified on Biotage^®^ with 0 to 6 % methanol gradient in dichloromethane to afford S4 as an light yellow solid (85mg, 62 %). ^1^H NMR (600 MHz, DMSO) δ 7.58 (s, 1H), 7.48 – 7.46 (s, 1H), 7.39 (d, *J* = 8.4 Hz, 1H), 7.30 – 7.24 (m, 2H), 7.24 – 7.18 (m, 2H), 7.18 – 7.14 MHz, DMSO) δ 155.74, 138.80, 135.49, 131.06, 129.91, 128.97, 128.72, 127.74, 127.35, 121.76, 119.77, 119.59, 112.05, 110.47, 101.37, 49.63, 46.68, 40.40, 40.22, 40.12, 39.94, 39.84, 39.71, 39.57. HRMS (ESI-TOF) calcd for C_23_H_21_N_4_, 353.1761 (M+H^+^), found 353.1750.

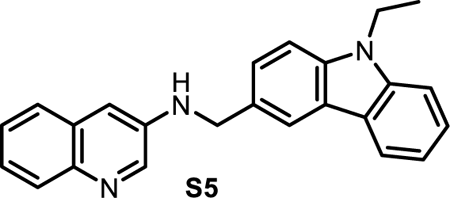

#### N-((9-ethyl-9H-carbazol-3-yl)methyl)quinolin-3-amine (S5)

Synthesized according to scheme 1 purified on Biotage^®^ with 0 to 6 % methanol gradient in dichloromethane to afford S5 as an brown solid (90 mg, 58 %). ^1^H NMR (400 MHz, CDCl_3_) δ 8.54 (dd, J = 2.8, 1.0 Hz, 1H), 8.20 – 8.08 (m, 2H), 8.00 (dt, J = 6.2, 3.1 Hz, 1H), 7.62 (dd, J = 6.3, 3.3 Hz, 1H), 7.52 (dd, J = 7.0, 1.4 Hz, 2H), 7.44 (td, J = 6.6, 3.1 Hz, 4H), 7.27 (dd, J = 7.0, 1.8, Hz, 1H), 7.14 (d, J = 2.7 Hz, 1H), 4.59 (d, J = 3.6 Hz, 2H), 4.47 (s, 1H), 4.43 – 4.33 (m, 2H), 1.46 (td, J = 7.2, 1.2 Hz, 3H). ^13^C NMR (101 MHz, CDCl3) δ 143.46, 143.45, 142.18, 141.63, 140.34, 139.53, 129.57, 129.03, 128.48, 128.46, 126.91, 126.07, 125.92, 125.56, 124.92, 123.26, 122.65, 120.52, 119.76, 118.97, 110.38, 108.76, 108.62, 48.53, 37.64, 13.84. HRMS (ESI-TOF) calcd for C_24_H_22_N_3_, 352.1804 (M+H^+^), found 352.1808.

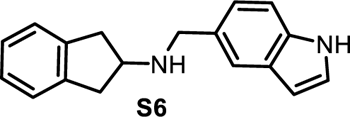

#### N-((1H-indol-5-yl)methyl)-2,3-dihydro-1H-inden-2-amine (S6)

Synthesized according to scheme 1general procedure 1 purified on Biotage^®^ with 0 to 6 % methanol gradient in dichloromethane to afford S6 as an brown solid (90 mg, 58 %). ^1^H NMR (400 MHz, CDCl3) δ 8.65 (s, 1H), 7.59 – 7.51 (m, 1H), 7.20 – 7.14 (m, 3H), 7.14 – 7.08 (m, 3H), 7.01 (dd, J = 3.2, 2.3 Hz, 1H), 6.44 (dd, J = 3.0, 1.9, Hz, 1H), 3.92 (s, 2H), 3.70 (p, J = 6.9 Hz, 1H), 3.16 (dd, J = 15.6, 7.2 Hz, 2H), 2.81 (dd, J = 15.6, 6.6 Hz, 2H). ^13^C NMR (101 MHz, CDCl_3_) δ 141.93, 135.21, 131.41, 128.10, 126.51, 124.84, 124.81, 122.73, 120.30, 111.28, 102.26, 58.85, 52.89, 40.11. HRMS (ESI-TOF) calcd for C_18_H_19_N_2_, 263.1543 (M+H^+^), found 263.1535.

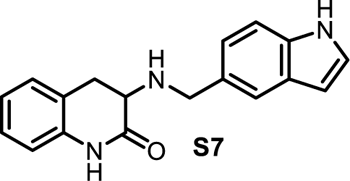

#### 3-(((1H-indol-5-yl)methyl)amino)-3,4-dihydroquinolin-2(1H)-one (S7)

Synthesized according to scheme 3, purified on Biotage^®^ with 5 to 80 % ethyl acetate gradient in hexane to afford S7 as an off white solid (90 mg, 58 %). ^1^H NMR (400 MHz, CD_3_OD_SPE) δ 7.53 (d, J = 1.6 Hz, 1H), 7.35 (d, J = 8.3 Hz, 1H), 7.20 (d, J = 3.1 Hz, 1H), 7.13 (td, J = 7.8, 2.8 Hz, 3H), 6.94 (td, J = 7.5, 1.2 Hz, 1H), 6.87 – 6.79 (m, 1H), 6.40 (dd, J = 3.2, 0.9 Hz, 1H), 4.01 (d, J = 12.6 Hz, 1H), 3.87 (d, J = 12.6 Hz, 1H), 3.40 (dd, J = 13.4, 6.2 Hz, 1H), 3.14 (dd, J = 15.3, 6.3 Hz, 1H), 128.21, 128.00, 127.37, 124.72, 122.79, 122.53, 121.70, 119.90, 114.98, 110.98, 100.95, 54.50, 51.37, 30.95. HRMS (ESI-TOF) calcd for C_18_H_19_N_3_O, 292.1445 (M+H^+^), found 292.1446

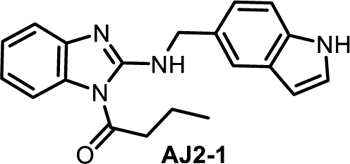

#### 1-(2-(((1H-indol-5-yl)methyl)amino)-1H-benzo[d]imidazol-1-yl)butan-1-one (AJ2-1)

Synthesized according to scheme 1 general procedure 1 and following general procedure 4, purified by Biotage^®^ with hexane: ethyl acetate, (6:4) to afford AJ2-1 as an off white solid (17 mg, 62 %). ^1^H NMR (400 MHz, CDCl_3_) δ 8.21 (s, 1H), 8.16 (s, 1H), 7.68 (dt, *J* = 1.6, 0.8 Hz, 1H), 7.47 (dd, *J* = 7.8, 1.2 Hz, 1H), 7.38 (ddd, *J* = 8.4, 2.5, 1.7 Hz, 2H), 7.26 – 7.20 (m, 4H), 7.06 (ddd, *J* = 8.6, 7.5, 1.3 Hz, 1H), 6.53 – 6.55 (m, 1H), 4.85 (d, *J* = 5.2 Hz, 2H), 2.99 (t, *J* = 7.2 Hz, 2H), 1.84 (p, *J* = 7.3 Hz, 2H), 1.08 (t, *J* = 7.4 Hz, 3H). ^13^C NMR (151 MHz, CDCl_3_) δ 174.57, 155.01, 143.89, 135.34, 130.14, 129.31, 128.08, 124.85, 124.74, 122.41, 120.21, 120.19, 117.13, 113.05, 111.33, 102.69, 47.60, 40.33, 17.27, 13.62. HRMS (ESI-TOF) calcd for C_20_H_21_N_4_O, 333.1710 (M+H^+^), found 333.1715.

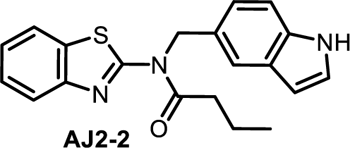

#### N-((1H-indol-5-yl)methyl)-N-(benzo[d]thiazol-2-yl)butyramide (AJ2-2)

Synthesized according to scheme 1 general procedure 1 and following general procedure 2, purified by PTLC with hexane: ethyl acetate, (7:3) to afford AJ2-2 as brown solid (8 mg, 62 %) ^1^H NMR (400 MHz, CDCl_3_) δ 8.17 (s, 1H), 7.84 (dd, *J* = 7.7, 1.1 Hz, 1H), 7.79 (dt, *J* = 8.2, 0.9 Hz, 1H), 7.48 – 7.45 (m, 1H), 7.40 (dd, *J* = 7.2, 1.3 Hz, 1H), 7.36 – 7.27 (m, 2H), 7.20 (dd, *J* = 3.2, 2.4 Hz, 1H), 7.09 (dd, *J* = 8.5, 1.8 Hz, 1H), 6.48 (d, *J* = 2.0, 1H), 5.74 (s, 2H), 2.62 (t, *J* = 7.3 Hz, 2H), 1.72 (q, *J* = 7.4 Hz, 2H), 0.92 (t, *J* = 7.4 Hz, 3H). ^13^C NMR (151 MHz, DMSO) δ 173.98, 159.69, 147.59, 135.12, 132.74, 127.74, 126.99, 126.05, 125.92, 123.78, 121.47, 120.90, 119.35, 117.03, 111.77, 100.99, 50.22, 35.73, 17.35, 13.43. HRMS (ESI-TOF) calcd for C_20_H_20_N_3_OS, 350.1322 (M+H^+^), found 350.1329.

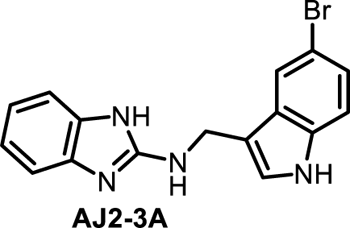

#### N-((5-bromo-1H-indol-3-yl)methyl)-1H-benzo[d]imidazol-2-amine (AJ2-3A)

Synthesized according to scheme 1, general procedure 1 purified on Biotage^®^ with dichloromethane: methanol (9:1) to afford AJ2-3A as light brown solid (160 mg, 64 %); ^1^H NMR (400 MHz, CD_3_OD) δ 7.74 (s, 1H), 7.28 (s, 1H), 7.23 (dd, *J* = 8.6, 0.6 Hz, 1H), 7.20 – 7.12 (m, 3H), 6.94 (dd, *J* = 5.8, 3.2 Hz, 2H), 4.64 (d, *J* = 0.8 Hz, 2H). ^13^C NMR (151 MHz, CD_3_OD_SPE) δ 156.77, 136.92, 129.81, 126.02, 125.37, 122.27, 121.36, 114.00, 113.41, 113.19, 112.77, 39.38. HRMS (ESI-TOF) calcd for C_16_H_14_BrN_4_, 341.0397 (M+H^+^), found 341.0396.

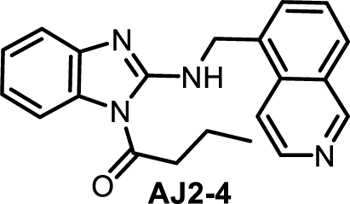

#### 1-(2-((isoquinolin-5-ylmethyl)amino)-1H-benzo[d]imidazol-1-yl)butan-1-one (AJ2-4)

Synthesized according to scheme 1, general procedure 1 and following general procedure 4, purified on Biotage^®^ with Hexane: Ethyl acetate (6:4) to afford AJ2-4 as light brown solid (7 mg, 54 %) ^1^H NMR (400 MHz, CD_3_OD) δ 9.17 (s, 1H), 8.40 (d, *J* = 6.1 Hz, 1H), 7.96 (dd, *J* = 7.2, 4.6 Hz, 2H), 7.76 (dd, *J* = 7.2, 1.2 Hz, 1H), 7.62 – 7.54 (m, 1H), 7.48 (d, *J* = 8.2 Hz, 1H), 7.24 – 7.20 (m, 1H), 7.12 (dd, *J* = 7.7, 1.0 Hz, 1H), 7.07 – 6.99 (m, 1H), 5.09 (s, 2H), 3.00 (t, *J* = 7.1 Hz, 2H), 1.73 (q, *J* = 7.3 Hz, 2H), 0.98 (t, *J* = 7.4 Hz, 3H). ^13^C NMR (151 MHz, CDCl_3_) δ 174.81, 154.76, 153.16, 143.67, 134.32, 132.84, 130.23, 130.17, 128.98, 128.01, 127.77, 127.35, 126.92, 125.05, 120.66, 117.37, 116.56, 113.17, 56.00, 44.15, 40.33, 17.21, 13.59. HRMS (ESI-TOF) calcd for C_21_H_21_N_4_O, 345.1710 (M+H^+^), found 345.1720.

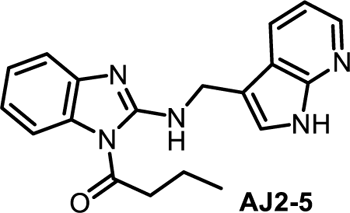

#### 1-(2-(((1H-pyrrolo[2,3-b]pyridin-3-yl)methyl)amino)-1H-benzo[d]imidazol-1-yl)butan-1-one (AJ2-5)

Synthesized according to scheme 1 general procedure 1 and following procedure 4, purified on Biotage^®^ with solvent Hexane: Ethyl acetate (6:4) to afford AJ2-5 as light brown solid to afford AJ2-5 as brown solid (12 mg, 62 %); ^1^H NMR (400 MHz, CDCl_3_) δ 10.42 (s, 1H), 8.33 (dd, *J* = 4.8, 1.5 Hz, 1H), 8.10 (t, *J* = 5.2 Hz, 1H), 8.05 (dd, *J* = 7.9, 1.5 Hz, 1H), 7.50 (dd, *J* = 8.0, 1.3 Hz, 1H), 7.44 – 7.38 (m, 2H), 7.30 – 7.24 (m, 2H), 7.13 – 7.05 (m, 2H), 4.93 (dd, *J* = 5.1, 0.8 Hz, 2H), 2.98 (t, *J* = 7.2 Hz, 2H), 1.83 (q, *J* = 7.3 Hz, 2H), 1.07 (t, *J* = 7.4 Hz, 3H). HRMS (ESI-TOF) calcd for C_19_H_20_N_5_O, 334.1663 (M+H^+^), found 334.1676.

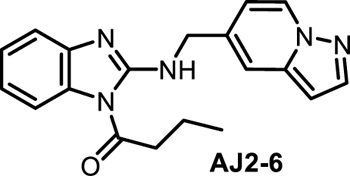

#### 1-(2-((pyrazolo[1,5-a]pyridin-5-ylmethyl)amino)-1H-benzo[d]imidazol-1-yl)butan-1-one (AJ2-6)

Synthesized according to scheme 1 general procedure 1 and following general procedure 4, purified on Biotage^®^ with hexane: ethyl acetate (6:4) to afford AJ2-6 as a brown solid (6 mg, 52 %) ^1^H NMR (400 MHz, CDCl_3_) δ 8.45 (d, *J* = 7.2 Hz, 1H), 8.34 (s, 1H), 7.95 (d, *J* = 2.3 Hz, 1H), 7.55 (s, 1H), 7.45 (t, *J* = 8.3 Hz, 2H), 7.29 (d, *J* = 0.9 Hz, 2H), 7.16 – 7.06 (m, 1H), 6.82 (dd, *J* = 7.2, 2.0 Hz, 1H), 6.49 (d, *J* = 2.3 Hz, 1H), 4.83 (d, *J* = 5.9 Hz, 2H), 3.06 (t, *J* = 7.2 Hz, 2H), 1.92 (q, *J* = 7.3 Hz, 2H), 1.14 (t, *J* = 7.4 Hz, 3H). ^13^C NMR (151 MHz, CDCl_3_) δ 174.89, 154.97, 143.45, 142.30, 139.95, 134.11, 130.18, 128.72, 125.02, 120.66, 117.39, 115.65, 113.14, 111.75, 96.82, 45.91, 40.36, 17.28, 13.62. HRMS (ESI-TOF) calcd for C_19_H_20_N_5_O, 334.1663 (M+H^+^), found 334.1676.

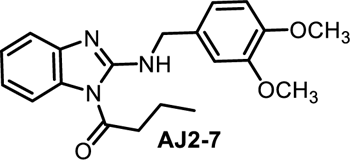

#### 1-(2-((3,4-dimethoxybenzyl)amino)-1H-benzo[d]imidazol-1-yl)butan-1-one (AJ2-7)

Synthesized according to scheme 1, general procedure 1 and following general procedure 4, purified on Biotage^®^ with hexane: ethyl acetate (6:4) to afford AJ2-7 as a brown solid (12 mg, 72%) ^1^H NMR (400 MHz, CDCl_3_) δ 8.14 (t, *J* = 5.5 Hz, 1H), 7.45 (dd, *J* = 7.9, 1.3 Hz, 1H), 7.38 (dt, *J* = 8.2, 0.8 Hz, 1H), 7.24 (dd, *J* = 7.7, 1.0 Hz, 1H), 7.06 (dd, *J* = 8.2, 1.3 Hz, 1H), 6.98 – 6.93 (m, 2H), 6.87 – 6.81 (m, 1H), 4.70 (d, *J* = 5.4 Hz, 2H), 3.88 (s, 3H), 3.87 (s, 3H), 2.99 (t, *J* = 7.2 Hz, 2H), 1.86 (q, *J* = 7.3 Hz, 2H), 1.09 (t, *J* = 7.4 Hz, 3H). ^13^C NMR (151 MHz, CDCl_3_) δ 174.68, 154.95, 149.13, 148.54, 143.67, 130.58, 130.10, 124.92, 120.36, 120.16, 117.15, 113.08, 111.31, 111.25, 55.96, 55.91, 46.79, 40.33, 17.26, 13.62. HRMS (ESI-TOF) calcd for C_20_H_24_N_3_O_3_, 354.1812 (M+H^+^), found 354.1822.

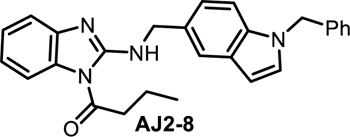

#### 1-(2-(((1-benzyl-1H-indol-5-yl)methyl)amino)-1H-benzo[d]imidazol-1-yl)butan-1-one (AJ2-8)

Synthesized according to scheme 1, general procedure 1 and following general procedure 4, purified on Biotage^®^ with hexane: ethyl acetate (6:4) to afford AJ2-8 as an off white solid (14 mg, 74 %) ^1^H NMR (400 MHz, CDCl_3_) δ 8.13 (s, 1H), 7.71 – 7.65 (m, 1H), 7.46 (dd, *J* = 7.9, 1.2 Hz, 1H), 7.38 (d, *J* = 8.1 Hz, 1H), 7.33 – 7.27 (m, 3H), 7.25 – 7.23 (m, 2H), 7.15 (d, *J* = 3.2 Hz, 1H), 7.10 (dd, *J* = 4.5, 2.1 Hz, 1H), 7.09 – 7.02 (m, 2H), 6.53 (dd, *J* = 3.1, 0.8 Hz, 1H), 5.32 (s, 2H), 4.84 (d, *J* = 5.1 Hz, 2H), 2.98 (t, *J* = 7.2 Hz, 2H), 1.84 (q, *J* = 7.4 Hz, 2H), 1.08 (t, *J* = 7.4 Hz, 3H). ^13^C NMR (151 MHz, CDCl_3_) δ 174.56, 154.98, 143.89, 137.46, 135.88, 130.13, 128.83, 128.80, 127.64, 126.77, 126.72, 124.85, 122.15, 121.45, 120.51, 120.18, 119.89, 117.14, 113.03, 110.38, 110.03, 101.74, 50.22, 47.59, 40.33, 17.26, 13.62. HRMS (ESI-TOF) calcd for C_27_H_27_N_4_O, 423.2180 (M+H^+^), found 423.2176.

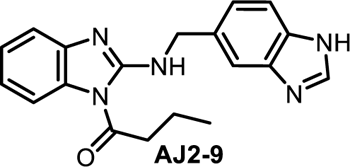

#### 1-(2-(((1H-benzo[d]imidazol-5-yl)methyl)amino)-1H-benzo[d]imidazol-1-yl)butan-1-one (AJ2-9)

Synthesized according to scheme 1, general procedure 1 and following general procedure 4, purified by PTLC with dichloromethane: methanol (9.5:0.5) to afford AJ2-9 as off white solid (6 mg, 48 %) ^1^H NMR (400 MHz, MeOD) δ 8.05 (s, 1H), 7.57 (d, *J* = 1.5 Hz, 1H), 7.50 (d, *J* = 8.3 Hz, 1H), 7.24 (dd, *J* = 8.3, 1.7 Hz, 1H), 7.15 ∼ 7.12 (m, 3H), 6.94 (dd, *J* = 5.9, 3.1 Hz, 2H), 4.62 (s, 2H), 2.12 (t, *J* = 7.4 Hz, 2H), 1.56 ∼ 1.47 (m, 2H), 0.84 (t, J = 7.4 Hz, 3H). ^13^C NMR (151 MHz, MeOD) δ 178.06, 154.19, 141.60, 135.66, 133.20, 122.19, 120.83, 120.72, 114.94, 111.28, 37.01, 29.36, 18.59, 12.75. HRMS (ESI-TOF) calcd for C_19_H_20_N_5_O, 334.1663 (M+H^+^), found 334.1674.

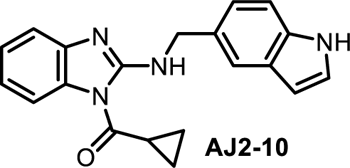

#### (2-(((1H-indol-5-yl)methyl)amino)-1H-benzo[d]imidazol-1-yl)(cyclopropyl)methanone (AJ2-10)

Synthesized according to scheme 1, general procedure 1 and following general procedure 4, purified by PTLC with hexane: ethyl acetate (6:4) to afford AJ2-10 as off white solid (11 mg, 54 %) ^1^H NMR (400 MHz, CDCl_3_) δ 8.30 (s, 1H), 7.68 (t, *J* = 5.2 Hz, 1H), 7.58 (s, 1H), 7.51 (d, *J* = 8.1 Hz, 1H), 7.40 (d, *J* = 7.9 Hz, 1H), 7.26 (d, *J* = 8.4 Hz, 1H), 7.15 (dd, *J* = 14.9, 2.5 Hz, 4H), 6.98 (t, *J* = 7.8 Hz, 1H), 6.44 (d, *J* = 3.2 Hz, 1H), 4.75 (d, *J* = 4.6 Hz, 2H), 2.41 (tt, *J* = 8.3, 4.6 Hz, 1H), 1.32 – 1.24 (m, 2H), 1.12 (dd, *J* = 7.8, 3.4 Hz, 2H). ^13^C NMR (151 MHz, CDCl_3_) δ 174.96, 154.41, 143.92, 135.36, 130.69, 129.28, 129.26, 128.07, 124.78, 124.73, 124.61, 122.32, 120.14, 117.07, 112.78, 111.36, 111.31, 102.62, 102.56, 47.56, 16.72, 10.26. HRMS (ESI-TOF) calcd for C_20_H_19_N_4_O, 331.1554 (M+H^+^), found 331.1554.

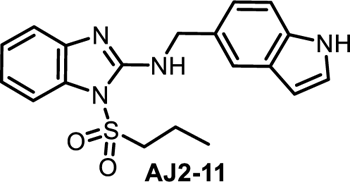

#### N-((1H-indol-5-yl)methyl)-1-(propylsulfonyl)-1H-benzo[d]imidazol-2-amine (AJ2-11)

Synthesized according to scheme 1, general procedure 1 and following general procedure 4 with 1-Propanesulfonyl chloride, purified by PTLC (Hexane/Ethyl acetate 7:3) to afford AJ2-11 as an off white solid (5 mg, 43 %) ^1^H NMR (400 MHz, DMSO) δ 11.05 (s, 1H), 7.56 (d, *J* = 1.6 Hz, 1H), 7.54 – 7.48 (m, 1H), 7.36 (d, *J* = 8.3 Hz, 1H), 7.34 – 7.28 (m, 2H), 7.22 – 7.14 (m, 2H), 7.09 (t, *J* = 5.9 Hz, 1H), 7.05 (td, *J* = 7.7, 1.2 Hz, 1H), 6.39 (dd, *J* = 2.0, 0.9 Hz, 1H), 4.69 (d, *J* = 5.8 Hz, 2H), 3.65 – 3.56 (m, 2H), 1.60 – 1.48 (m, 2H), 0.82 (t, *J* = 7.4 Hz, 3H). ^13^C NMR (151 MHz, DMSO) δ 152.60, 142.79, 135.63, 131.68, 129.76, 128.01, 126.12, 124.89, 121.49, 121.13, 119.41, 116.76, 112.22, 111.76, 101.42, 54.76, 47.20, 16.88, 12.49. HRMS (ESI-TOF) calcd for C_19_H_21_N_4_O_2_S, 369.1380 (M+H^+^), found 369.1381.

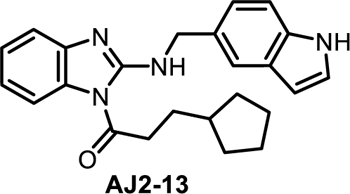

#### 1-(2-(((1H-indol-5-yl)methyl)amino)-1H-benzo[d]imidazol-1-yl)-3-cyclopentylpropan-1-one (AJ2-13)

Synthesized according to scheme 1, general procedure 1 and following general procedure 4, purified by PTLC with hexane: ethyl acetate (7:3) to afford AJ2-13 as off white solid (8 mg, 47 %) ^1^H NMR (400 MHz, CDCl_3_) δ 8.20 (s, 1H), 8.18 (t, *J* = 4.4 Hz, 1H), 7.69 – 7.66 (m, 1H), 7.47 (dd, *J* = 8.0, 1.2 Hz, 1H), 7.42 – 7.35 (m, 2H), 7.25 – 7.17 (m, 4H), 7.10 – 7.04 (m, 2H), 6.54 (dd, *J* = 2.0, 1.0 Hz, 1H), 4.84 (d, *J* = 5.1 Hz, 2H), 3.08 – 2.97 (m, 2H), 2.37 (s, 2H), 1.85 – 1.77 (m, 4H), 1.69 – 1.63 (m, 5H). ^13^C NMR (151 MHz, CDCl_3_) δ 155.39, 135.65, 134.62, 127.96, 127.89, 120.49, 120.05, 116.70, 109.02, 108.80, 65.90, 48.02, 37.63, 36.04, 34.97, 30.05, 28.04, 19.58, 15.51, 15.30. HRMS (ESI-TOF) calcd for C_24_H_27_N_4_O, 387.2180 (M+H^+^), found 387.2186.

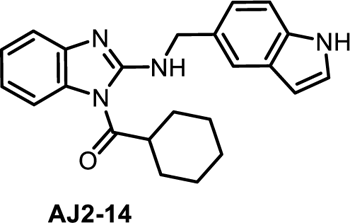

#### (2-(((1H-indol-5-yl)methyl)amino)-1H-benzo[d]imidazol-1-yl)(cyclohexyl)methanone (AJ2-14)

Synthesized according to scheme 1, general procedure 1 and following general procedure 4, purified by PTLC with hexane: ethyl acetate (6:4) to afford AJ2-14 as off white solid (6 mg, 47%) ^1^H NMR (400 MHz, CDCl_3_) δ 8.32 (s, 1H), 8.15 (t, *J* = 5.2 Hz, 1H), 7.58 (d, *J* = 1.6 Hz, 1H), 7.38 (dd, *J* = 7.9, 1.2 Hz, 1H), 7.27 (d, *J* = 8.3 Hz, 1H), 7.22 – 7.13 (m, 4H), 7.04 – 6.96 (m, 2H), 6.49 – 6.42 (m, 1H), 4.74 (d, *J* = 5.1 Hz, 2H), 3.12 – 3.07 (m, 1H), 2.02 – 1.93 (m, 2H), 1.87 – 1.81 (m, 2H), 1.75 – 1.65 (m, 2H), 1.57 – 1.50 (m, 2H), 1.37 (dt, *J* = 12.7, 3.3 Hz, 2H). ^13^C NMR (151 MHz, CDCl_3_) δ 178.08, 155.28, 143.83, 135.37, 129.83, 129.18, 128.08, 124.87, 124.78, 122.35, 120.39, 120.17, 117.12, 112.82, 111.36, 102.62, 47.67, 44.79, 29.04, 28.73, 25.63, 25.47, 25.42. HRMS (ESI-TOF) calcd for C_23_H_25_N_4_O, 373.2023 (M+H^+^), found 373.2035.

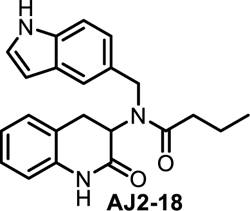

#### N-((1H-indol-5-yl)methyl)-N-(2-oxo-1,2,3,4-tetrahydroquinolin-3-yl)butyramide (AJ2-18)

Synthesized according to scheme 2 and general procedure 4, purified on Biotage^®^ with hexane: ethyl acetate (6:4) to afford AJ2-18 as off white solid (14 mg, 62 %); ^1^H NMR (400 MHz, CDCl_3_) δ 8.46 (s, 1H), 8.31 (s, 1H), 7.48 (d, *J* = 1.6 Hz, 1H), 7.32 (d, *J* = 8.3 Hz, 1H), 7.19 – 7.13 (m, 1H), 7.02 (dd, *J* = 8.3, 1.7 Hz, 2H), 6.87 (d, *J* = 7.6 Hz, 1H), 6.79 (td, *J* = 7.4, 1.1 Hz, 1H), 6.62 (dd, *J* = 7.9, 1.1 Hz, 1H), 6.49 – 6.42 (m, 1H), 5.00 – 4.85 (m, 1H), 4.77 (d, *J* = 17.1 Hz, 1H), 4.63 (d, *J* = 17.0 Hz, 1H), 3.34 (t, *J* = 14.8 Hz, 1H), 2.69 (dd, *J* = 15.3, 6.7 Hz, 1H), 2.51 – 2.32 (m, 2H), 1.68 (q, *J* = 7.4 Hz, 2H), 0.88 (t, *J* = 7.4 Hz, 3H). ^13^C NMR (101 MHz, CDCl_3_) δ 174.69, 169.14, 136.32, 135.33, 128.58, 128.32, 128.13, 127.62, 125.12, 122.95, 122.50, 120.65, 118.54, 115.14, 111.61, 102.50, 55.30, 51.88, 35.64, 30.32, 18.75, 13.90. HRMS (ESI-TOF) calcd for C_22_H_23_N_3_NaO_2_, 384.1682 (M+Na^+^), found 384.1697.

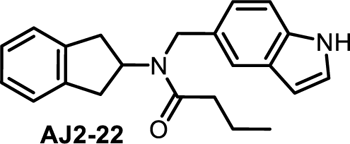

#### N-((1H-indol-5-yl)methyl)-N-(2,3-dihydro-1H-inden-2-yl)butyramide (AJ2-22)

Synthesized according to scheme 1, general procedure 1 and following and general procedure 4, purified on Biotage^®^ with hexane: ethyl acetate (6:4) to afford AJ2-22 as brown viscous liquid (17 mg, 68%); ^1^H NMR (400 MHz, CDCl_3_) δ 8.79 (s, 1H), 8.55 (s, 0.39 H), 7.41 (d, *J* = 1.8 Hz, 1H), 7.39 – 7.32 (m, 1.61 H), 7.26 (d, *J* = 8.4 Hz, 0.5 H), 7.19 (t, *J* = 2.8 Hz, 1.12 H), 7.17 – 7.12 (m, 1.93 H), 7.00 (d, *J* = 8.4 Hz, 0.45 H), 6.95 – 6.93 (dd, *J* = 8.4, 1.9 Hz, 1.19 H), 6.50 (t, *J* = 2.7 Hz, 1.10 H), 6.45 (s, 0.42 H), 5.58 – 5.50 (m, 1.14 H), 4.90 (t, *J* = 8.2 Hz, 0.43 H), 4.74 (s, 0.89 H), 4.64 (s, 2.20 H), 3.17 – 2.93 (m, 6.63 H), 2.57 (t, *J* = 7.6 Hz, 0.93 H), 2.32 (t, *J* = 7.5 Hz, 2.28 H), 1.83 (q, *J* = 7.5 Hz, 1.01H), 1.74 – 1.65 (m, 2.49 H), 1.04 (t, *J* = 7.4 Hz, 1.43 H), 0.89 (t, *J* = 7.4 Hz, 3.53H). ^13^C NMR (151 MHz, DMSO) δ 173.33, 172.89, 141.56, 141.13, 135.53, 135.29, 130.38, 129.33, 128.30, 128.07, 126.93, 126.75, 126.22, 125.83, 124.78, 124.75, 120.51, 119.71, 117.97, 117.38, 112.08, 111.54, 101.47, 101.30, 58.09, 56.00, 55.38, 48.80, 45.07, 36.88, 36.15, 35.58, 35.31, 19.11, 18.68, 14.37, 14.20. Note: rotomeric isomers observed. HRMS (ESI-TOF) calcd for C_22_H_25_N_2_O, 333.1962 (M+H^+^), found 333.1960.

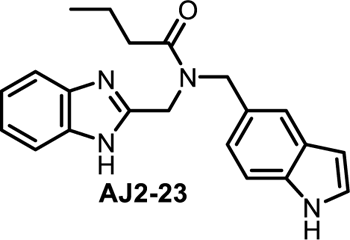

#### N-((1H-benzo[d]imidazol-2-yl)methyl)-N-((1H-indol-5-yl)methyl)butyramide (AJ2-23)

Synthesized according to scheme 1, general procedure 1 and following general procedure 4, purified by PTLC with hexane: ethyl acetate (6:4) to afford AJ2-23 as viscous liquid (13 mg, 57%); ^1^H NMR (400 MHz, CDCl_3_) δ 10.50 (s, 1H), 9.17 (s, 1H), 7.79 – 7.69 (m, 1H), 7.47 – 7.39 (m, 2H), 7.28 – 7.23 (m, 4H), 6.89 (dd, *J* = 8.3, 1.7 Hz, 1H), 6.53 – 6.47 (m, 1H), 4.70 (s, 2H), 4.69 (s, 2H), 2.48 (t, *J* = 7.5 Hz, 2H), 1.74 (h, *J* = 7.4 Hz, 2H), 0.96 (t, *J* = 7.4 Hz, 3H). ^13^C NMR (151 MHz, CDCl_3_) δ 175.42, 151.21, 135.50, 128.25, 126.69, 125.16, 120.85, 118.71, 111.72, 102.48, 52.27, 44.34, 35.10, 18.73, 13.95. HRMS (ESI-TOF) calcd for C_21_H_23_N_4_O, 347.1867 (M+H^+^), found 347.1853.

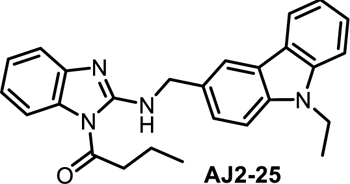

#### 1-(2-(((9-ethyl-9H-carbazol-3-yl)methyl)amino)-1H-benzo[d]imidazol-1-yl)butan-1-one (AJ2-25)

Synthesized according to scheme 1, general procedure 1 and following general procedure 4, purified on Biotage^®^ with hexane: ethyl acetate (6:4) to afford AJ2-25 as a yellow solid (16 mg, 68 %); ^1^H NMR (400 MHz, CDCl_3_) δ 8.21 (s, 1H), 8.14 (s, 1H), 8.09 (d, *J* = 7.8 Hz, 1H), 7.54 (dd, *J* = 8.4, 1.5 Hz, 1H), 7.51 – 7.44 (m, 2H), 7.44 – 7.37 (m, 3H), 7.26 – 7.19 (m, 2H), 7.11 – 7.03 (m, 1H), 4.94 (d, *J* = 5.2 Hz, 2H), 4.38 (q, *J* = 7.2 Hz, 2H), 3.00 (t, *J* = 7.2 Hz, 2H), 1.85 (h, *J* = 7.4 Hz, 2H), 1.49 – 1.37 (m, 3H), 1.14 – 1.04 (m, 3H). ^13^C NMR (151 MHz, CDCl_3_) δ 174.62, 155.00, 143.88, 140.29, 139.51, 130.16, 128.35, 125.96, 125.75, 124.88, 123.13, 122.74, 120.55, 120.24, 120.18, 118.84, 117.19, 113.06, 108.68, 108.51, 47.49, 40.35, 37.62, 17.26, 13.82, 13.62. HRMS (ESI-TOF) calcd for C_26_H_27_N_4_O, 411.2180 (M+H^+^), found 411.2180.

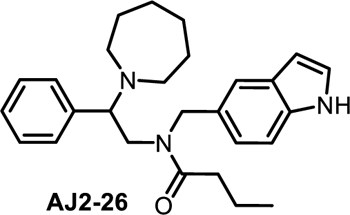

#### N-((1H-indol-5-yl)methyl)-N-(2-(azepan-1-yl)-2-phenylethyl)butyramide (AJ2-26)

Synthesized according to scheme 1, general procedure 1 and following general procedure 4, purified PTLC with Hexane: Ethyl acetate (7:3) to afford AJ2-26 as a colorless liquid (22 mg, 56 %); ^1^H NMR (400 MHz, CD_2_Cl_2_) δ 8.97 (s, 1H), 7.67 – 7.48 (m, 2H), 7.42 – 7.31 (m, 2H), 7.25 (d, *J* = 8.3 Hz, 2H), 7.13 (t, *J* = 2.6 Hz, 2H), 6.69 (dd, *J* = 8.4, 1.7 Hz, 1H), 6.36 (t, *J* = 2.4 Hz, 1H), 4.41 – 4.29 (m, 2H), 3.62 – 3.51 (m, 2H), 3.05 (s, 2H), 2.29 – 2.09 (m, 3H), 1.79 – 1.69 (m, 3H), 1.50 (dt, *J* = 14.8, 9.3 Hz, 9H), 0.80 (d, *J* = 7.4 Hz, 3H). ^13^C NMR (151 MHz, CDCl_3_) δ 173.89, 173.54, 135.13, 128.88, 128.57, 128.27, 128.10, 125.69, 125.44, 124.86, 124.48, 122.66, 120.59, 120.31, 118.27, 111.38, 111.11, 102.56, 68.89, 66.49, 52.69, 52.56, 52.02, 48.67, 35.53, 35.00, 29.72, 26.92, 26.84, 19.00, 18.86, 14.07, 14.03. Note: rotomeric isomers observed, HRMS (ESI-TOF) calcd for C_27_H_36_N_3_O, 418.2853 (M+H^+^), found 418.2853.

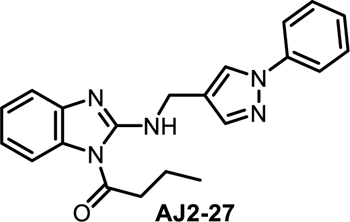

#### 1-(2-(((1-phenyl-1H-pyrazol-4-yl)methyl)amino)-1H-benzo[d]imidazol-1-yl)butan-1-one (AJ2-27)

Synthesized according to scheme 1, general procedure 1 and following general procedure 4, purified PTLC with Hexane: Ethyl acetate (6:4) to afford AJ2-27 as a colorless liquid (16 mg, 62 %); ^1^H NMR (400 MHz, CDCl_3_) δ 8.11 (t, *J* = 5.6 Hz, 1H), 8.01 (d, *J* = 0.8 Hz, 1H), 7.77 (d, *J* = 0.7 Hz, 1H), 7.69 – 7.63 (m, 2H), 7.49 – 7.37 (m, 4H), 7.31 – 7.24 (m, 3H), 7.08 (dd, *J* = 8.5, 1.3 Hz, 1H), 4.71 (d, *J* = 5.5 Hz, 2H), 3.00 (t, *J* = 7.2 Hz, 2H), 1.87 (h, *J* = 7.3 Hz, 2H), 1.09 (t, *J* = 7.4 Hz, 3H). ^13^C NMR (151 MHz, CDCl_3_) δ 174.67, 154.76, 143.61, 140.78, 140.06, 130.12, 129.42, 126.51, 126.17, 124.93, 120.46, 120.41, 119.15, 117.21, 113.11, 40.31, 37.23, 17.25, 13.62. HRMS (ESI-TOF) calcd for C_21_H_22_N_5_O, 360.1819 (M+H^+^), found 360.1816.

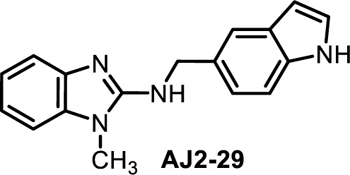

#### N-((1H-indol-5-yl)methyl)-1-methyl-1H-benzo[d]imidazol-2-amine (AJ2-29)

Synthesized according to scheme 1, purified on Biotage^®^ with Dichloromethane: ethyl acetate (6:4) to afford biotage (DCM/MeOH; 9:1) to afford AJ2-29 as a brown solid (165 mg, 72 %); ^1^H NMR (400 MHz, CDCl_3_) δ 8.40 (s, 1H), 7.70 (d, *J* = 1.6 Hz, 1H), 7.53 (dt, *J* = 7.7, 1.0 Hz, 1H), 7.42 – 7.36 (m, 1H), 7.31 – 7.24 (m, 3H), 7.13 (ddd, *J* = 7.7, 5.0, 3.7 Hz, 1H), 7.10 – 7.05 (m, 2H), 6.55 (d, *J* = 1.1 Hz, 1H), 4.81 (d, *J* = 5.1 Hz, 2H), 4.24 (d, *J* = 5.5 Hz, 1H), 3.46 (s, 3H). ^13^C NMR (151 MHz, DMSO) δ 155.74, 142.91, 135.78, 135.52, 130.85, 127.99, 125.93, 121.62, 120.66, 119.22, 118.73, 115.32, 111.57, 107.63, 101.37, 46.85, 28.69. HRMS (ESI-TOF) calcd for C_17_H_17_N_4_, 277.1448 (M+H^+^), found 277.1441.

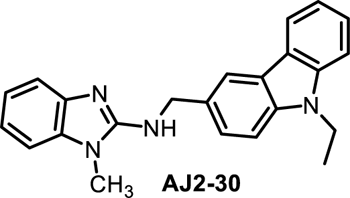

#### N-((9-ethyl-9H-carbazol-3-yl)methyl)-1-methyl-1H-benzo[d]imidazol-2-amine (AJ2-30)

Synthesized according to scheme 1 and general procedure 1, purified on Biotage^®^ with 0 to 6 % methanol gradient in dichloromethane to afford AJ2-30 as a light yellow solid (248 mg, 76 %); ^1^H NMR (400 MHz, DMSO) δ 8.17 (t, *J* = 1.1 Hz, 1H), 8.12 (dt, *J* = 7.8, 1.0 Hz, 1H), 7.61 – 7.53 (m, 3H), 7.43 (dd, *J* = 8.3, 1.2 Hz, 1H), 7.23 (t, *J* = 5.9 Hz, 1H), 7.23 – 7.13 (m, 3H), 7.00 – 6.84 (m, 2H), 4.75 (d, *J* = 5.8 Hz, 2H), 4.42 (q, *J* = 7.1 Hz, 2H), 3.55 (s, 3H), 1.28 (t, *J* = 7.1 Hz, 3H). ^13^C NMR (101 MHz, CDCl_3_) δ 154.38, 142.32, 140.31, 139.55, 135.03, 128.90, 126.27, 125.89, 123.13, 122.67, 121.24, 120.52, 120.38, 119.61, 118.96, 116.53, 108.68, 108.59, 107.05, 48.26, 37.63, 28.24, 13.82. HRMS (ESI-TOF) calcd for C_23_H_23_N_4_, 355.1917 (M+H^+^), found 355.1925.

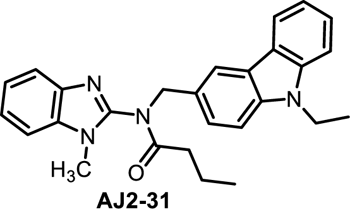

#### N-((9-ethyl-9H-carbazol-3-yl)methyl)-N-(1-methyl-1H-benzo[d]imidazol-2-yl)butyramide (AJ2-31)

Synthesized according to scheme 1, general procedure 1 and following general procedure 3, purified on Biotage^®^ with 0 to 5 % methanol gradient in dichloromethane to afford AJ2-31 as a white solid (64 mg, 52 %); ^1^H NMR (400 MHz, DMSO) δ 8.15 – 7.86 (m, 2H), 7.64 (d, *J* = 7.6 Hz, 1H), 7.57 (d, *J* = 8.2 Hz, 1H), 7.50 (dd, *J* = 11.8, 8.1 Hz, 2H), 7.45 – 7.43 (m, 1H), 7.35 (d, *J* = 8.5 Hz, 1H), 7.25 (p, *J* = 7.4 Hz, 2H), 7.19 – 7.12 (m, 1H), 5.07 (s, 2H), 4.40 (q, *J* = 7.1 Hz, 2H), 3.39 (s, 3H), 2.00 (s, 2H), 1.62 – 1.48 (m, 2H), 1.28 (t, *J* = 7.1 Hz, 3H), 0.81 (d, *J* = 7.7 Hz, 3H).^13^C NMR (151 MHz, DMSO) δ 171.61, 147.34, 139.80, 139.16, 138.30, 133.94, 126.53, 125.82, 125.15, 122.23, 121.58, 121.29, 119.82, 119.61, 118.74, 118.12, 110.28, 108.53, 108.33, 50.42, 36.34, 34.55, 28.66, 17.24, 13.05, 12.90. Note: rotomeric isomers observed, HRMS (ESI-TOF) calcd for C_27_H_29_N_4_O, 425.2336 (M+H^+^), found 425.2353.

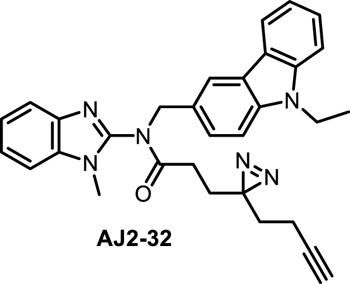

#### 3-(3-(but-3-yn-1-yl)-3H-diazirin-3-yl)-N-((9-ethyl-9H-carbazol-3-yl)methyl)-N-(1-methyl-1H-benzo[d]imidazol-2-yl)propanamide (AJ2-32)

Synthesized according to scheme 1 and general procedure 1 following general procedure 3, purified on Biotage^®^ with 0 to 5 % methanol gradient in dichloromethane to afford AJ2-32 as a light brown viscous liquid (12 mg, 46 %); ^1^H NMR (400 MHz, CDCl_3_) δ 8.00 (d, *J* = 7.8 Hz, 1H), 7.97 (d, *J* = 1.6 Hz, 1H), 7.83 (dd, *J* = 6.9, 2.1 Hz, 1H), 7.50 – 7.46 (m, 1H), 7.41 (dt, *J* = 8.3, 1.0 Hz, 1H), 7.39 – 7.26 (m, 4H), 7.25 – 7.18 (m, 2H), 5.19 (s, 2H), 4.34 (q, *J* = 7.2 Hz, 2H), 3.05 (s, 3H), 1.99 (td, *J* = 7.4, 2.6 Hz, 3H), 1.95 – 1.81 (m, 4H), 1.62 (t, *J* = 7.4 Hz, 2H), 1.42 (t, *J* = 7.2 Hz, 3H). ^13^C NMR (151 MHz, CDCl_3_) δ171.28, 147.65, 140.97, 140.25, 139.62, 134.48, 127.08, 126.62, 125.93, 123.58, 123.05, 122.96, 122.63, 121.50, 120.61, 120.32, 118.99, 110.05, 108.57, 108.53, 82.69, 69.19, 52.72, 37.62, 32.42, 29.24, 28.16, 27.76, 27.67, 13.80, 13.22. Note: rotomeric isomers observed, HRMS (ESI-TOF) calcd for C_31_H_31_N_6_O, 503.2554 (M+H^+^), found 503.2556.

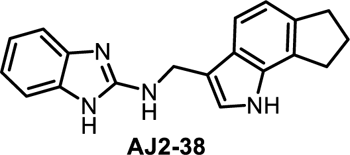

#### N-((1,6,7,8-tetrahydrocyclopenta[g]indol-3-yl)methyl)-1H-benzo[d]imidazol-2-amine (AJ2-38)

Synthesized according to scheme 1 and general procedure 1, purified on purified on Biotage^®^ with 0 to 5 % methanol gradient in dichloromethane to afford AJ2-38 as an off white solid (64 mg, 72 %); ^1^H NMR (400 MHz, CD_3_OD) δ 7.38 (d, *J* = 8.0 Hz, 1H), 7.22 (dd, *J* = 5.8, 3.2 Hz, 2H), 7.17 (s, 1H), 6.99 (dd, *J* = 5.8, 3.2 Hz, 2H), 6.91 (d, *J* = 8.0 Hz, 1H), 4.70 (s, 2H), 2.97 (dt, *J* = 23.9, 7.3 Hz, 4H), 2.11 (p, *J* = 7.4 Hz, 2H). ^13^C NMR (101 MHz, CD_3_OD) δ 154.74, 137.63, 136.51, 133.87, 125.45, 125.30, 122.36, 120.45, 116.22, 115.69, 112.07, 111.32, 47.52, 47.31, 47.09, 38.70, 32.64, 29.49, 25.02. HRMS (ESI-TOF) calcd for C_19_H_19_N_4_, 303.1604 (M+H^+^), found 303.1602.

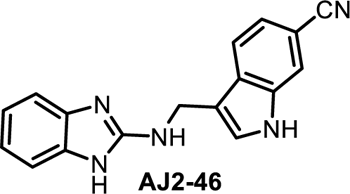

#### 3-(((1H-benzo[d]imidazol-2-yl)amino)methyl)-1H-indole-6-carbonitrile (AJ2-46)

Synthesized according to scheme 1 and general procedure 1, purified on Biotage^®^ with 0 to 70 % ethyl acetate gradient in hexane to afford AJ2-46 as an brown solid (35 mg, 58%); ^1^H NMR (400 MHz, CD_3_OD) δ 7.83 – 7.75 (m, 2H), 7.58 (d, *J* = 0.9 Hz, 1H), 7.30 (dd, *J* = 8.2, 1.5 Hz, 1H), 7.23 (dd, *J* = 5.8, 3.2 Hz, 2H), 7.00 (dd, *J* = 5.8, 3.2 Hz, 2H), 4.77 (d, *J* = 0.8 Hz, 2H). ^13^C NMR (151 MHz, DMSO) δ 155.98, 135.58, 135.55, 130.16, 128.91, 121.68, 121.20, 120.54, 116.88, 114.67, 102.96, 37.89. HRMS (ESI-TOF) calcd for C_17_H_14_N_5_, 288.1244 (M+H^+^), found 288.1239.

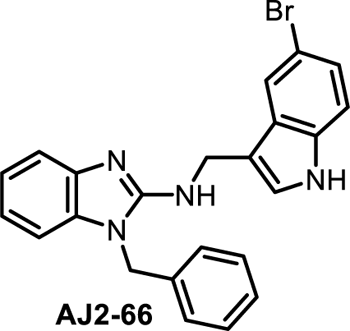

#### 1-Benzyl-N-((5-bromo-1H-indol-3-yl)methyl)-1H-benzo[d]imidazol-2-amine (AJ2-66)

Synthesized according to scheme 1 and general procedure 1, purified purified on Biotage^®^ with 0 to 70 % ethyl acetate gradient in hexane to afford AJ2-66 as a brown solid (48 mg, 63%), ^1^H NMR (400 MHz, CDCl_3_) δ 9.64 – 9.52 (m, 1H), 7.38 – 7.30 (m, 2H), 7.09 (dd, *J* = 5.0, 1.9 Hz, 3H), 7.03 – 6.97 (m, 3H), 6.94 (ddd, *J* = 6.6, 4.9, 2.3 Hz, 4H), 6.83 (d, *J* = 2.3 Hz, 1H), 4.92 (s, 2H), 4.51 (s, 2H). ^13^C NMR (101 MHz, CDCl_3_) δ 153.72, 140.70, 135.15, 134.96, 134.32, 129.16, 128.24, 128.10, 126.53, 124.88, 124.68, 121.86, 121.07, 120.48, 115.77, 113.02, 112.76, 111.71, 107.89, 45.80, 39.24. HRMS (ESI-TOF) calcd for C_23_H_20_BrN_4_, 431.0866 (M+H^+^), found 431.0850.

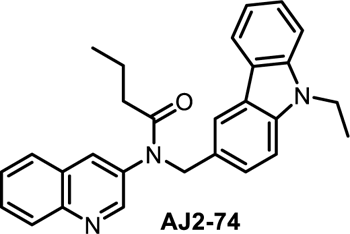

#### N-((9-ethyl-9H-carbazol-3-yl)methyl)-N-(quinolin-3-yl)butyramide (AJ2-74)

Synthesized according to Scheme 1 and general procedure 1 followed by general procedure 4, purified on PTLC with Hexane: Ethyl acetate (6:4) to afford AJ2-74 as a brown solid (17 mg, 68%), ^1^H NMR (400 MHz, CDCl_3_) δ 8.61 – 8.32 (m, 1H), 8.02 (d, *J* = 8.5 Hz, 1H), 7.92 (d, *J* = 7.7 Hz, 1H), 7.85 (d, *J* = 1.2 Hz, 1H), 7.71 – 7.57 (m, 3H), 7.46 (ddd, *J* = 8.1, 6.8, 1.1 Hz, 1H), 7.41 – 7.28 (m, 2H), 7.21 – 7.16 (m, 1H), 7.10 (ddd, *J* = 7.9, 7.0, 1.1 Hz, 1H), 5.10 (s, 2H), 4.26 (q, *J* = 7.2 Hz, 2H), 2.06 – 1.92 (m, 2H), 1.61 (q, *J* = 7.4 Hz, 2H), 1.34 (t, *J* = 7.2 Hz, 3H), 0.77 (t, *J* = 7.4 Hz, 3H). ^13^C NMR (151 MHz, CDCl_3_) δ 172.80, 150.98, 147.07, 140.23, 139.50, 135.81, 134.68, 130.14, 129.37, 127.91, 127.83, 127.46, 127.42, 126.90, 125.78, 123.07, 122.69, 121.15, 120.55, 118.85, 108.51, 108.47, 53.44, 37.61, 36.77, 18.87, 13.84, 13.82. HRMS (ESI-TOF) calcd for C_28_H_28_N_3_O, 422.2227 (M+H^+^), found 422.2217.

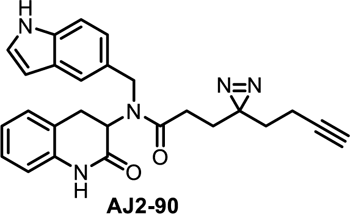

#### N-((1H-indol-5-yl)methyl)-3-(3-(but-3-yn-1-yl)-3H-diazirin-3-yl)-N-(2-oxo-1,2,3,4-tetrahydroquinolin-3-yl)propenamide (AJ2-90)

Synthesized according to general scheme 2 and general procedure 2, purified by PTLC (Hexane/Ethyl acetate; 5:5) to afford AJ2-90 as white solid (12 mg, 46%), ^1^H NMR (400 MHz, CDCl_3_) δ 8.27 (s, 1H), 7.87 (s, 1H), 7.48 (d, *J* = 1.7 Hz, 1H), 7.34 (d, *J* = 8.4 Hz, 1H), 7.19 (d, *J* = 2.4 Hz, 1H), 7.04 (td, *J* = 6.7, 3.4 Hz, 2H), 6.91 (d, *J* = 7.4 Hz, 1H), 6.82 (td, *J* = 7.5, 1.1 Hz, 1H), 6.61 (dd, *J* = 7.9, 1.1 Hz, 1H), 6.48 – 6.49 (m, 1H), 4.88 – 4.83 (m, 1H), 4.72 – 4.59 (m, 2H), 3.42 – 3.34 (m, 1H), 2.74 – 2.70 (m, 1H), 2.27 – 2.15 (m, 2H), 1.89 – 1.77 (m, 4H), 1.56 – 1.49 (m, 2H).). ^13^C NMR (151 MHz, CDCl_3_) δ 172.65, 168.47, 136.18, 135.33, 128.43, 128.18, 127.71, 125.10, 123.06, 122.39, 120.73, 118.69, 114.96, 111.66, 102.69, 102.62, 82.86, 69.09, 55.58, 51.99, 32.49, 30.26, 29.72, 28.01, 27.72, 13.29. HRMS (ESI-TOF) calcd for C_26_H_26_N_5_NaO_2_, 462.1900 (M+Na^+^), found 462.1908.

## II. NMR spectra

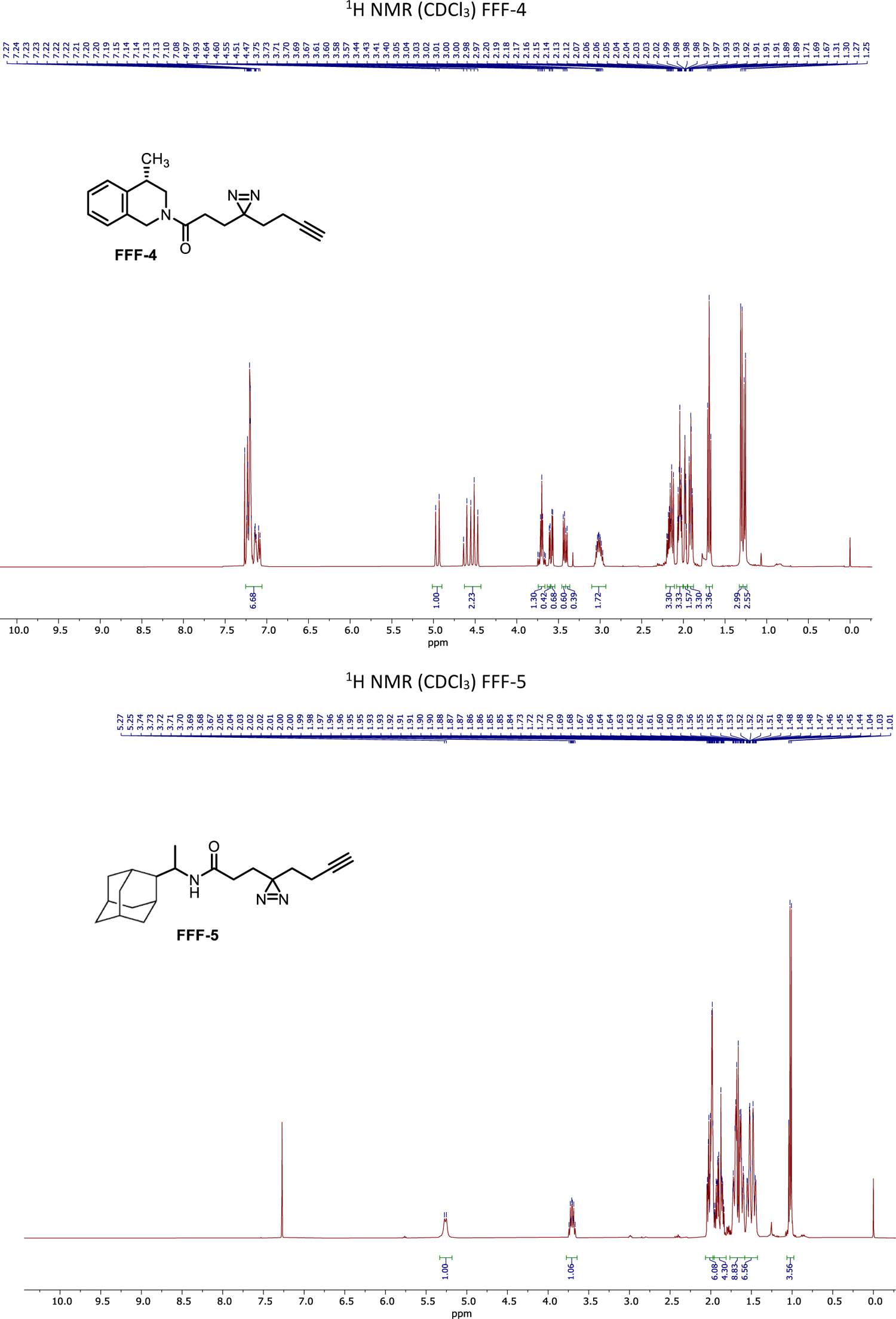
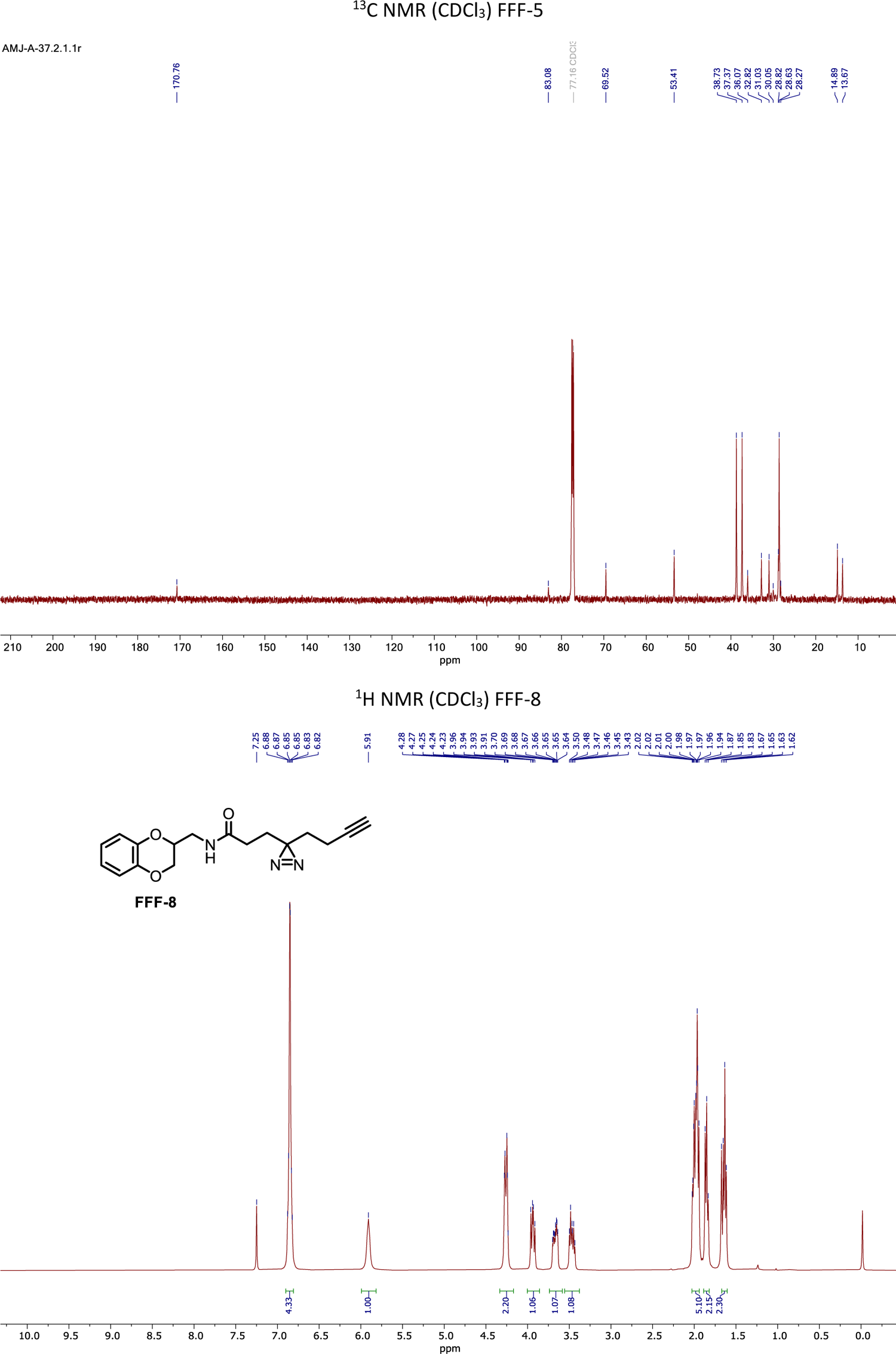
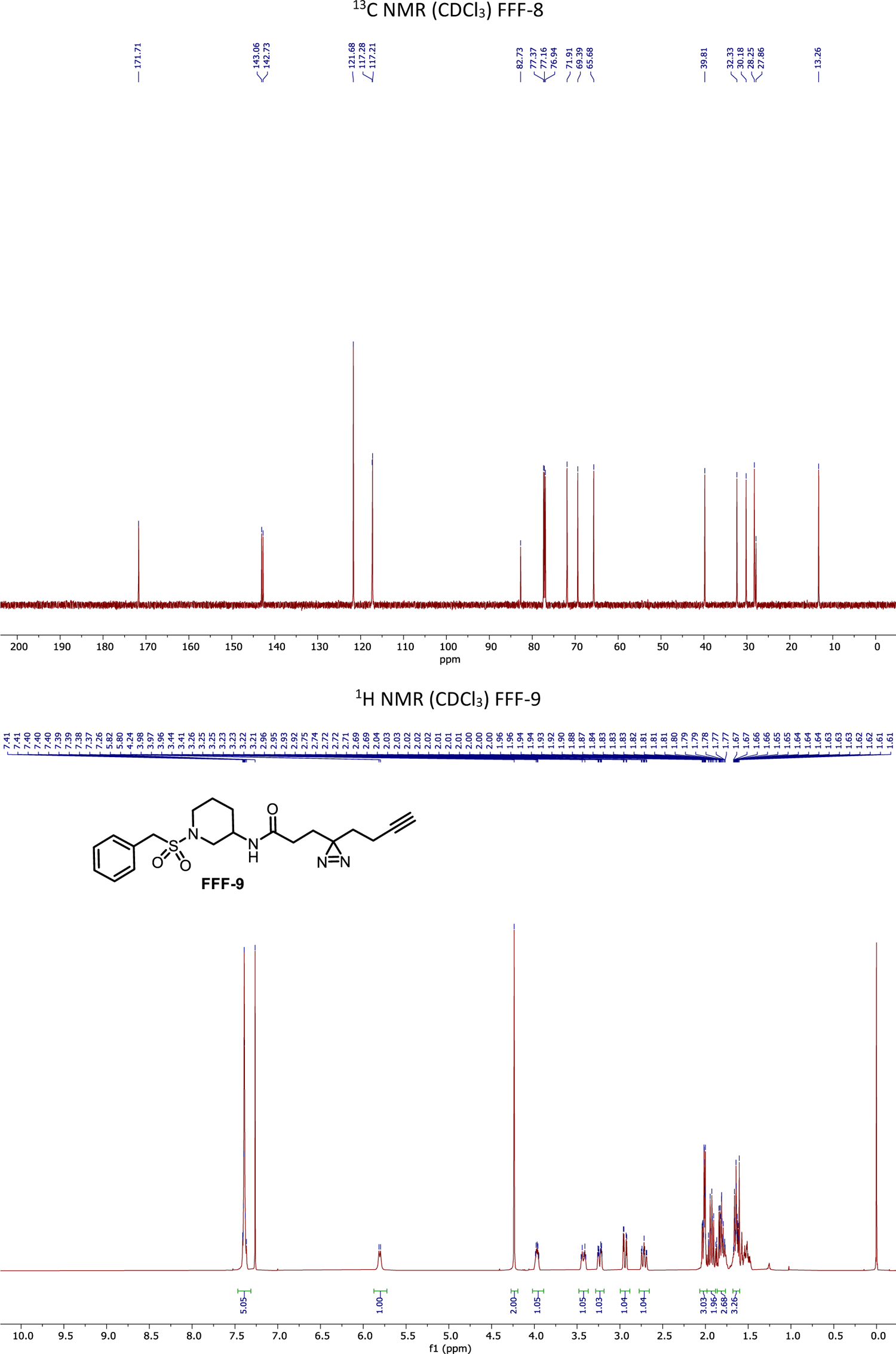
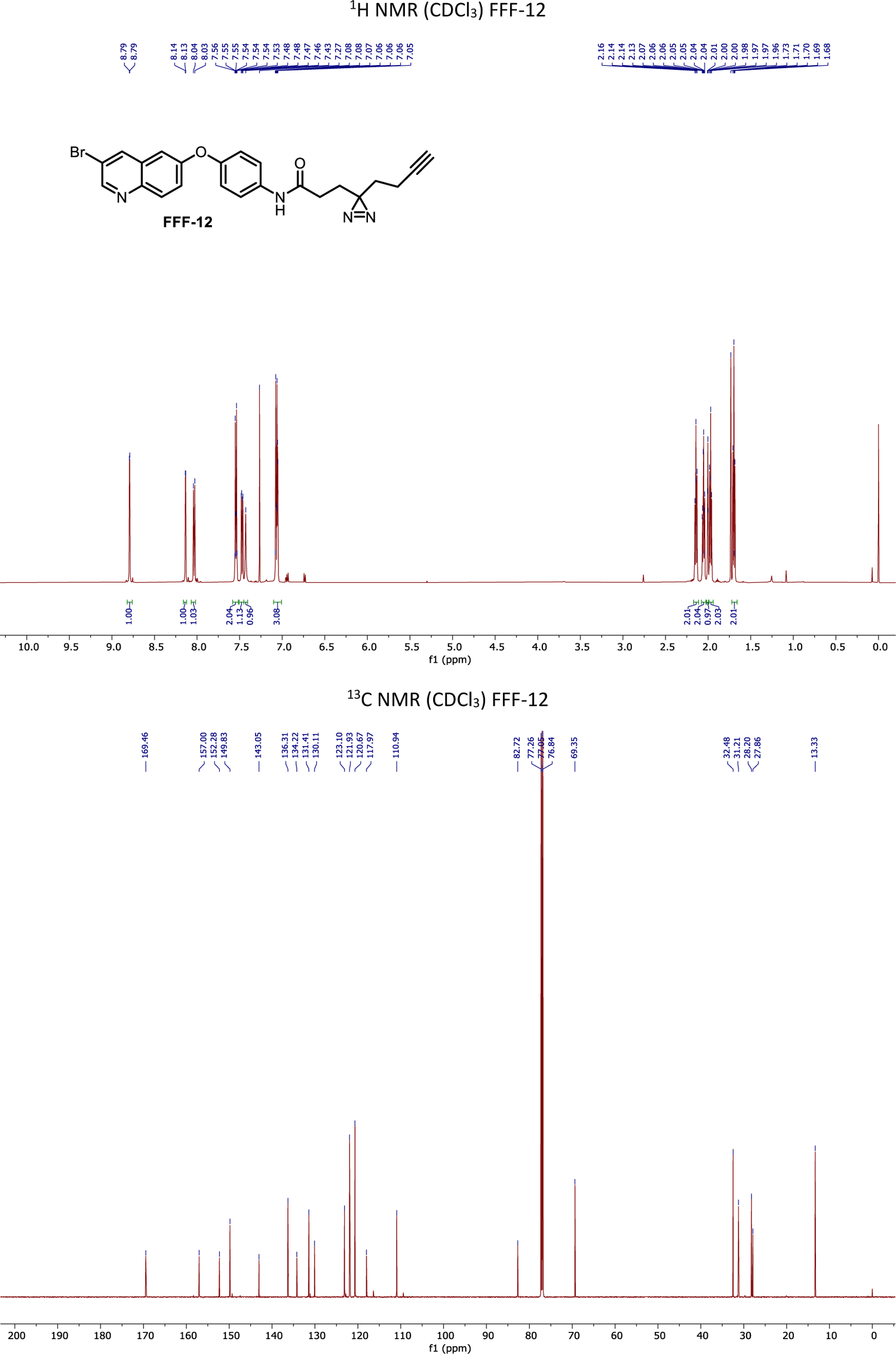
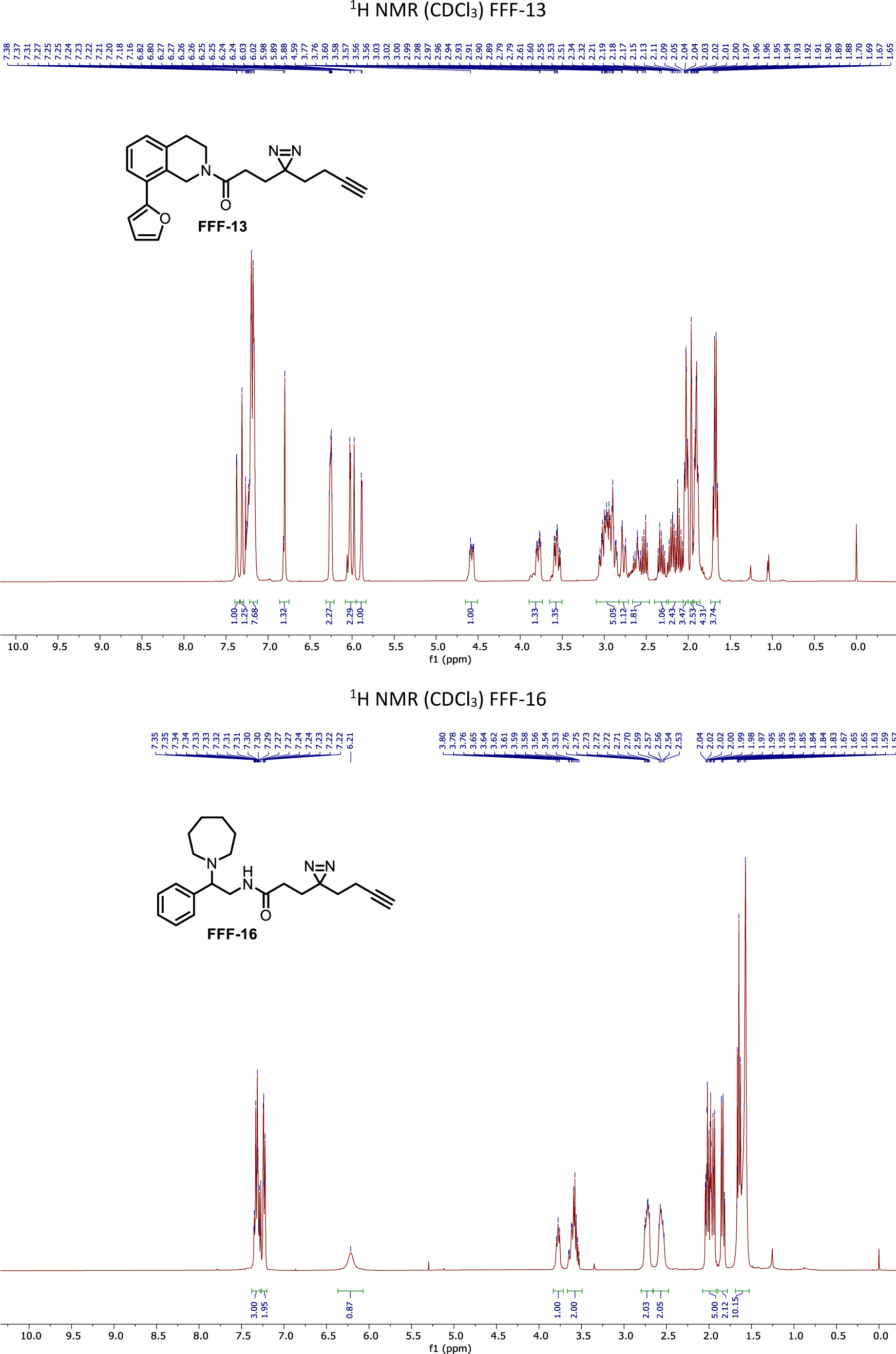
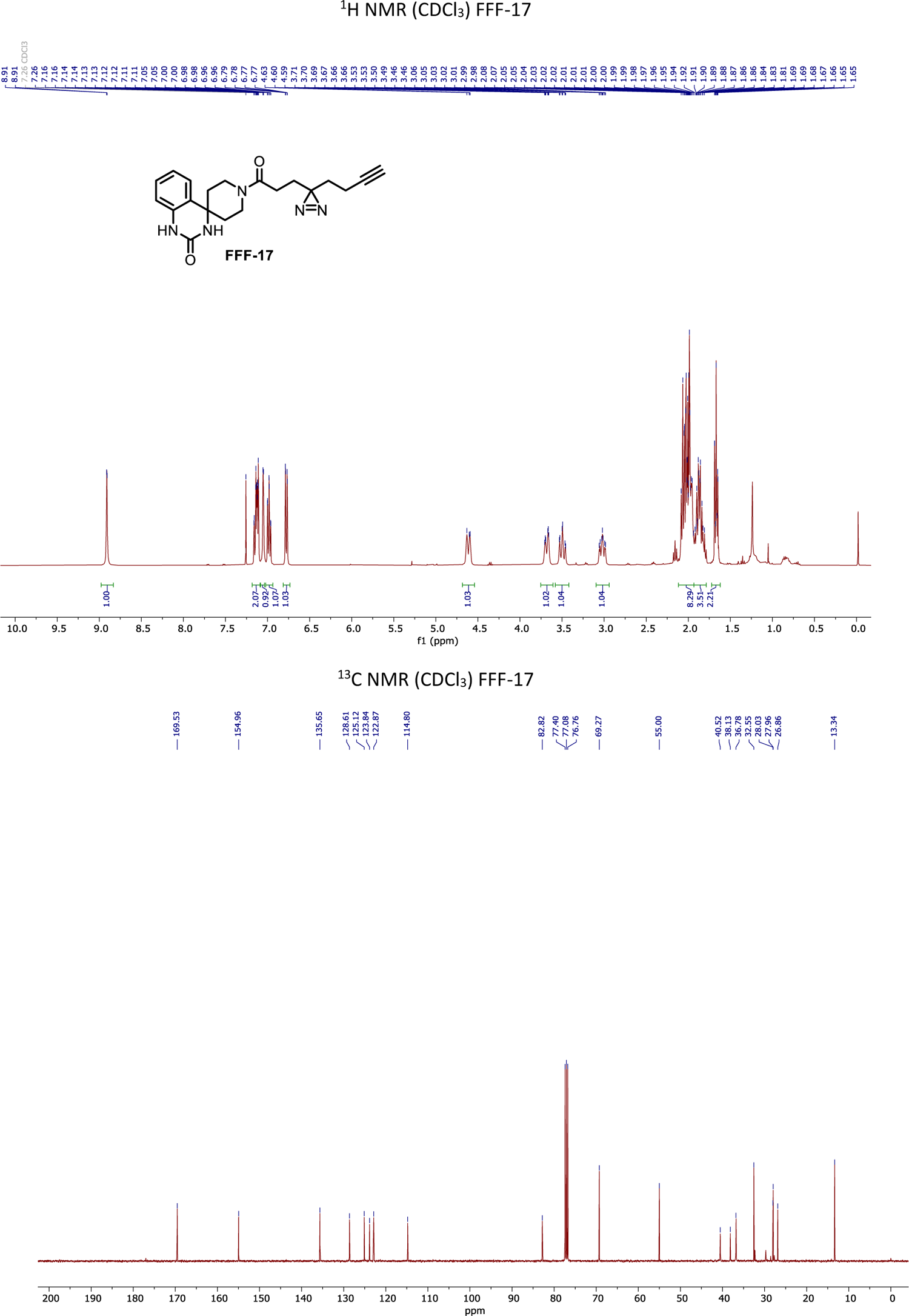
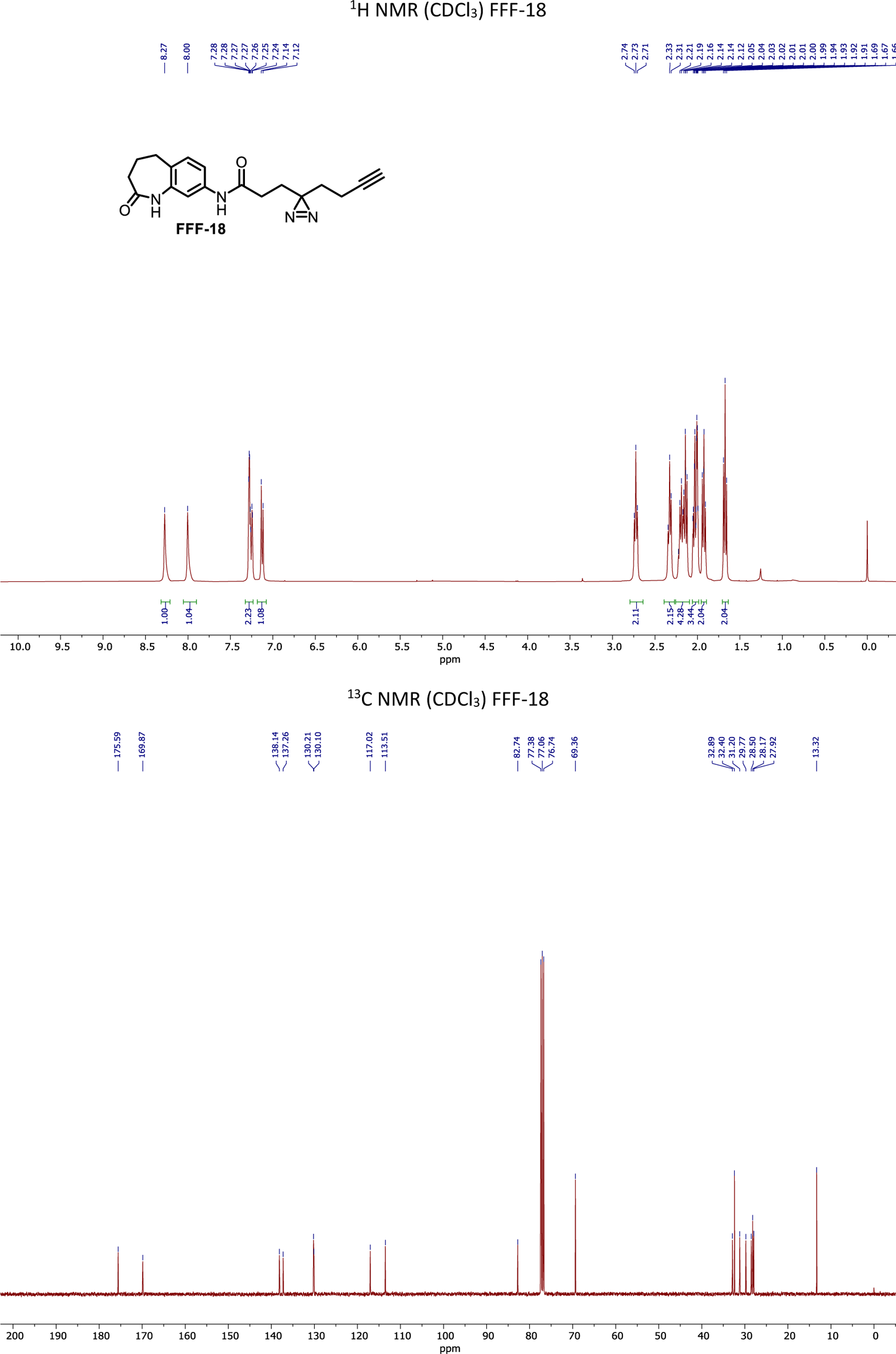
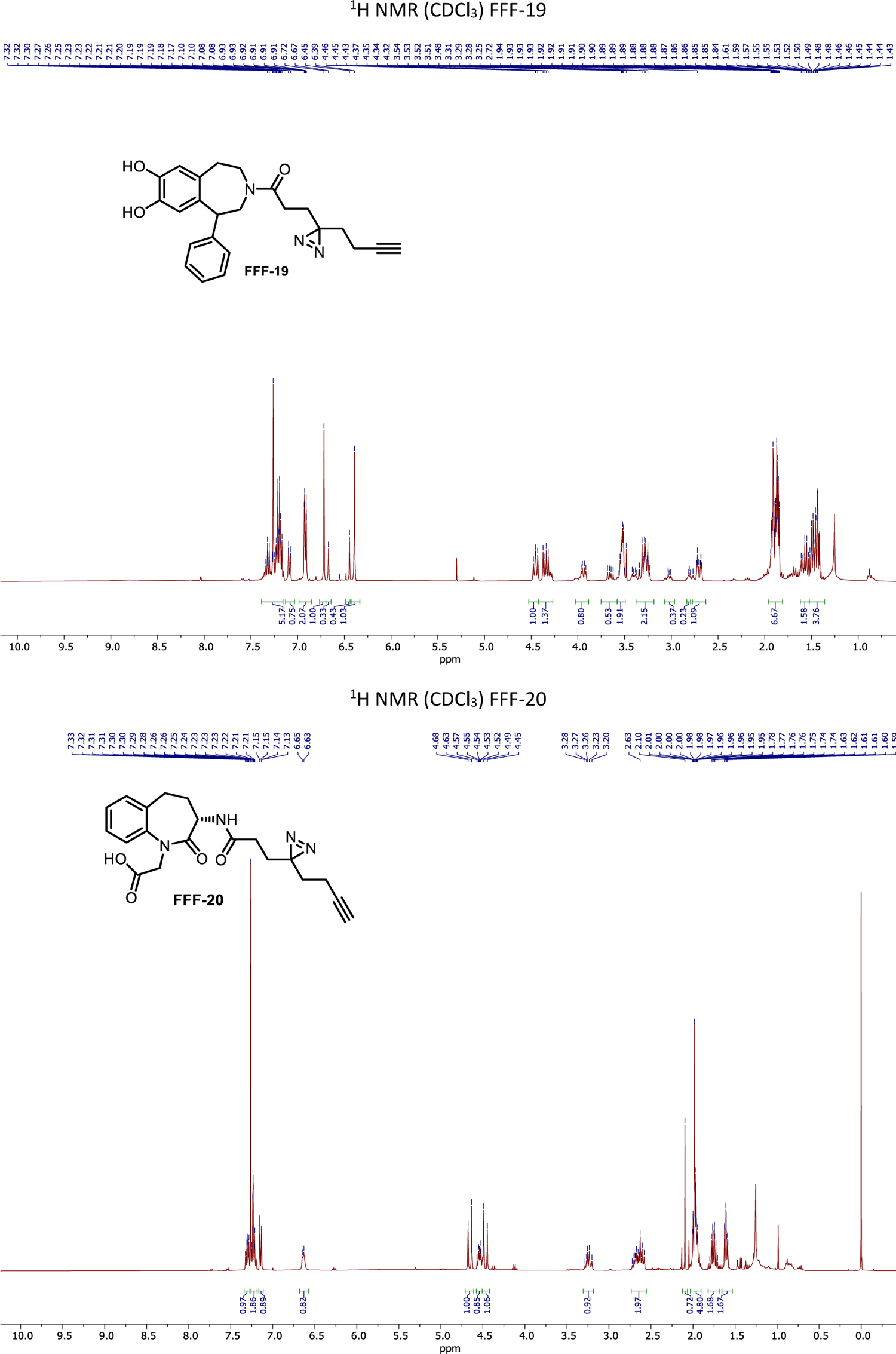
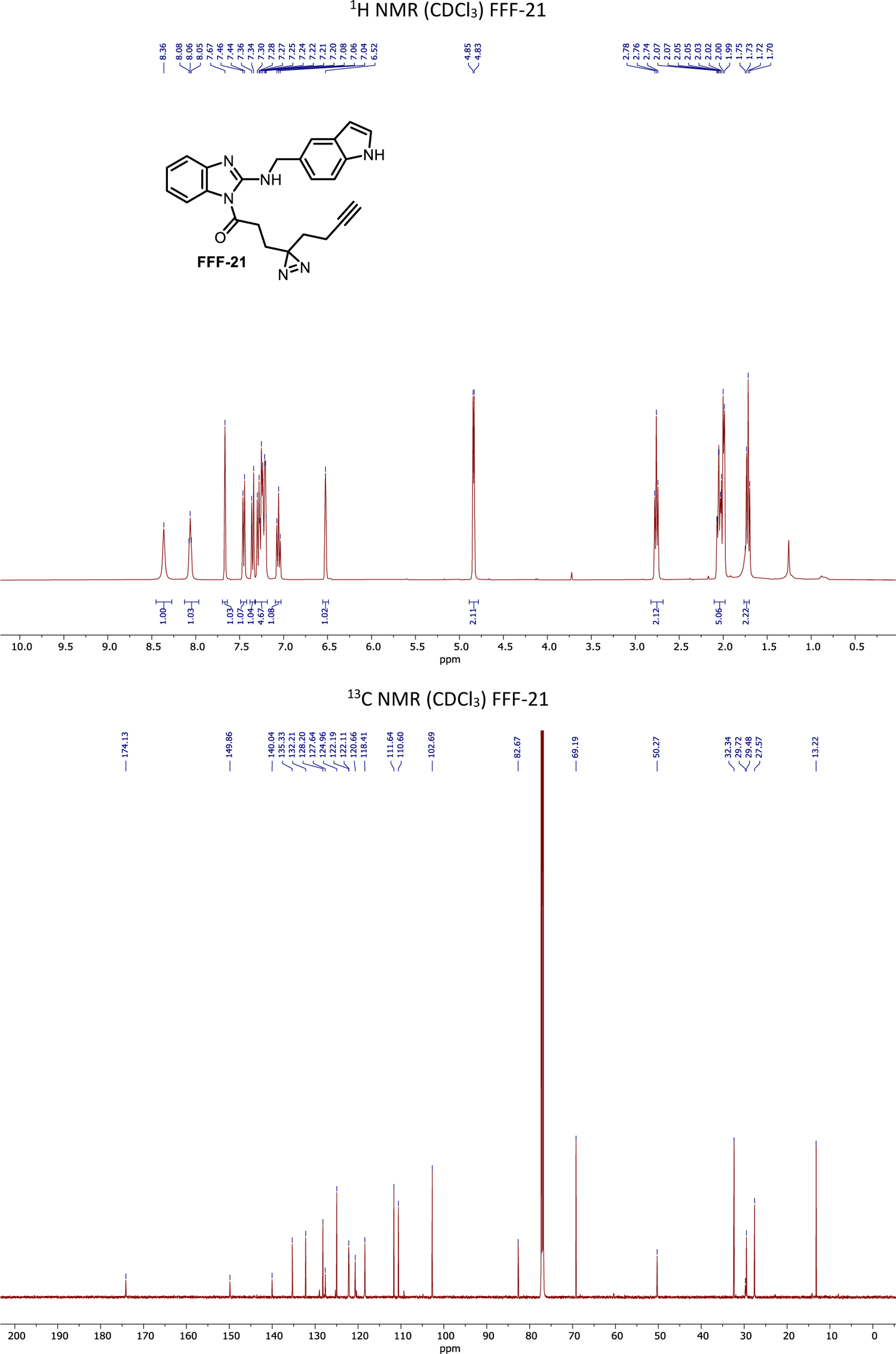
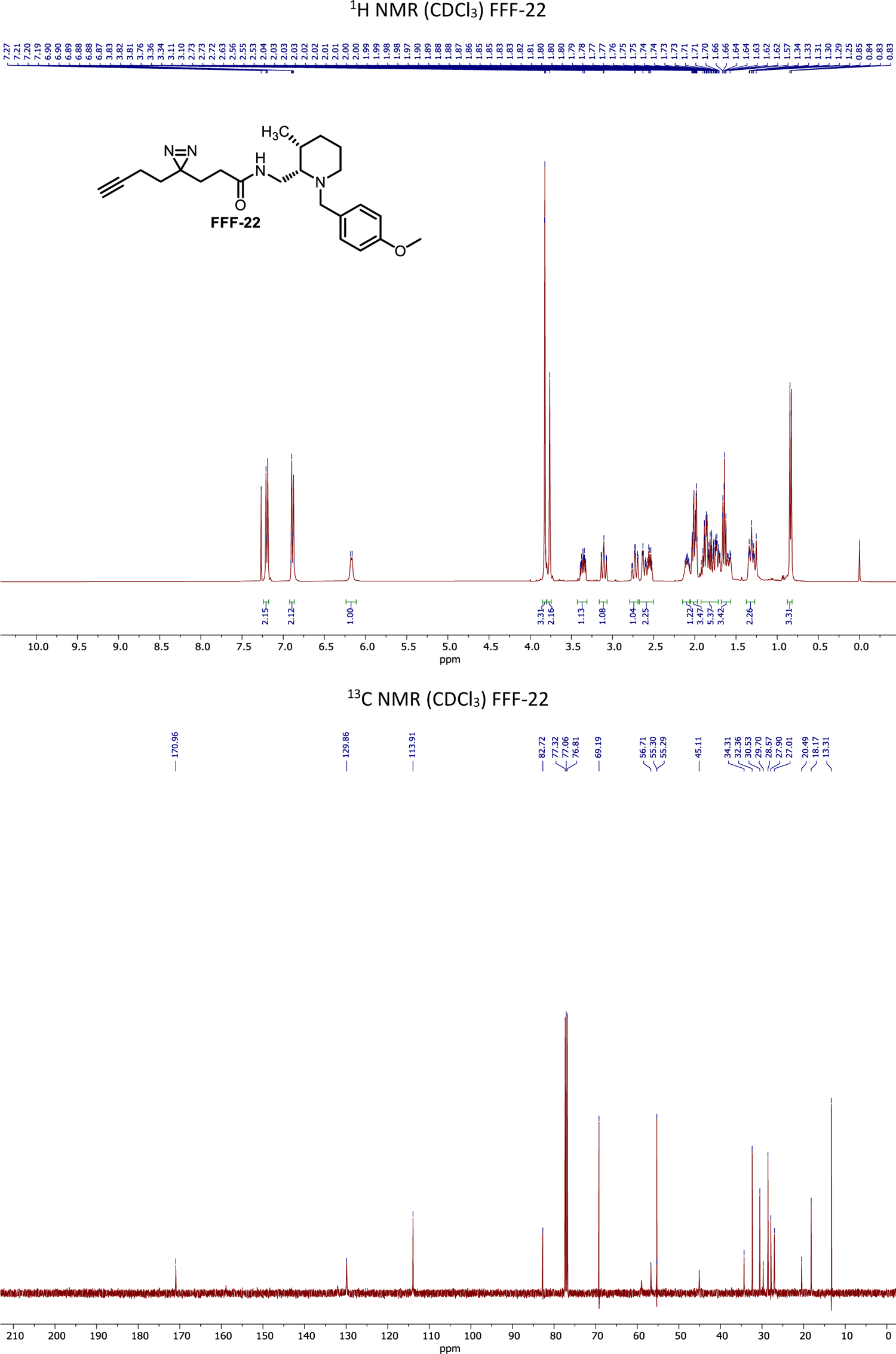
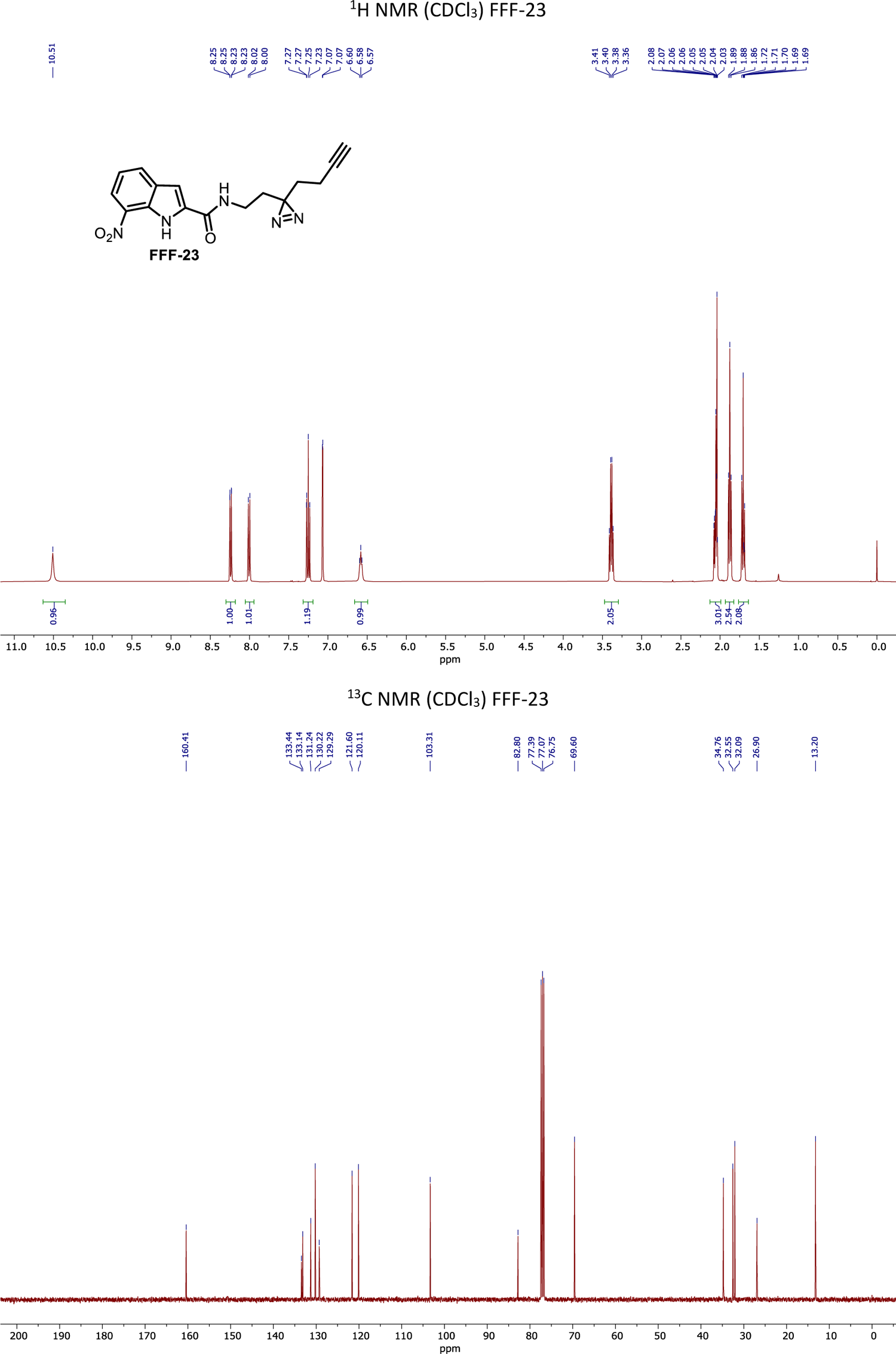
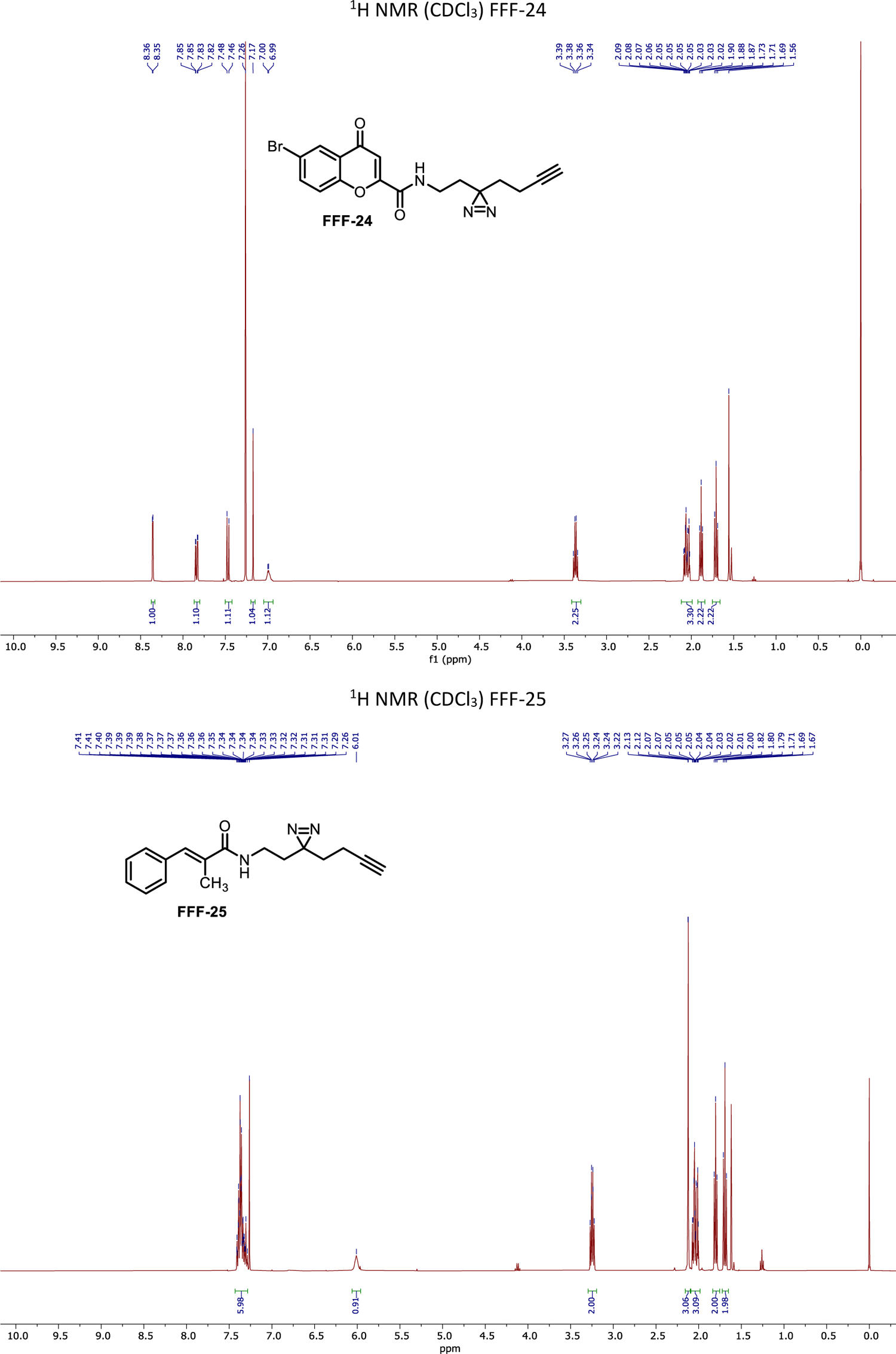
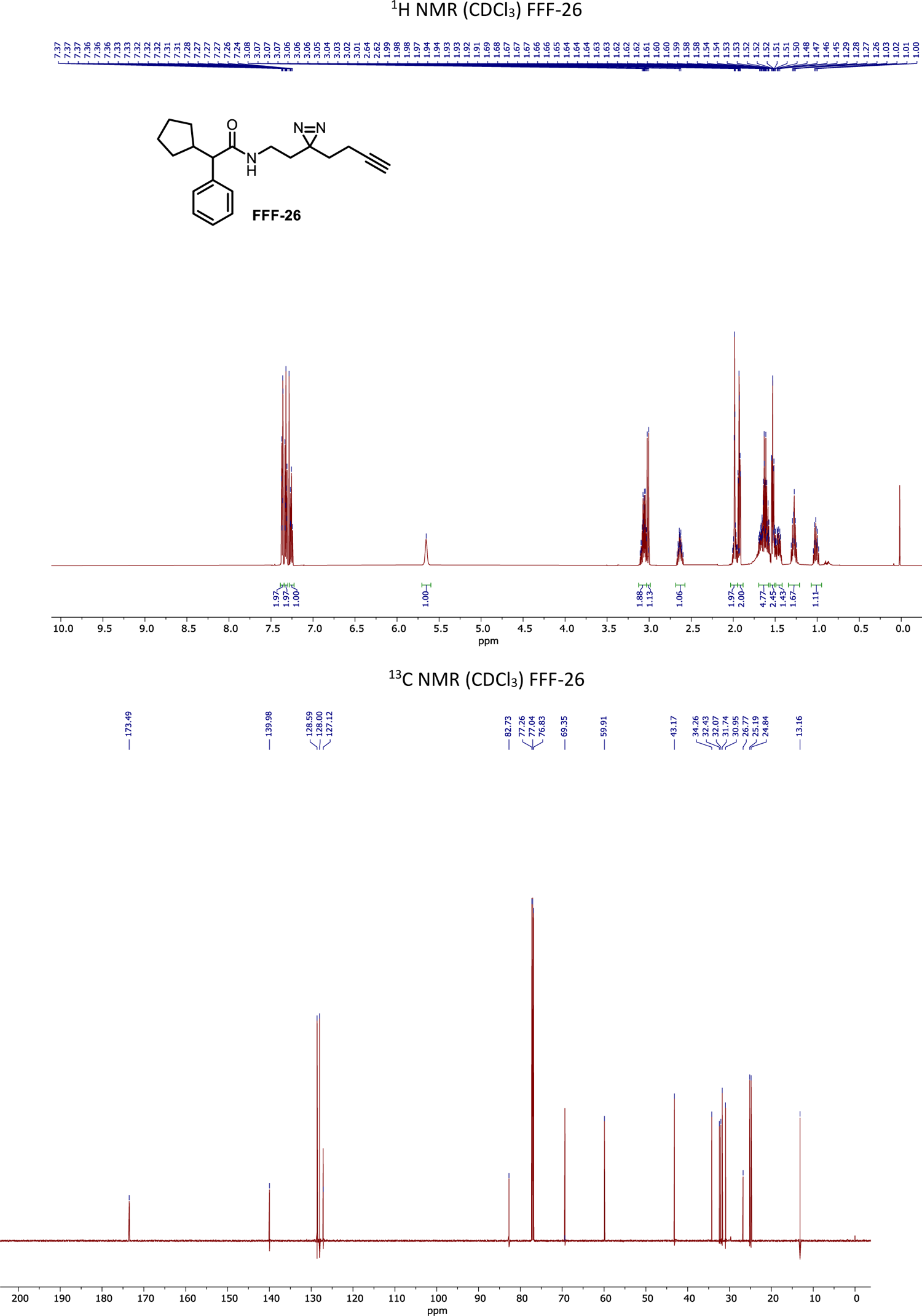
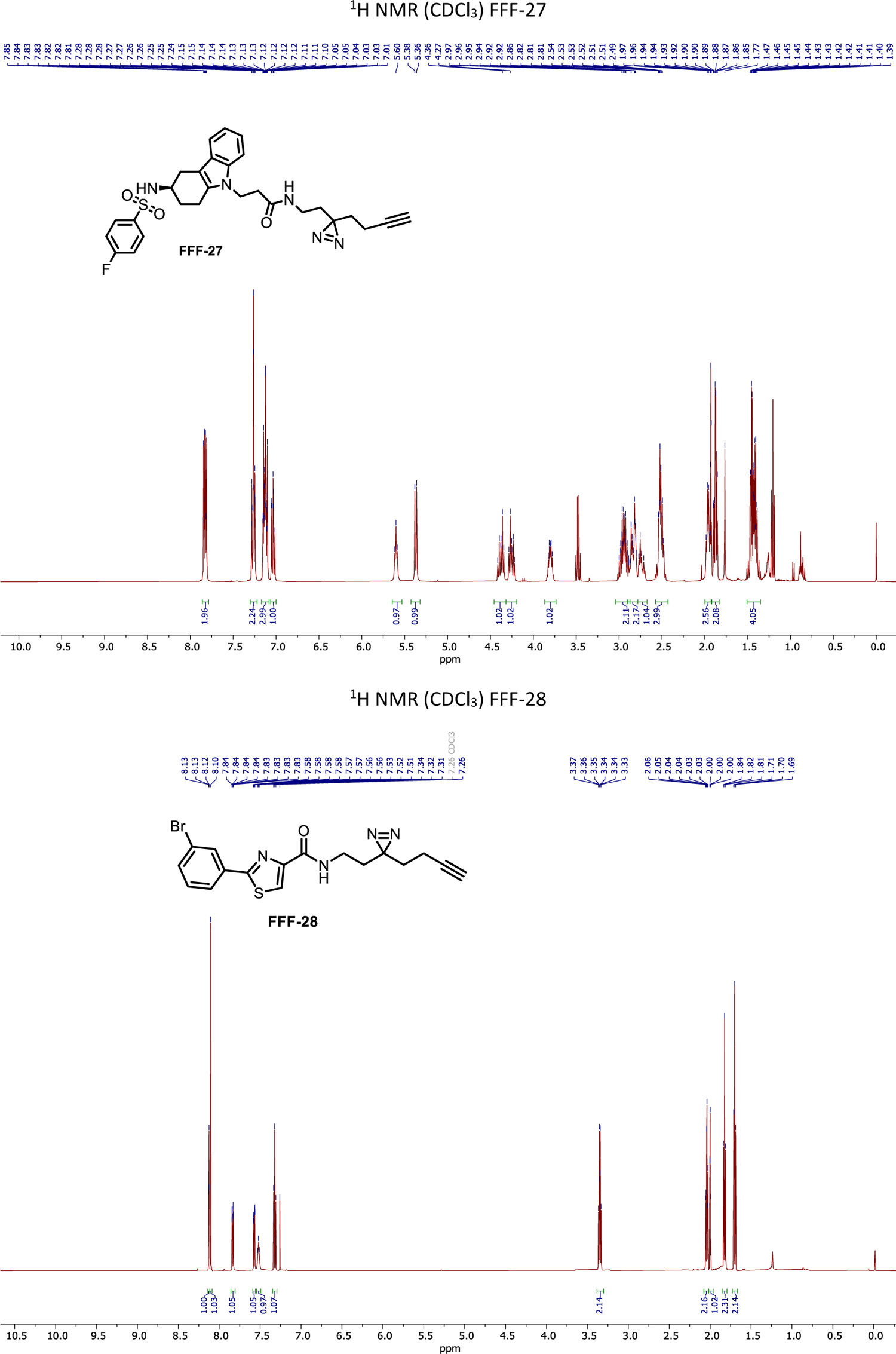
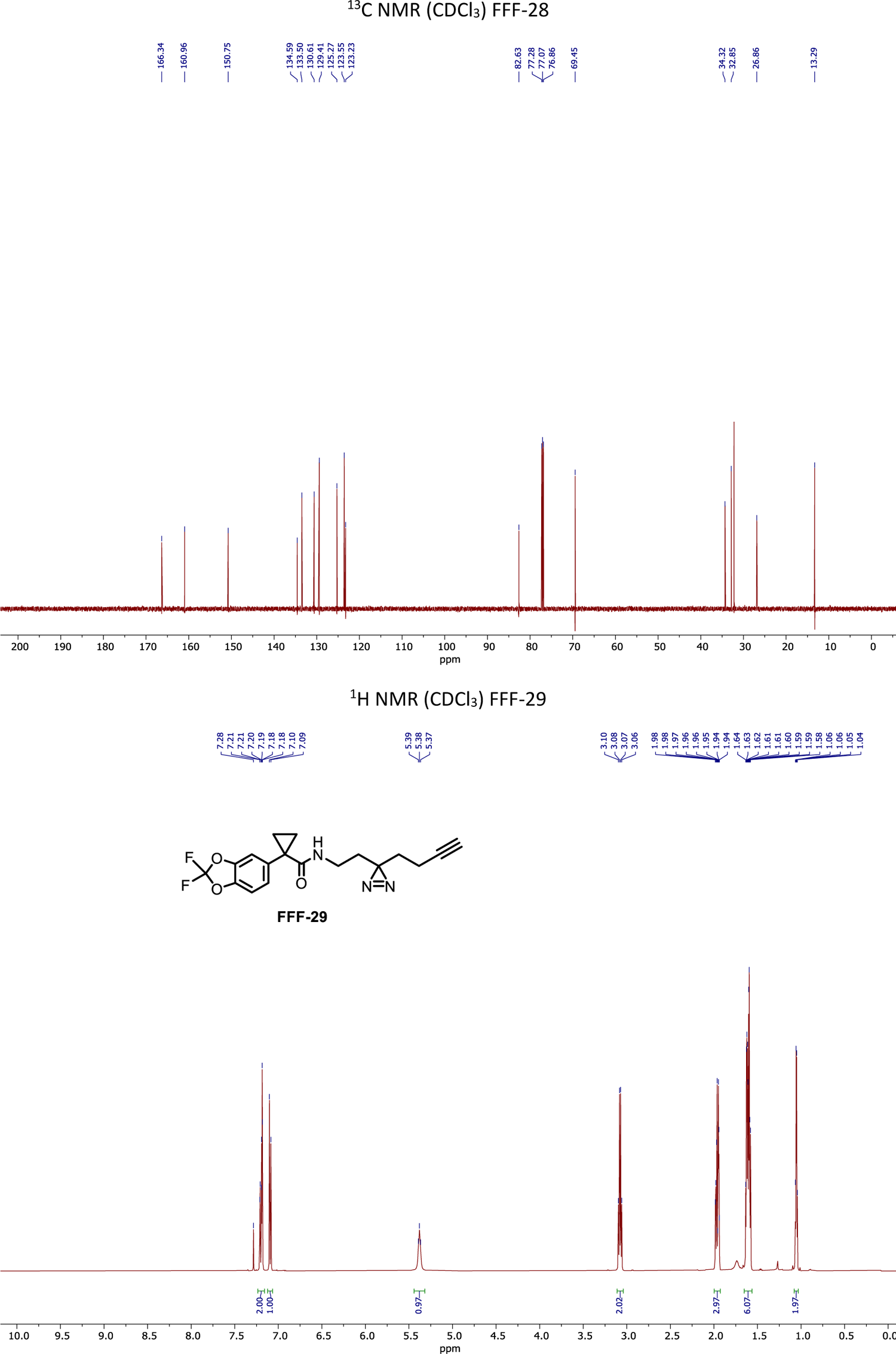
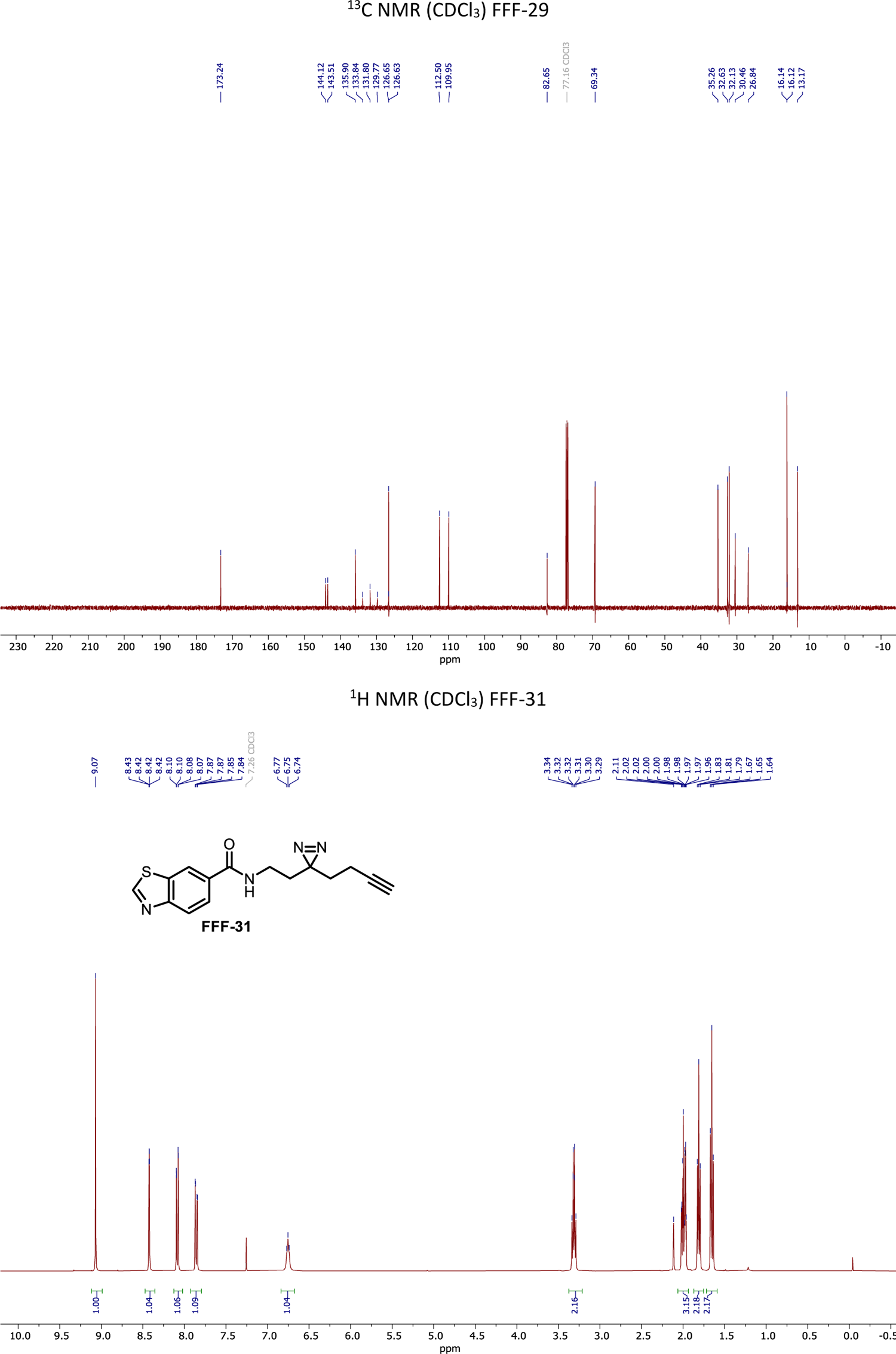
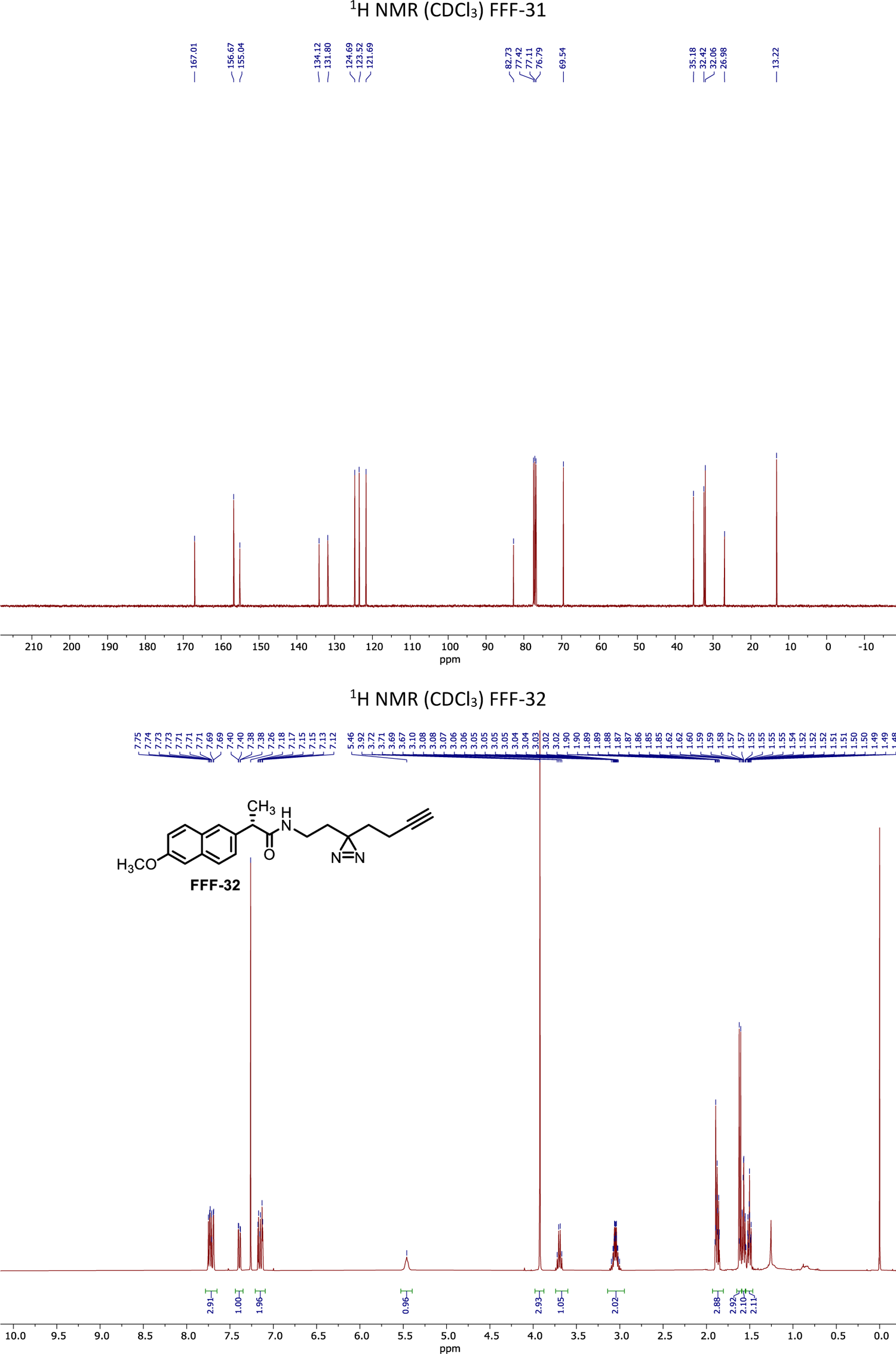
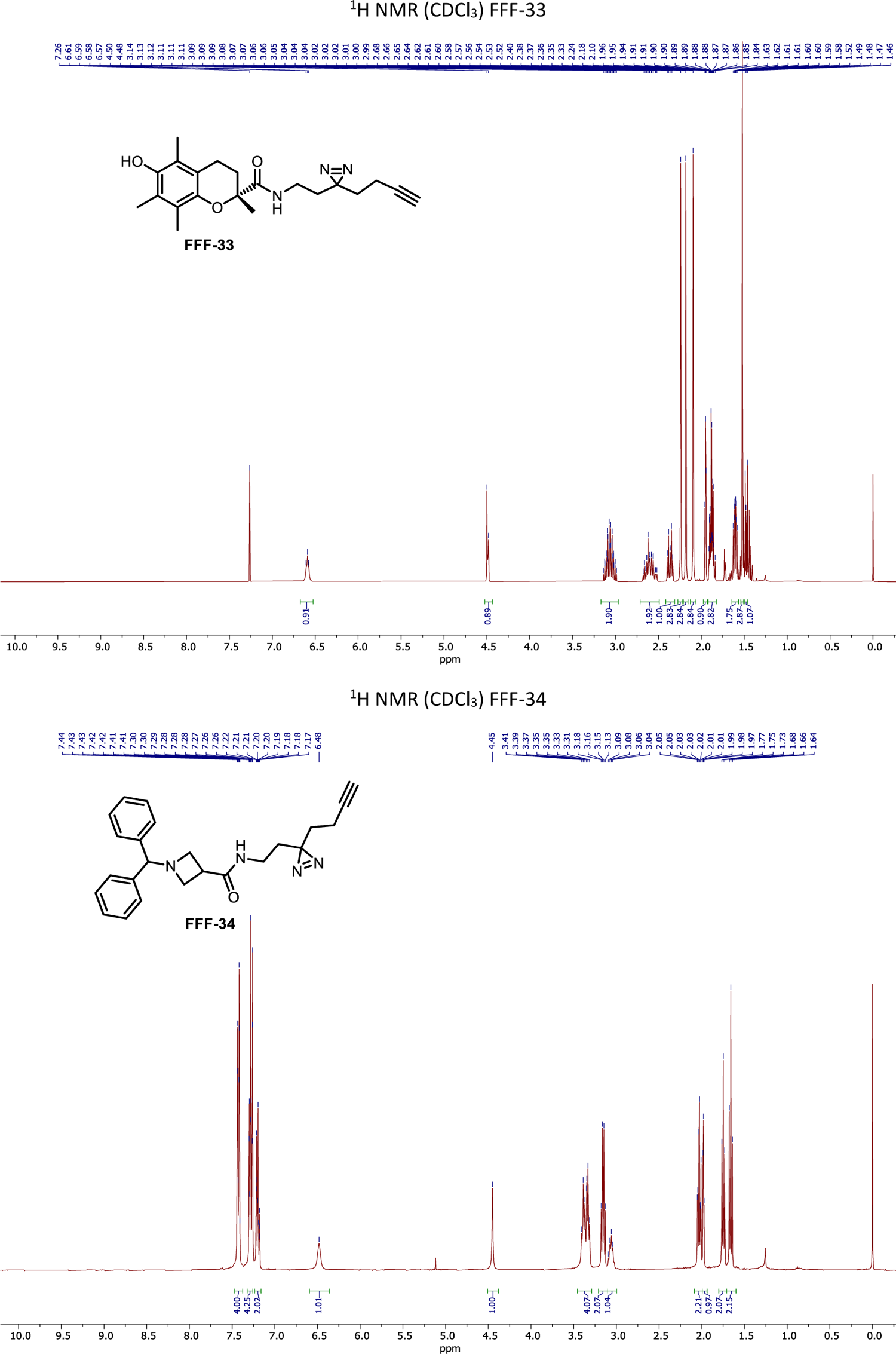
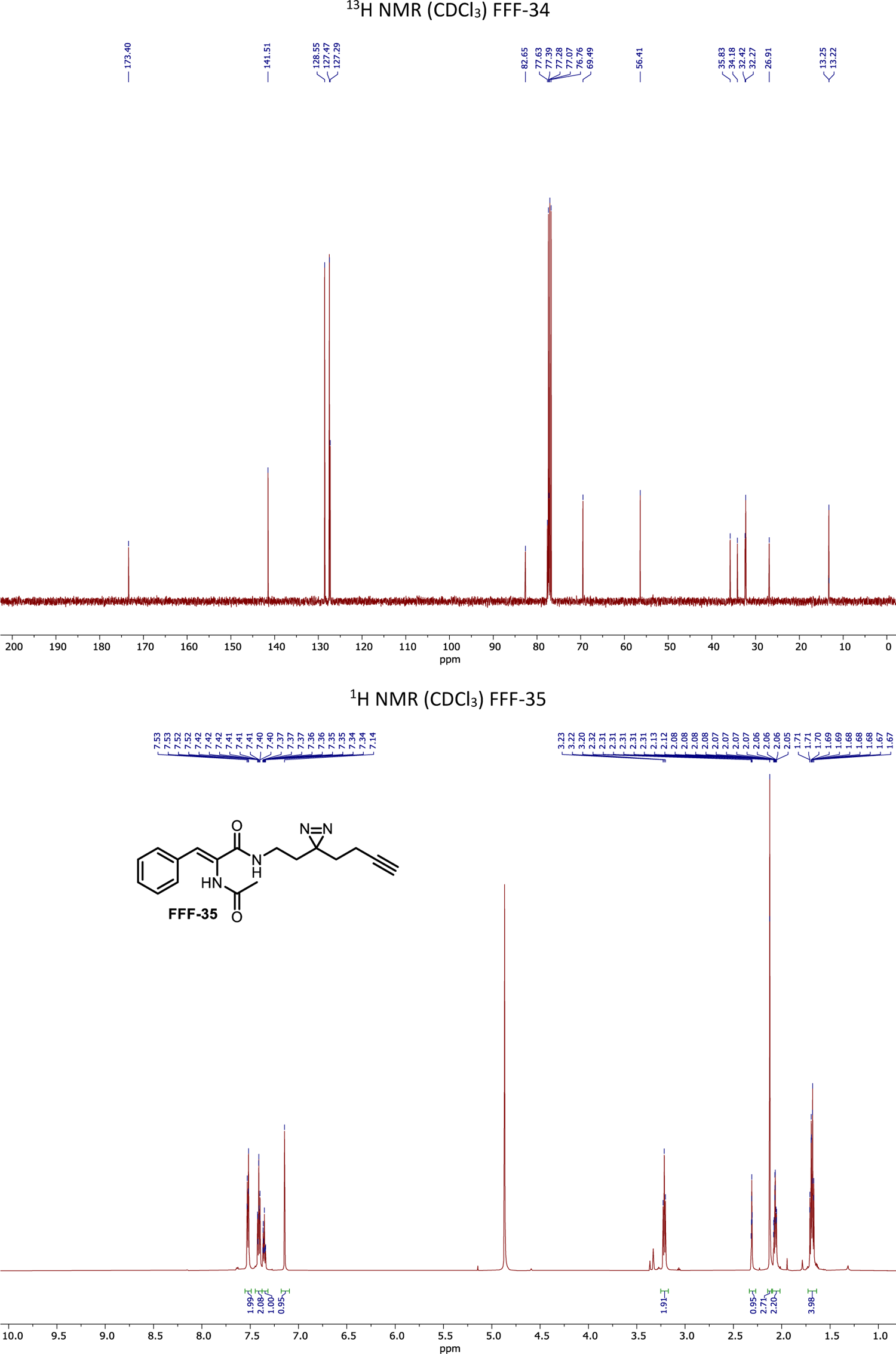
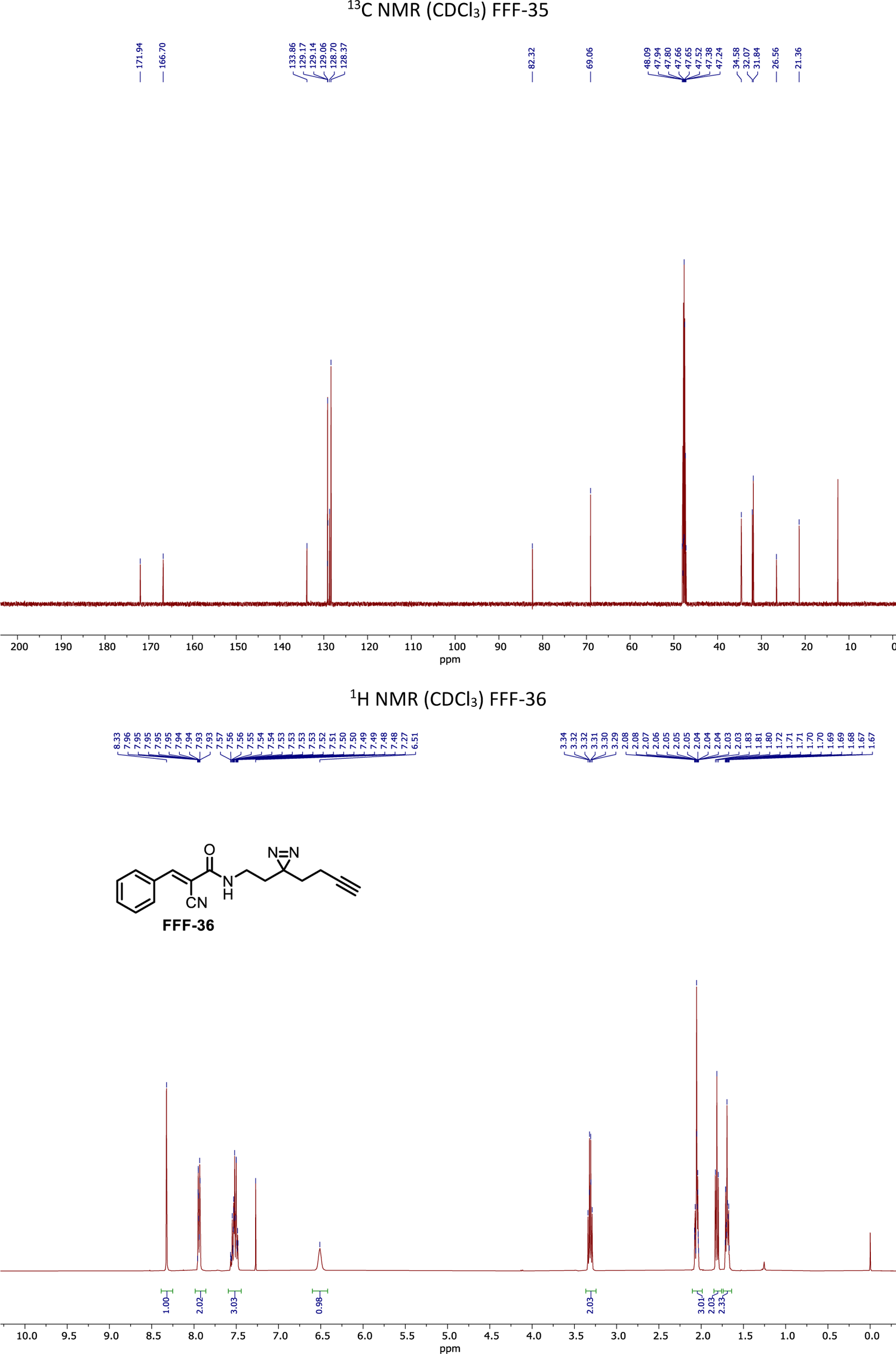
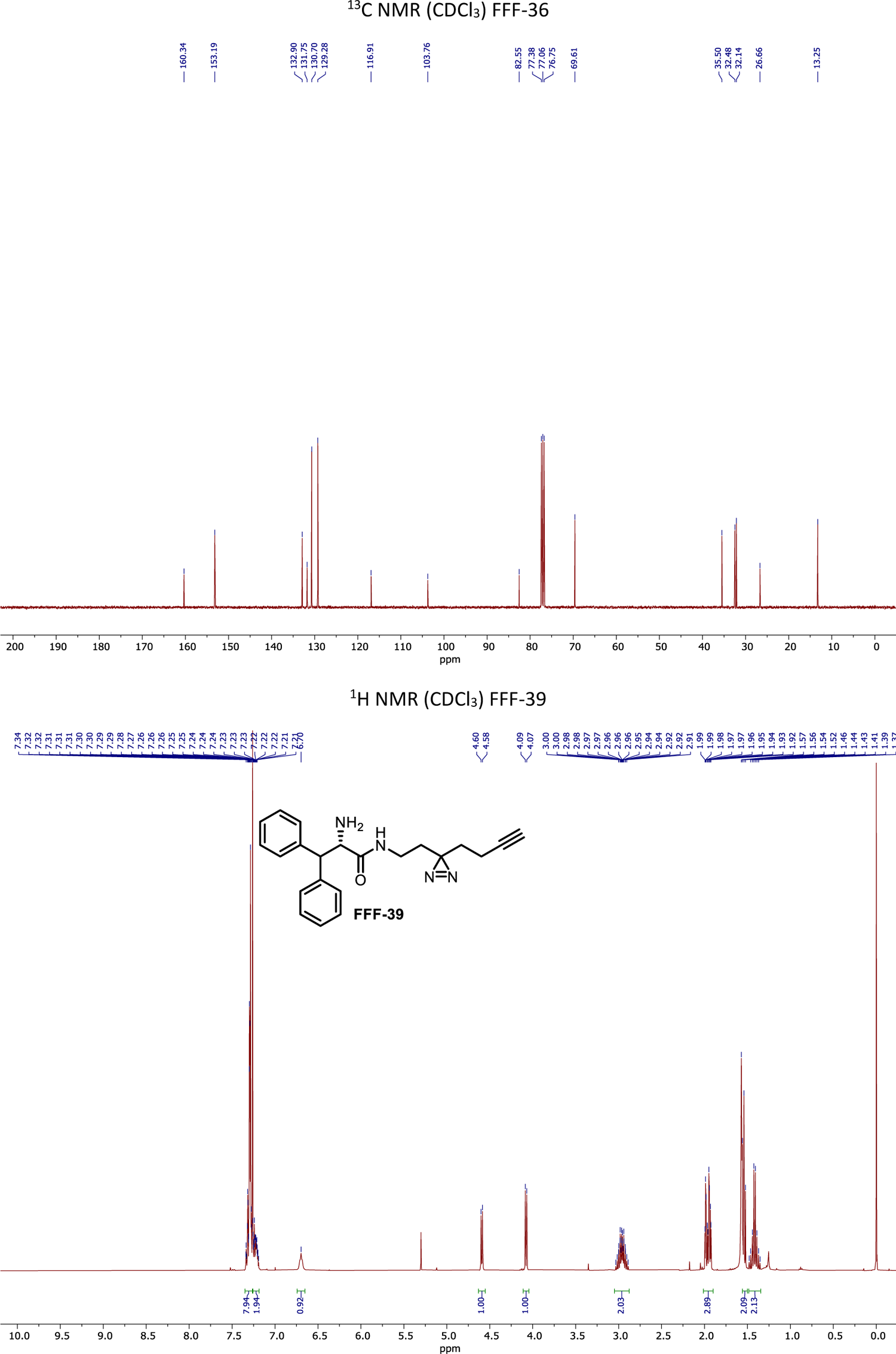
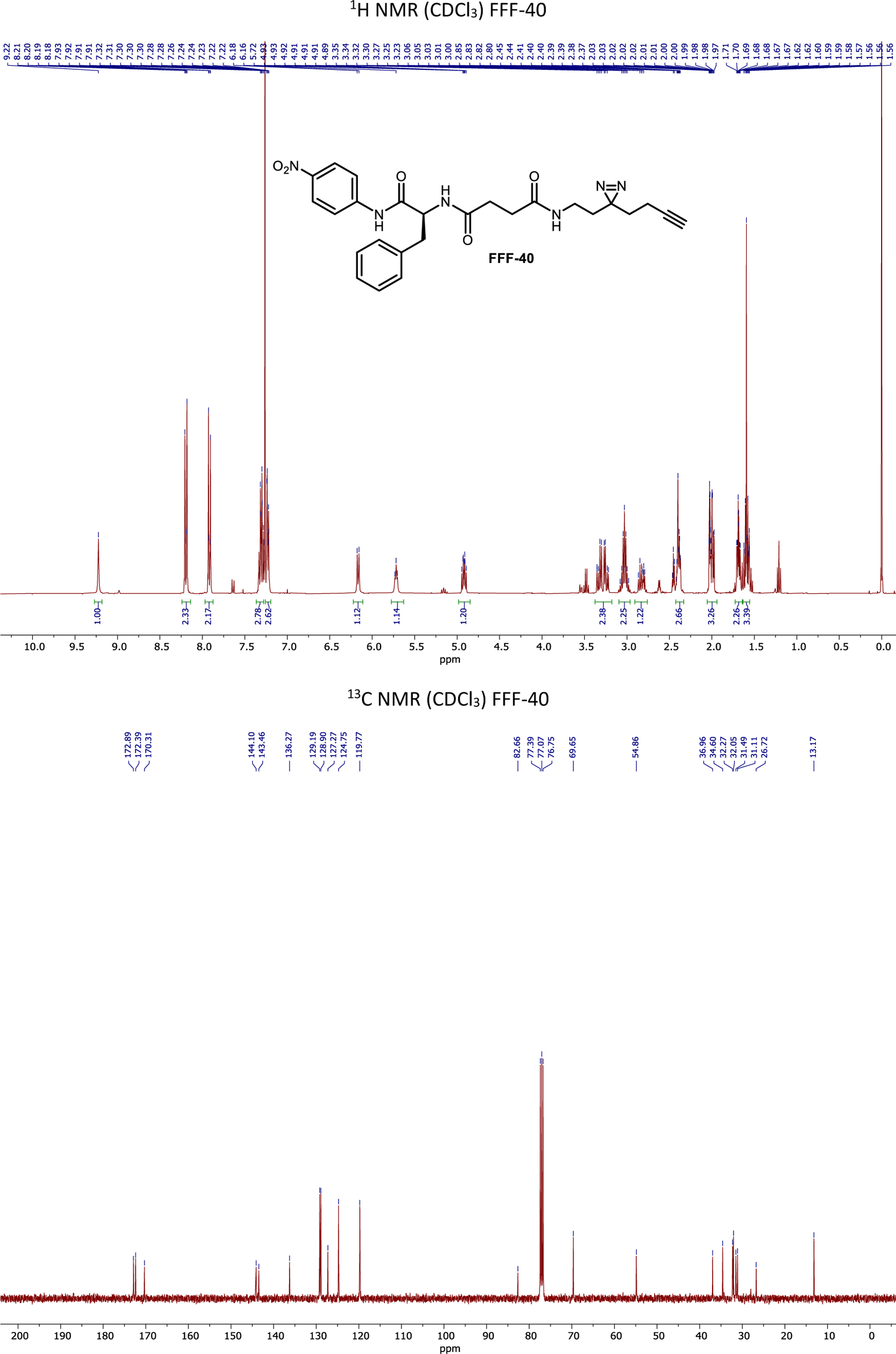
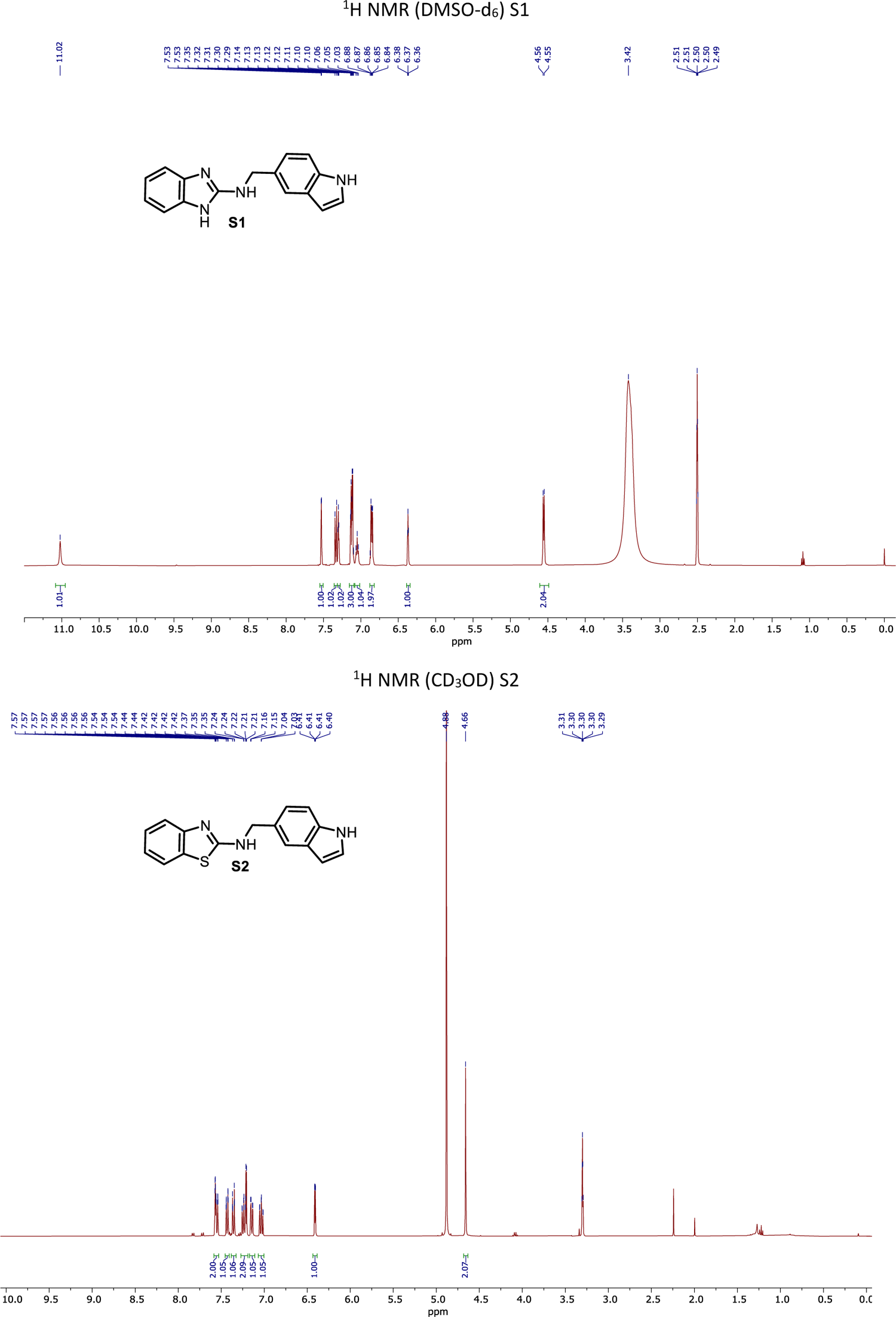
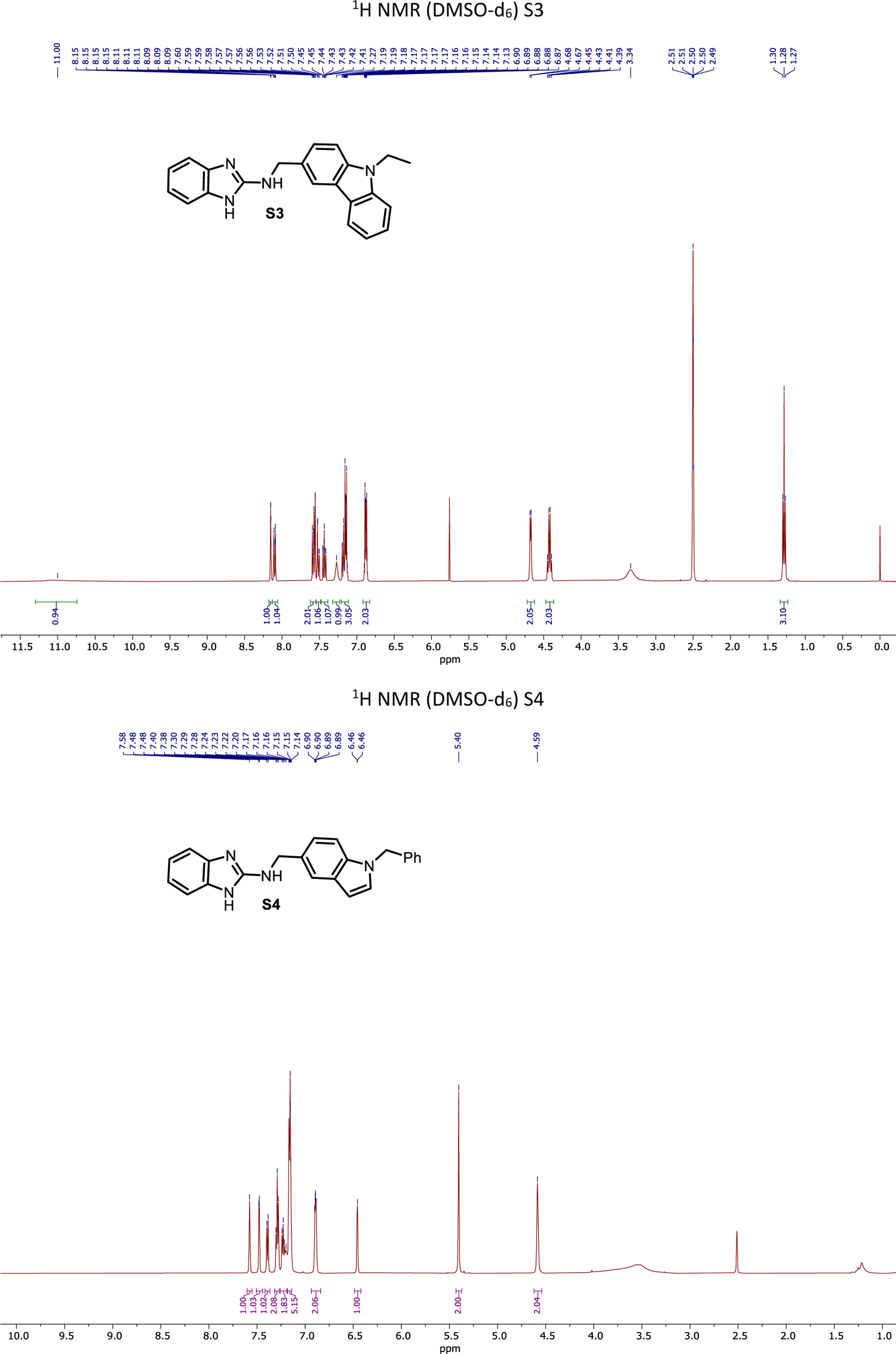
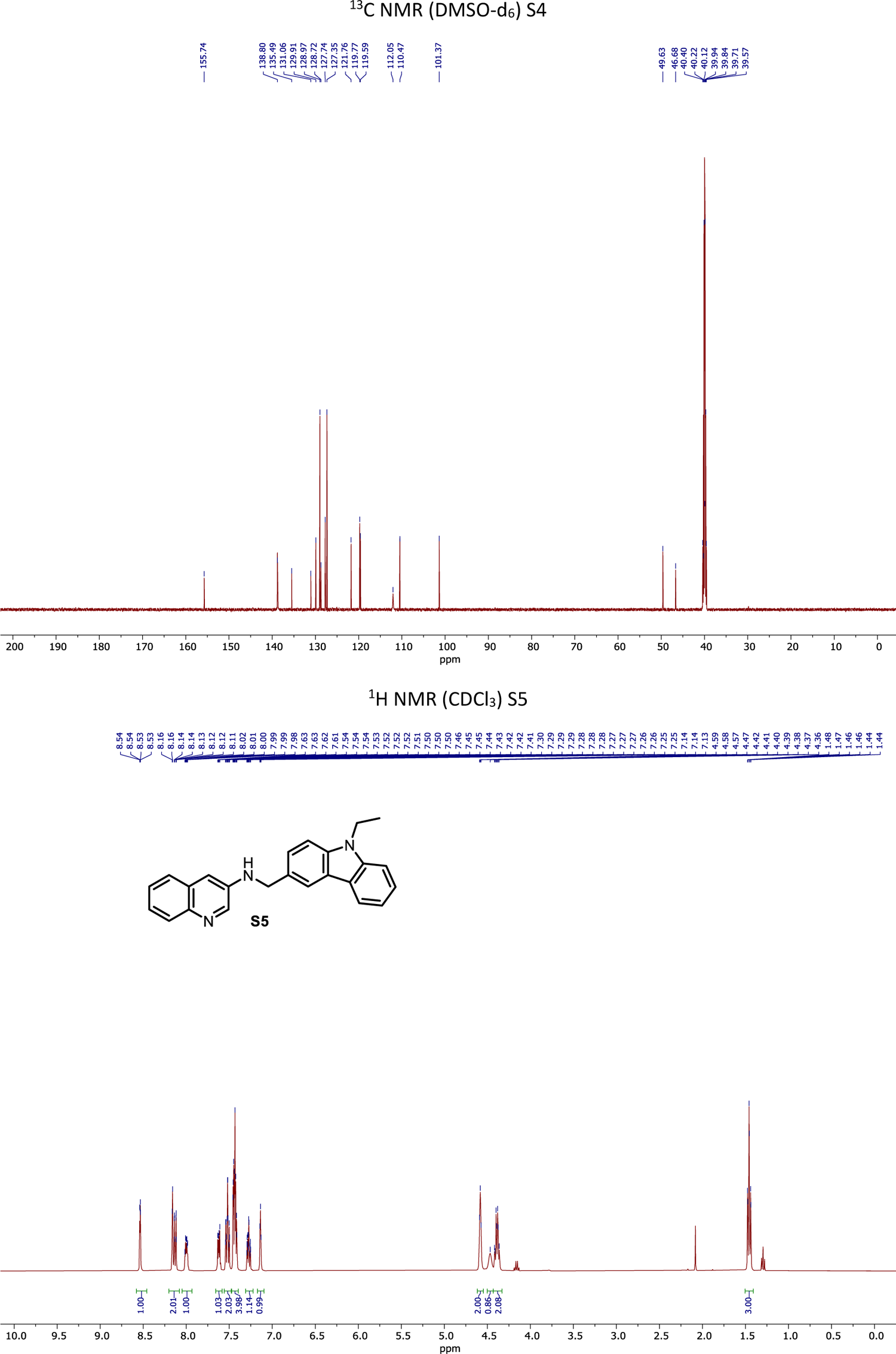
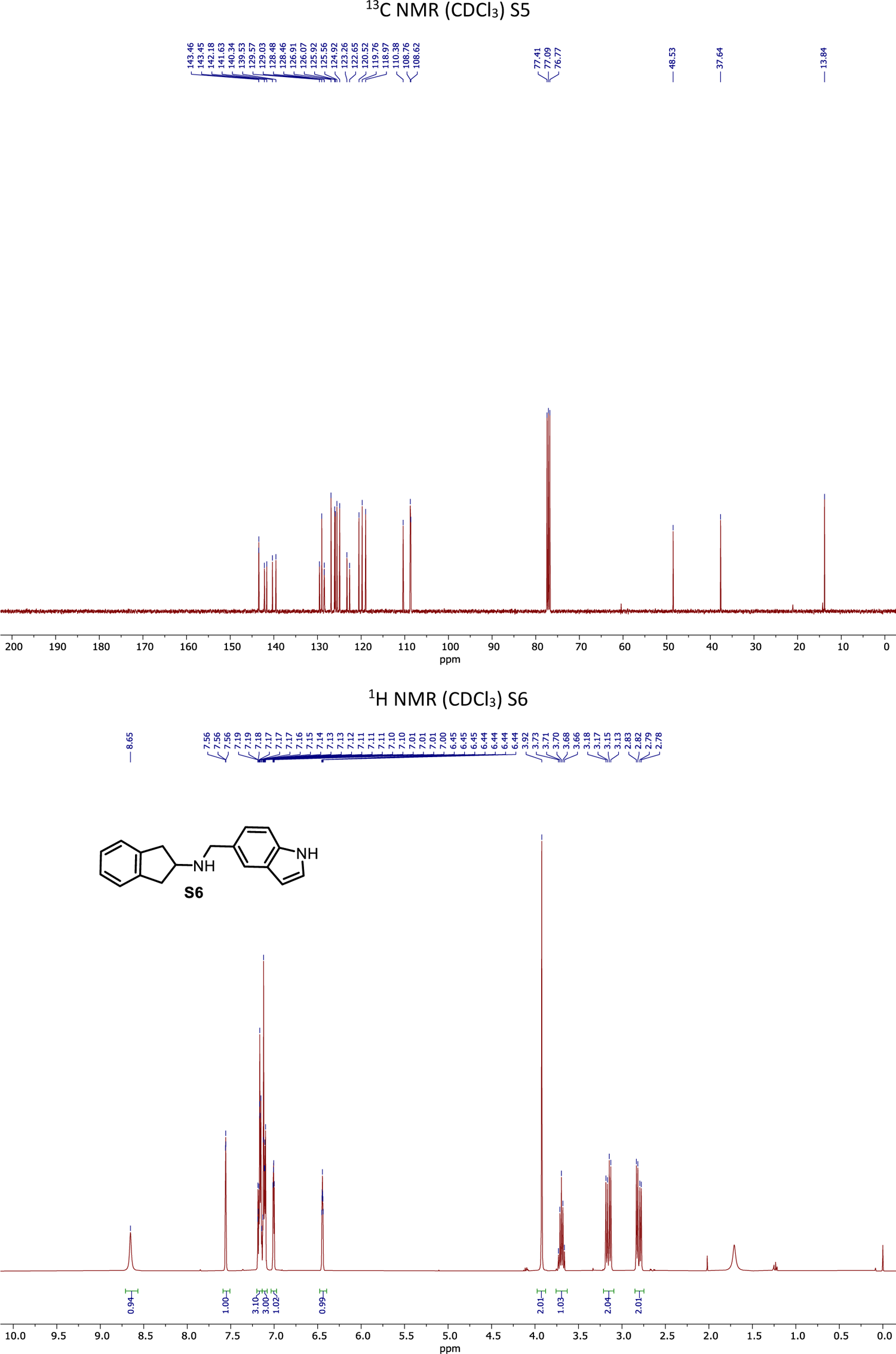
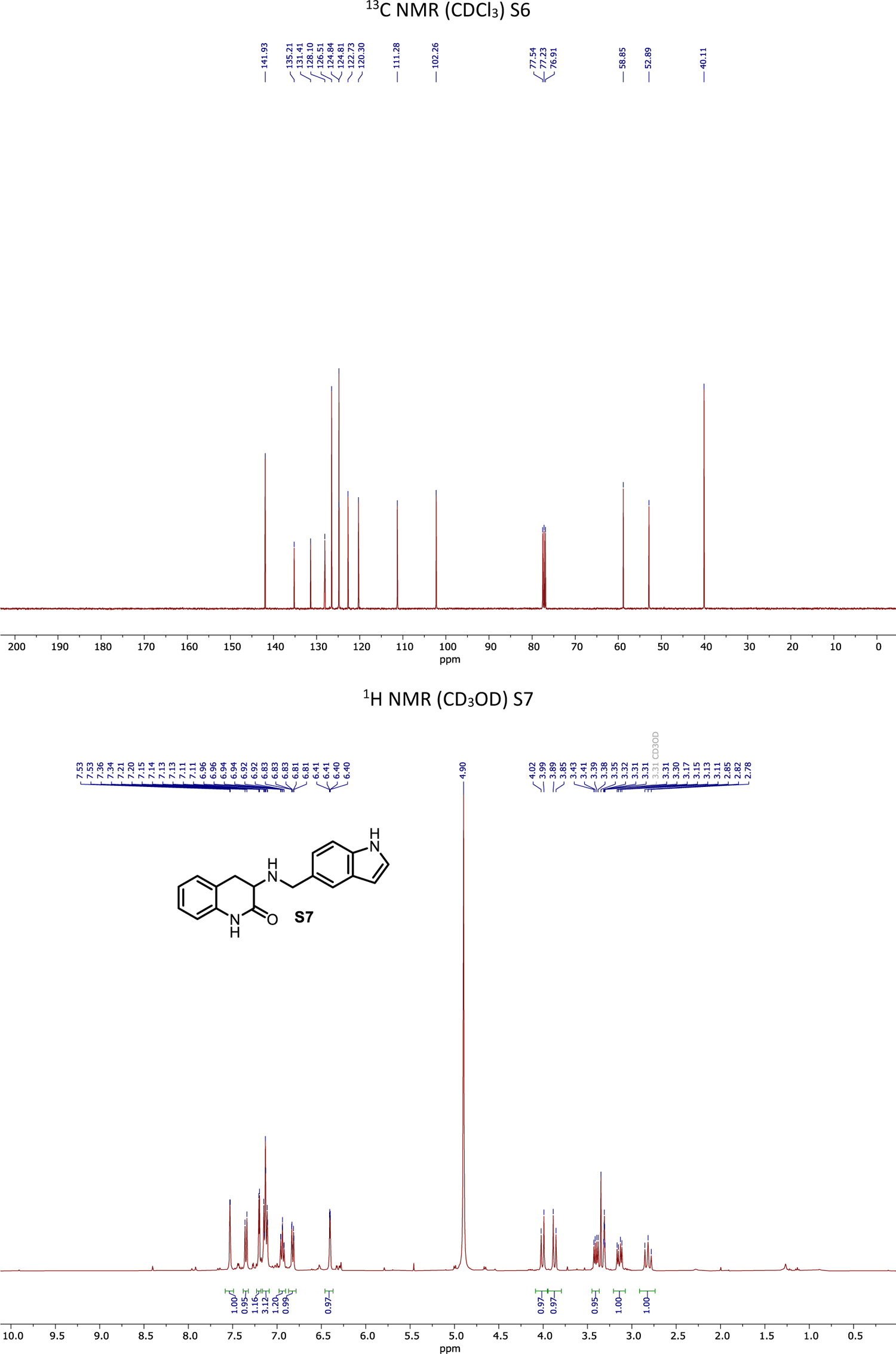
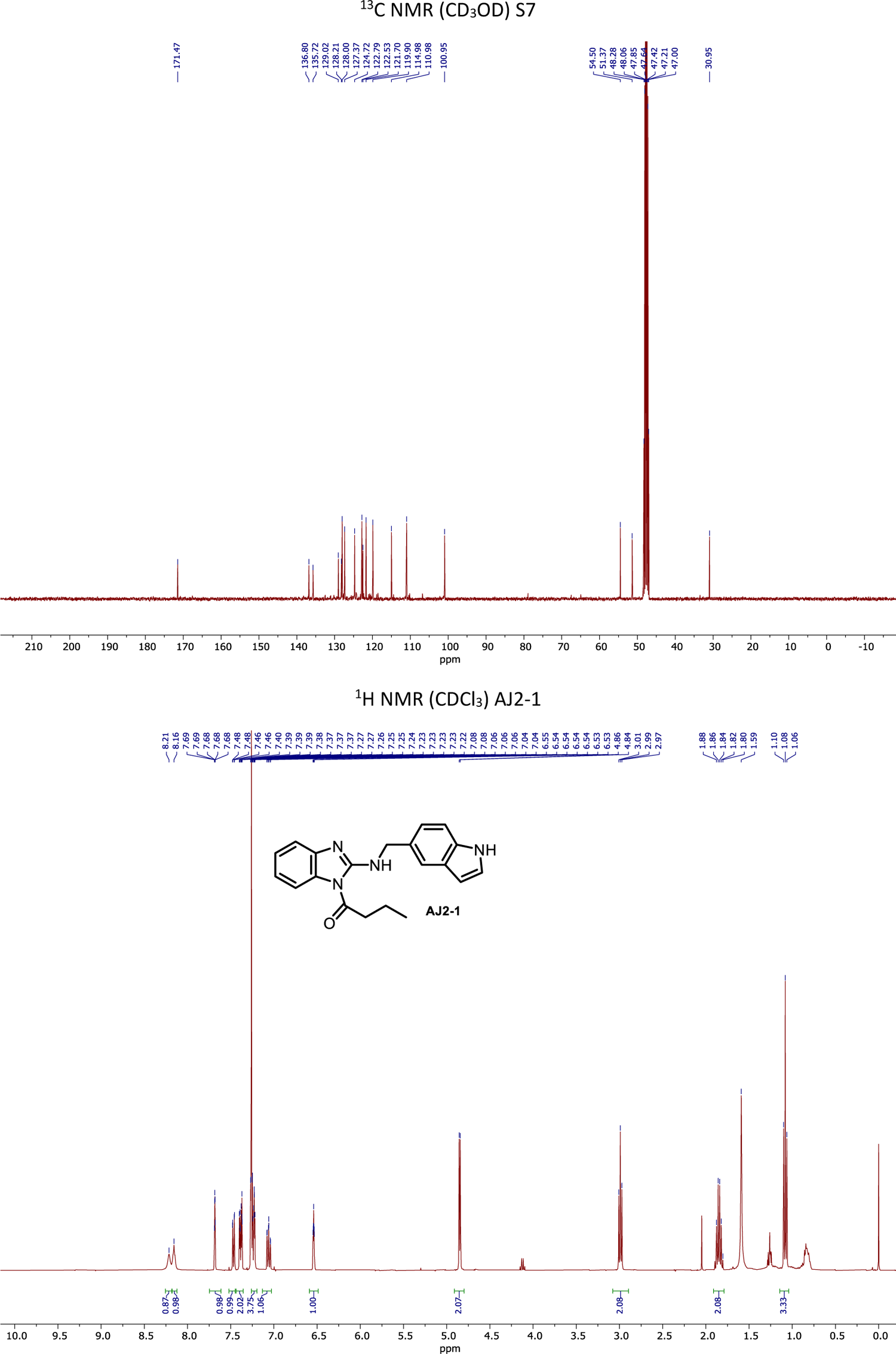
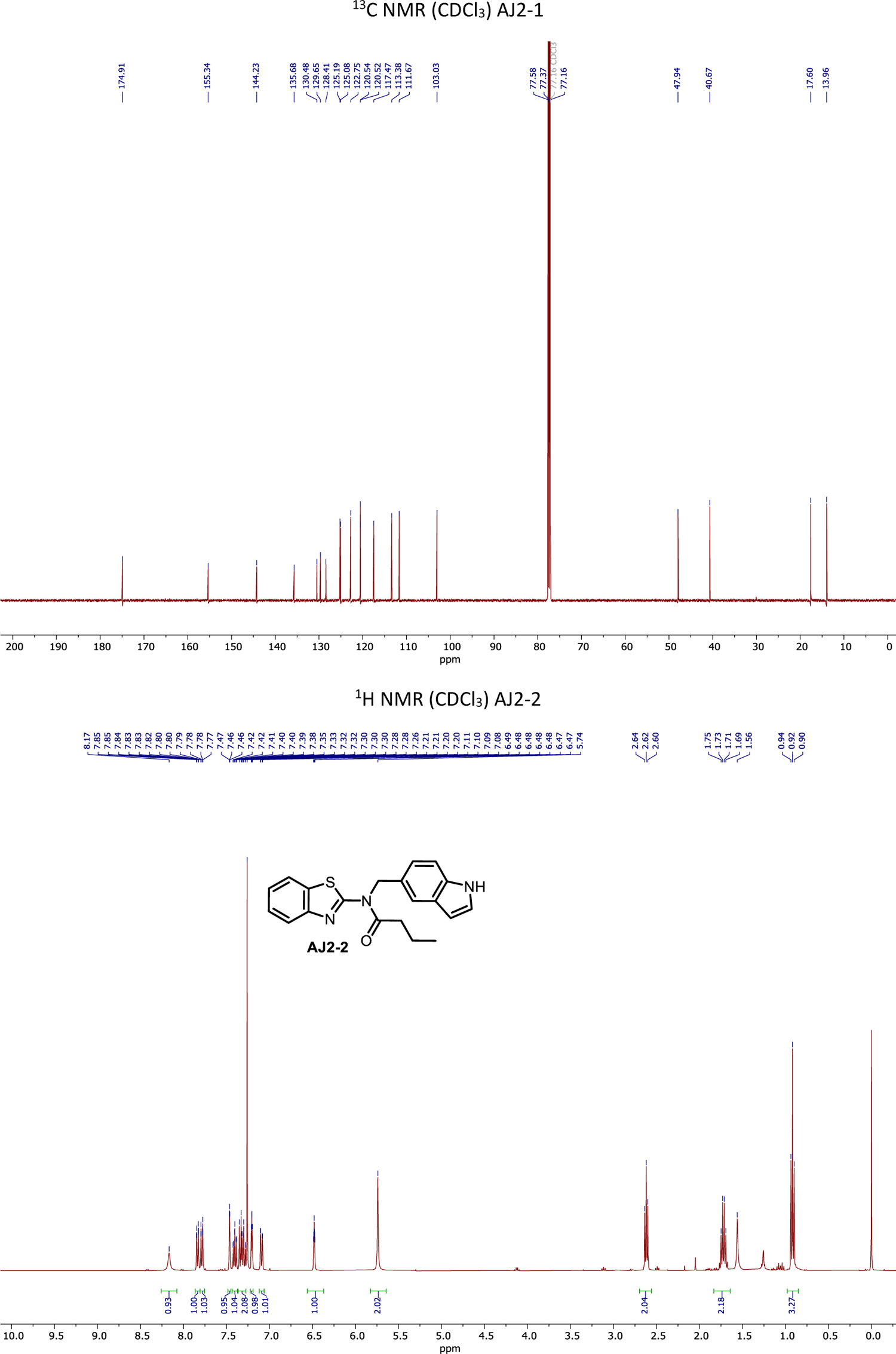
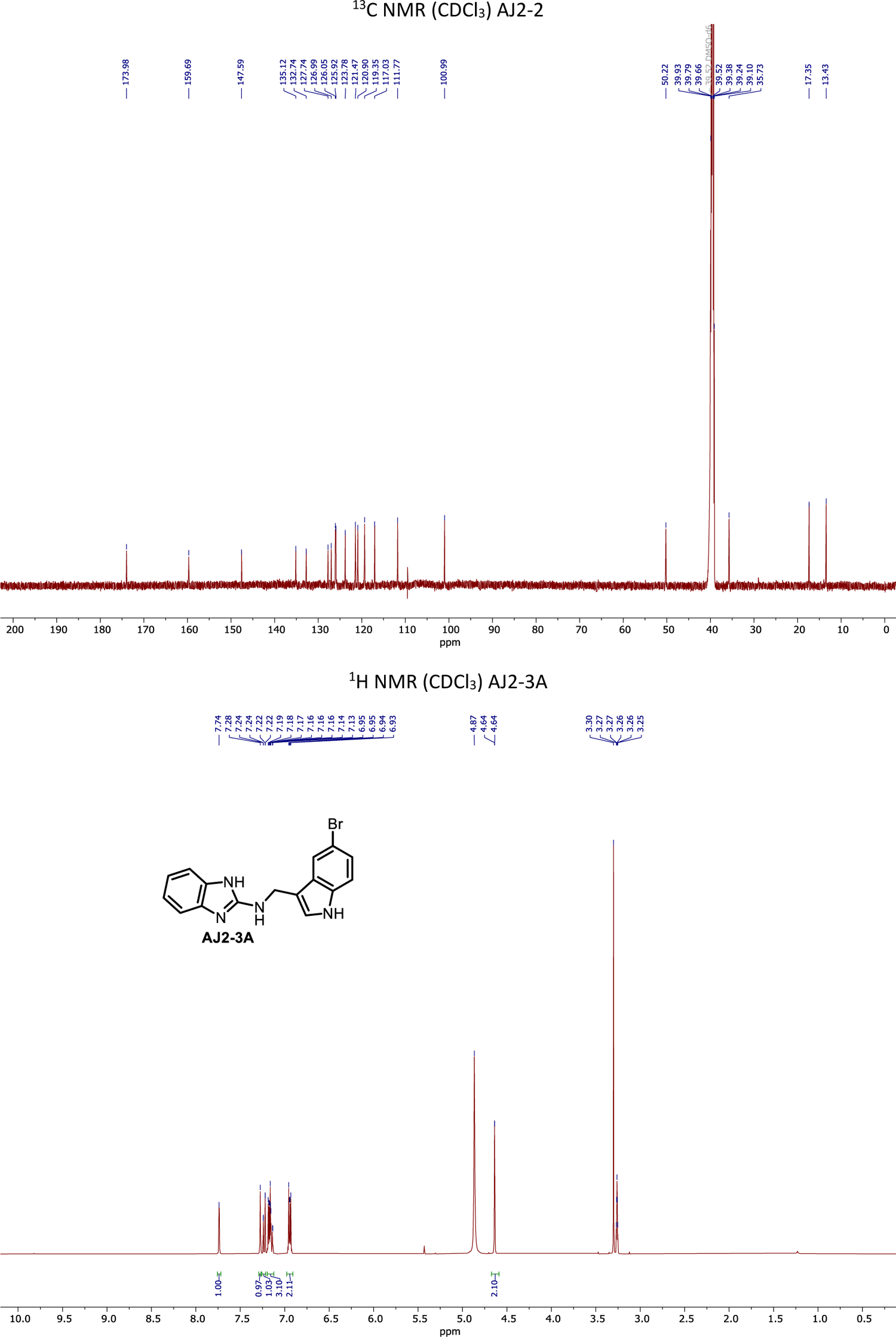
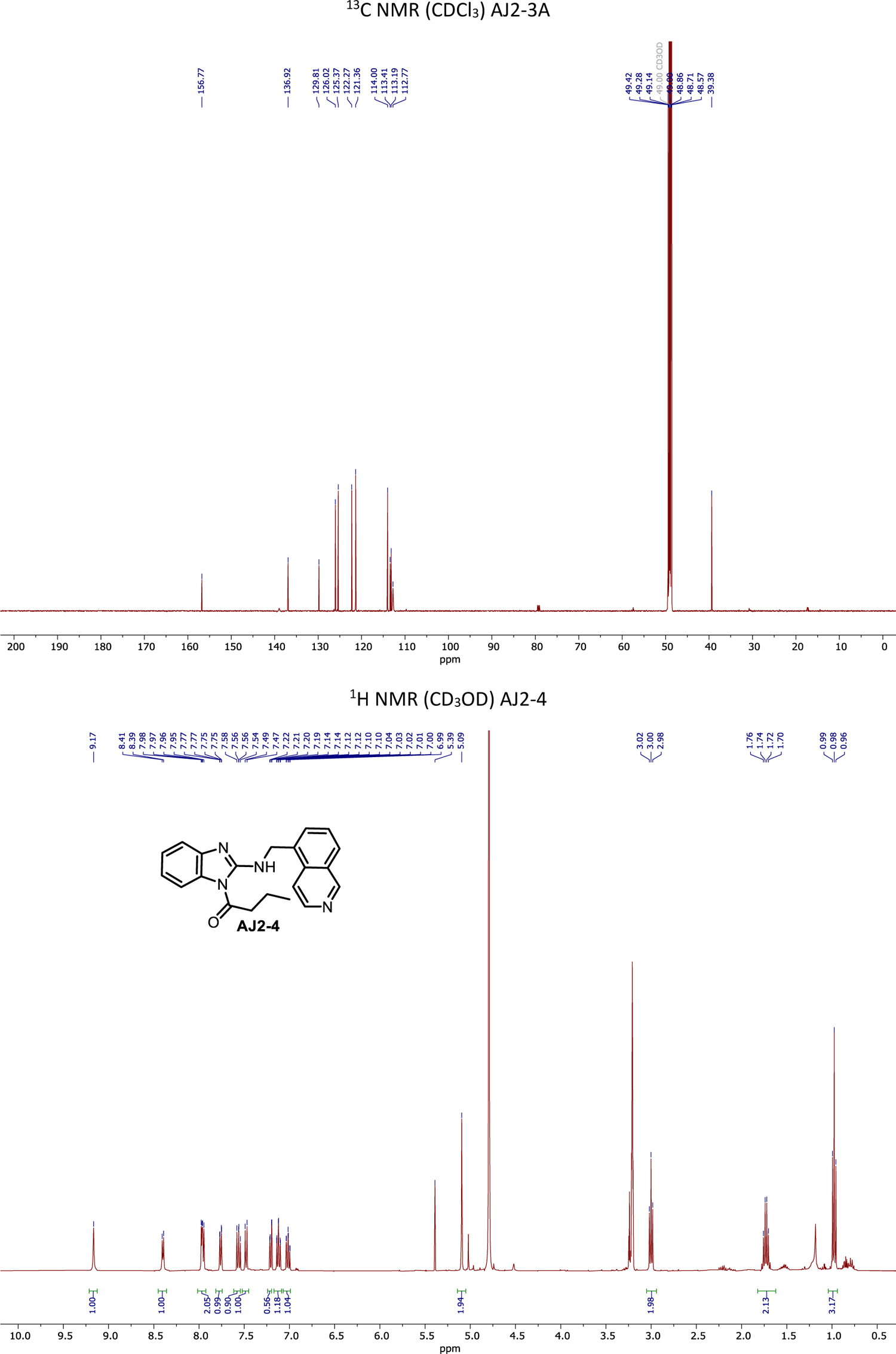
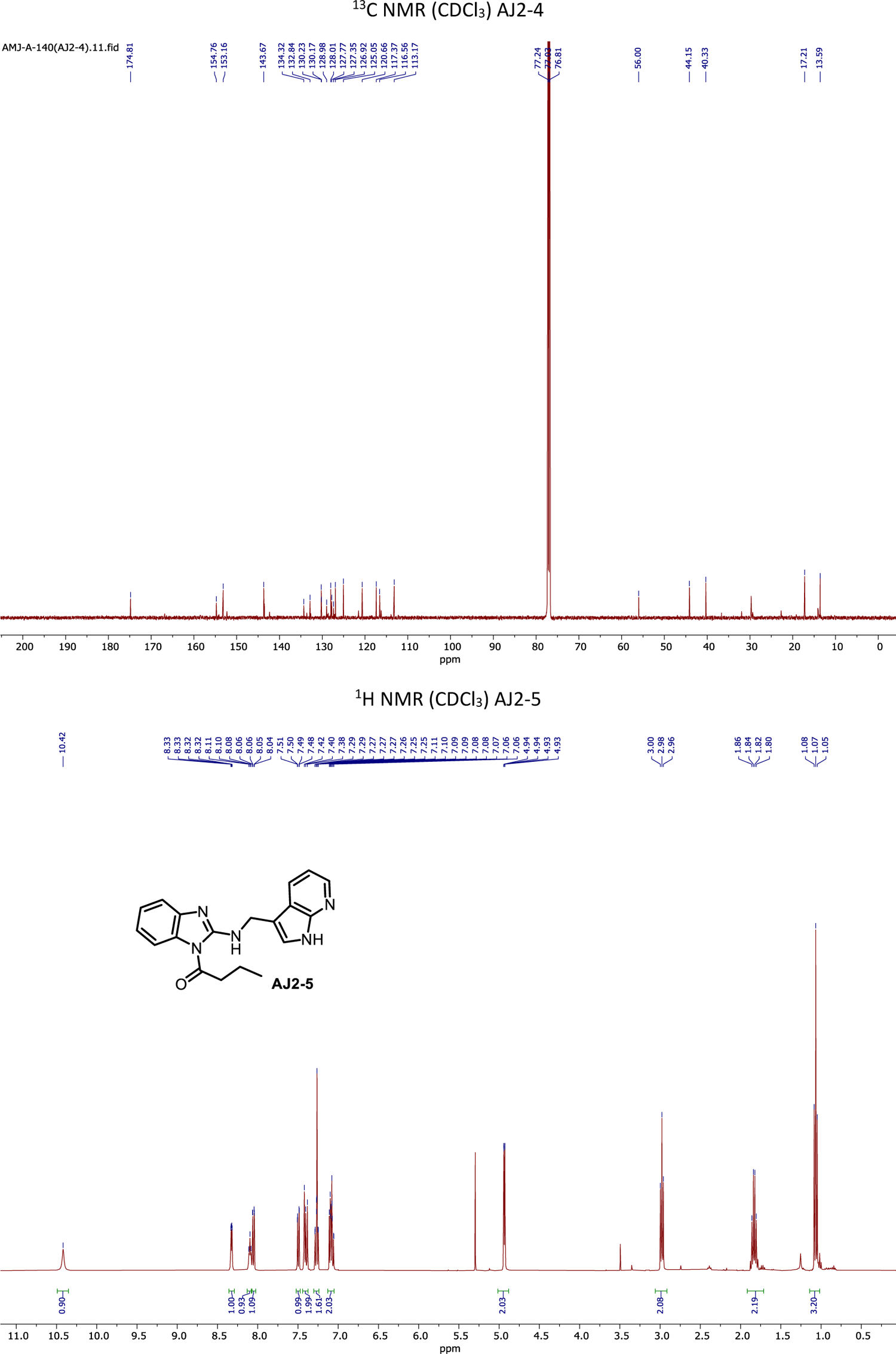
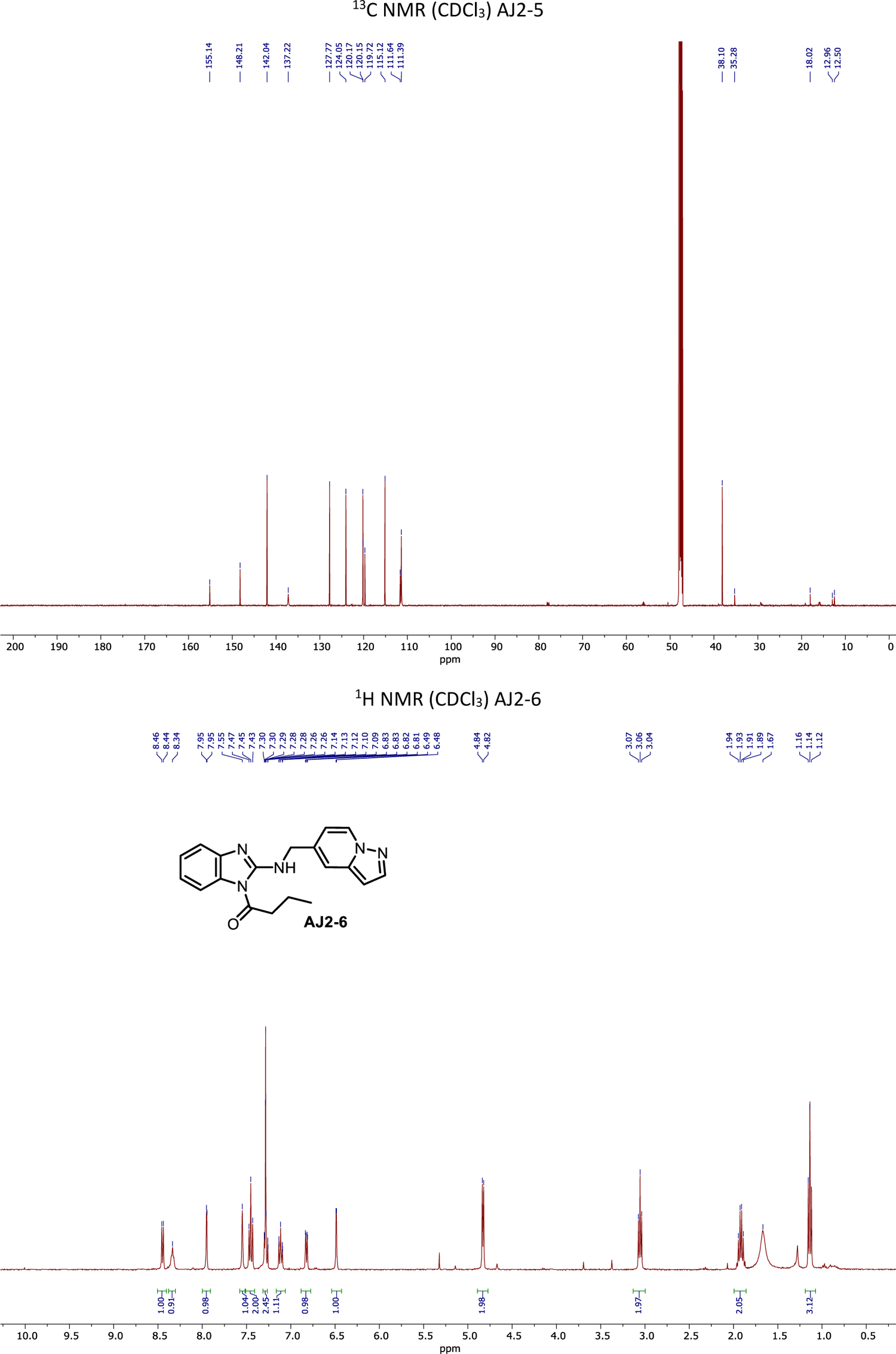
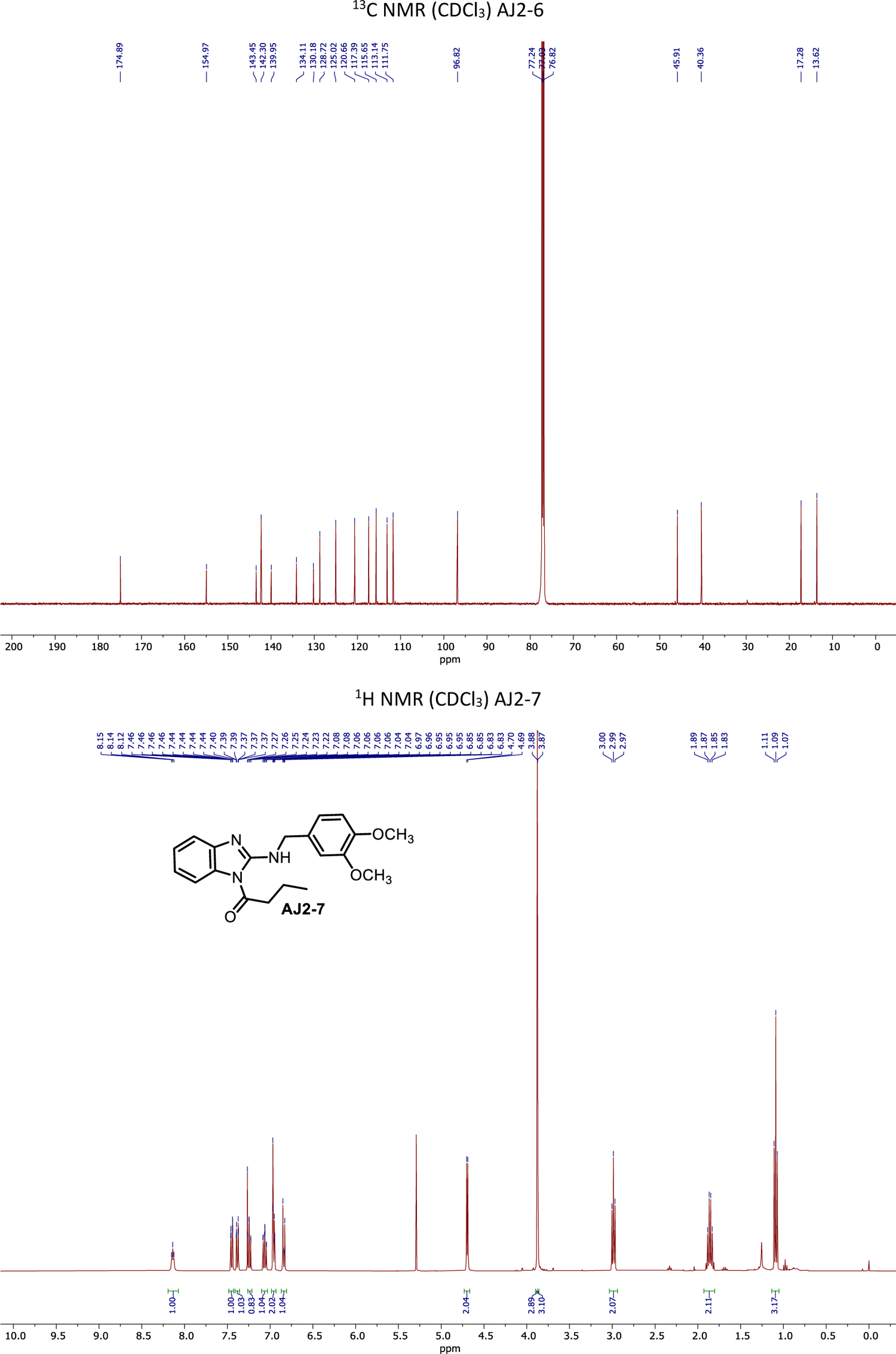
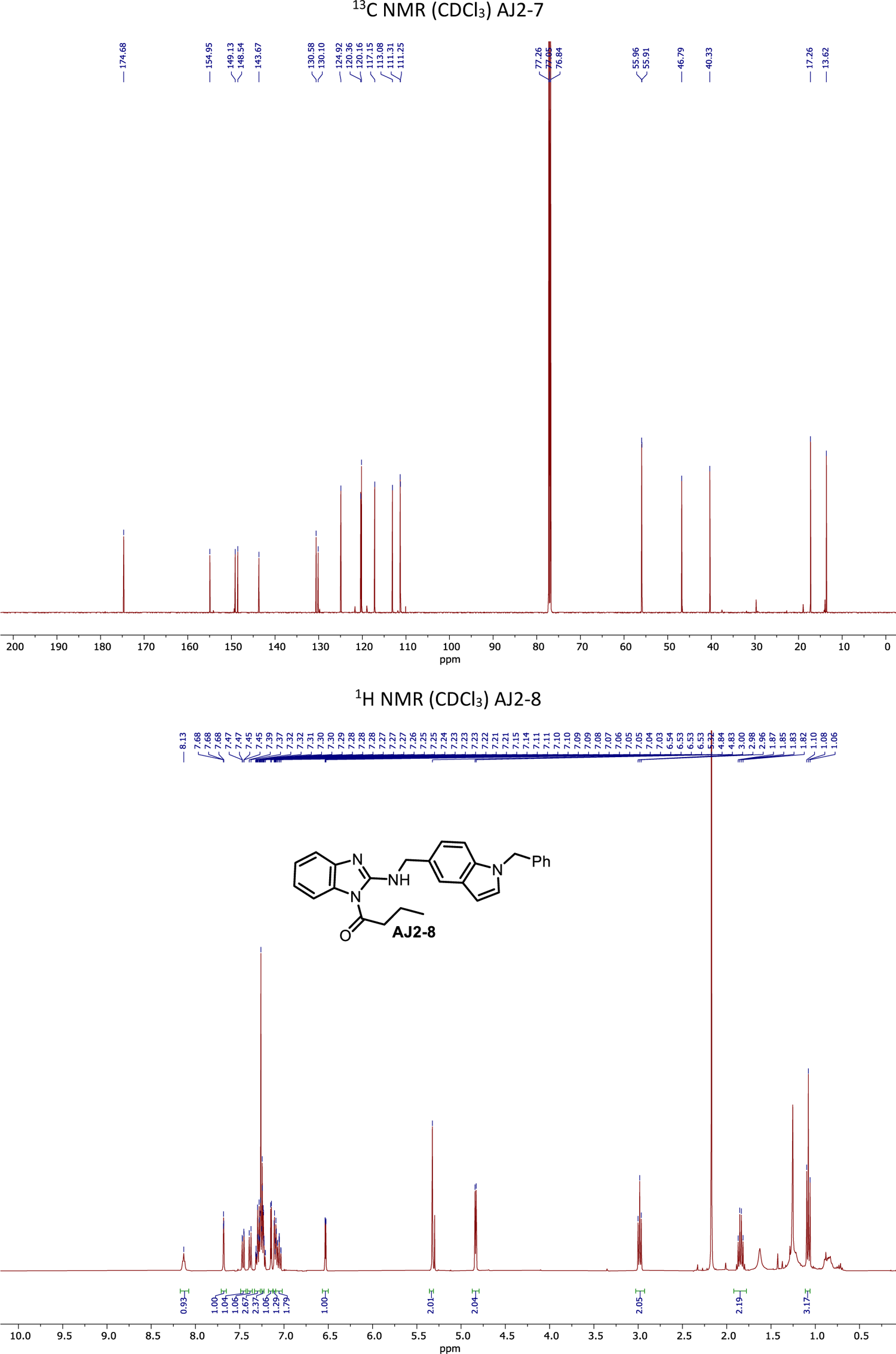
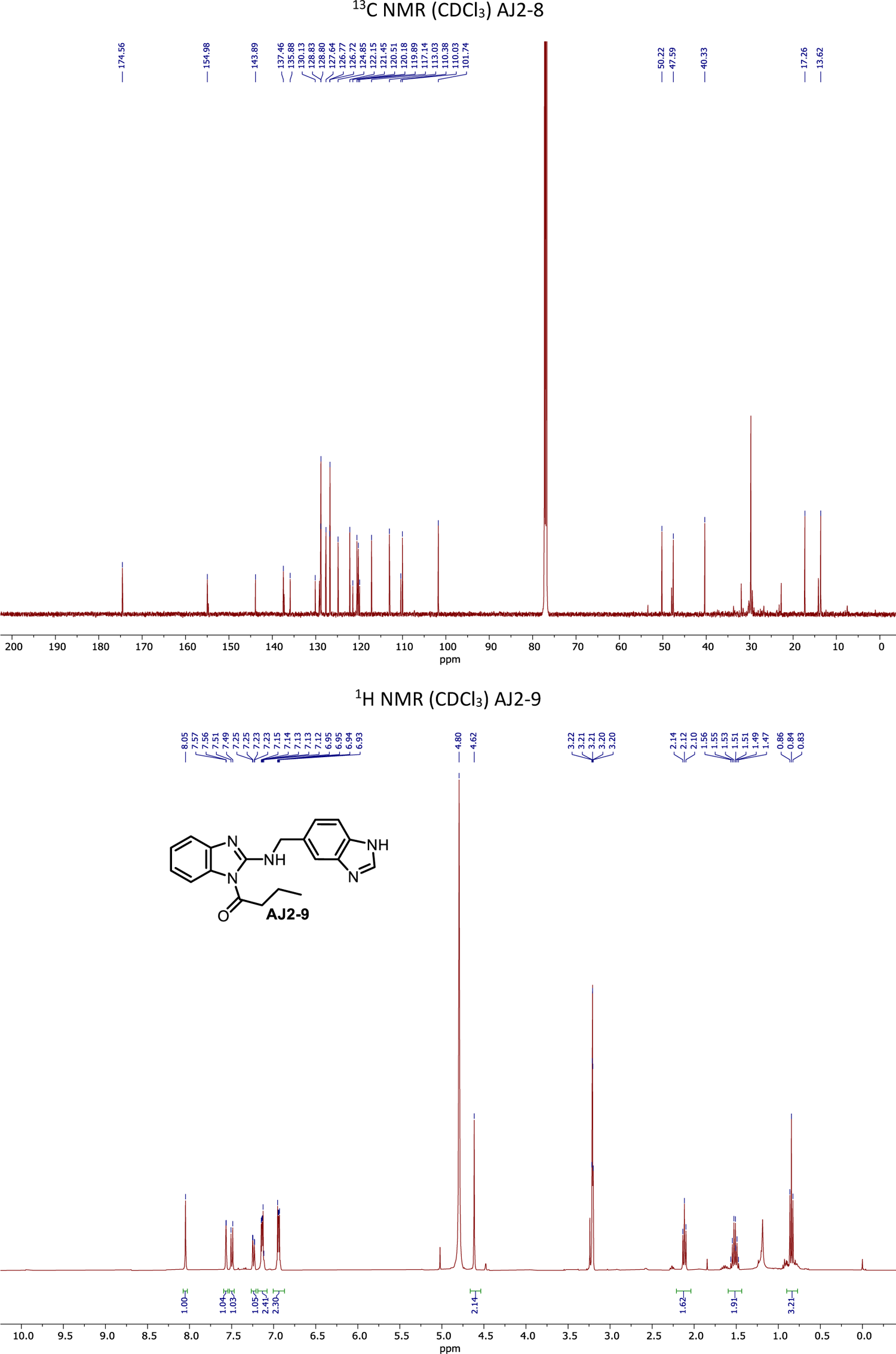
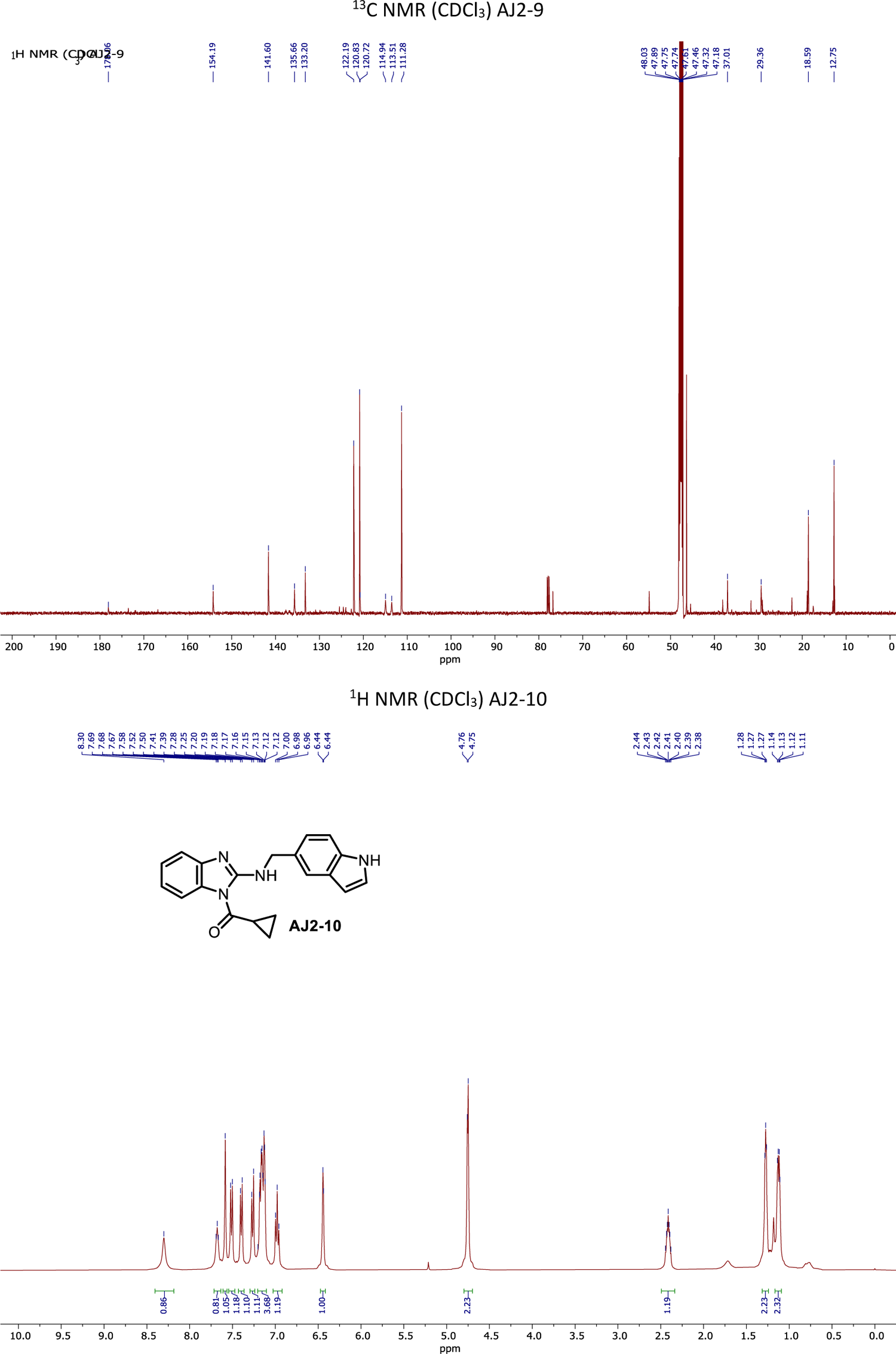
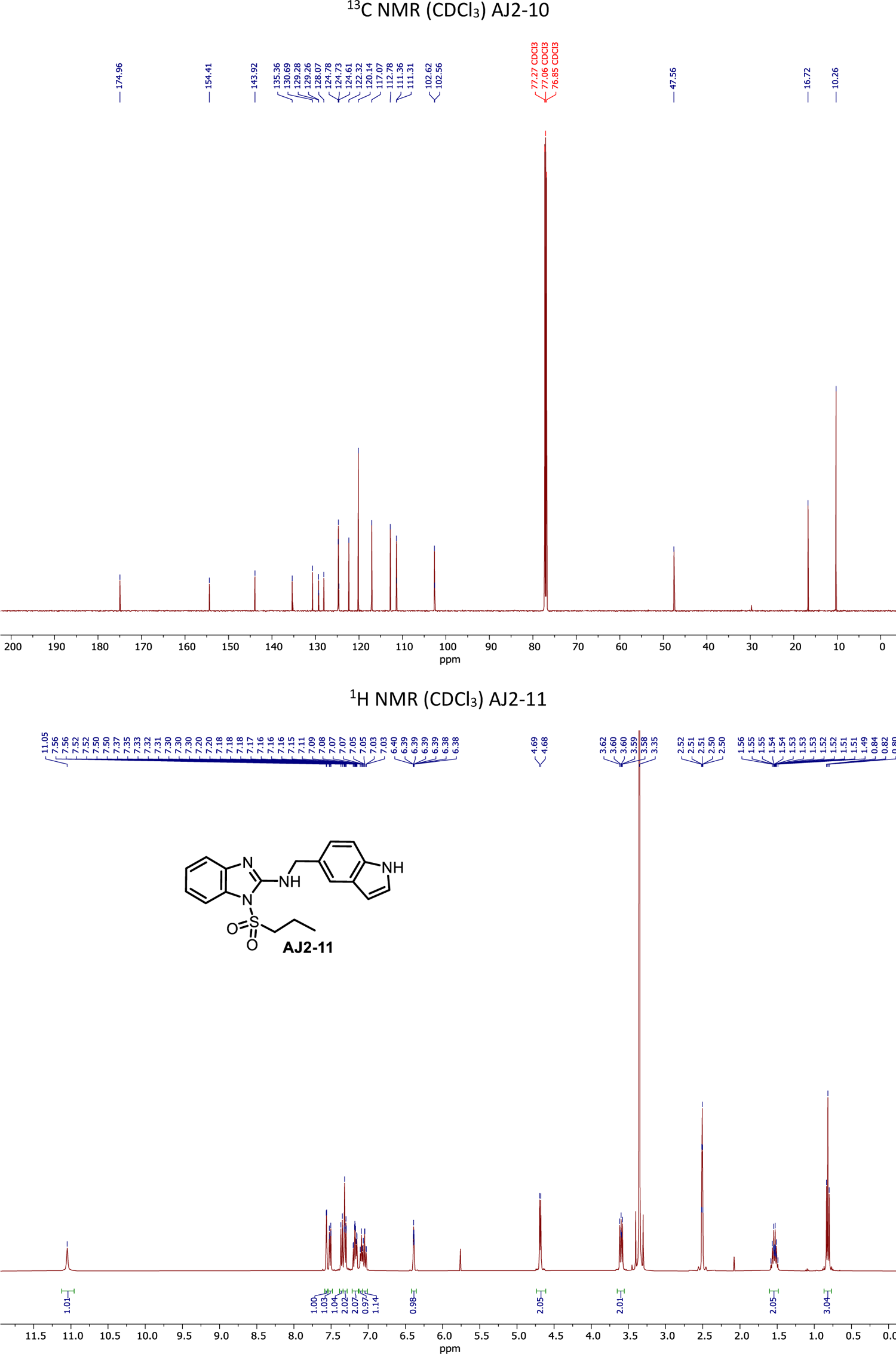
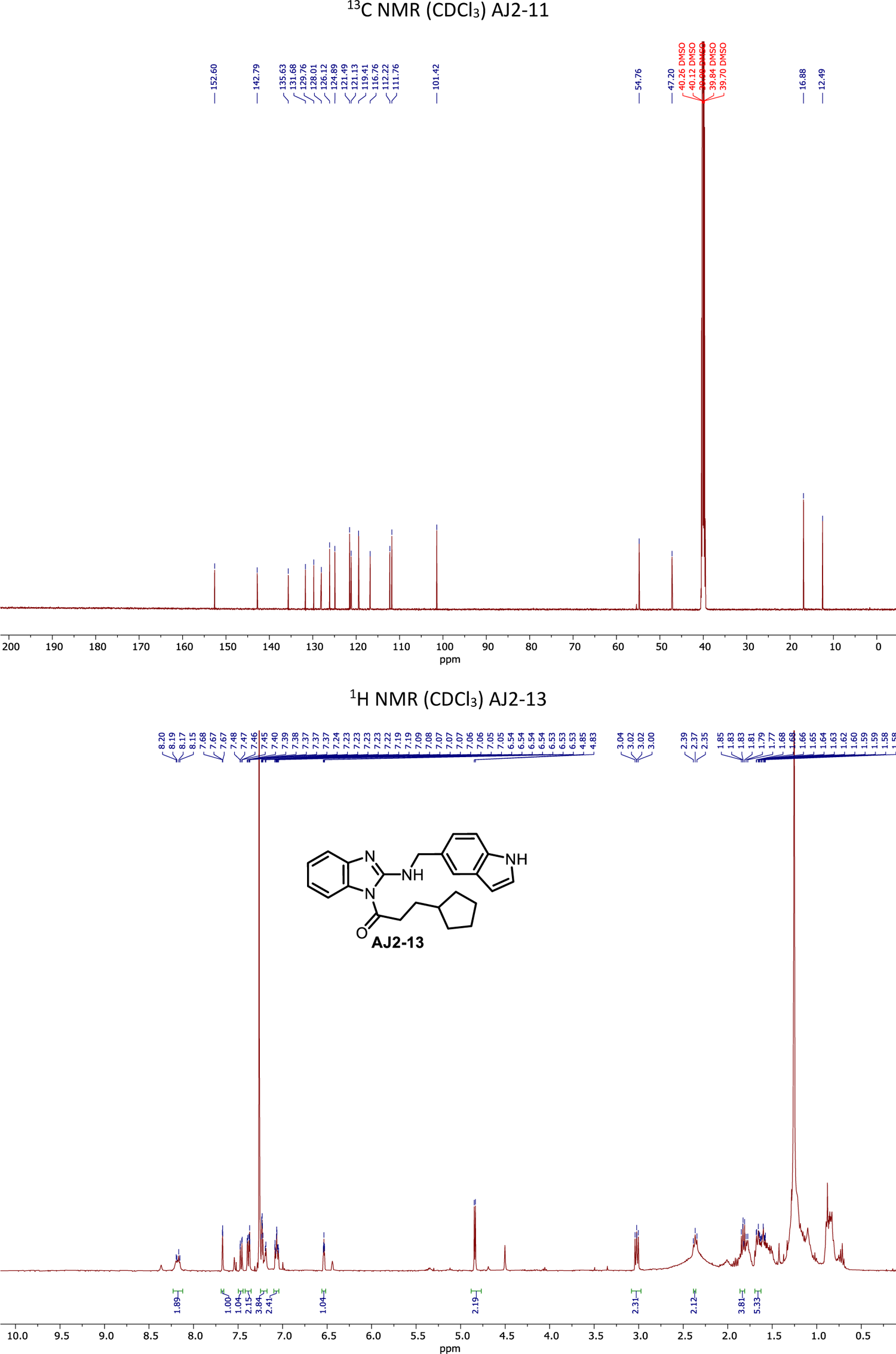
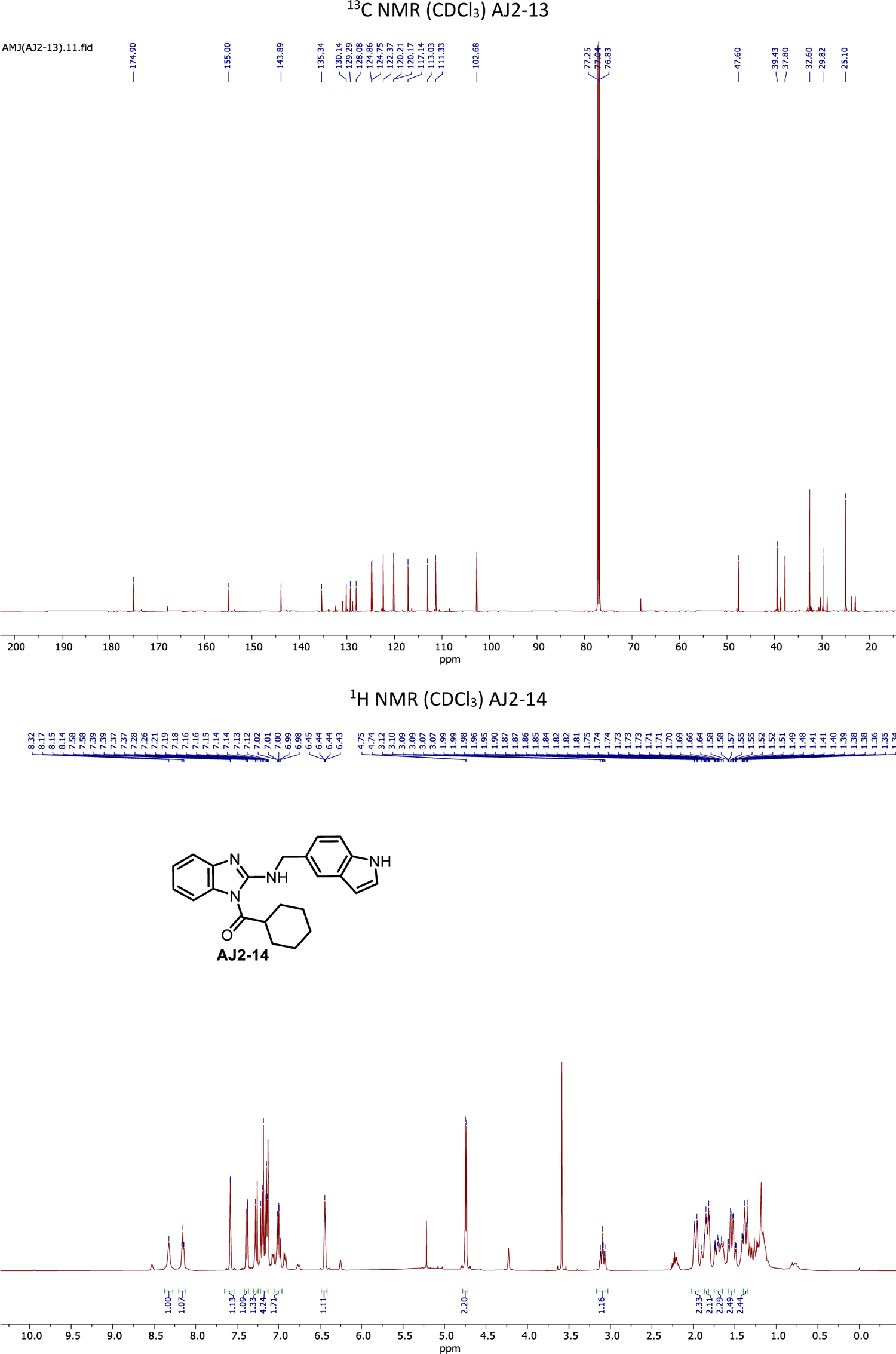
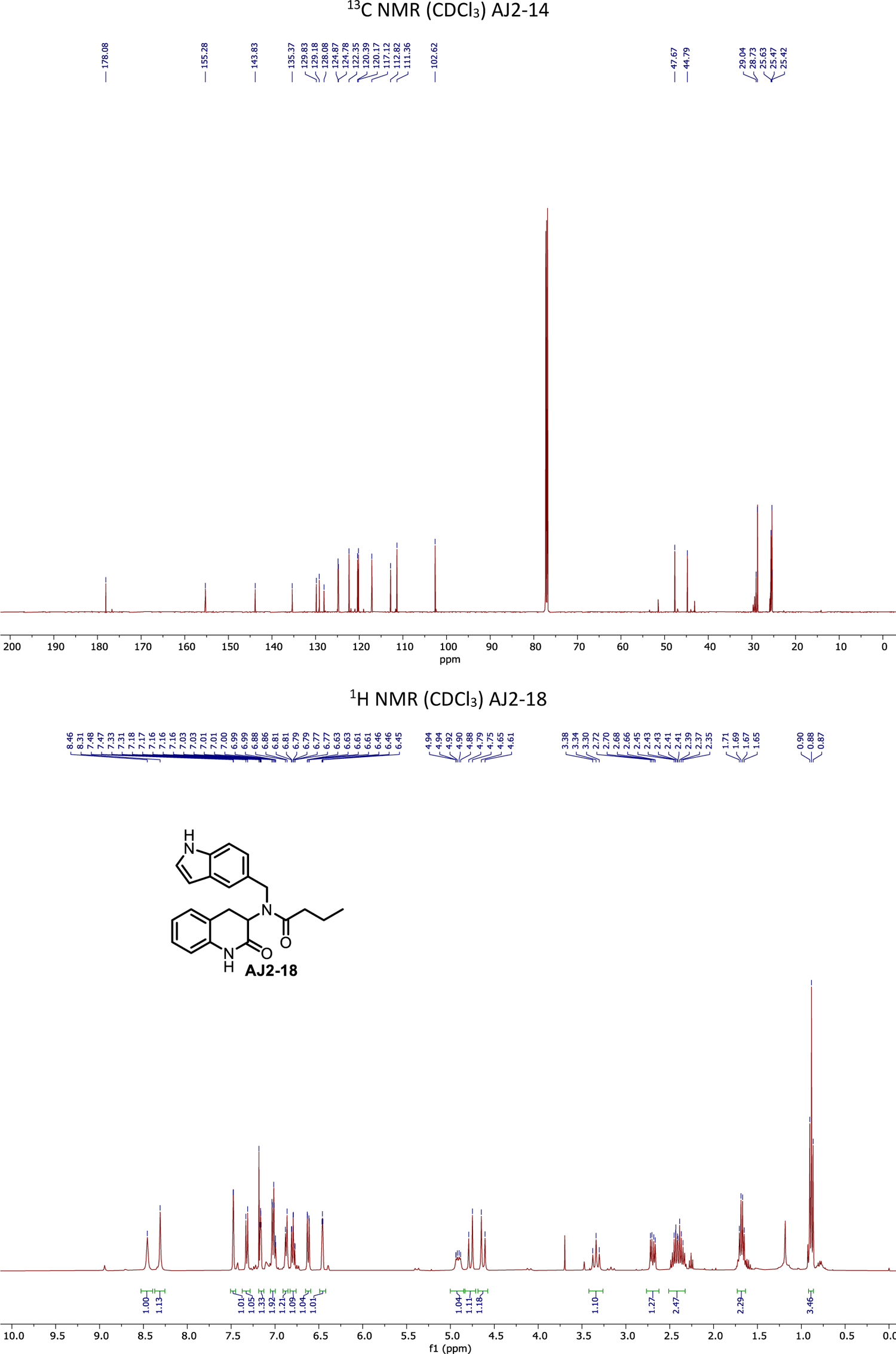
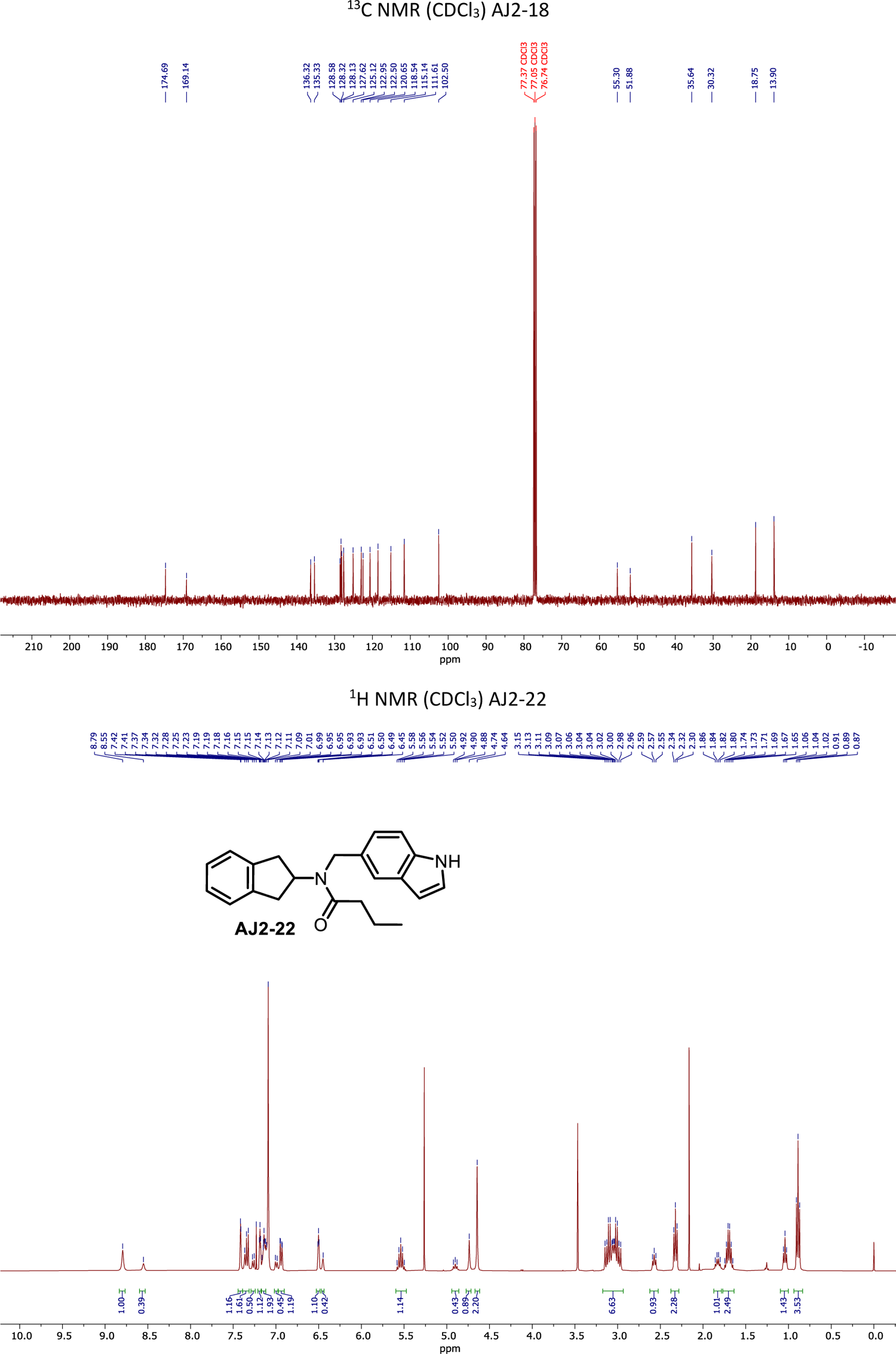
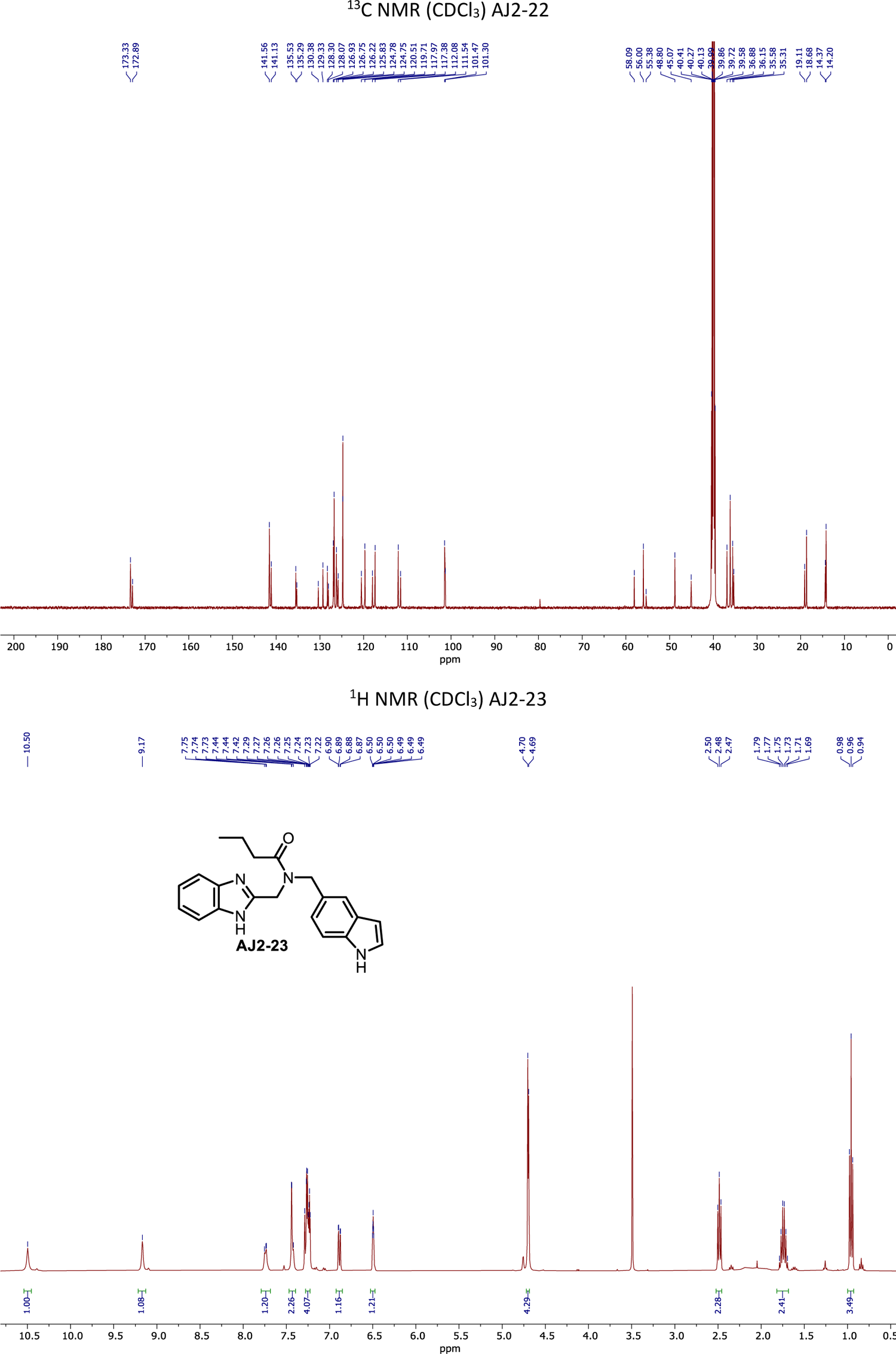
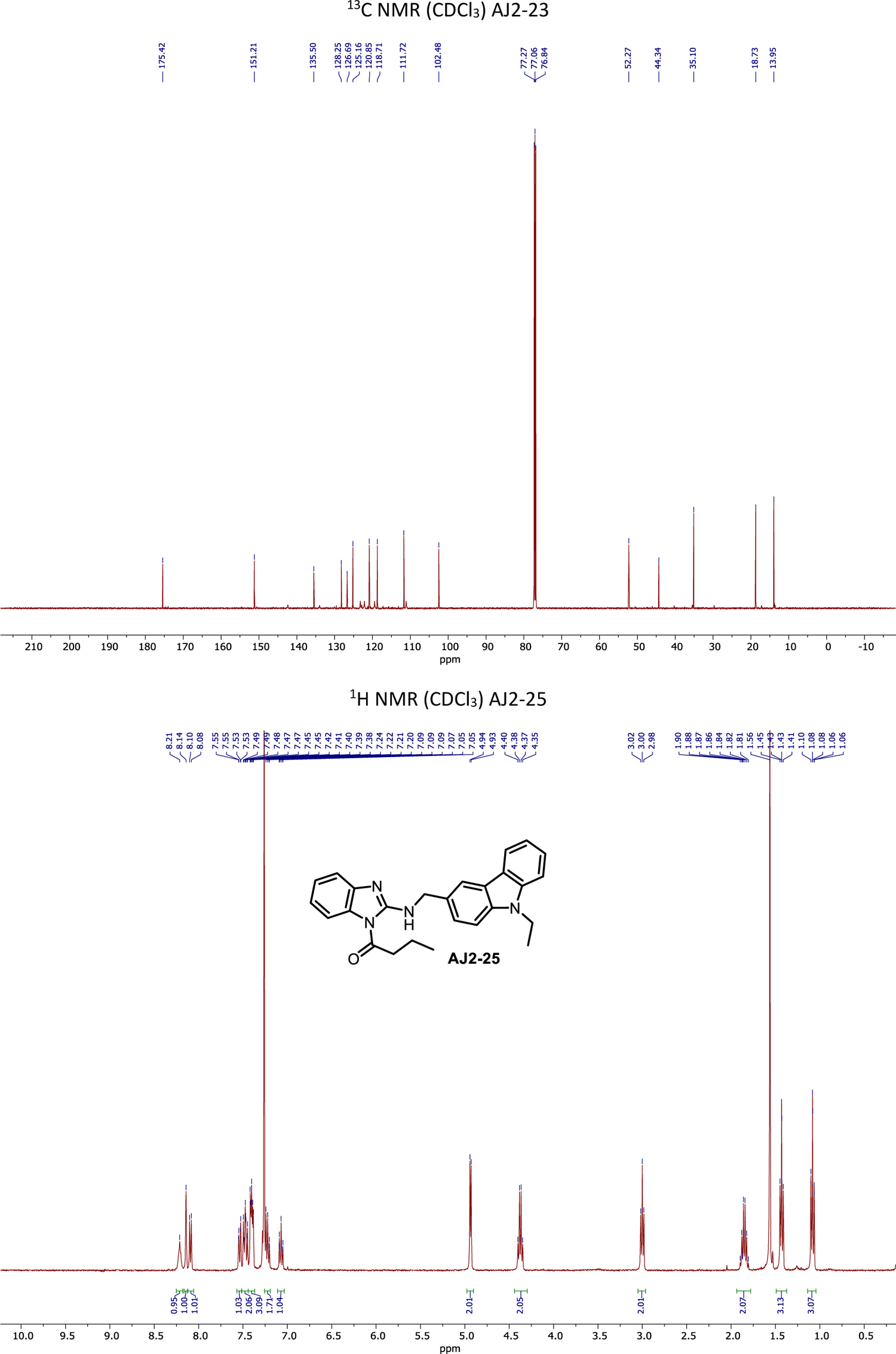
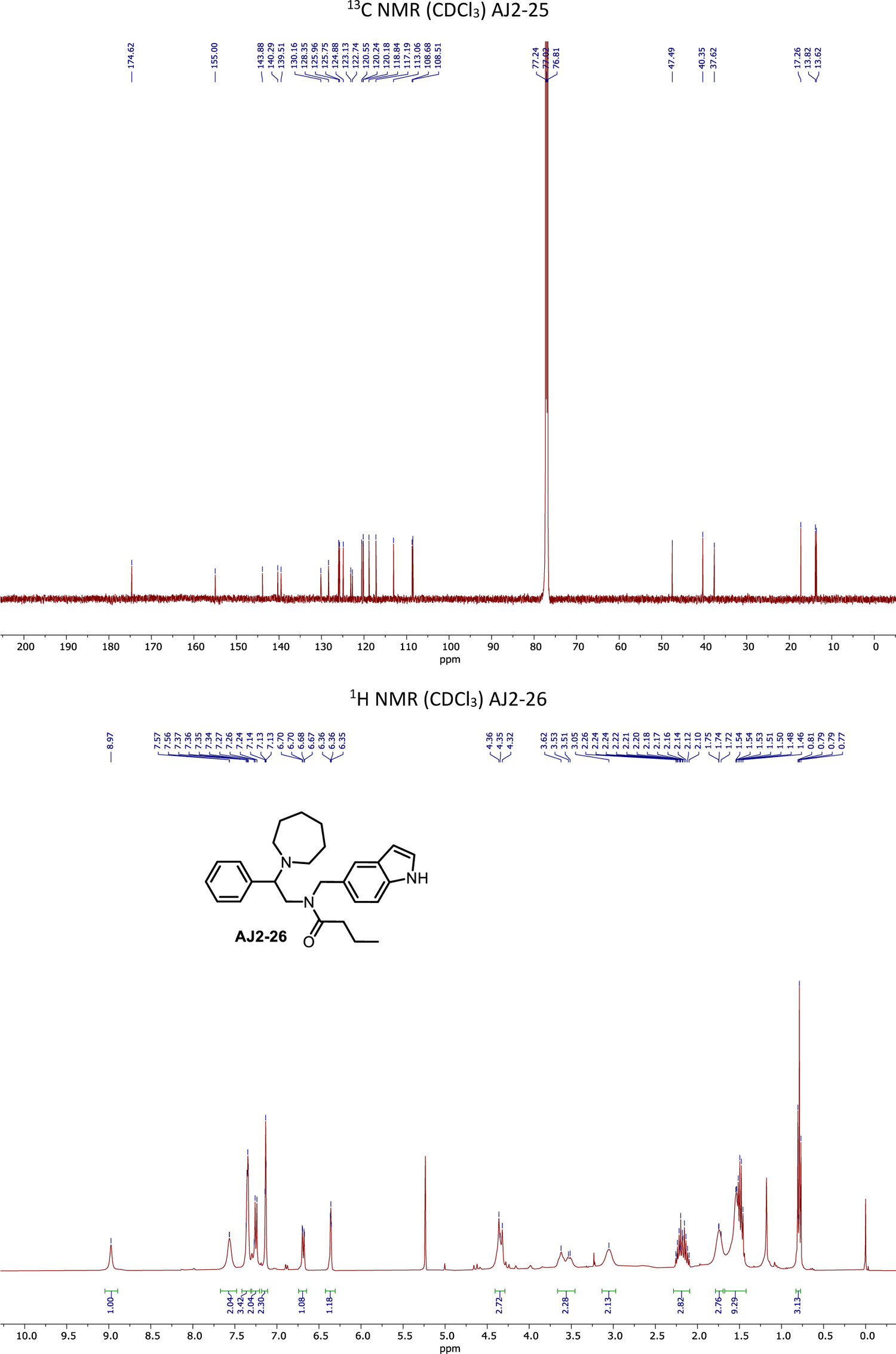
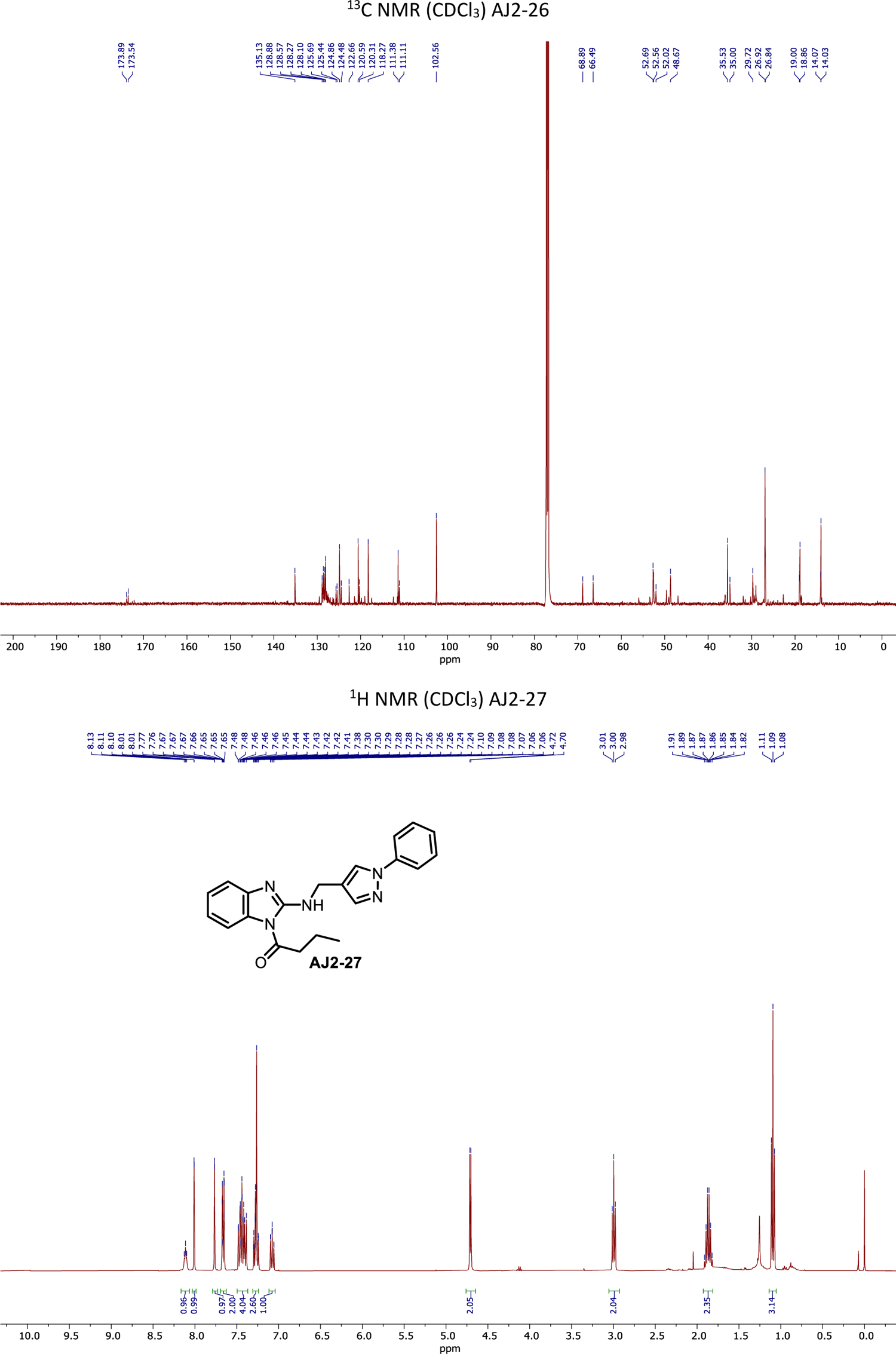
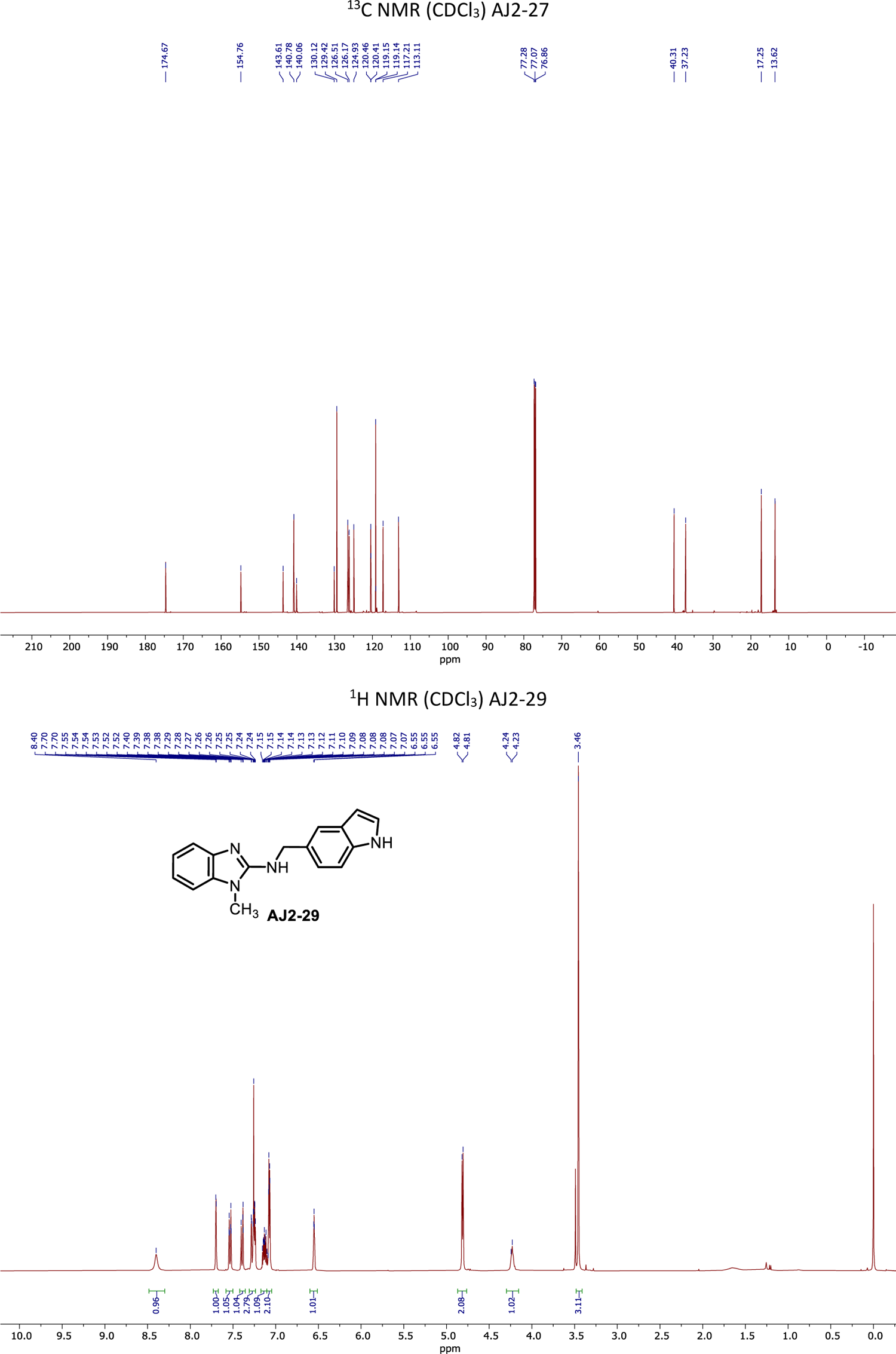
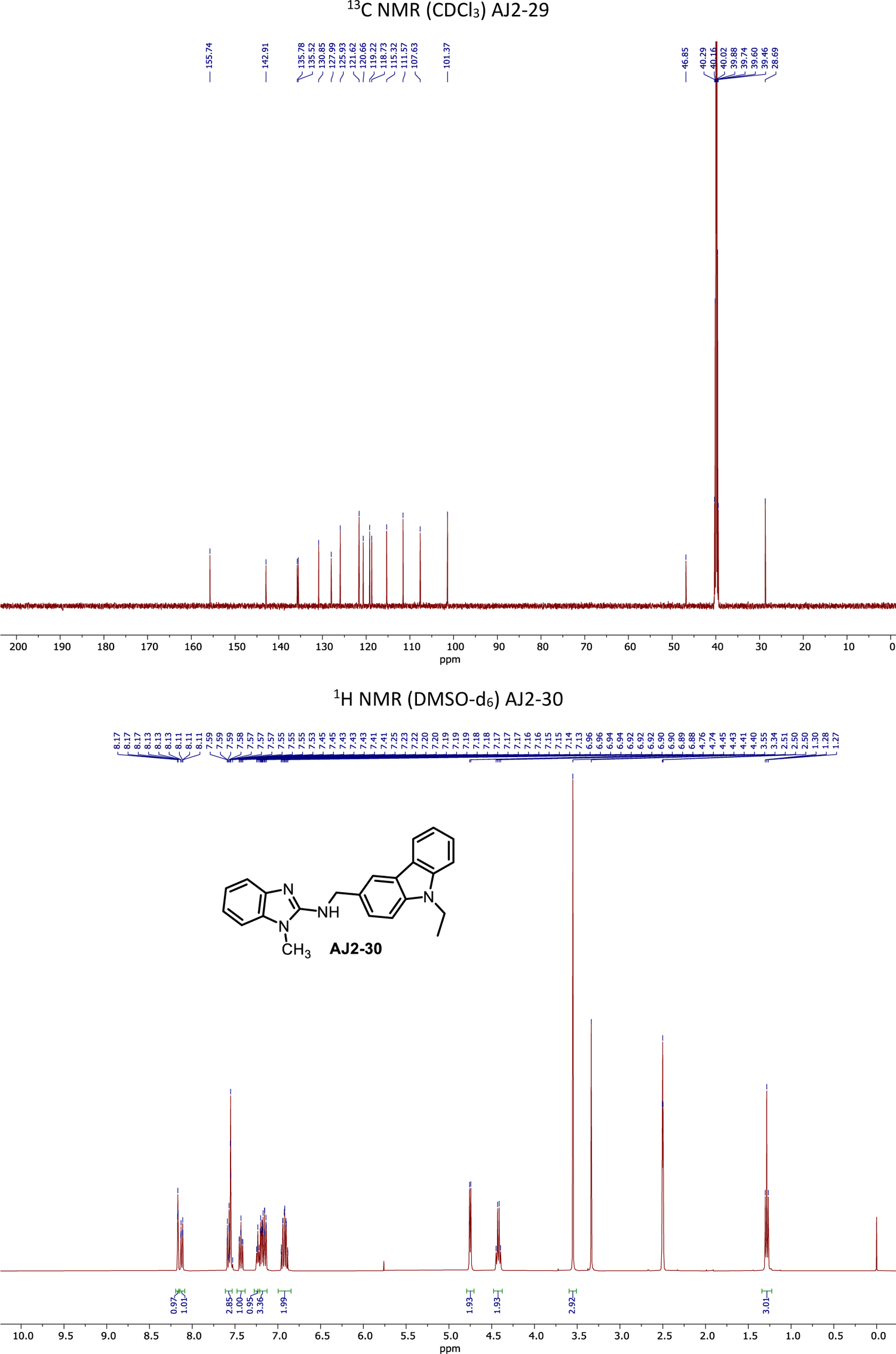
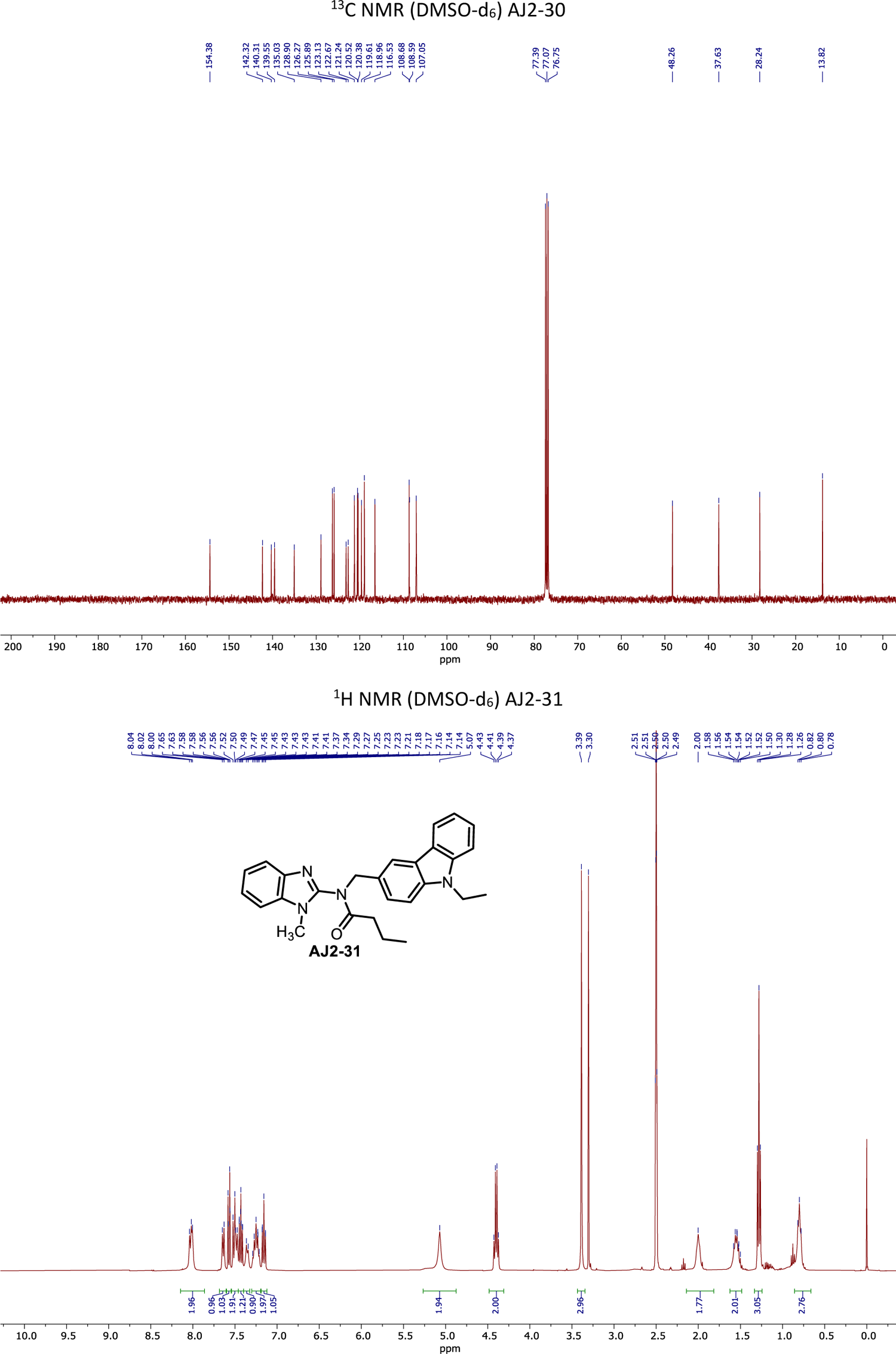
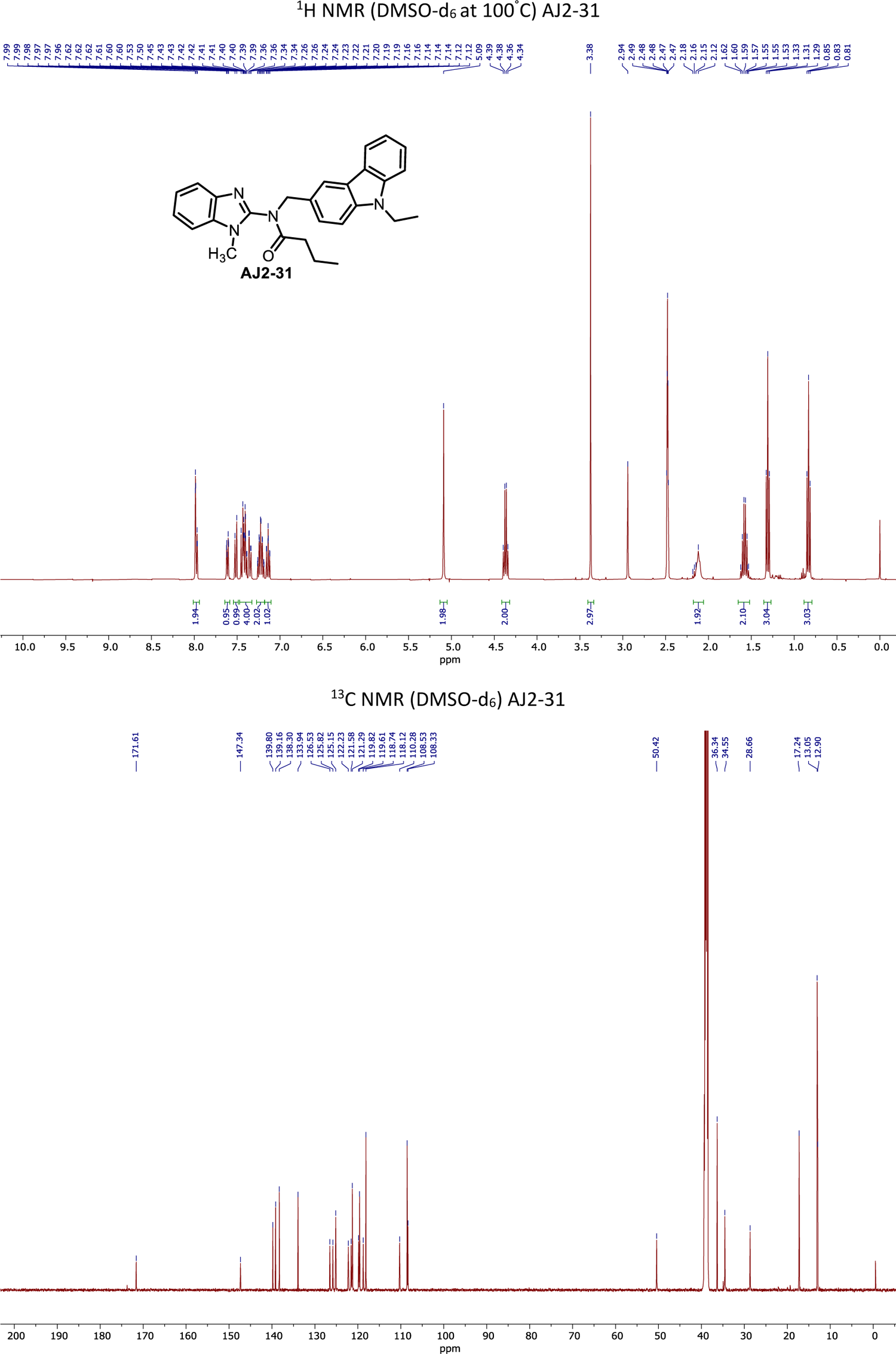
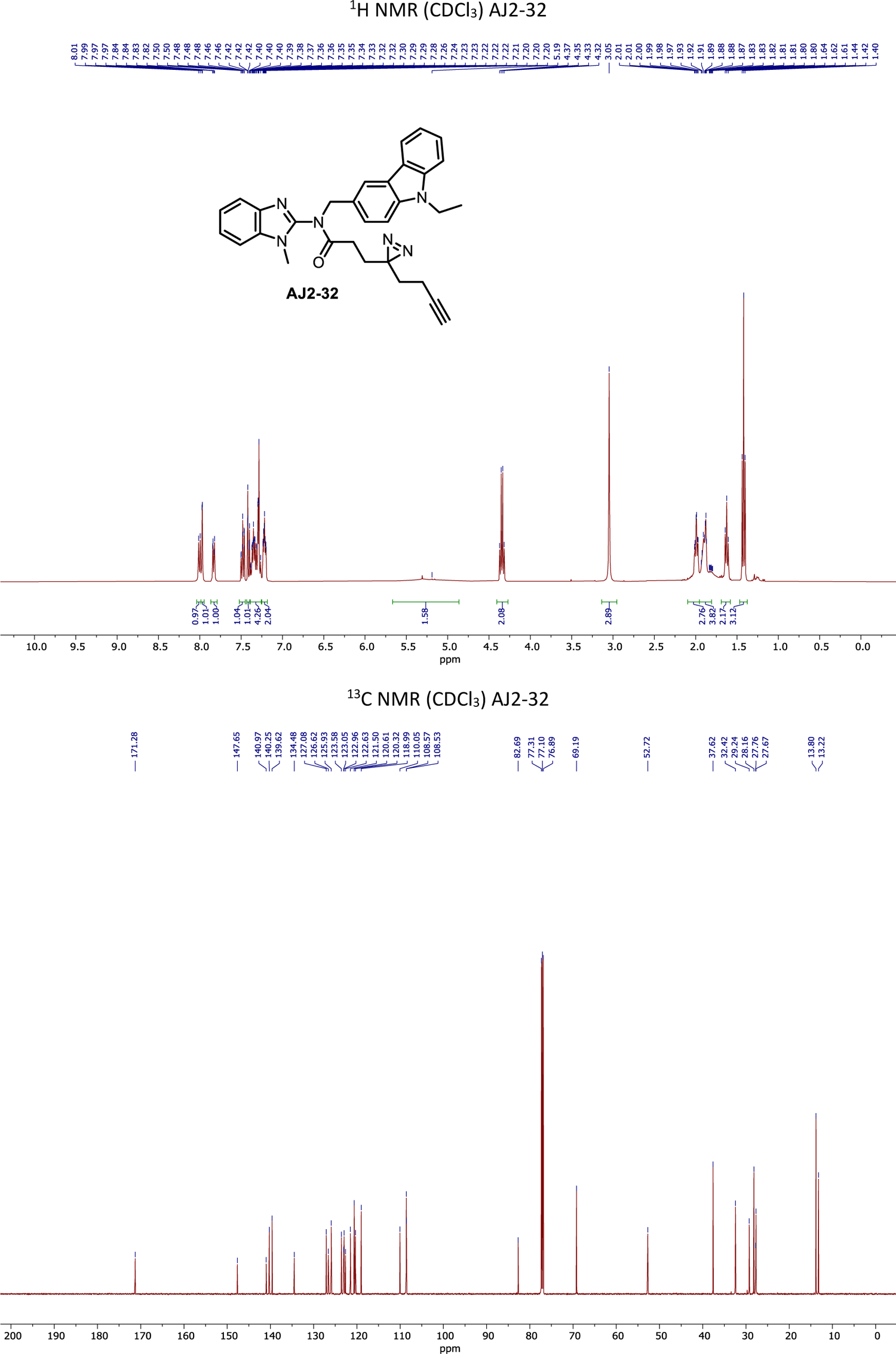
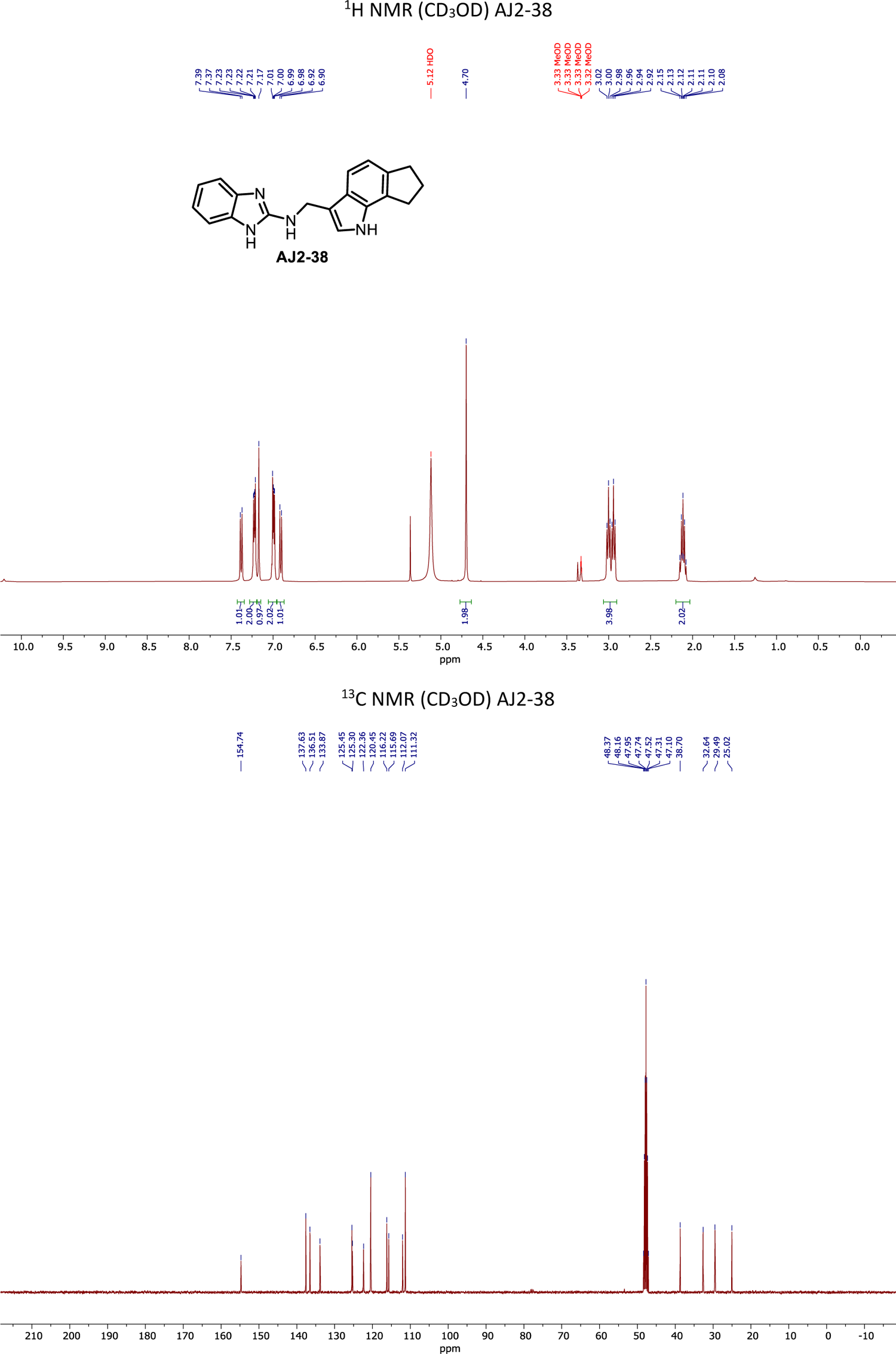
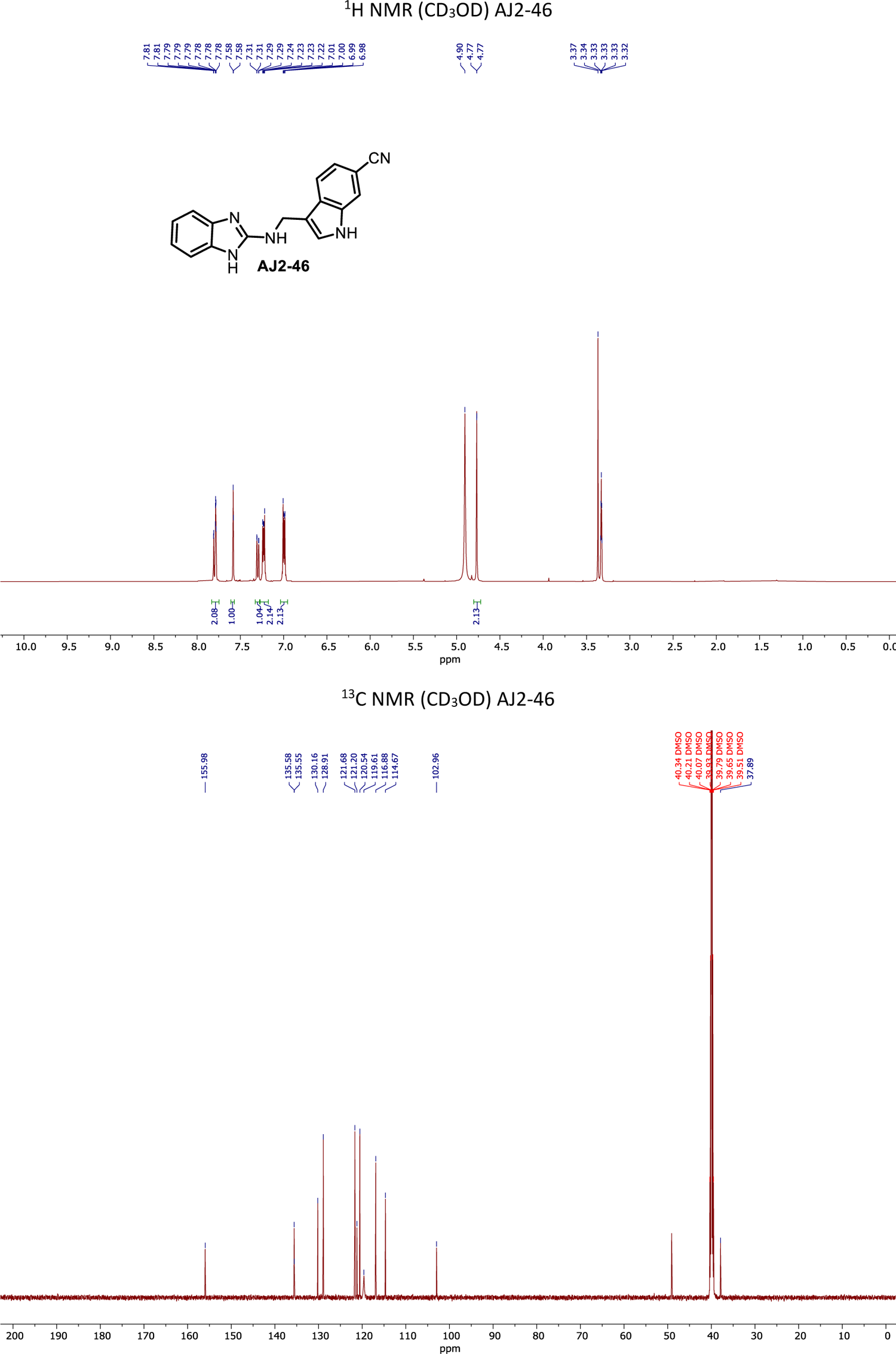
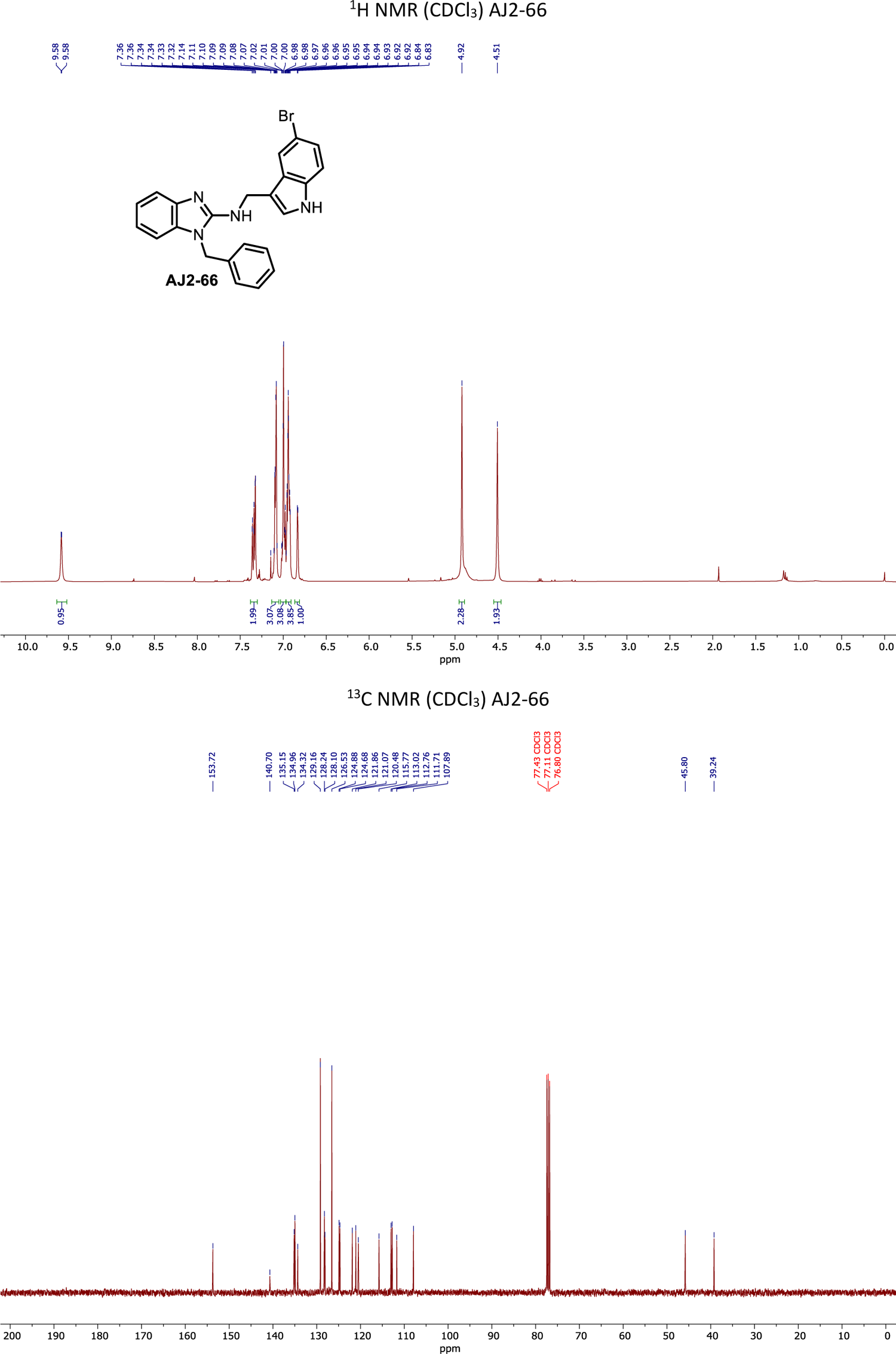
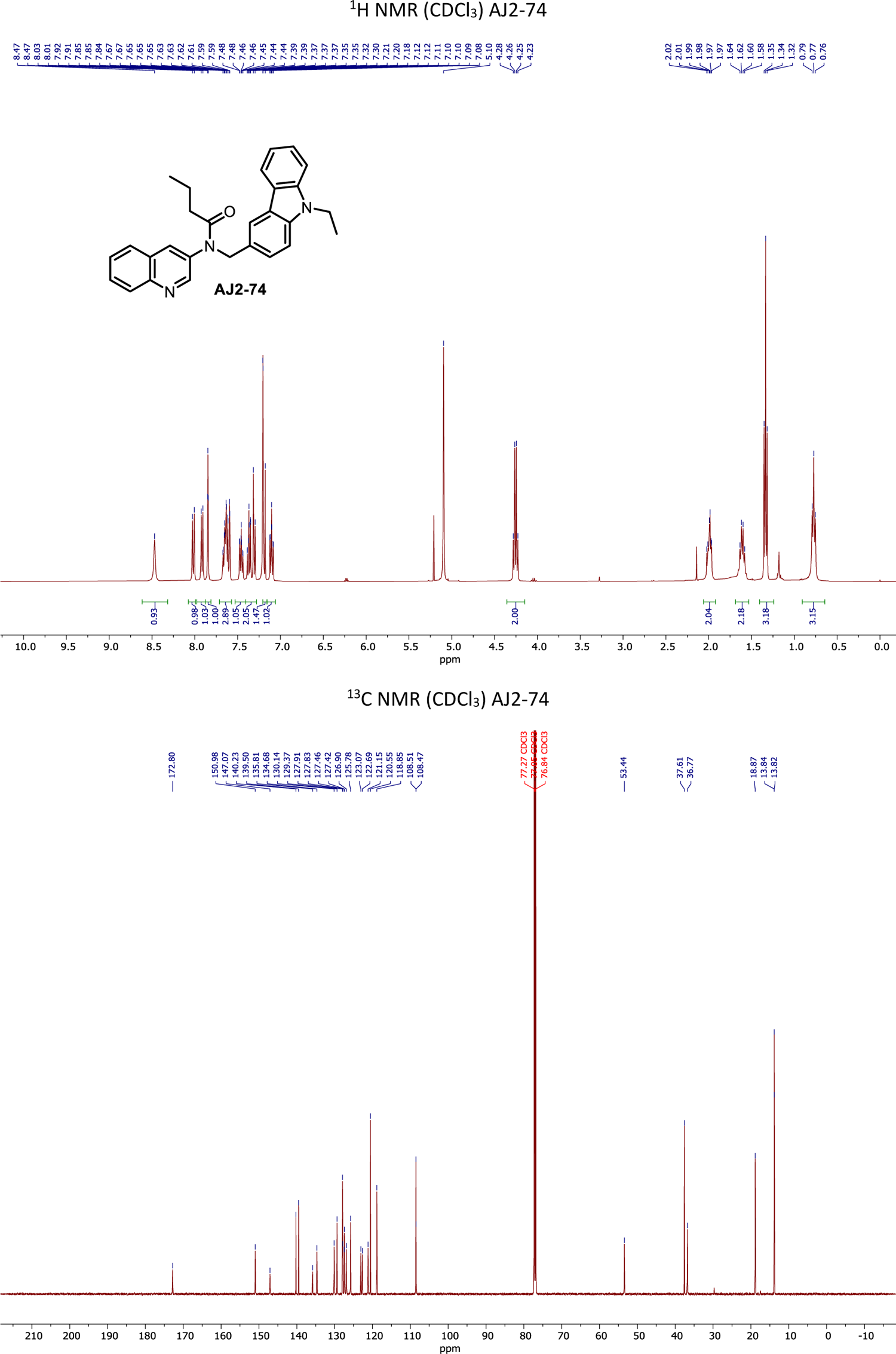
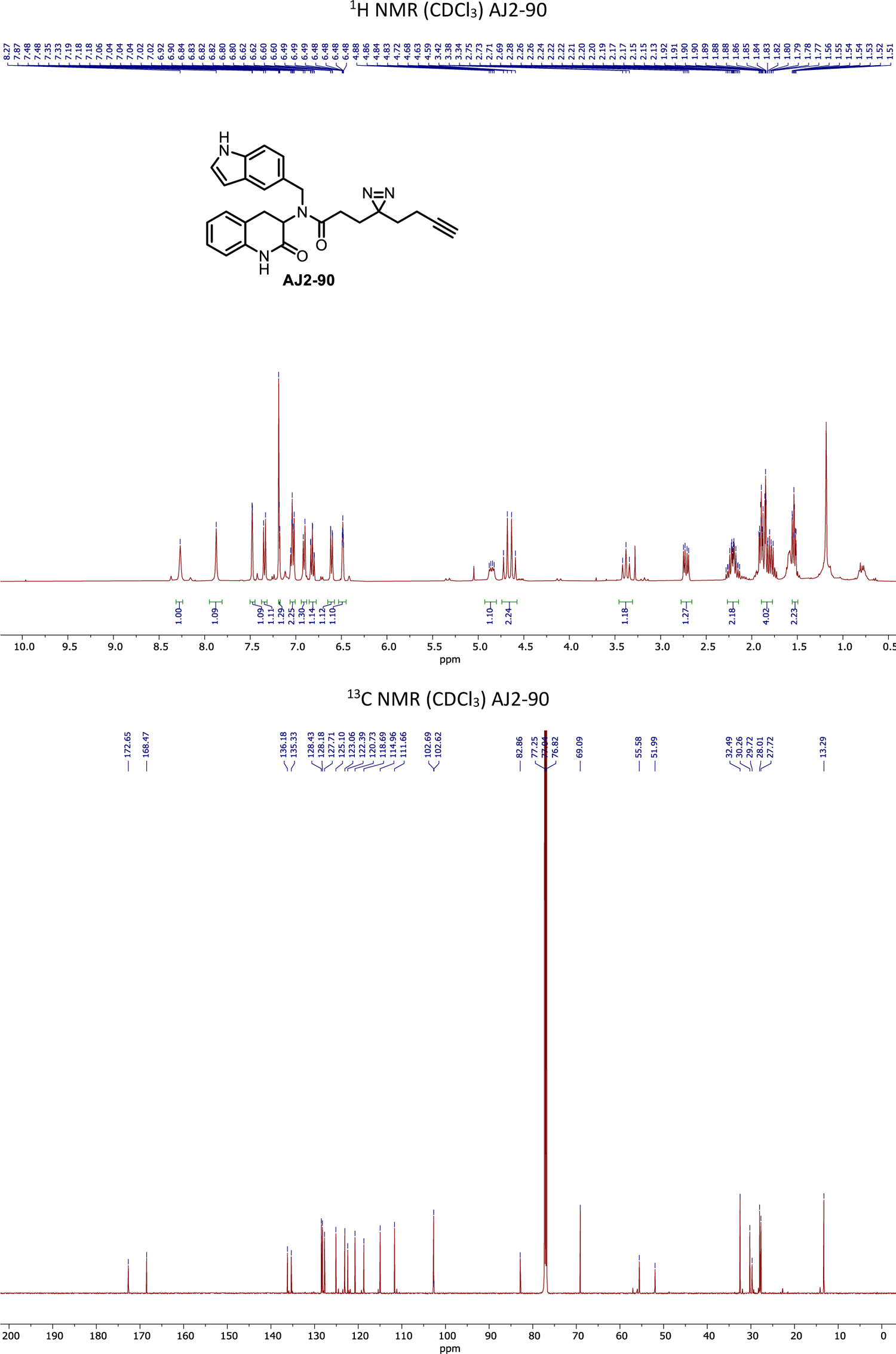

